# Diversification of the restriction–modification system of *Streptococcus pyogenes* through its acquisition of mobile elements

**DOI:** 10.1101/2020.06.30.179317

**Authors:** Atsushi Ota, Yukiko Nishiuchi, Noriko Nakanishi, Yoshio Iijima, Tomotada Iwamoto, Ken Osaki, Yoshitoshi Ogura, Atsushi Toyoda, Yutaka Suzuki, Tetsuya Hayashi, Hiroki Ohge, Hirotada Mori, Manabu Ato, Fumito Maruyama

## Abstract

Restriction–modification (RM) systems are typically regarded as “primitive immune systems” in bacteria. The roles of methylation in gene regulation, segregation, and mismatch repair are increasingly recognized. To analyze methyltransferase (MTase) diversity in *Streptococcus pyogenes*, we compared the RM system distribution in eight new complete genome sequences obtained here and in the database-deposited complete genome sequences of 51 strains. The MTase gene distribution showed that type I MTases often change DNA sequence specificity via switching target recognition domains between strains. The type II MTases in the included strains fell into two groups: a prophage-dominant one and a CRISPR-dominant one. Some highly variable type II MTases were found in the prophage region, suggesting that MTases acquired from phage DNA can generate methylome diversity. Additionally, to investigate the possible contribution of DNA methylation to phenotype, we compared the methylomes and transcriptomes from the four most closely related strains, the results of which suggest that phage-derived methylases possibly regulate the methylome, and, hence, regulate expression levels in *S*. *pyogenes*. Our findings will benefit further experimental work on the relationship between virulence genes and pathogenicity in *S*. *pyogenes*.

## INTRODUCTION

In bacteria, DNA methylation is associated with restriction–modification (RM) systems. RM systems usually encode restriction endonuclease (REase) and DNA methyltransferase (MTase). The REase recognizes and cleaves foreign DNA sequences at specific sites, while MTase activity distinguishes “self” from “non-self” DNA by transferring a methyl from S-adenosyl-methionine to the same specific DNA sequence within the host genome (1, 2). The methylated bases found in bacteria are N6-methyl-adenine (6mA), C5-methyl-cytosine (5mC), and N4-methyl-cytosine (4mC) (3-5).

RM systems can be classified into four types (I, II, III, and IV) based on their molecular structure, cofactor requirements, mechanism of action, target sequence, and cleavage position (2, 4, 5). The most abundant type of RM system is type II (69.5% of all 26,582 RM systems in 5,568 analyzed bacterial genomes) (6). In addition to having an MTase and REase, some type II RM systems also contain various subclasses of enzymes (e.g., IIB, IIG, IIL, and some IIH) that possess both methylation and endonuclease activity (2). Type I RM systems are the second-most-abundant RM systems in prokaryotic genomes (17.4% of all 26,582 RM systems in 5,568 analyzed bacterial genomes) (6, 7). Typical multi-subunit-complex type I RM systems contain an REase, MTase, and a specificity domain. These complexes either fully methylate the recognition sites occurring on only one DNA strand or cleave the DNA several bases upstream or downstream of recognition sites that are unmethylated on both strands (2). Type III RM systems are comprised of a Mod (MTase) protein, which recognizes asymmetric target sequences, and an REase protein. There are relatively few type III MTases in prokaryotic genomes (6.8% of all 26,582 RM systems in 5,568 analyzed bacterial genomes) (7). Finally, type IV RM systems lack methyltransferase activity and consist of only one or two DNA methylation-specific REases.

The roles of the RM system besides its function in the prokaryotic immune system have become more apparent in recent years. The type I specificity subunits of some bacterial species have been found to become switched with one another, thereby producing heterogeneity in gene expression (8-12). This “phase variation” can also contribute to control bacterial virulence, immune evasion, and niche adaptation (13). “Solitary methyltransferases” can also mediate DNA methylation independently of an active cognate REase. Solitary MTase, a type II RM system, has no REase but is nevertheless involved in the global gene regulation of several bacterial species (14, 15). Two examples of solitary bacterial MTases are DNA adenosine methyltransferase (Dam) and cell cycle-regulated methyltransferase (CcrM). Dam methylation of GATC sites in *Escherichia coli* has been shown to regulate the timing of DNA replication, whereas in *Caulobacter crescentus*, CcrM regulates the cell cycle (16). These studies have extended our understanding, such that we now view DNA methylation as an epigenetic signaling mechanism used by bacteria to help with their ongoing adaptation.

*Streptococcus pyogenes* causes uncomplicated pharyngitis, scarlet fever, rheumatic fever, and streptococcal toxic shock syndrome (STSS). Invasive strains of *S*. *pyogenes* are often reported as “flesh-eating” bacteria because infections with these pathogens can progress rapidly and dramatically after illness onset, with an approximate mortality rate of 40% (18). This is why many researchers study *S*. *pyogenes* despite it not being a model organism. The incidence of *S*. *pyogenes* infection has risen since 2011 (17-19), and elucidating its invasion mechanism has become a pressing issue. It is thought that the virulence factors encoded by phages are involved in bacterial virulence to some extent (20). Comparative genome analyses have suggested the pathogenic diversity of *S*. *pyogenes* might result from single-nucleotide polymorphisms (SNPs) in genes such as those encoding two-component systems (TCSs) (21). However, the presence or absence of some specific genes and SNPs in TCSs were not found to contribute to the pathogenesis widely (22). Thus, other factors, like epigenetic function, should be investigated for their potential to define the pathogenicity of *S*. *pyogenes*.

To analyze the MTase gene diversity in *S*. *pyogenes*, the eight *S*. *pyogenes* strains we sequenced herein were compared with 51 complete genome database-deposited sequences to produce an updated view of the phylogenetic diversity of this species. The results of our methylome and RNA-Seq analyses prompt us to suggest that phage-derived methylases regulate the methylome, and hence, regulate expression levels in *S*. *pyogenes*. Our findings contribute to the current knowledge base about the effects of epigenetic modification on the pathological diversification of *S*. *pyogenes*.

## MATERIALS AND METHODS

### Strains and DNA preparation

For genome sequence determination, we obtained four *S*. *pyogenes* strains from the American Type Culture Collection (ATCC) (ATCC14918, ATCC14919, MGAS9429, and MGAS2096) and four strains from the Department of Infectious Diseases, Kobe Institute of Health (KIH05-6, KIH06-45, KIH06-201, and KIH08-6) (Table 1). ATCC14918 and ATCC14919 were isolated from a human pharynx in 1986. ATCC14919 is an indicator strain lysed by a phage of ATCC14918. MGAS9429 was isolated from an animal swab (23). MGAS2096 was isolated from a patient with acute post-streptococcal glomerulonephritis in 1993 (23). The *emm* type of all four of these strains is *emm*12. KIH05-6, KIH06-45, KIH06-201, and KIH08-45 were isolated in 2005–2008 from patients with pharyngitis in Japan, and their serotypes are *emm*3, *emm*6, *emm*6, and *emm*156, respectively. These strains were cultured in Todd–Hewitt broth (BD Diagnostic Systems; Sparks, MD, USA) supplemented with 0.2% yeast extract, as previously described (24).

**Table 1.** The complete genome sequences determined in this study.

| Strain | *emm* type | Isolation source | Accession # | # of<br>contig | Total read length | Genome size | Genome<br>coverage | n50 | GC content<br>(%) | gene | CDS | tRNA | rRNA |
| --- | --- | --- | --- | --- | --- | --- | --- | --- | --- | --- | --- | --- | --- |
| ATCC14918 | 229 | Pharynx | DRX060373 | 1 | 2,259,100,157 | 1,796,820 | 1,257.3 | 1,796,820 | 38.56 | 1,768 | 1,682 | 67 | 18 |
| ATCC14919 | 229 | Pharynx | DRX060374 | 1 | 2,411,644,106 | 1,757,745 | 1,372.0 | 1,757,745 | 38.56 | 1,709 | 1,623 | 67 | 18 |
| MGAS9429_2 | 229 | Pharyngeal swab - animal | DRX060376 | 1 | 2,010,397,268 | 1,836,296 | 1,094.8 | 1,836,296 | 38.55 | 1,824 | 1,738 | 67 | 18 |
| MGAS2096_2 | 229 | Human clinical specimen | DRX060375 | 1 | 1,628,396,143 | 1,860,246 | 875.4 | 1,860,246 | 38.73 | 1,824 | 1,738 | 67 | 18 |
| KIH05-6 | 6 | pharyngitis | DRX173148 | 1 | 311,592,050 | 1,866,843 | 166.9 | 1,866,843 | 38.58 | 1,914 | 1,828 | 67 | 18 |
| KIH06-45 | 3 | pharyngitis | DRX173149 | 1 | 315,098,671 | 1,906,193 | 165.3 | 1,906,193 | 38.62 | 1,979 | 1,880 | 77 | 21 |
| KIH06-201 | 156 | pharyngitis | DRX173150 | 1 | 152,246,458 | 1,982,968 | 76.8 | 1,982,968 | 38.5 | 2,023 | 1,937 | 67 | 18 |
| KIH08-6 | 6 | pharyngitis | DRX173151 | 1 | 610,902,547 | 1,905,820 | 320.5 | 1,905,820 | 38.6 | 1,969 | 1,883 | 67 | 18 |

DNA was extracted from these eight *S*. *pyogenes* isolates, as previously described (24). The extracted DNA concentrations were determined using the Quant-iT PicoGreen dsDNA Assay Kit (Life Technologies Corporation, CA, USA), and DNA quality was checked using a NanoDrop 1000 instrument (Thermo Fisher Scientific, DE, USA). The 51 complete *S*. *pyogenes* genome sequences from the PATRIC genome online database (http://www.patricbrc.org) were obtained on 11 July 2016 (Supplementary Table S1). They were used as references for the comparative genomics analyses.

### SMRT sequencing and sequence annotation

Complete genome sequences were determined for the strains using the PacBio RSII platform (Pacific Biosciences, Menlo Park, CA, USA), a 20-kb insert-sized library, and P4C2 chemistry (Table 1). *De novo* assembly was conducted using Falcon 0.5.0 with the default parameters (25) for strain MGAS9429. Canu ver. 1.2 with the default parameters (26) was used for the other seven strains. Genome annotation was performed by using Prokka ver. 1.11 with the default parameters (27).

### Characterizing RM systems

RM systems were detected using a sequence similarity search (on May 21, 2015) with the RM genes registered in REBASE (http://rebase.neb.com/rebase/rebase.html) (28). Component genes from the RM systems were annotated by BLASTP searching (29) against the protein sequences using the gold-standard library for RM systems obtained from the database as queries. All hits were clustered with Blastclust in the NCBI Blast package (29), and the resulting clusters were merged when they showed over 90% sequence similarity in the alignable regions.

Target recognition domain (TRD) sequences were determined and classified based on the BLASTP results. Sequences lacking similarity in the BLASTP search (e-value: <1e^−5^) against known TRDs in REBASE were classified as new TRD variants. Besides the gold standard, we used information on the methylation patterns of the *S*. *pyogenes* strains available at REBASE PacBio (http://rebase.neb.com/cgi-bin/pacbio; last accessed March 16, 2016) to predict the recognition sequences of the type I and II RM systems.

### Characterizing prophage and CRISPR regions

Prophages in each genome were predicted using PhiSpy ver. 2.3 (30) run with the default parameters, and the prophage sequences were re-annotated using Prokka ver. 1.11. The prophage region sequences were represented from Easyfig (http://mjsull.github.io/Easyfig/). Each CDS was compared by BLASTN. The percentage average nucleotide identity (ANI) was calculated by applying the ANIm tool in the JSpecies package(31). We also analyzed the number of spacer arrays and prophages in each genome. Acquired CRISPR loci were predicted below.

### Determining *emm* types

Coding sequences (CDSs) were searched using BLASTN against the *emm* database in the Centers for Disease Control and Prevention (ftp://ftp.cdc.gov/pub/infectious_diseases/biotech/emmsequ/) (29). CDSs that perfectly matched specific *emm* genes in the database were considered to be *emm* genes from the corresponding strain.

### Gene content and genome structure analyses

Pan and core genomes were defined using Roary v3.6 with the default parameters (32). Proteins with similar sequences were clustered using CD-Hit. The BLASTP cut-off value was set at 95%. Core- and pan-genome protein family numbers were estimated by genome sampling up to the number of input genomes at the default setting. The chromosomal locations of these genes were inspected to determine whether they were located in the recombination tracts predicted in a previous report by fastGEAR (33-34).

### Constructing phylogenetic trees

A maximum parsimony phylogenetic tree for the 59 *S*. *pyogenes* isolates (51 from databases and eight from this study) was constructed using kSNP3 v3.0 with a k-mer length of 19 (37). The Kchooser script in kSNP3 was used to estimate the optimum k-mer values. Constructed trees were visualized using FigTree v1.4.2 (http://tree.bio.ed.ac.uk/software/figtree).

### Predicting CRISPR/Cas loci

CRISPR loci were predicted using the CRISPR Recognition Tool ver. 1.2 with a minimum repeat length of 30, maximum repeat length of 48, and maximum spacer length of 80 (38). To characterize the spacer sequences, the spacer list was subjected to a BLASTN search against the following four databases: 1) the “gold standards” dataset of REBASE (accessed October 2016), 2) 59 *S*. *pyogenes* genome sequences, 3) predicted RM systems in 59 *S*. *pyogenes* strains, and 4) prophage region sequences in 59 *S*. *pyogenes* strains. The BLASTP cut-off value was set at 95% for identity and 90% for query coverage.

### Methylome analysis

Genomic DNA was sequenced on the PacBio RS II platform. Base modifications were analyzed using the RS_Modification_Detection.1 tool from the SMRT analysis package version 2.3.0. Briefly, after the reads were mapped to the genome sequences, the interpulse durations were measured for all nucleotide positions in the genomes and compared with the expected durations in a kinetic model of the polymerase (39) to identify significant associations. The number of methylated bases in a 1-kb window was counted with a sliding 500-bp window for each strand to overview the methylation distribution.

### Transcriptome analysis

RNA was extracted from three biological replicates per condition. RNA extraction and DNase treatment were performed using the RiboPure-Bacteria kit (Ambion; Austin, TX, USA), and rRNA was depleted with the Illumina Ribo-Zero rRNA Removal kit (Epicentre). RNA integrity was checked with an RNA nano chip (Agilent Technologies, catalog # 5067–1511) using the Agilent 2100 Bioanalyzer. Sequencing libraries were prepared and barcoded with the SureSelect Strand Specific RNA Library Prep system (Agilent Technologies, catalog # G9691A and G9691B). The libraries were sequenced with Hiseq3000 to generate 100 base-pair paired-end reads. Read quality was evaluated using FastQC v0.67 (http://www.bioinformatics.babraham.ac.uk/projects/fastqc/). The sequences were quality filtered and adapter trimmed using Trimmomatic v0.3625 under the default parameters. The reads were mapped to each genome by HISAT run with the default parameters. Mapped reads were visualized by the Integrative Genomics Viewer (IGV) v2.5 (http://www.broadinstitute.org/igv).

## RESULTS

### General features of the genome, RM system, and methylome

We sequenced eight *S*. *pyogenes* strains with a coverage range of 76–1372. The final assemblies were gathered into each one contig, and determined as complete sequences (Table 1). On average, a total of 1876 genes with 68 tRNA genes and 18 rRNA operons were predicted. We analyzed the whole genome data for all 59 *S*. *pyogenes* strains. The pan-genome size and the number of core genes were estimated, and we confirmed the previously proposed assertion (40) that *S*. *pyogenes* is an open pan-genome species (Supplementary Figure S1). The general characteristics of the genomes are shown in Table 1 and Supplementary Table S1.

We compared the RM systems from all 59 *S*. *pyogenes* strains using REBASE, which is a comprehensive and fully curated database of RM systems. We found type I and II RM systems in these 59 *S*. *pyogenes* strains (Table 2); none of the strains in the present study had type III or IV systems. Our methylome analysis of the eight newly sequenced strains showed that each strain has different methylation motifs (Table 3). A DNA synthesis kinetics analysis revealed that the predominant modification type was m6A, whereas other types (e.g., m4C and m5C) were infrequent but did exist (Supplementary Figure S2). By counting the number of methylated bases per 1,000 bases (Supplementary Figure S3), we found that the distribution of methylated bases across the genome differed on each strand. Pathogenesis-related genes, such as the fibronectin-binding protein, were identified in the densely methylated region (Supplementary Table S2). A motif analysis showed that all the motifs targeted by type 1 RMs on each genome were 100% methylated. In contrast, the other motifs were not. For example, only 51.4% of 5’-GMGGTAVYR motifs on the ATCC14918 genome were methylated. Finally, we clustered the methylated bases to identify motifs related to recognition sequences (Table 3).

**Table 2.**
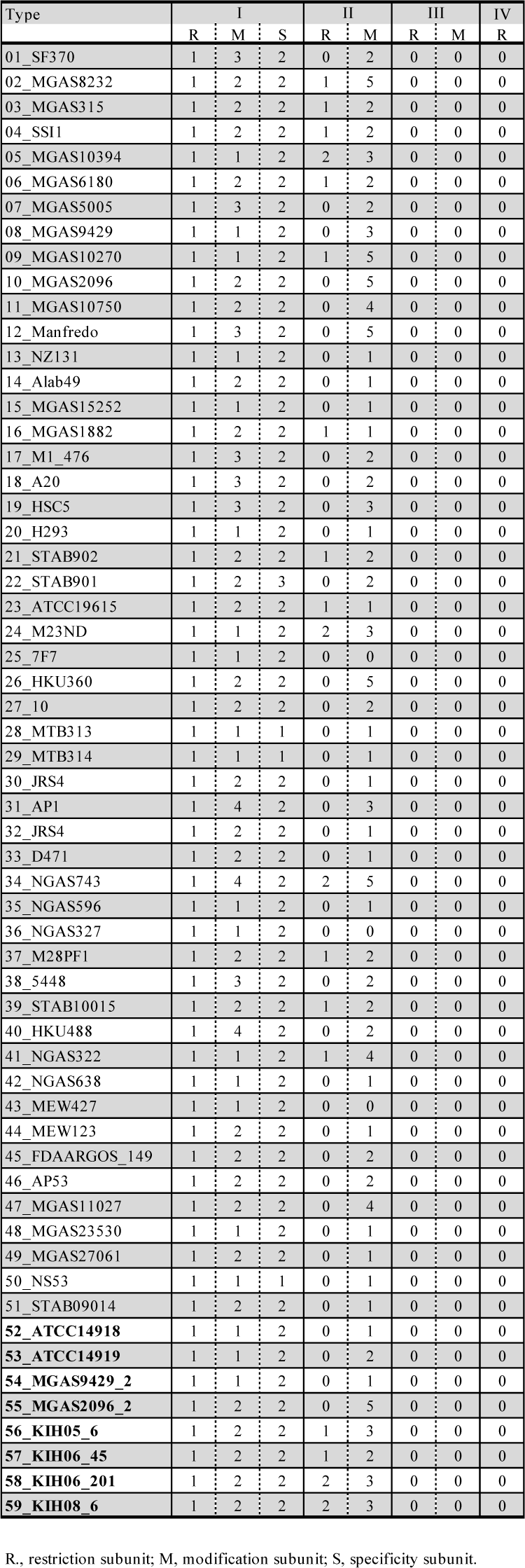
Distribution of the RM systems in the *S*. *pyogenes* strains used in this study.

**Table 3.**
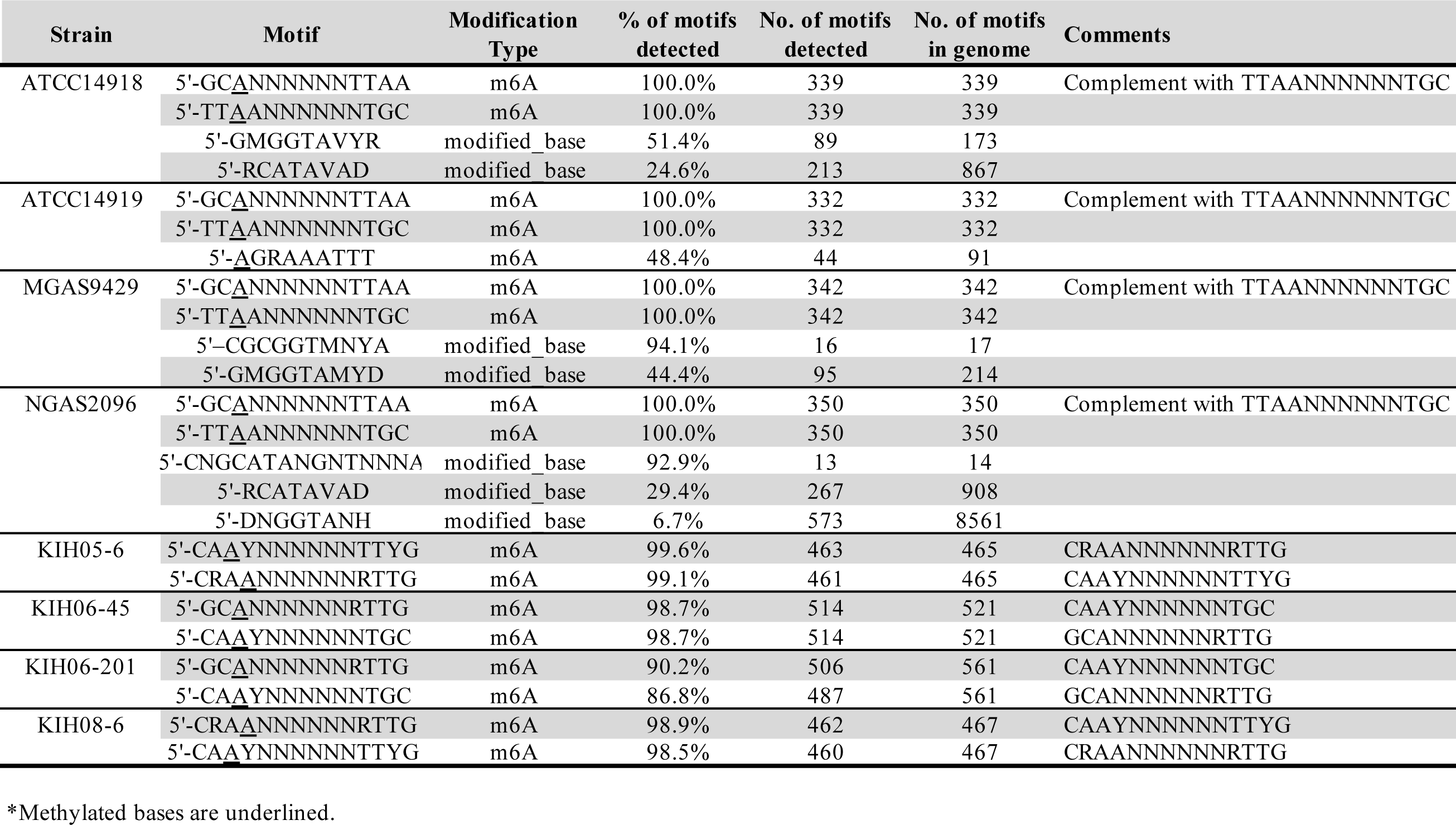
The methylation motifs in each strain.

### Relationship between type I RM systems and genome phylogeny

The S subunit of a type I RM system contains two tandem TRD sequences, TRD1 and TRD2 (Figure 1). In each TRD pair of the eight strains whose genome sequences were determined here, the combination of TRD1 and TRD2 in the four *emm*12 strains was the same and recognized GCANNNNNNTTAA (Figure 1, Table 3, and Supplementary Figure S4). Similarly, the TRD combination in the *emm*6 strains (KIH05-6, KIH08-6) was the same and recognized CAAYNNNNNNTTYG. The *emm* type differed between KIH06-45 and KIH06-201, but the TRD combination in these strains was the same and recognized GCANNNNNNRTTG.

**Figure 1.**
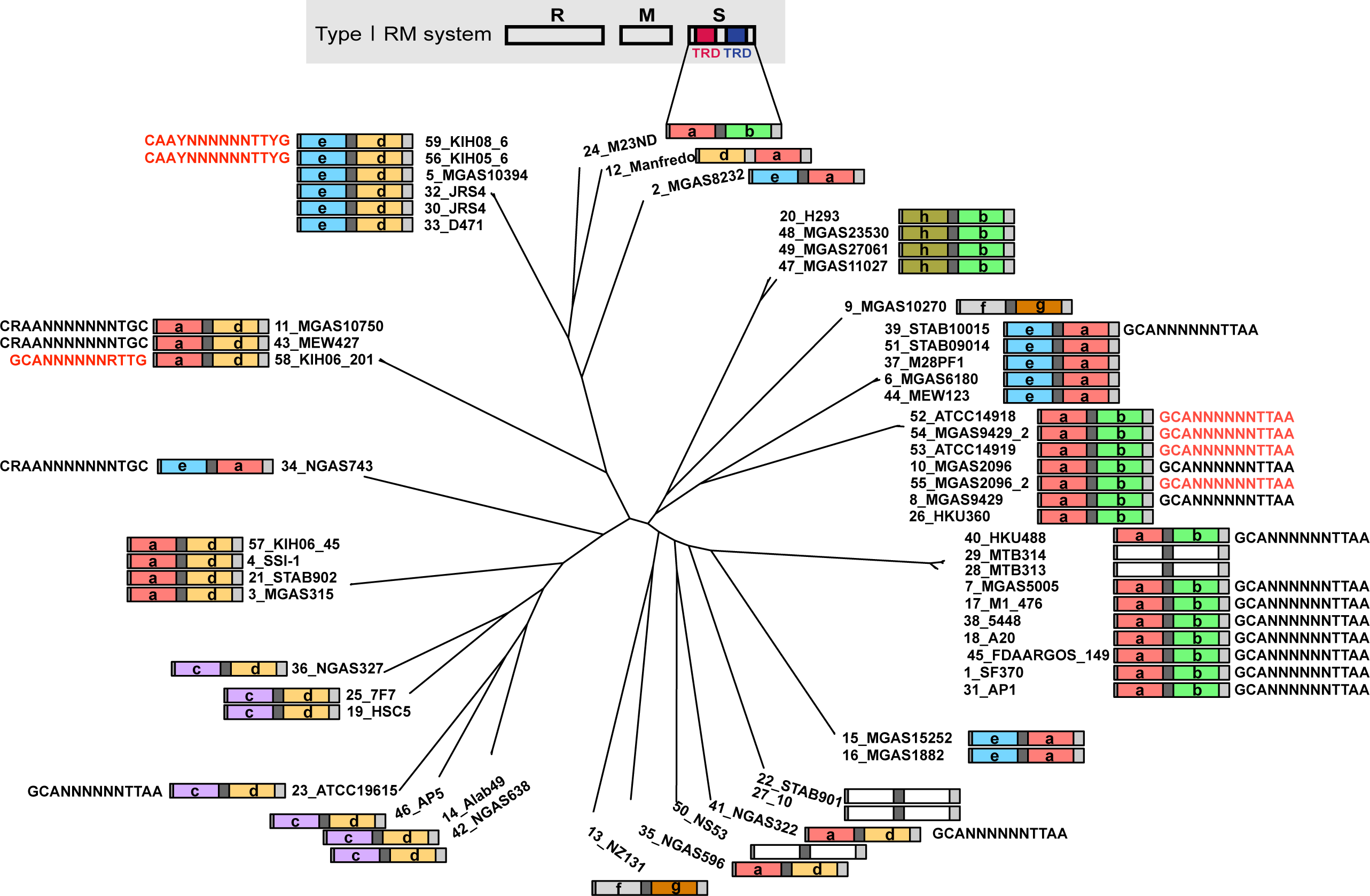
Distribution of type I target recognition domains (TRDs). A genomic tree of 59 *S*. *pyogenes* isolates and the TRD combinations for each isolate. The letters a, b, c, e, f, g, and h represent the TRD types. The recognition sequences of each type I RM system are shown where they are known. The recognition sequences detected in this study are colored in red.

We compared the maximum parsimony tree constructed using genome-wide SNPs with the type I TRD combination of *S*. *pyogenes* (Figure 1). With the clades represented by the *emm*12 strains, the TRD combinations were all a and b, meaning that they share the same recognition sequences (Supplementary Figure S4). The TRD combination appears to be mostly consistent with the phylogeny; however, despite being close in phylogeny, the 12_Manfredo strain TRD combination was different from that of the 24_M23ND strain.

### Features and distributions of the type II RM systems

We identified 127 type II RM systems in the 59 strains, and 104 of the genes from 41 of the strains were solitary MTases (Table 2). The MTase-encoding genes were found in close proximity to the REase gene or both MTase and REase were encoded by a single gene. When no closely located partners were predicted, the MTases were considered solitary MTases. Despite passing quality control and having high genome coverages, the motifs detected in *S*. *pyogenes* were not specific (Table 3). Compared with the type I motifs, these candidate type II motifs were not 100% methylated. For example, in ATCC14918, 51.4% of the 5’-GMGGTAVYRs were methylated, and 24.6% of the 5’-RCATAVADs were methylated. Hence, type II RM systems could change the methylome in *S*. *pyogenes* to a large extent.

We found that some type II MTases are located in the prophage region. As shown in Figure 2, the prophage sequences from four of the *emm*12 strains were aligned and compared. Strain ATCC14918 contains one prophage, whereas strains ATCC14919, MGAS9429_2, and MGAS2096_2 contain two prophage regions; however, they lack high sequence similarity values despite their close phylogenetic relationship. The prophage regions in ATCC14918 and ATCC14919 share 98% similarity by ANI (Table 4), but the only prophage in ATCC14918 encodes an MTase. We detected 121 type II MTase genes in the 59 *S*. *pyogenes* genomes, with 58 of them encoded in the prophage region as solitary genes (Figure 3). The phylogenetic tree we constructed for the *S*. *pyogenes* genomes fell into two clades: a prophage-dominant clade and a CRISPR-dominant one. In the prophage-dominant group, the strains were found to contain various MTases, 69% of which are on prophages. In the CRISPR-dominant group, 31% of the MTases were found on the prophage. While 19 strains were found to lack a CRISPR-Cas system completely, 18 belong to prophage-dominant groups.

**Table 4.** Average Nucleotide Identity (ANI) (%) based on prophage sequences.

| ANIb | ATCC14918-Φ1 | ATCC14919-Φ1 | MGAS9429 2-Φ1 | MGAS9429 2-Φ2 | MGAS2096 2-Φ1 | MGAS2096 2-Φ2 |
| --- | --- | --- | --- | --- | --- | --- |
| ATCC14918-Φ1 | --- | 96.56 | 100 | NaN | 91.06 | 96.56 |
| ATCC14919-Φ1 | 96.38 | --- | 96.38 | NaN | 99.46 | 100 |
| MGAS9429 2-Φ1 | 100 | 96.56 | --- | NaN | 91.06 | 96.56 |
| MGAS9429 2-Φ2 | NaN | NaN | NaN | --- | NaN | 77.59 |
| MGAS2096 2-Φ1 | 88.32 | 100 | 88.32 | 80.41 | --- | 100 |
| MGAS2096 2-Φ2 | 96.29 | 100 | 96.29 | 77.01 | 100 | --- |

**Figure 2.**
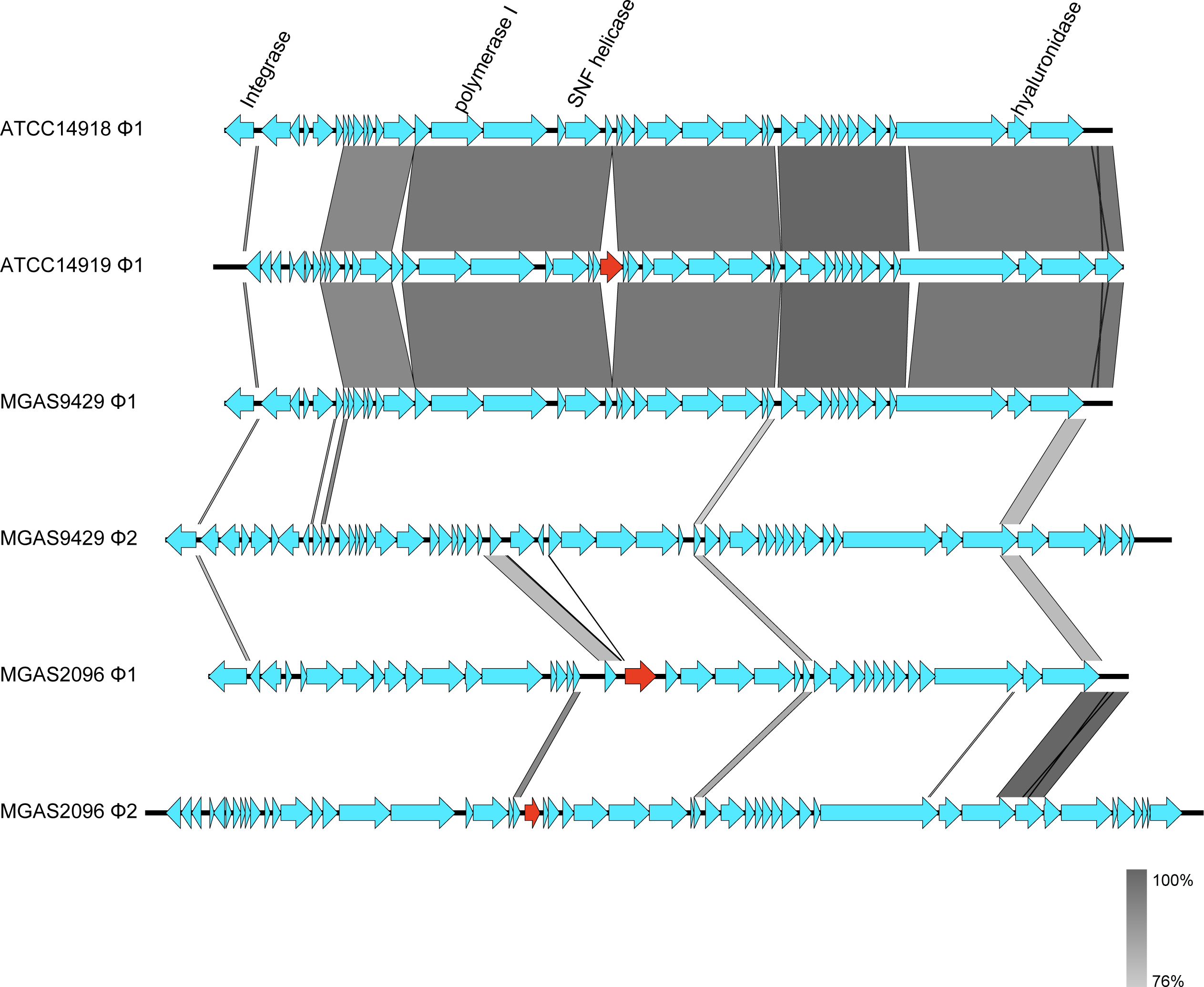
Sequence comparison of prophage regions integrated on the chromosome in *S*. *pyogenes* strains ATCC14918, ATCC14919, MGAS9429, and MGAS2096. Genome organization representation from Easyfig (http://mjsull.github.io/Easyfig/). Each CDS was compared by BLASTN. The type II Methylases are shown as red arrow.

**Figure 3.**
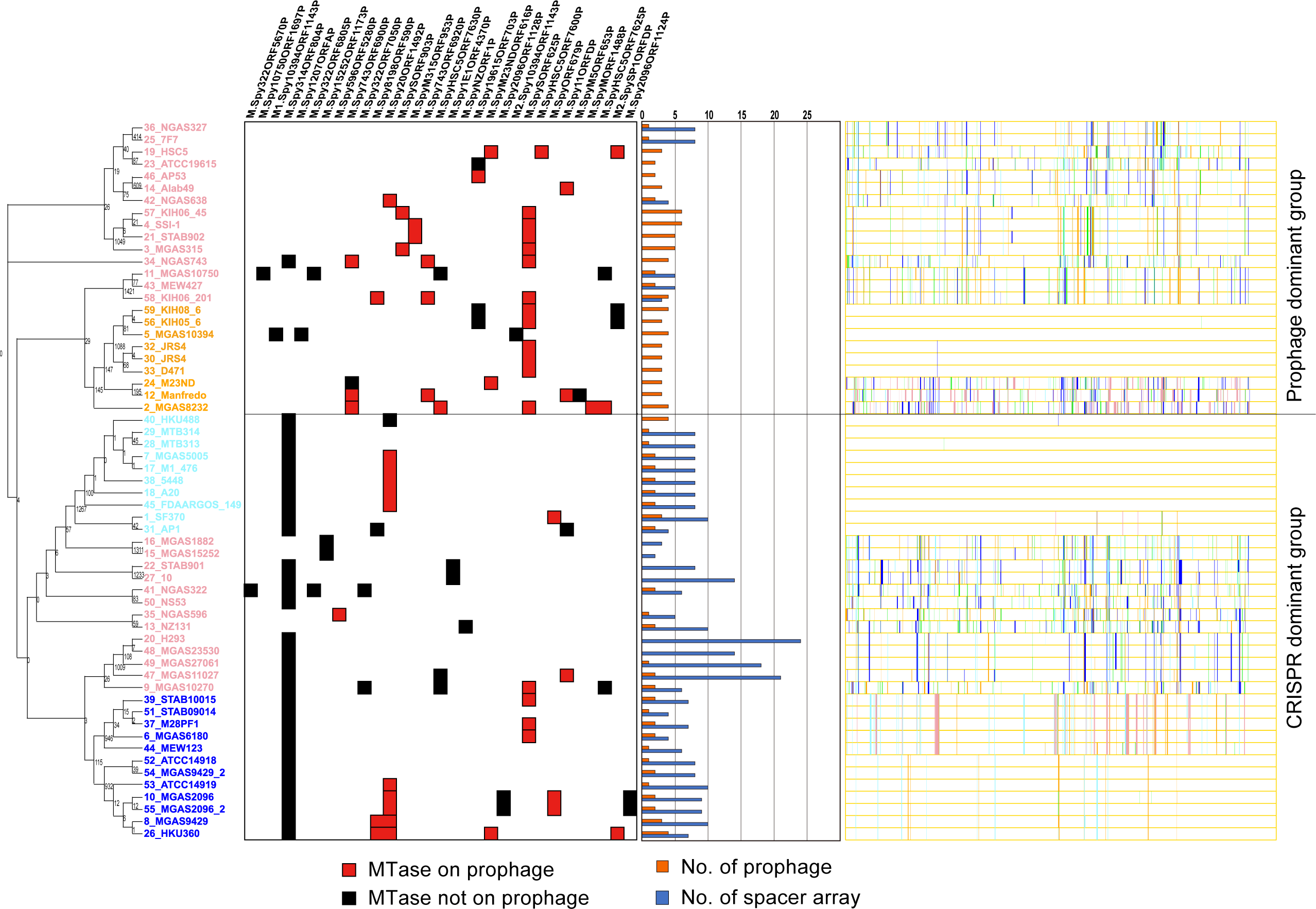
Distribution of type II MTases and the numbers of prophages and CRISPR spacers. The vertical column names are the MTases detected in this study. Left: An SNP-based genomic tree and the isolate names are shown on the left. The number of SNPs is shown on each node. The strains that we sequenced in this study are listed in red. Middle: Black boxes indicate the presence of corresponding type II MTases. Red boxes indicate the presence of MTases in the prophage regions. The bar chart shows the number of spacers (blue) and prophages (orange) in each genome. Right: Colors indicate donor lineage (population group) of the recent genomic imports. *S*. *pyogenes* lineage and recombination were inferred using fastGEAR software (Mostowy et al. 2017).

### CRISPR spacers exhibiting high nucleotide similarity in the genome

We identified 317 CRISPR spacers in the 59 isolates. The number of spacers in all the loci varied among the isolates, ranging from 0 to 24. We obtained unique spacer lists for each CRISPR locus. Nucleotide similarity searches enabled us to also analyze the spacers from the genomes of the 59 *S*. *pyogenes* isolates (Supplementary Tables S3-S6). Four predicted MTases (M.Spy19615ORF703P, M.Spy743ORF6920P, M.SpyHSC5ORF7630P, and M.SpyM23NDORF616P) were found to have high similarity with the spacers from ten strains, and two others (M.SpyHSC5ORF7625P and M2.SpySP1ORFDP) have high similarity with the spacers from eight strains. All these type II MTases were encoded on the prophage region rather than in the chromosome. We found that 206 spacers share high nucleotide similarity with six MTase sequences from the 59 assessed *S*. *pyogenes* strains.

### Comparison of the methylation and transcriptome of virulence genes between *emm*12 strains

To investigate the possible role of DNA methylation in *S*. *pyogenes*, we examined the RNA-Seq results for six virulence genes, *speB, speC, csrR, csrA, mga*, and *nga*, in four *emm*12 strains and conducted a motif analysis. For each region and its upper 20 bases, methylated motifs and mapped reads were compared for both strands (Figure 4). The IGV showed that RNA mapping of *speB* in the four strains clearly differed, with the mRNA expressed on only one strand in ATCC14918. Although the RNA was isolated from the exponential phase, the coverage of each gene differed. The transcriptional level of a region showed no correlation with the location or type of methylated motifs. In ATCC14918 and MGAS9429, *csrR* and *csrA* genes are encoded by two-component systems.

**Figure 4.**
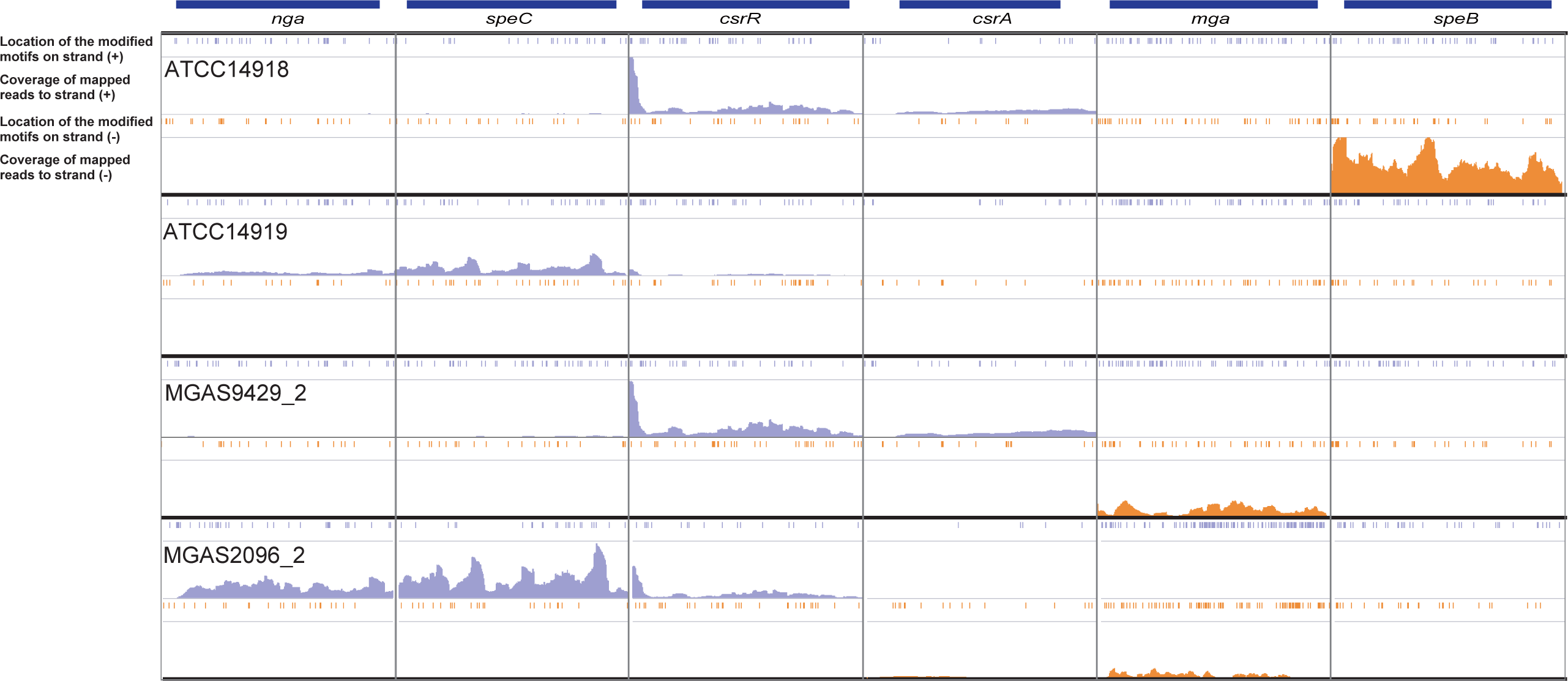
Integrative Genomics Viewer (IGV) representation of mapped Illumina strand-specific RNA-Seq reads versus the locations of methylated motifs in six loci. The RNA-Seq coverage map shows six gene regions from four *emm*12 strains. The location of each motif and the mapping coverage are shown on each strand from the four isolates. The coverage range is 0 to 2000.

## DISCUSSION

In bacteria, the number of type I RM systems varies among species, and the RM systems are usually encoded by 5 to 10 genes in the genome (41). For example, in *Helicobacter pylori*, about 30 type I RM systems are encoded in the chromosomal region of each genome. In contrast, only one or two type I RM system genes are encoded in the *S*. *pyogenes* genome, which is a smaller amount than that in other streptococcal species, and these genes are not highly variable (Table 2).

Shuffling of TRDs as “phase variation” has been described in several species over the last 20 years (42-44). One example of phase variation is the switching of TRDs in the type I RM systems of some species, which produces multiple methyltransferase variants and, consequently, variable methylation (11). Another example is type III TRD switching in *Haemophilus influenzae, Neisseria gonorrhoeae*, and *Neisseria meningitidis*, where such switching can regulate the expression of multiple genes. Recently, a type I TRD *S*. *pyogenes* knockout strain was found to show different methylation and expression profiles from those of the wildtype, but the strain lacked TRD switching, and the phenotype of the knockout was not different from that of the wildtype (45). Hence, the switching of different TRD combinations is considered to make a relatively small contribution to phase variation in *S*. *pyogenes*.

In the present study, although the type II MTases of the four *emm*12 strains were all prophage-encoded” solitary MTases”, we found that their positions and types were dissimilar. Among our 59 *S*. *pyogenes* strains, many of the solitary MTases are on prophages, and the distribution of the type II RM systems is not consistent with the phylogeny (Figure 3). The genome tree was not directly consistent with the distribution of the type II MTases. The clustering determined by fastGEAR divided the genomic tree into two groups: a prophage-dominant group and a CRISPR-dominant group (Figure 3). In the prophage-dominant group, more MTases tended to be encoded in the prophage region compared with in the CRISPR-dominant group. It is already known that phage can disseminate various genes to bacteria (e.g., virulence genes), and the same may also apply to MTases. The similarity and structure analysis of the prophage region from the four closely-related strains revealed the presence of multiple MTases in this region (Figure 2, Supplementary Figure S5). All these MTases were type II solitary MTases, and type I RM was not encoded. Although the chromosomes are closely related in these four strains, their prophages did not share any sequence similarities (Table 4), suggesting that genetic diversification in *S*. *pyogenes* has frequently occurred via prophage acquisition.

It is known that the CRISPR spacers in *S*. *pyogenes* share sequence similarity with prophage sequences. It is also known that the presence of few CRISPR spacers in the genome increases the chance of *S*. *pyogenes* acquiring and integrating more prophages (46). This suggests that *S*. *pyogenes* use CRISPR to control prophage invasion, and this facilitates genetic diversification. In this study, the sequences of some spacers showed high similarity to those of MTases, suggesting that genetic diversification in *S*. *pyogenes* has accelerated via acquiring MTase-encoding phages. It is thought that the main reason phages have MTases is to avoid the host defense mechanism, i.e., the RM system, which is incorporated into the host genome. However, it is possible that the host is actively incorporating phages that contain MTase, which in turn can diversify its methylome. That MTase-targeted spacers exist in bacteria supports the notion that they are involved in an evolutionary arms race between bacteria and phages (47). Similarly, in *S*. *pyogenes*, the number of spacers in its genome is negatively correlated with the number of phages (46) (48), and the number of MTases encoded on phages in the prophage-dominant group is also significantly higher than that in the CRISPR-dominant group, suggesting that an evolutionary arms race also exists in *S*. *pyogenes*.

Methylome diversification, as driven by changes in the TRD combinations present in a genome, has been reported in some species (41). The present study is the first to analyze the methylase distribution in *S*. *pyogenes* strains comprehensively. Further analysis of four closely related strains enabled us to identify some virulence factors and various methylation sites (and their sequences) among the strains. Additionally, pathogenesis-related genes, such as the fibronectin-binding protein, were identified in the densely methylated region (Supplementary Table S2), but the biological significance of these instances of hypermethylation is not yet clear. These results suggest that phage-derived methylome diversification may affect the pathogenicity profiles of *S*. *pyogenes* strains.

An active methylase encoded on a prophage in *S*. *pyogenes* has already been reported (49). Expression levels and methylome changes in *S*. *pyogenes* have also been described quite recently (45). The authors of that study elucidated the contribution of the type I RM system to the methylome and phenotype. The present study differs in that the RM systems in multiple *S*. *pyogenes* strains were comprehensively detected and compared with the methylome. Our methylome analysis was conducted in four closely related strains so that we could compare the transcriptome and methylome analyses. The mRNA expression level of each allele was dissimilar, the type and position of the recognition sequence in the genome also differed (Figure 4), and the four strains fell within the CRISPR-dominant group (Figure 3).

## CONCLUSION

To understand the effect of the RM system as related to genome diversification in *S*. *pyogenes*, we acquired and compared the complete genome sequences of 59 *S*. *pyogenes* strains. We found that the *S*. *pyogenes* RM system is diversely distributed in the genome, likely at least partially due to phage-driven gene transfer. Diversification of DNA methylation was also observed in the strains. These findings support the hypothesis that phages can be incorporated into the *S*. *pyogenes* genome by circumventing the biological defense mechanism of the restriction system by retaining their methylation enzymes. Furthermore, it seems likely that the diversity of DNA modification seen in our strains is regulated by methylases derived from external factors, or phages. It will be necessary to confirm that expression levels and phenotype changes have occurred by using a gene disruption strain in the future. Furthermore, to avoid possible bias relating to the isolation sources of the database strains, the inclusion of additional strains isolated from various situations will be necessary to provide informational and statistical robustness to similar studies. It would also be worthwhile to clarify the effect of DNA modification in *S*. *pyogenes* on the disease condition by further analyzing its epigenome under various disease states.

## ACCESSION NUMBERS

The genome sequences described in this study have been deposited in DDBJ/DRA/GenBank under the accession no. DRA004914 (BioProject PRJDB4542). The data described here can be freely and openly accessed in GenBank.

## ACKNOWLEDGMENTS

We thank Sandra Cheesman, Ph.D., and Katie Oakley, Ph.D., from Edanz Group (www.edanzediting.com/ac) for editing drafts of this manuscript.

## REFERENCES

1. Vasu, K., Vasu, K., Nagaraja, V. and Nagaraja, V. (2013) Diverse functions of restriction-modification systems in addition to cellular defense. Microbiol. Mol. Biol. Rev., 77, 53–72.

2. Ershova, A.S., Rusinov, I.S., Spirin, S.A., Karyagina, A.S. and Alexeevski, A.V. (2015) Role of restriction-modification systems in prokaryotic evolution and ecology. Biochemistry Mosc., 80, 1373–1386.

3. Donczew, R., Zakrzewska-Czerwinska, J. and Zawilak-Pawlik, A. (2014) Beyond dnaA: the role of DNA topology and DNA methylation in bacterial replication initiation. J. Mol. Biol., 426, 2269–2282.

4. Adhikari, S. and Curtis, P.D. (2016) DNA methyltransferases and epigenetic regulation in bacteria. FEMS Microbiol. Rev., 40, 575–591.

5. Sánchez-Romero, M.A. and Casadesús, J. (2019) The bacterial epigenome. Nat. Rev. Microbiol., 1, 76–14.

6. Oliveira, P.H. and Fang, G. (2020) Conserved DNA methyltransferases: A window into fundamental mechanisms of epigenetic regulation in bacteria. Trends Microbiol., 10.1016/j.tim.2020.04.007.

7. Oliveira, P.H., Touchon, M. and Rocha, E.P.C. (2014) The interplay of restriction-modification systems with mobile genetic elements and their prokaryotic hosts. Nucleic Acids Res., 42, 10618–10631.

8. Manso, A.S., Chai, M.H., Atack, J.M., Furi, L., De Ste Croix, M., Haigh, R., Trappetti, C., Ogunniyi, A.D., Shewell, L.K., Boitano, M., et al. (2014) A random six-phase switch regulates pneumococcal virulence via global epigenetic changes. Nat. Commun., 5, 5055.

9. Furuta, Y., Konno, M., Osaki, T., Yonezawa, H., Ishige, T., Imai, M., Shiwa, Y., Shibata-Hatta, M., Kanesaki, Y., Yoshikawa, H., et al. (2015) Microevolution of virulence-related genes in *Helicobacter pylori* familial infection. PLoS ONE, 10, e0127197.

10. Zheng, H., Dietrich, C., Hongoh, Y. and Brune, A. (2016) Restriction-modification systems as mobile genetic elements in the evolution of an intracellular symbiont. Mol. Bio. Evol., 33, 721–725.

11. Atack, J.M., Tan, A., Bakaletz, L.O., Jennings, M.P. and Seib, K.L. (2018) Phasevarions of bacterial pathogens: Methylomics sheds new light on old enemies. Trends Microbiol., 26, 715–726.

12. Phillips, Z.N., Tram, G., Seib, K.L. and Atack, J.M. (2019) Phase-variable bacterial loci: how bacteria gamble to maximise fitness in changing environments. Biochem. Soc. Trans., 47, 1131–1141.

13. Atack, J.M., Srikhanta, Y.N., Fox, K.L., Jurcisek, J.A., Brockman, K.L., Clark, T.A., Boitano, M., Power, P.M., Jen, F.E.-C., McEwan, A.G., et al. (2015) A biphasic epigenetic switch controls immunoevasion, virulence and niche adaptation in non-typeable *Haemophilus influenzae*. Nat. Commun., 6, 7828.

14. Fang, G., Munera, D., Friedman, D.I., Mandlik, A., Chao, M.C., Banerjee, O., Feng, Z., Losic, B., Mahajan, M.C., Jabado, O.J., et al. (2012) Genome-wide mapping of methylated adenine residues in pathogenic *Escherichia coli* using single-molecule real-time sequencing. Nat. Biotechnol., 30, 1232–1239.

15. Murray, I.A., Clark, T.A., Morgan, R.D., Boitano, M., Anton, B.P., Luong, K., Fomenkov, A., Turner, S.W., Korlach, J. and Roberts, R.J. (2012) The methylomes of six bacteria. Nucleic Acids Res., 40, 11450–11462.

16. Low, D.A. and Casadesús, J. (2008) Clocks and switches: bacterial gene regulation by DNA adenine methylation. Curr. Opin. Microbiol., 11, 106–112.

17. Ben Zakour, N.L., Davies, M.R., You, Y., Chen, J.H.K., Forde, B.M., Stanton-Cook, M., Yang, R., Cui, Y., Barnett, T.C., Venturini, C., et al. (2015) Transfer of scarlet fever-associated elements into the group A Streptococcus M1T1 clone. Sci. Rep., 5, 15877.

18. Chalker, V., Jironkin, A., Coelho, J., Al-Shahib, A., Platt, S., Kapatai, G., Daniel, R., Dhami, C., Laranjeira, M., Chambers, T., et al. (2017) Genome analysis following a national increase in scarlet fever in England 2014. BMC Genomics, 18, 224.

19. You, Y., Davies, M.R., Protani, M., McIntyre, L., Walker, M.J. and Zhang, J. (2018) Scarlet fever epidemic in China caused by *Streptococcus pyogenes* serotype M12: Epidemiologic and molecular analysis. EBioMedicine, 28, 128–135.

20. Hamada, S., Kawabata, S. and Nakagawa, I. (2015) Molecular and genomic characterization of pathogenic traits of group A *Streptococcus pyogenes*. Proc. Jpn. Acad., Ser. B, Phys. Biol. Sci., 91, 539–559.

21. Kreikemeyer, B., McIver, K.S. and Podbielski, A. (2003) Virulence factor regulation and regulatory networks in *Streptococcus pyogenes* and their impact on pathogen-host interactions. Trends Microbiol., 11, 224–232.

22. Kachroo, P., Eraso, J.M., Beres, S.B., Olsen, R.J., Zhu, L., Nasser, W., Bernard, P.E., Cantu, C.C., Saavedra, M.O., Arredondo, M.J., et al. (2019) Integrated analysis of population genomics, transcriptomics and virulence provides novel insights into *Streptococcus pyogenes* pathogenesis. Nat. Genet., 7, e00403–16.

23. Beres, S.B., Richter, E.W., Nagiec, M.J., Sumby, P., Porcella, S.F., DeLeo, F.R. and Musser, J.M. (2006) Molecular genetic anatomy of inter- and intraserotype variation in the human bacterial pathogen group A Streptococcus. Proc. Natl. Acad. Sci. U.S.A., 103, 7059–7064.

24. Maruyama, F., Kobata, M., Kurokawa, K., Nishida, K., Sakurai, A., Nakano, K., Nomura, R., Kawabata, S., Ooshima, T., Nakai, K., et al. (2009) Comparative genomic analyses of *Streptococcus mutans* provide insights into chromosomal shuffling and species-specific content. BMC Genomics, 10, 358.

25. Chin, C.-S., Peluso, P., Sedlazeck, F.J., Nattestad, M., Concepcion, G.T., Clum, A., Dunn, C., O’Malley, R., Figueroa-Balderas, R., Morales-Cruz, A., et al. (2016) Phased diploid genome assembly with single-molecule real-time sequencing. Nat. Methods, 13, 1050–1054.

26. Koren, S., Walenz, B.P., Berlin, K., Miller, J.R., Bergman, N.H. and Phillippy, A.M. (2017) Canu: scalable and accurate long-read assembly via adaptive k-mer weighting and repeat separation. Genome Res., 27, 722–736.

27. Seemann, T. (2014) Prokka: rapid prokaryotic genome annotation. Bioinformatics, 30, 2068–2069.

28. Roberts, R.J., Vincze, T., Posfai, J. and Macelis, D. (2015) REBASE--a database for DNA restriction and modification: enzymes, genes and genomes. Nucleic Acids Res., 43, D298–9.

29. Altschul, S.F., Gish, W., Miller, W., Myers, E.W. and Lipman, D.J. (1990) Basic local alignment search tool. J. Mol. Biol., 215, 403–410.

30. Akhter, S., Aziz, R.K. and Edwards, R.A. (2012) PhiSpy: a novel algorithm for finding prophages in bacterial genomes that combines similarity- and composition-based strategies. Nucleic Acids Res., 40, e126–e126.

31. Richter, M. and Rosselló-Móra, R. (2009) Shifting the genomic gold standard for the prokaryotic species definition. Proc. Natl. Acad. Sci. U.S.A., 106, 19126–19131.

32. Page, A.J., Cummins, C.A., Hunt, M., Wong, V.K., Reuter, S., Holden, M.T.G., Fookes, M., Falush, D., Keane, J.A. and Parkhill, J. (2015) Roary: rapid large-scale prokaryote pan genome analysis. Bioinformatics, 31, 3691–3693.

33. Yano, H., Iwamoto, T., Nishiuchi, Y., Nakajima, C., Starkova, D.A., Mokrousov, I., Narvskaya, O., Yoshida, S., Arikawa, K., Nakanishi, N., et al. (2017) Population structure and local adaptation of MAC lung disease agent *Mycobacterium avium* subsp. hominissuis. Genome Bio. Evol., 9, 2403–2417.

34. Mostowy, R., Croucher, N.J., Andam, C.P., Corander, J., Hanage, W.P. and Marttinen, P. (2017) Efficient inference of recent and ancestral recombination within bacterial populations. Mol. Bio. Evol., 34, 1167–1182.

35. Gardner, S.N., Slezak, T. and Hall, B.G. (2015) kSNP3.0: SNP detection and phylogenetic analysis of genomes without genome alignment or reference genome. Bioinformatics, 31, 2877–2878.

36. Bland, C., Ramsey, T.L., Sabree, F., Lowe, M., Brown, K., Kyrpides, N.C. and Hugenholtz, P. (2007) CRISPR recognition tool (CRT): a tool for automatic detection of clustered regularly interspaced palindromic repeats. BMC Bioinformatics, 8, 209.

37. Schadt, E.E., Banerjee, O., Fang, G., Feng, Z., Wong, W.H., Zhang, X., Kislyuk, A., Clark, T.A., Luong, K., Keren-Paz, A., et al. (2013) Modeling kinetic rate variation in third generation DNA sequencing data to detect putative modifications to DNA bases. Genome Res., 23, 129–141.

38. Maruyama, Fumito, et al. (2016) “Streptococcus pyogenes genomics.” Streptococcus pyogenes: Basic biology to clinical manifestations, edited by Joseph J. Ferretti et. al., University of Oklahoma Health Sciences Center.

39. Furuta, Y., Namba-Fukuyo, H., Shibata, T.F., Nishiyama, T., Shigenobu, S., Suzuki, Y., Sugano, S., Hasebe, M. and Kobayashi, I. (2014) Methylome diversification through changes in DNA methyltransferase sequence specificity. PLoS Genet., 10, e1004272.

40. Furuta, Y. and Kobayashi, I. (2012) Movement of DNA sequence recognition domains between non-orthologous proteins. Nucleic Acids Res., 40, 9218–9232.

41. De Ste Croix, M., Vacca, I., Kwun, M.J., Ralph, J.D., Bentley, S.D., Haigh, R., Croucher, N.J. and Oggioni, M.R. (2017) Phase-variable methylation and epigenetic regulation by type I restriction-modification systems. FEMS Microbiol. Rev., 41, S3–S15.

42. Oliver, M.B., Basu Roy, A., Kumar, R., Lefkowitz, E.J. and Swords, W.E. (2017) *Streptococcus pneumoniae* TIGR4 phase-locked opacity variants differ in virulence phenotypes. mSphere, 2, 1543.

43. Nye, T.M., Jacob, K.M., Holley, E.K., Nevarez, J.M., Dawid, S., Simmons, L.A. and Watson, M.E. (2019) DNA methylation from a type I restriction modification system influences gene expression and virulence in *Streptococcus pyogenes*. PLoS Pathog., 15, e1007841.

44. Nozawa, T., Furukawa, N., Aikawa, C., Watanabe, T., Haobam, B., Kurokawa, K., Maruyama, F. and Nakagawa, I. (2011) CRISPR inhibition of prophage acquisition in *Streptococcus pyogenes*. PLoS ONE, 6, e19543.

45. Endo, A., Watanabe, T., Ogata, N., Nozawa, T., Aikawa, C., Arakawa, S., Maruyama, F., Izumi, Y. and Nakagawa, I. (2015) Comparative genome analysis and identification of competitive and cooperative interactions in a polymicrobial disease. ISME J., 9, 629–642.

46. Yamada, S., Shibasaki, M., Murase, K., Watanabe, T., Aikawa, C., Nozawa, T. and Nakagawa, I. (2019) Phylogenetic relationship of prophages is affected by CRISPR selection in Group A Streptococcus. BMC Microbiol., 19, 24.

47. Euler, C.W., Ryan, P.A., Martin, J.M. and Fischetti, V.A. (2007) M.SpyI, a DNA methyltransferase encoded on a mefA chimeric element, modifies the genome of *Streptococcus pyogenes*. J. Bacteriol., 189, 1044–1054.

