## Supplemental Figures and Tables for "Diversification of the restriction–modification system of *Streptococcus pyogenes* through its acquisition of mobile elements"

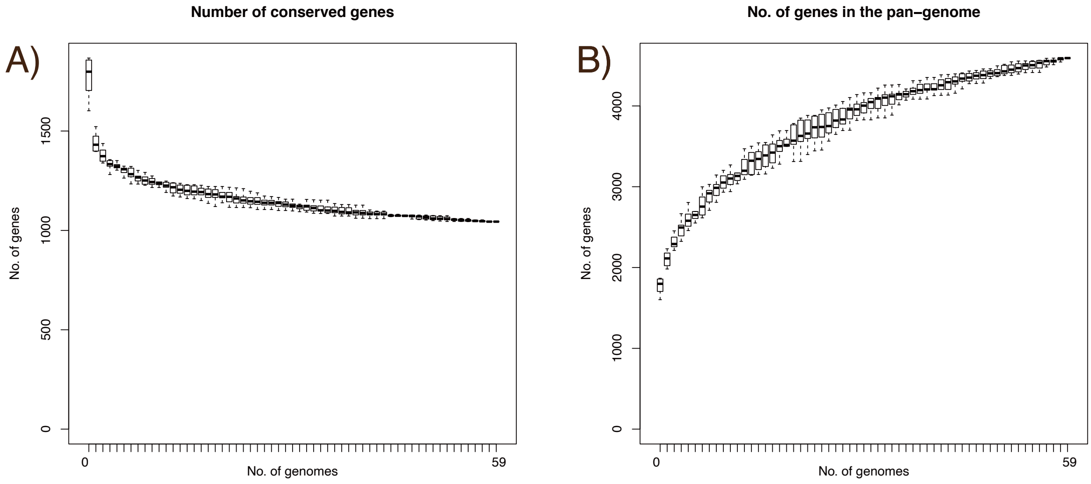


**Supplementary Figure S1: Pan-genome plots of *S. pyogenes* gene numbers.** The number of specific genes is plotted as a function of the number of sequentially added strains. The right curve is the calculated pan-genome size.

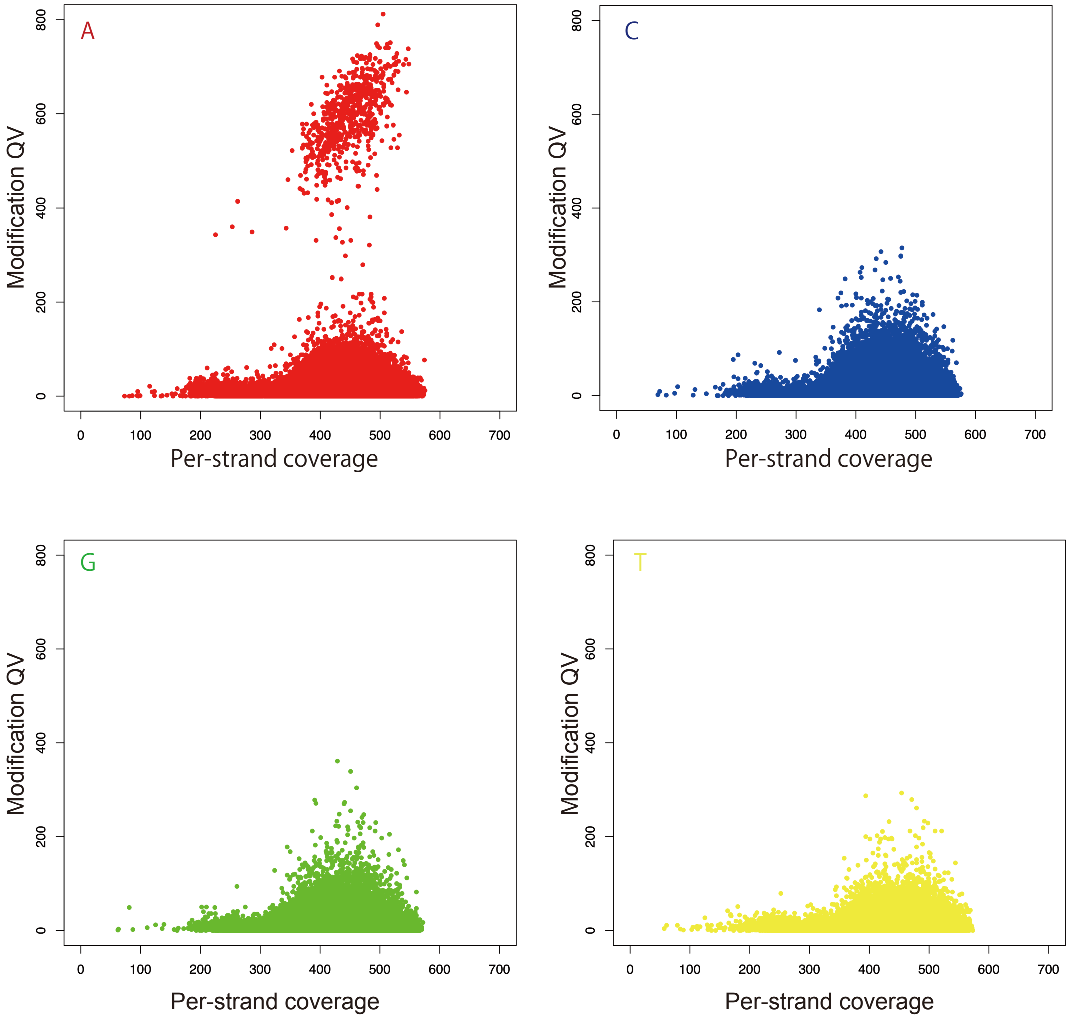


**Supplementary Figure S2: Detection of N6-methyladenine (m6A) base modification in ATCC14918.** The peak of the scatter plots indicates an increase in methylated A, C, G, and T bases in ATCC14918, an indication of 6mA sites being present in ATCC14918.

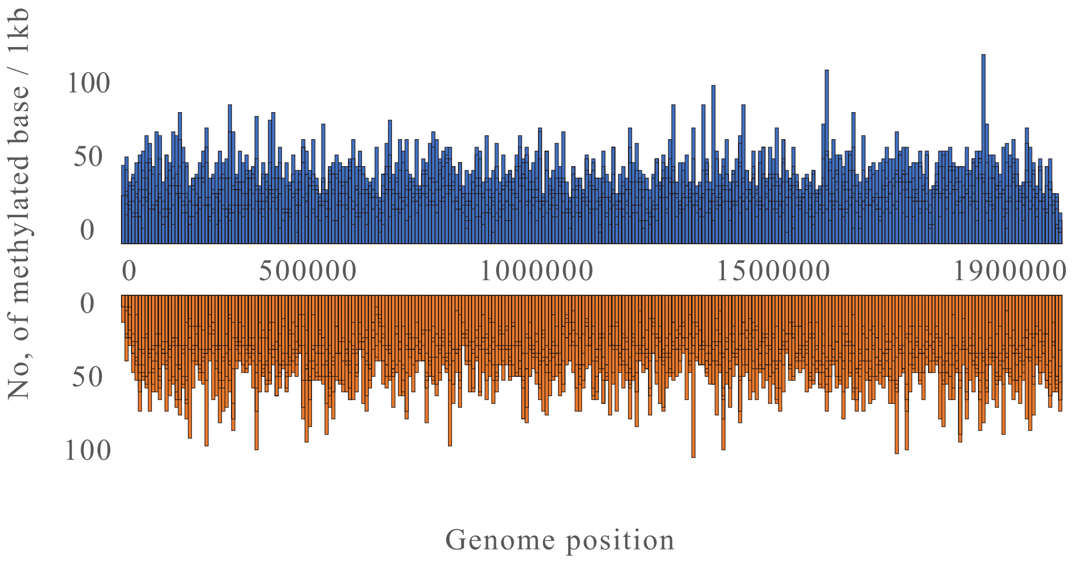


**Supplementary Figure S3: Distribution of methylated bases in genomes and hypermethylated genes.** Density of methylated bases on each strand of the *S. pyogenes* genome.

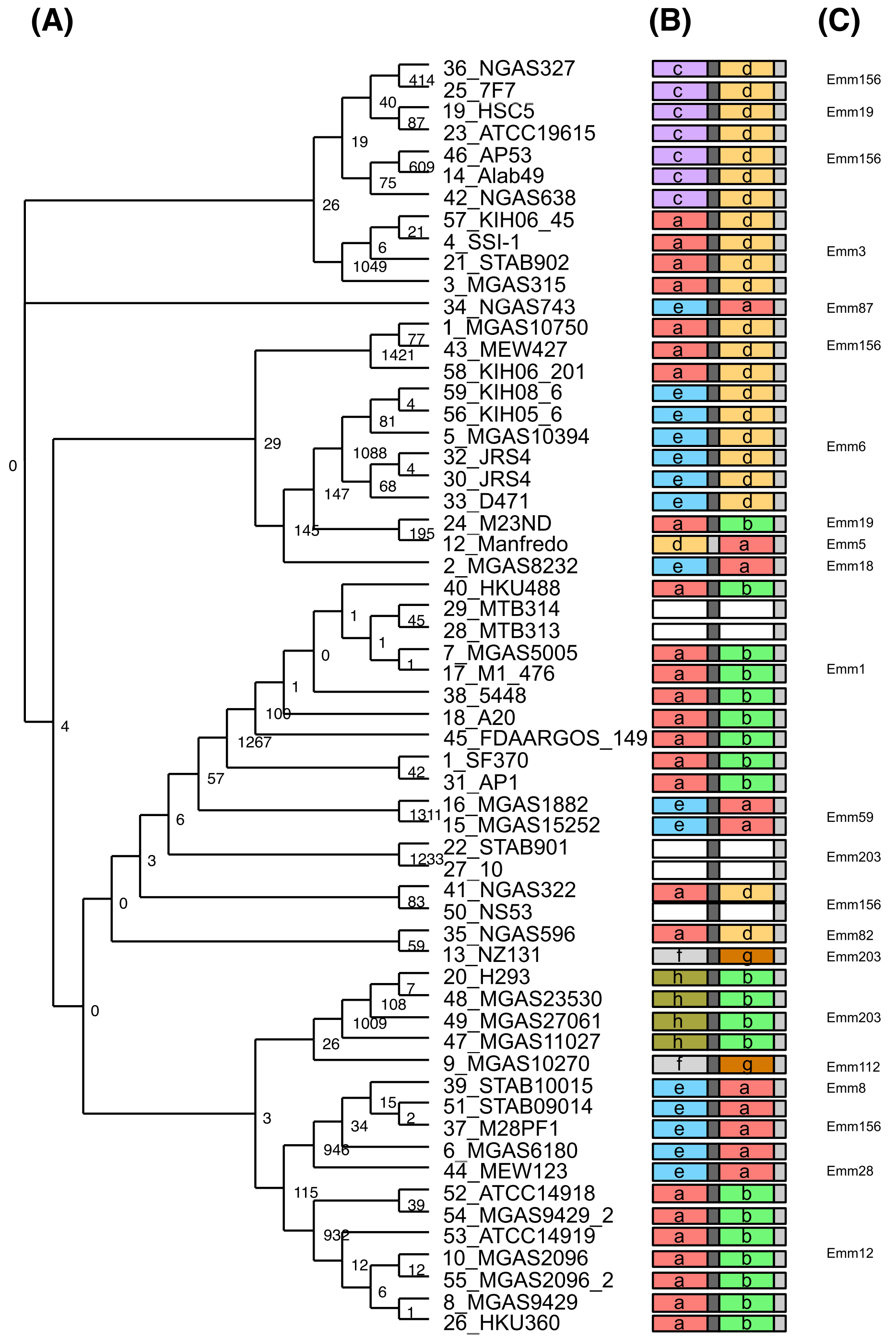


**Supplementary Figure S4: Distribution of type I RM systems compared with *emm* type.** (A) Genomic tree of TRDs of type I RM systems on 59 *S. pyogenes* strains. (B) Organization of TRDs in each strain. The colors correspond to type. (C) *emm* types on each clade.

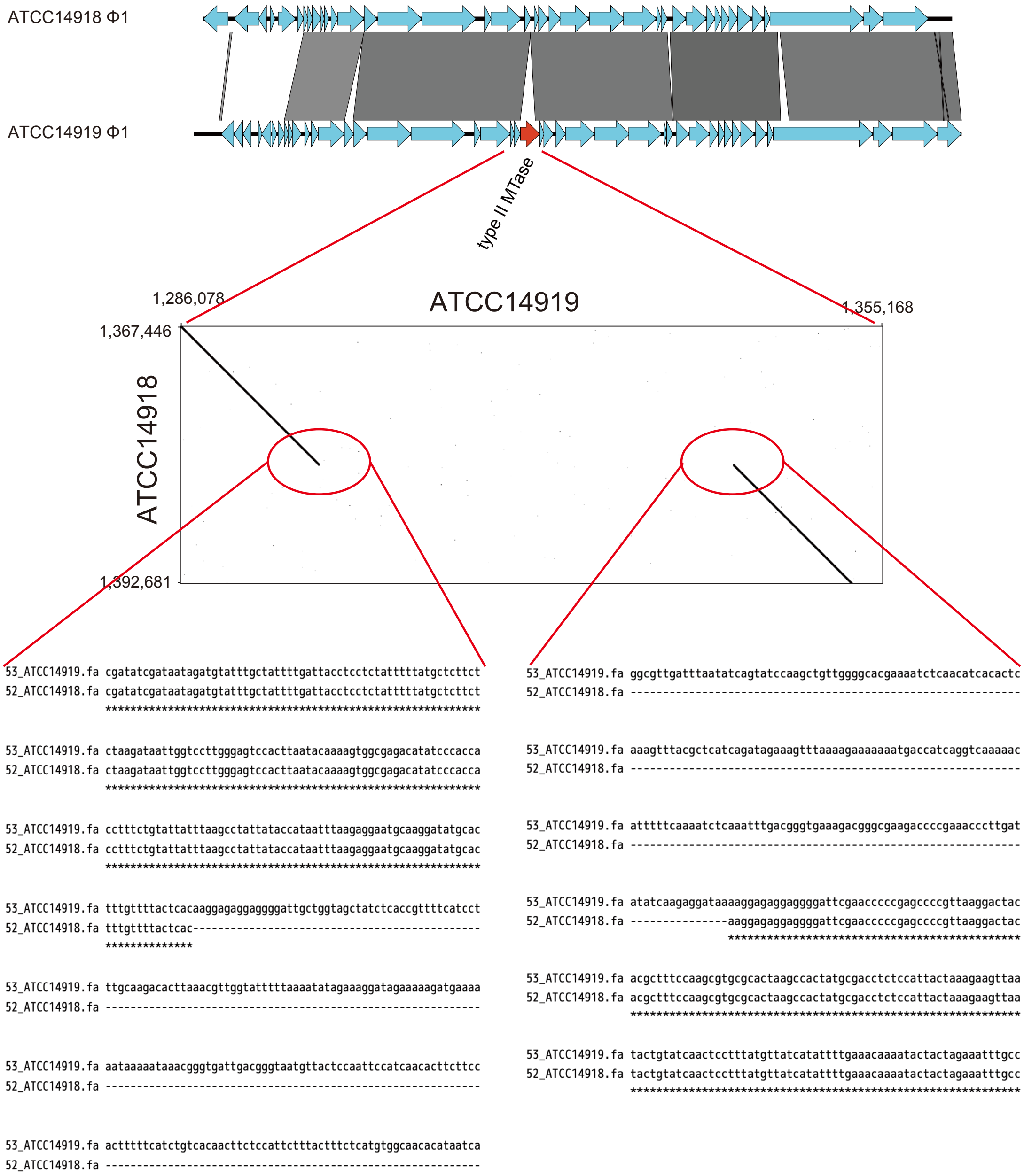


**Supplementary Figure S5: Sequence alignment of type II MTase coding region of the prophage on ATCC14919 and ATCC14918.**

### Supplementary Table S1. Strains downloaded from PATRIC.

| #Organism/Name | Strain | CladeID | BioSample | BioProject | Group | SubGroup | Assembly | Size (Mb) | GC% | Replicons | WGS | Scaffolds | Genes | Proteins | Release Date | Modify Date | Level | RefSeq FTP | GenBank FTP |
|---|---|---|---|---|---|---|---|---|---|---|---|---|---|---|---|---|---|---|---|
| Streptococcus pyogenes M1 GAS | SF370 | 20126 | SAMN02604089 | PRJNA269 | Terrabacteria | Firmicutes | GCA_000006785.2 | 1.85243 | 38.5 | chromosome:NC_002737.2/AE004092.2 | - | 1 | 1801 | 1693 | 2001/4/10 | 2016/4/18 | Complete Genome | ftp://ftp.ncbi.nlm.nih.gov/genomes/all/GCF_000006785.2_ASM678v2 | ftp://ftp.ncbi.nlm.nih.gov/genomes/all/GCA_000006785.2_ASM678v2 |
| Streptococcus pyogenes MGAS8232 | MGAS8232 | 20126 | SAMN02603495 | PRJNA286 | Terrabacteria | Firmicutes | GCA_000007285.1 | 1.89502 | 38.5 | chromosome:NC_003485.1/AE009949.1 | - | 1 | 1920 | 1794 | 2002/1/31 | 2015/8/12 | Complete Genome | ftp://ftp.ncbi.nlm.nih.gov/genomes/all/GCF_000007285.1_ASM728v1 | ftp://ftp.ncbi.nlm.nih.gov/genomes/all/GCA_000007285.1_ASM728v1 |
| Streptococcus pyogenes MGAS315 | MGAS315 | 20126 | SAMN02603496 | PRJNA311 | Terrabacteria | Firmicutes | GCA_000007425.1 | 1.90052 | 38.6 | chromosome:NC_004070.1/AE014074.1 | - | 1 | 1923 | 1804 | 2002/7/18 | 2016/3/2 | Complete Genome | ftp://ftp.ncbi.nlm.nih.gov/genomes/all/GCF_000007425.1_ASM742v1 | ftp://ftp.ncbi.nlm.nih.gov/genomes/all/GCA_000007425.1_ASM742v1 |
| Streptococcus pyogenes str. Manfredo | Manfredo | 20126 | SAMEA1705956 | PRJNA270 | Terrabacteria | Firmicutes | GCA_000009385.1 | 1.84127 | 38.6 | chromosome:NC_009332.1/AM295007.1 | - | 1 | 1841 | 1720 | 2007/2/5 | 2015/8/12 | Complete Genome | ftp://ftp.ncbi.nlm.nih.gov/genomes/all/GCF_000009385.1_ASM938v1 | ftp://ftp.ncbi.nlm.nih.gov/genomes/all/GCA_000009385.1_ASM938v1 |
| Streptococcus pyogenes SSI-1 | SSI-1 | 20126 | - | PRJNA301 | Terrabacteria | Firmicutes | GCA_000011285.1 | 1.89427 | 38.6 | chromosome:NC_004606.1/BA000034.2 | - | 1 | 1912 | 1808 | 2003/3/4 | 2015/8/12 | Complete Genome | ftp://ftp.ncbi.nlm.nih.gov/genomes/all/GCF_000011285.1_ASM1128v1 | ftp://ftp.ncbi.nlm.nih.gov/genomes/all/GCA_000011285.1_ASM1128v1 |
| Streptococcus pyogenes MGAS10394 | MGAS10394 | 20126 | SAMN02603497 | PRJNA12469 | Terrabacteria | Firmicutes | GCA_000011665.1 | 1.89988 | 38.7 | chromosome:NC_006086.1/CP000003.1 | - | 1 | 1875 | 1738 | 2004/8/3 | 2015/8/12 | Complete Genome | ftp://ftp.ncbi.nlm.nih.gov/genomes/all/GCF_000011665.1_ASM1166v1 | ftp://ftp.ncbi.nlm.nih.gov/genomes/all/GCA_000011665.1_ASM1166v1 |
| Streptococcus pyogenes MGAS5005 | MGAS5005 | 20126 | SAMN02603500 | PRJNA13888 | Terrabacteria | Firmicutes | GCA_000011765.2 | 1.83856 | 38.5 | chromosome:NC_007297.2/CP000017.2 | - | 1 | 1835 | 1724 | 2005/8/5 | 2016/3/2 | Complete Genome | ftp://ftp.ncbi.nlm.nih.gov/genomes/all/GCF_000011765.3_ASM1176v2 | ftp://ftp.ncbi.nlm.nih.gov/genomes/all/GCA_000011765.2_ASM1176v2 |
| Streptococcus pyogenes MGAS6180 | MGAS6180 | 20126 | SAMN02603499 | PRJNA13887 | Terrabacteria | Firmicutes | GCA_000012165.1 | 1.89757 | 38.4 | chromosome:NC_007296.1/CP000056.1 | - | 1 | 1870 | 1748 | 2005/8/4 | 2016/3/2 | Complete Genome | ftp://ftp.ncbi.nlm.nih.gov/genomes/all/GCF_000012165.1_ASM1216v1 | ftp://ftp.ncbi.nlm.nih.gov/genomes/all/GCA_000012165.1_ASM1216v1 |
| Streptococcus pyogenes MGAS9429 | MGAS9429 | 20126 | SAMN02603501 | PRJNA16363 | Terrabacteria | Firmicutes | GCA_000013485.1 | 1.83647 | 38.5 | chromosome:NC_008021.1/CP000259.1 | - | 1 | 1800 | 1680 | 2006/5/4 | 2016/3/2 | Complete Genome | ftp://ftp.ncbi.nlm.nih.gov/genomes/all/GCF_000013485.1_ASM1348v1 | ftp://ftp.ncbi.nlm.nih.gov/genomes/all/GCA_000013485.1_ASM1348v1 |
| Streptococcus pyogenes MGAS10270 | MGAS10270 | 20126 | SAMN02603502 | PRJNA16364 | Terrabacteria | Firmicutes | GCA_000013505.1 | 1.92825 | 38.4 | chromosome:NC_008022.1/CP000260.1 | - | 1 | 1923 | 1778 | 2006/5/4 | 2016/3/2 | Complete Genome | ftp://ftp.ncbi.nlm.nih.gov/genomes/all/GCF_000013505.1_ASM1350v1 | ftp://ftp.ncbi.nlm.nih.gov/genomes/all/GCA_000013505.1_ASM1350v1 |
| Streptococcus pyogenes MGAS2096 | MGAS2096 | 20126 | SAMN02603503 | PRJNA16365 | Terrabacteria | Firmicutes | GCA_000013525.1 | 1.86035 | 38.7 | chromosome:NC_008023.1/CP000261.1 | - | 1 | 1809 | 1609 | 2006/5/4 | 2016/3/2 | Complete Genome | ftp://ftp.ncbi.nlm.nih.gov/genomes/all/GCF_000013525.1_ASM1352v1 | ftp://ftp.ncbi.nlm.nih.gov/genomes/all/GCA_000013525.1_ASM1352v1 |
| Streptococcus pyogenes MGAS10750 | MGAS10750 | 20126 | SAMN02603504 | PRJNA16366 | Terrabacteria | Firmicutes | GCA_000013545.1 | 1.93711 | 38.3 | chromosome:NC_008024.1/CP000262.1 | - | 1 | 1924 | 1770 | 2006/5/4 | 2016/3/2 | Complete Genome | ftp://ftp.ncbi.nlm.nih.gov/genomes/all/GCF_000013545.1_ASM1354v1 | ftp://ftp.ncbi.nlm.nih.gov/genomes/all/GCA_000013545.1_ASM1354v1 |
| Streptococcus pyogenes NZ131 | NZ131; ATCC BAA-1633 | 20126 | SAMN02604236 | PRJNA20707 | Terrabacteria | Firmicutes | GCA_000018125.1 | 1.81578 | 38.6 | chromosome:NC_011375.1/CP000829.1 | - | 1 | 1791 | 1626 | 2008/10/16 | 2016/3/2 | Complete Genome | ftp://ftp.ncbi.nlm.nih.gov/genomes/all/GCF_000018125.1_ASM1812v1 | ftp://ftp.ncbi.nlm.nih.gov/genomes/all/GCA_000018125.1_ASM1812v1 |
| Streptococcus pyogenes Alab49 | Alab49 | 20126 | SAMN02603353 | PRJNA73399 | Terrabacteria | Firmicutes | GCA_000230295.1 | 1.82731 | 38.6 | chromosome:NC_017596.1/CP003068.1 | - | 1 | 1816 | 1699 | 2011/10/7 | 2015/8/13 | Complete Genome | ftp://ftp.ncbi.nlm.nih.gov/genomes/all/GCF_000230295.1_ASM23029v1 | ftp://ftp.ncbi.nlm.nih.gov/genomes/all/GCA_000230295.1_ASM23029v1 |
| Streptococcus pyogenes MGAS15252 | MGAS15252 | 20126 | SAMN02603944 | PRJNA49669 | Terrabacteria | Firmicutes | GCA_000250905.1 | 1.75083 | 38.5 | chromosome:NC_017040.1/CP003116.1 | - | 1 | 1693 | 1587 | 2012/3/1 | 2015/8/13 | Complete Genome | ftp://ftp.ncbi.nlm.nih.gov/genomes/all/GCF_000250905.1_ASM25090v1 | ftp://ftp.ncbi.nlm.nih.gov/genomes/all/GCA_000250905.1_ASM25090v1 |
| Streptococcus pyogenes MGAS1882 | MGAS1882 | 20126 | SAMN02603945 | PRJNA66081 | Terrabacteria | Firmicutes | GCA_000250925.1 | 1.78103 | 38.5 | chromosome:NC_017053.1/CP003121.1 | - | 1 | 1728 | 1624 | 2012/3/1 | 2015/8/13 | Complete Genome | ftp://ftp.ncbi.nlm.nih.gov/genomes/all/GCF_000250925.1_ASM25092v1 | ftp://ftp.ncbi.nlm.nih.gov/genomes/all/GCA_000250925.1_ASM25092v1 |
| Streptococcus pyogenes A20 | A20 | 20126 | SAMN02603640 | PRJNA175952 | Terrabacteria | Firmicutes | GCA_000307535.1 | 1.83728 | 38.5 | chromosome:NC_018936.1/CP003901.1 | - | 1 | 1833 | 1724 | 2012/10/22 | 2015/8/14 | Complete Genome | ftp://ftp.ncbi.nlm.nih.gov/genomes/all/GCF_000307535.1_ASM30753v1 | ftp://ftp.ncbi.nlm.nih.gov/genomes/all/GCA_000307535.1_ASM30753v1 |
| Streptococcus pyogenes M1 476 | M1 476 | 20126 | - | PRJDB141 | Terrabacteria | Firmicutes | GCA_000349925.2 | 1.83113 | 38.5 | chromosome 1:NC_020540.2/AP012491.2 | - | 1 | 1820 | 1721 | 2012/7/14 | 2015/8/14 | Complete Genome | ftp://ftp.ncbi.nlm.nih.gov/genomes/all/GCF_000349925.2_ASM34992v2 | ftp://ftp.ncbi.nlm.nih.gov/genomes/all/GCA_000349925.2_ASM34992v2 |
| Streptococcus pyogenes HSC5 | HSC5 | 20126 | SAMN02604317 | PRJNA205050 | Terrabacteria | Firmicutes | GCA_000422045.1 | 1.81835 | 38.5 | chromosome:NC_021807.1/CP006366.1 | - | 1 | 1806 | 1686 | 2013/7/11 | 2015/8/18 | Complete Genome | ftp://ftp.ncbi.nlm.nih.gov/genomes/all/GCF_000422045.1_ASM42204v1 | ftp://ftp.ncbi.nlm.nih.gov/genomes/all/GCA_000422045.1_ASM42204v1 |
| Streptococcus pyogenes STAB902 | STAB902 | 20126 | SAMN03081524 | PRJNA231895 | Terrabacteria | Firmicutes | GCA_000732425.1 | 1.89212 | 38.5 | chromosome:NZ_CP007041.1/CP007041.1 | - | 1 | 1912 | 1804 | 2014/7/22 | 2015/8/18 | Complete Genome | ftp://ftp.ncbi.nlm.nih.gov/genomes/all/GCF_000732425.1_ASM73242v1 | ftp://ftp.ncbi.nlm.nih.gov/genomes/all/GCA_000732425.1_ASM73242v1 |
| Streptococcus pyogenes | ATCC 19615 | 20126 | SAMN02905033 | PRJNA244349 | Terrabacteria | Firmicutes | GCA_000743015.1 | 1.8448 | 38.5 | chromosome:NZ_CP008926.1/CP008926.1 | - | 1 | 1835 | 1702 | 2014/8/22 | 2015/8/16 | Complete Genome | ftp://ftp.ncbi.nlm.nih.gov/genomes/all/GCF_000743015.1_ASM74301v1 | ftp://ftp.ncbi.nlm.nih.gov/genomes/all/GCA_000743015.1_ASM74301v1 |
| Streptococcus pyogenes | M23ND | 20126 | SAMN02800743 | PRJNA248257 | Terrabacteria | Firmicutes | GCA_000756485.1 | 1.84648 | 38.6 | chromosome:NZ_CP008695.1/CP008695.1 | - | 1 | 1839 | 1719 | 2014/9/22 | 2015/8/17 | Complete Genome | ftp://ftp.ncbi.nlm.nih.gov/genomes/all/GCF_000756485.1_ASM75648v1 | ftp://ftp.ncbi.nlm.nih.gov/genomes/all/GCA_000756485.1_ASM75648v1 |
| Streptococcus pyogenes | 7F7 | 20126 | SAMN02644974 | PRJNA238516 | Terrabacteria | Firmicutes | GCA_000767505.1 | 1.70979 | 38.6 | chromosome:NZ_CP007240.1/CP007240.1 | - | 1 | 1653 | 1546 | 2014/10/20 | 2015/8/17 | Complete Genome | ftp://ftp.ncbi.nlm.nih.gov/genomes/all/GCF_000767505.1_ASM76750v1 | ftp://ftp.ncbi.nlm.nih.gov/genomes/all/GCA_000767505.1_ASM76750v1 |
| Streptococcus pyogenes | HKU360 | 20126 | SAMN02980885 | PRJNA257934 | Terrabacteria | Firmicutes | GCA_000772185.1 | 1.94454 | 38.5 | chromosome:NZ_CP009612.1/CP009612.1 | - | 1 | 1916 | 1809 | 2014/11/5 | 2015/8/17 | Complete Genome | ftp://ftp.ncbi.nlm.nih.gov/genomes/all/GCF_000772185.1_ASM77218v1 | ftp://ftp.ncbi.nlm.nih.gov/genomes/all/GCA_000772185.1_ASM77218v1 |
| Streptococcus pyogenes | 10 | 20126 | SAMN02649402 | PRJNA238987 | Terrabacteria | Firmicutes | GCA_000772245.1 | 1.79615 | 38.5 | chromosome:NZ_CP007241.1/CP007241.1 | - | 1 | 1741 | 1614 | 2014/11/5 | 2015/8/17 | Complete Genome | ftp://ftp.ncbi.nlm.nih.gov/genomes/all/GCF_000772245.1_ASM77224v1 | ftp://ftp.ncbi.nlm.nih.gov/genomes/all/GCA_000772245.1_ASM77224v1 |
| Streptococcus pyogenes | AP1 | 20126 | SAMN02716635 | PRJNA242701 | Terrabacteria | Firmicutes | GCA_000993765.1 | 1.90829 | 38.5 | chromosome:NZ_CP007537.1/CP007537.1 | - | 1 | 1930 | 1830 | 2015/5/7 | 2015/8/20 | Complete Genome | ftp://ftp.ncbi.nlm.nih.gov/genomes/all/GCF_000993765.1_ASM99376v1 | ftp://ftp.ncbi.nlm.nih.gov/genomes/all/GCA_000993765.1_ASM99376v1 |
| Streptococcus pyogenes JRS4 | JRS4 | 20126 | SAMN03610013 | PRJNA283215 | Terrabacteria | Firmicutes | GCA_001014285.1 | 1.81197 | 38.6 | chromosome:NZ_CP011414.1/CP011414.1 | - | 1 | 1788 | 1667 | 2015/5/28 | 2015/8/20 | Complete Genome | ftp://ftp.ncbi.nlm.nih.gov/genomes/all/GCF_001014285.1_ASM101428v1 | ftp://ftp.ncbi.nlm.nih.gov/genomes/all/GCA_001014285.1_ASM101428v1 |
| Streptococcus pyogenes | D471 | 20126 | SAMN03610015 | PRJNA283214 | Terrabacteria | Firmicutes | GCA_001014305.1 | 1.81197 | 38.6 | chromosome:NZ_CP011415.1/CP011415.1 | - | 1 | 1788 | 1667 | 2015/5/28 | 2015/8/20 | Complete Genome | ftp://ftp.ncbi.nlm.nih.gov/genomes/all/GCF_001014305.1_ASM101430v1 | ftp://ftp.ncbi.nlm.nih.gov/genomes/all/GCA_001014305.1_ASM101430v1 |
| Streptococcus pyogenes | NGAS743 | 20126 | SAMN02715744 | PRJNA243328 | Terrabacteria | Firmicutes | GCA_001019635.1 | 1.91555 | 38.5 | chromosome:NZ_CP007560.1/CP007560.1 | - | 1 | 1924 | 1805 | 2015/6/3 | 2015/8/20 | Complete Genome | ftp://ftp.ncbi.nlm.nih.gov/genomes/all/GCF_001019635.1_ASM101963v1 | ftp://ftp.ncbi.nlm.nih.gov/genomes/all/GCA_001019635.1_ASM101963v1 |
| Streptococcus pyogenes | NGAS596 | 20126 | SAMN02715758 | PRJNA243328 | Terrabacteria | Firmicutes | GCA_001019675.1 | 1.79131 | 38.5 | chromosome:NZ_CP007561.1/CP007561.1 | - | 1 | 1739 | 1617 | 2015/6/3 | 2015/8/20 | Complete Genome | ftp://ftp.ncbi.nlm.nih.gov/genomes/all/GCF_001019675.1_ASM101967v1 | ftp://ftp.ncbi.nlm.nih.gov/genomes/all/GCA_001019675.1_ASM101967v1 |
| Streptococcus pyogenes | NGAS327 | 20126 | SAMN02715759 | PRJNA243328 | Terrabacteria | Firmicutes | GCA_001019695.1 | 1.70205 | 38.6 | chromosome:NZ_CP007562.1/CP007562.1 | - | 1 | 1644 | 1537 | 2015/6/3 | 2015/8/20 | Complete Genome | ftp://ftp.ncbi.nlm.nih.gov/genomes/all/GCF_001019695.1_ASM101969v1 | ftp://ftp.ncbi.nlm.nih.gov/genomes/all/GCA_001019695.1_ASM101969v1 |
| Streptococcus pyogenes | M28PF1 | 20126 | SAMN03733610 | PRJNA284654 | Terrabacteria | Firmicutes | GCA_001020185.2 | 1.89698 | 38.4 | chromosome:NZ_CP011535.2/CP011535.2 | - | 1 | 1874 | 1752 | 2015/6/5 | 2016/7/20 | Complete Genome | ftp://ftp.ncbi.nlm.nih.gov/genomes/all/GCF_001020185.2_ASM102018v2 | ftp://ftp.ncbi.nlm.nih.gov/genomes/all/GCA_001020185.2_ASM102018v2 |
| Streptococcus pyogenes | 5448 | 20126 | SAMN02866032 | PRJNA252999 | Terrabacteria | Firmicutes | GCA_001021955.1 | 1.82952 | 38.5 | chromosome:NZ_CP008776.1/CP008776.1 | - | 1 | 1817 | 1725 | 2015/6/8 | 2015/8/20 | Complete Genome | ftp://ftp.ncbi.nlm.nih.gov/genomes/all/GCF_001021955.1_ASM102195v1 | ftp://ftp.ncbi.nlm.nih.gov/genomes/all/GCA_001021955.1_ASM102195v1 |
| Streptococcus pyogenes | STAB10015 | 20126 | SAMN03418291 | PRJNA278400 | Terrabacteria | Firmicutes | GCA_001023495.1 | 1.95045 | 38.2 | chromosome:NZ_CP011068.1/CP011068.1 | - | 1 | 1917 | 1800 | 2015/6/9 | 2015/8/21 | Complete Genome | ftp://ftp.ncbi.nlm.nih.gov/genomes/all/GCF_001023495.1_ASM102349v1 | ftp://ftp.ncbi.nlm.nih.gov/genomes/all/GCA_001023495.1_ASM102349v1 |
| Streptococcus pyogenes | H293 | 20126 | SAMEA3865279 | PRJEB1935 | Terrabacteria | Firmicutes | GCA_001039695.2 | 1.72625 | 38.6 | chromosome:NZ_HG316453.1/HG316453.2 | - | 1 | 1661 | 1539 | 2014/5/13 | 2016/2/4 | Complete Genome | ftp://ftp.ncbi.nlm.nih.gov/genomes/all/GCF_001039695.1_emm89-1 | ftp://ftp.ncbi.nlm.nih.gov/genomes/all/GCA_001039695.2_emm89-1 |
| Streptococcus pyogenes | HKU488 | 20126 | SAMN03846812 | PRJNA289181 | Terrabacteria | Firmicutes | GCA_001051095.1 | 1.94341 | 38.5 | chromosome:NZ_CP012045.1/CP012045.1 | - | 1 | 1944 | 1832 | 2015/7/9 | 2015/8/21 | Complete Genome | ftp://ftp.ncbi.nlm.nih.gov/genomes/all/GCF_001051095.1_ASM105109v1 | ftp://ftp.ncbi.nlm.nih.gov/genomes/all/GCA_001051095.1_ASM105109v1 |
| Streptococcus pyogenes | NGAS322 | 20126 | SAMN03274509 | PRJNA243328 | Terrabacteria | Firmicutes | GCA_001267805.1 | 1.95047 | 38.3 | chromosome:NZ_CP010449.1/CP010449.1 | - | 1 | 1929 | 1809 | 2015/8/17 | 2015/8/24 | Complete Genome | ftp://ftp.ncbi.nlm.nih.gov/genomes/all/GCF_001267805.1_ASM126780v1 | ftp://ftp.ncbi.nlm.nih.gov/genomes/all/GCA_001267805.1_ASM126780v1 |
| Streptococcus pyogenes | NGAS638 | 20126 | SAMN03274510 | PRJNA243328 | Terrabacteria | Firmicutes | GCA_001267845.1 | 1.7914 | 38.6 | chromosome:NZ_CP010450.1/CP010450.1 | - | 1 | 1758 | 1650 | 2015/8/17 | 2015/8/24 | Complete Genome | ftp://ftp.ncbi.nlm.nih.gov/genomes/all/GCF_001267845.1_ASM126784v1 | ftp://ftp.ncbi.nlm.nih.gov/genomes/all/GCA_001267845.1_ASM126784v1 |
| Streptococcus pyogenes | MEW427 | 20126 | SAMN04419118 | PRJNA308988 | Terrabacteria | Firmicutes | GCA_001535505.1 | 1.81445 | 38.5 | chromosome:NZ_CP014138.1/CP014138.1 | - | 1 | 1767 | 1554 | 2016/1/25 | 2016/4/7 | Complete Genome | ftp://ftp.ncbi.nlm.nih.gov/genomes/all/GCF_001535505.1_ASM153550v1 | ftp://ftp.ncbi.nlm.nih.gov/genomes/all/GCA_001535505.1_ASM153550v1 |
| Streptococcus pyogenes | MEW123 | 20126 | SAMN04419117 | PRJNA308987 | Terrabacteria | Firmicutes | GCA_001535565.1 | 1.8787 | 38.3 | chromosome:NZ_CP014139.1/CP014139.1 | - | 1 | 1827 | 1653 | 2016/1/25 | 2016/4/7 | Complete Genome | ftp://ftp.ncbi.nlm.nih.gov/genomes/all/GCF_001535565.1_ASM153556v1 | ftp://ftp.ncbi.nlm.nih.gov/genomes/all/GCA_001535565.1_ASM153556v1 |
| Streptococcus pyogenes JRS4 | JRS4 | 20126 | - | PRJDB114 | Terrabacteria | Firmicutes | GCA_001547715.1 | 1.81112 | 38.6 | chromosome Unknown:NZ_AP012335.1/AP012335.1 | - | 1 | 1785 | 1657 | 2015/4/4 | 2016/2/4 | Complete Genome | ftp://ftp.ncbi.nlm.nih.gov/genomes/all/GCF_001547715.1_ASM154771v1 | ftp://ftp.ncbi.nlm.nih.gov/genomes/all/GCA_001547715.1_ASM154771v1 |
| Streptococcus pyogenes | MTB313 | 20126 | SAMD00000328 | PRJDB1654 | Terrabacteria | Firmicutes | GCA_001547815.1 | 1.74533 | 38.5 | chromosome Unknown:AP014572.1 | - | 1 | 1843 | 1758 | 2015/2/24 | 2015/6/4 | Complete Genome | - | ftp://ftp.ncbi.nlm.nih.gov/genomes/all/GCA_001547815.1_ASM154781v1 |
| Streptococcus pyogenes | MTB314 | 20126 | SAMD00000332 | PRJDB1668 | Terrabacteria | Firmicutes | GCA_001547835.1 | 1.74483 | 38.5 | chromosome Unknown:NZ_AP014585.1/AP014585.1 | - | 1 | 1722 | 1588 | 2015/2/24 | 2016/2/25 | Complete Genome | ftp://ftp.ncbi.nlm.nih.gov/genomes/all/GCF_001547835.1_ASM154783v1 | ftp://ftp.ncbi.nlm.nih.gov/genomes/all/GCA_001547835.1_ASM154783v1 |
| Streptococcus pyogenes | FDAARGOS_149 | 20126 | SAMN03996295 | PRJNA231221 | Terrabacteria | Firmicutes | GCA_001559175.1 | 1.83964 | 38.5 | chromosome:NZ_CP014027.1/CP014027.1 | - | 1 | 1831 | 1720 | 2016/2/11 | 2016/2/12 | Complete Genome | ftp://ftp.ncbi.nlm.nih.gov/genomes/all/GCF_001559175.1_ASM155917v1 | ftp://ftp.ncbi.nlm.nih.gov/genomes/all/GCA_001559175.1_ASM155917v1 |
| Streptococcus pyogenes | AP53 | 20126 | SAMN04334485 | PRJNA305806 | Terrabacteria | Firmicutes | GCA_001620285.1 | 1.86055 | 38.6 | chromosome:NZ_CP013672.1/CP013672.1 | - | 1 | 1862 | 1742 | 2016/4/18 | 2016/4/19 | Complete Genome | ftp://ftp.ncbi.nlm.nih.gov/genomes/all/GCF_001620285.1_ASM162028v1 | ftp://ftp.ncbi.nlm.nih.gov/genomes/all/GCA_001620285.1_ASM162028v1 |
| Streptococcus pyogenes | MGAS11027 | 20126 | SAMN03792044 | PRJNA167314 | Terrabacteria | Firmicutes | GCA_001635895.1 | 1.78687 | 38.6 | chromosome:NZ_CP013838.1/CP013838.1 | - | 1 | 1738 | 1617 | 2016/5/2 | 2016/5/16 | Complete Genome | ftp://ftp.ncbi.nlm.nih.gov/genomes/all/GCF_001635895.1_ASM163589v1 | ftp://ftp.ncbi.nlm.nih.gov/genomes/all/GCA_001635895.1_ASM163589v1 |
| Streptococcus pyogenes | MGAS23530 | 20126 | SAMN03775331 | PRJNA287028 | Terrabacteria | Firmicutes | GCA_001635935.1 | 1.70939 | 38.5 | chromosome:NZ_CP013839.1/CP013839.1 | - | 1 | 1640 | 1527 | 2016/5/2 | 2016/5/16 | Complete Genome | ftp://ftp.ncbi.nlm.nih.gov/genomes/all/GCF_001635935.1_ASM163593v1 | ftp://ftp.ncbi.nlm.nih.gov/genomes/all/GCA_001635935.1_ASM163593v1 |
| Streptococcus pyogenes | MGAS27061 | 20126 | SAMN03161558 | PRJNA266333 | Terrabacteria | Firmicutes | GCA_001635975.1 | 1.74135 | 38.5 | chromosome:NZ_CP013840.1/CP013840.1 | - | 1 | 1675 | 1565 | 2016/5/2 | 2016/5/16 | Complete Genome | ftp://ftp.ncbi.nlm.nih.gov/genomes/all/GCF_001635975.1_ASM163597v1 | ftp://ftp.ncbi.nlm.nih.gov/genomes/all/GCA_001635975.1_ASM163597v1 |
| Streptococcus pyogenes | NS53 | 20126 | SAMN04625972 | PRJNA317786 | Terrabacteria | Firmicutes | GCA_001638845.1 | 1.76512 | 38.4 | chromosome:NZ_CP015238.1/CP015238.1 | - | 1 | 1693 | 1587 | 2016/5/6 | 2016/5/12 | Complete Genome | ftp://ftp.ncbi.nlm.nih.gov/genomes/all/GCF_001638845.1_ASM163884v1 | ftp://ftp.ncbi.nlm.nih.gov/genomes/all/GCA_001638845.1_ASM163884v1 |
| Streptococcus pyogenes | STAB09014 | 20126 | SAMN03418290 | PRJNA278400 | Terrabacteria | Firmicutes | GCA_001678885.1 | 1.86249 | 38.4 | chromosome:NZ_CP011069.1/CP011069.1 | - | 1 | 1823 | 1697 | 2016/7/5 | 2016/7/6 | Complete Genome | ftp://ftp.ncbi.nlm.nih.gov/genomes/all/GCF_001678885.1_ASM167888v1 | ftp://ftp.ncbi.nlm.nih.gov/genomes/all/GCA_001678885.1_ASM167888v1 |
| Streptococcus pyogenes STAB901 | STAB901 | 20126 | SAMN03081523 | PRJNA232121 | Terrabacteria | Firmicutes | GCA_000732385.1 | 1.79561 | 38.5 | chromosome:CP007024.1 | - | 1 | 1812 | 1358 | 2014/7/22 | 2014/10/1 | Complete Genome | - | ftp://ftp.ncbi.nlm.nih.gov/genomes/all/GCA_000732385.1_ASM73238v1 |

**Supplementary Table S2. Top 5 hypermethylated regions on each strand in ATCC14918 genome.**

| No. of Methylated base | Strand | Annotation |
|---|---|---|
| 40 | + | Glutaredoxin-like protein NrdH, required for reduction of ribonucleotide reductase |
| 34 | + | Fibronectin-binding protein |
| 33 | - | Mobile element protein |
| 31 | + | SSU ribosomal protein S10p (S20e) |
| 30 | + | NADH peroxidase |
| 29 | + | DNA polymerase III beta subunit |
| 29 | - | ABC transporter, permease protein |
| 28 | - | SWF/SNF family helicase |
| 28 | - | Ribonuclease Z |
| 28 | - | DNA mismatch repair protein MutS |

**Supplementary Table S3. BLASTN outputs of spacer sequences against the "gold standards" dataset of REBASE (accessed October 2016)**

| query_name | subject_name | query_length | subject_length | score | E-value | identity | identity_per | gap | gap_per | query_start | query_end | coverage | subject_start | subject_end | subject_strand | subject_index |
|---|---|---|---|---|---|---|---|---|---|---|---|---|---|---|---|---|
| 10_MGAS2096.crt.1 | NOHIT | 38 | | | | | | | | | | | | | | |
| 10_MGAS2096.crt.2 | NOHIT | 37 | | | | | | | | | | | | | | |
| 10_MGAS2096.crt.3 | NOHIT | 0 | | | | | | | | | | | | | | |
| 10_MGAS2096.crt.4 | NOHIT | 33 | | | | | | | | | | | | | | |
| 10_MGAS2096.crt.5 | NOHIT | 35 | | | | | | | | | | | | | | |
| 10_MGAS2096.crt.6 | NOHIT | 35 | | | | | | | | | | | | | | |
| 10_MGAS2096.crt.7 | NOHIT | 35 | | | | | | | | | | | | | | |
| 10_MGAS2096.crt.8 | NOHIT | 36 | | | | | | | | | | | | | | |
| 10_MGAS2096.crt.9 | NOHIT | 34 | | | | | | | | | | | | | | |
| 11_MGAS10750.crt.1 | NOHIT | 33 | | | | | | | | | | | | | | |
| 11_MGAS10750.crt.2 | NOHIT | 34 | | | | | | | | | | | | | | |
| 11_MGAS10750.crt.3 | NOHIT | 35 | | | | | | | | | | | | | | |
| 11_MGAS10750.crt.4 | NOHIT | 33 | | | | | | | | | | | | | | |
| 11_MGAS10750.crt.5 | NOHIT | 34 | | | | | | | | | | | | | | |
| 13_NZ131.crt.1 | NOHIT | 30 | | | | | | | | | | | | | | |
| 13_NZ131.crt.2 | NOHIT | 31 | | | | | | | | | | | | | | |
| 13_NZ131.crt.3 | NOHIT | 30 | | | | | | | | | | | | | | |
| 13_NZ131.crt.4 | NOHIT | 30 | | | | | | | | | | | | | | |
| 13_NZ131.crt.5 | NOHIT | 0 | | | | | | | | | | | | | | |
| 13_NZ131.crt.6 | NOHIT | 35 | | | | | | | | | | | | | | |
| 13_NZ131.crt.7 | NOHIT | 35 | | | | | | | | | | | | | | |
| 13_NZ131.crt.8 | NOHIT | 35 | | | | | | | | | | | | | | |
| 13_NZ131.crt.9 | NOHIT | 35 | | | | | | | | | | | | | | |
| 13_NZ131.crt.10 | NOHIT | 33 | | | | | | | | | | | | | | |
| 15_MGAS15252.crt.1 | NOHIT | 37 | | | | | | | | | | | | | | |
| 15_MGAS15252.crt.2 | NOHIT | 37 | | | | | | | | | | | | | | |
| 16_MGAS1882.crt.1 | NOHIT | 30 | | | | | | | | | | | | | | |
| 16_MGAS1882.crt.2 | NOHIT | 30 | | | | | | | | | | | | | | |
| 16_MGAS1882.crt.3 | NOHIT | 30 | | | | | | | | | | | | | | |
| 17_M1_476.crt.1 | NOHIT | 30 | | | | | | | | | | | | | | |
| 17_M1_476.crt.2 | NOHIT | 30 | | | | | | | | | | | | | | |
| 17_M1_476.crt.3 | NOHIT | 30 | | | | | | | | | | | | | | |
| 17_M1_476.crt.4 | NOHIT | 0 | | | | | | | | | | | | | | |
| 17_M1_476.crt.5 | NOHIT | 33 | | | | | | | | | | | | | | |
| 17_M1_476.crt.6 | NOHIT | 34 | | | | | | | | | | | | | | |
| 17_M1_476.crt.7 | NOHIT | 33 | | | | | | | | | | | | | | |
| 17_M1_476.crt.8 | NOHIT | 35 | | | | | | | | | | | | | | |
| 18_A20.crt.1 | NOHIT | 30 | | | | | | | | | | | | | | |
| 18_A20.crt.2 | NOHIT | 30 | | | | | | | | | | | | | | |
| 18_A20.crt.3 | NOHIT | 30 | | | | | | | | | | | | | | |
| 18_A20.crt.4 | NOHIT | 0 | | | | | | | | | | | | | | |
| 18_A20.crt.5 | NOHIT | 33 | | | | | | | | | | | | | | |
| 18_A20.crt.6 | NOHIT | 34 | | | | | | | | | | | | | | |
| 18_A20.crt.7 | NOHIT | 33 | | | | | | | | | | | | | | |
| 18_A20.crt.8 | NOHIT | 35 | | | | | | | | | | | | | | |
| 1_SF370.crt.1 | NOHIT | 30 | | | | | | | | | | | | | | |
| 1_SF370.crt.2 | NOHIT | 30 | | | | | | | | | | | | | | |
| 1_SF370.crt.3 | NOHIT | 30 | | | | | | | | | | | | | | |
| 1_SF370.crt.4 | NOHIT | 30 | | | | | | | | | | | | | | |
| 1_SF370.crt.5 | NOHIT | 30 | | | | | | | | | | | | | | |
| 1_SF370.crt.6 | NOHIT | 30 | | | | | | | | | | | | | | |
| 1_SF370.crt.7 | NOHIT | 0 | | | | | | | | | | | | | | |
| 1_SF370.crt.8 | NOHIT | 33 | | | | | | | | | | | | | | |
| 1_SF370.crt.9 | NOHIT | 32 | | | | | | | | | | | | | | |
| 1_SF370.crt.10 | NOHIT | 34 | | | | | | | | | | | | | | |
| 20_H293.crt.1 | NOHIT | 30 | | | | | | | | | | | | | | |
| 20_H293.crt.2 | NOHIT | 30 | | | | | | | | | | | | | | |
| 20_H293.crt.3 | NOHIT | 30 | | | | | | | | | | | | | | |
| 20_H293.crt.4 | NOHIT | 30 | | | | | | | | | | | | | | |
| 20_H293.crt.5 | NOHIT | 30 | | | | | | | | | | | | | | |
| 20_H293.crt.6 | NOHIT | 0 | | | | | | | | | | | | | | |
| 20_H293.crt.7 | NOHIT | 19 | | | | | | | | | | | | | | |
| 20_H293.crt.8 | NOHIT | 28 | | | | | | | | | | | | | | |
| 20_H293.crt.9 | NOHIT | 19 | | | | | | | | | | | | | | |
| 20_H293.crt.10 | NOHIT | 19 | | | | | | | | | | | | | | |
| 20_H293.crt.11 | NOHIT | 0 | | | | | | | | | | | | | | |
| 20_H293.crt.12 | NOHIT | 34 | | | | | | | | | | | | | | |
| 20_H293.crt.13 | NOHIT | 34 | | | | | | | | | | | | | | |
| 20_H293.crt.14 | NOHIT | 37 | | | | | | | | | | | | | | |
| 20_H293.crt.15 | NOHIT | 35 | | | | | | | | | | | | | | |
| 20_H293.crt.16 | NOHIT | 34 | | | | | | | | | | | | | | |
| 20_H293.crt.17 | NOHIT | 35 | | | | | | | | | | | | | | |
| 20_H293.crt.18 | NOHIT | 33 | | | | | | | | | | | | | | |
| 20_H293.crt.19 | NOHIT | 34 | | | | | | | | | | | | | | |
| 20_H293.crt.20 | NOHIT | 37 | | | | | | | | | | | | | | |
| 20_H293.crt.21 | NOHIT | 33 | | | | | | | | | | | | | | |
| 20_H293.crt.22 | NOHIT | 34 | | | | | | | | | | | | | | |
| 20_H293.crt.23 | NOHIT | 33 | | | | | | | | | | | | | | |
| 20_H293.crt.24 | NOHIT | 34 | | | | | | | | | | | | | | |
| 22_STAB901.crt.1 | NOHIT | 30 | | | | | | | | | | | | | | |
| 22_STAB901.crt.2 | NOHIT | 30 | | | | | | | | | | | | | | |
| 22_STAB901.crt.3 | NOHIT | 30 | | | | | | | | | | | | | | |
| 22_STAB901.crt.4 | NOHIT | 30 | | | | | | | | | | | | | | |
| 22_STAB901.crt.5 | NOHIT | 0 | | | | | | | | | | | | | | |
| 22_STAB901.crt.6 | NOHIT | 29 | | | | | | | | | | | | | | |
| 22_STAB901.crt.7 | NOHIT | 31 | | | | | | | | | | | | | | |
| 22_STAB901.crt.8 | NOHIT | 32 | | | | | | | | | | | | | | |
| 25_7F7.crt.1 | NOHIT | 30 | | | | | | | | | | | | | | |
| 25_7F7.crt.2 | NOHIT | 30 | | | | | | | | | | | | | | |
| 25_7F7.crt.3 | NOHIT | 30 | | | | | | | | | | | | | | |
| 25_7F7.crt.4 | NOHIT | 30 | | | | | | | | | | | | | | |
| 25_7F7.crt.5 | NOHIT | 30 | | | | | | | | | | | | | | |
| 25_7F7.crt.6 | NOHIT | 30 | | | | | | | | | | | | | | |
| 25_7F7.crt.7 | NOHIT | 30 | | | | | | | | | | | | | | |
| 25_7F7.crt.8 | NOHIT | 30 | | | | | | | | | | | | | | |
| 26_HKU360.crt.1 | NOHIT | 38 | | | | | | | | | | | | | | |
| 26_HKU360.crt.2 | NOHIT | 37 | | | | | | | | | | | | | | |
| 26_HKU360.crt.3 | NOHIT | 0 | | | | | | | | | | | | | | |
| 26_HKU360.crt.4 | NOHIT | 32 | | | | | | | | | | | | | | |
| 26_HKU360.crt.5 | NOHIT | 34 | | | | | | | | | | | | | | |
| 26_HKU360.crt.6 | NOHIT | 34 | | | | | | | | | | | | | | |
| 26_HKU360.crt.7 | NOHIT | 33 | | | | | | | | | | | | | | |
| 27_I0.crt.1 | NOHIT | 30 | | | | | | | | | | | | | | |
| 27_I0.crt.2 | NOHIT | 30 | | | | | | | | | | | | | | |
| 27_I0.crt.3 | NOHIT | 30 | | | | | | | | | | | | | | |
| 27_I0.crt.4 | NOHIT | 30 | | | | | | | | | | | | | | |
| 27_I0.crt.5 | NOHIT | 30 | | | | | | | | | | | | | | |
| 27_I0.crt.6 | NOHIT | 30 | | | | | | | | | | | | | | |
| 27_I0.crt.7 | NOHIT | 30 | | | | | | | | | | | | | | |
| 27_I0.crt.8 | NOHIT | 30 | | | | | | | | | | | | | | |
| 27_I0.crt.9 | NOHIT | 30 | | | | | | | | | | | | | | |
| 27_I0.crt.10 | NOHIT | 30 | | | | | | | | | | | | | | |
| 27_I0.crt.11 | NOHIT | 0 | | | | | | | | | | | | | | |
| 27_I0.crt.12 | NOHIT | 29 | | | | | | | | | | | | | | |
| 27_I0.crt.13 | NOHIT | 31 | | | | | | | | | | | | | | |
| 27_I0.crt.14 | NOHIT | 32 | | | | | | | | | | | | | | |
| 28_MTB313.crt.1 | NOHIT | 29 | | | | | | | | | | | | | | |
| 28_MTB313.crt.2 | NOHIT | 31 | | | | | | | | | | | | | | |
| 28_MTB313.crt.3 | NOHIT | 29 | | | | | | | | | | | | | | |
| 28_MTB313.crt.4 | NOHIT | 0 | | | | | | | | | | | | | | |
| 28_MTB313.crt.5 | NOHIT | 33 | | | | | | | | | | | | | | |
| 28_MTB313.crt.6 | NOHIT | 34 | | | | | | | | | | | | | | |
| 28_MTB313.crt.7 | NOHIT | 33 | | | | | | | | | | | | | | |
| 28_MTB313.crt.8 | NOHIT | 35 | | | | | | | | | | | | | | |
| 29_MTB314.crt.1 | NOHIT | 30 | | | | | | | | | | | | | | |
| 29_MTB314.crt.2 | NOHIT | 30 | | | | | | | | | | | | | | |
| 29_MTB314.crt.3 | NOHIT | 30 | | | | | | | | | | | | | | |
| 29_MTB314.crt.4 | NOHIT | 0 | | | | | | | | | | | | | | |
| 29_MTB314.crt.5 | NOHIT | 33 | | | | | | | | | | | | | | |
| 29_MTB314.crt.6 | NOHIT | 33 | | | | | | | | | | | | | | |
| 29_MTB314.crt.7 | NOHIT | 33 | | | | | | | | | | | | | | |
| 29_MTB314.crt.8 | NOHIT | 35 | | | | | | | | | | | | | | |
| 31_AP1.crt.1 | NOHIT | 35 | | | | | | | | | | | | | | |
| 31_AP1.crt.2 | NOHIT | 33 | | | | | | | | | | | | | | |
| 31_AP1.crt.3 | NOHIT | 34 | | | | | | | | | | | | | | |
| 31_AP1.crt.4 | NOHIT | 33 | | | | | | | | | | | | | | |
| 35_NGAS596.crt.1 | NOHIT | 32 | | | | | | | | | | | | | | |
| 35_NGAS596.crt.2 | NOHIT | 33 | | | | | | | | | | | | | | |
| 35_NGAS596.crt.3 | NOHIT | 34 | | | | | | | | | | | | | | |
| 35_NGAS596.crt.4 | NOHIT | 33 | | | | | | | | | | | | | | |
| 35_NGAS596.crt.5 | NOHIT | 33 | | | | | | | | | | | | | | |
| 36_NGAS327.crt.1 | NOHIT | 30 | | | | | | | | | | | | | | |
| 36_NGAS327.crt.2 | NOHIT | 30 | | | | | | | | | | | | | | |
| 36_NGAS327.crt.3 | NOHIT | 30 | | | | | | | | | | | | | | |
| 36_NGAS327.crt.4 | NOHIT | 30 | | | | | | | | | | | | | | |
| 36_NGAS327.crt.5 | NOHIT | 30 | | | | | | | | | | | | | | |
| 36_NGAS327.crt.6 | NOHIT | 30 | | | | | | | | | | | | | | |
| 36_NGAS327.crt.7 | NOHIT | 30 | | | | | | | | | | | | | | |
| 36_NGAS327.crt.8 | NOHIT | 30 | | | | | | | | | | | | | | |
| 37_M28PF1.crt.1 | NOHIT | 35 | | | | | | | | | | | | | | |
| 37_M28PF1.crt.2 | NOHIT | 34 | | | | | | | | | | | | | | |
| 37_M28PF1.crt.3 | NOHIT | 0 | | | | | | | | | | | | | | |
| 37_M28PF1.crt.4 | NOHIT | 30 | | | | | | | | | | | | | | |
| 37_M28PF1.crt.5 | NOHIT | 30 | | | | | | | | | | | | | | |
| 37_M28PF1.crt.6 | NOHIT | 30 | | | | | | | | | | | | | | |
| 37_M28PF1.crt.7 | NOHIT | 30 | | | | | | | | | | | | | | |
| 38_5448.crt.1 | NOHIT | 35 | | | | | | | | | | | | | | |
| 38_5448.crt.2 | NOHIT | 33 | | | | | | | | | | | | | | |
| 38_5448.crt.3 | NOHIT | 34 | | | | | | | | | | | | | | |
| 38_5448.crt.4 | NOHIT | 33 | | | | | | | | | | | | | | |
| 38_5448.crt.5 | NOHIT | 0 | | | | | | | | | | | | | | |
| 38_5448.crt.6 | NOHIT | 30 | | | | | | | | | | | | | | |
| 38_5448.crt.7 | NOHIT | 30 | | | | | | | | | | | | | | |
| 38_5448.crt.8 | NOHIT | 30 | | | | | | | | | | | | | | |
| 39_STAB10015.crt.1 | NOHIT | 30 | | | | | | | | | | | | | | |
| 39_STAB10015.crt.2 | NOHIT | 30 | | | | | | | | | | | | | | |
| 39_STAB10015.crt.3 | NOHIT | 30 | | | | | | | | | | | | | | |
| 39_STAB10015.crt.4 | NOHIT | 30 | | | | | | | | | | | | | | |
| 39_STAB10015.crt.5 | NOHIT | 0 | | | | | | | | | | | | | | |
| 39_STAB10015.crt.6 | NOHIT | 34 | | | | | | | | | | | | | | |
| 39_STAB10015.crt.7 | NOHIT | 35 | | | | | | | | | | | | | | |
| 40_HKU488.crt.1 | NOHIT | 30 | | | | | | | | | | | | | | |
| 40_HKU488.crt.2 | NOHIT | 30 | | | | | | | | | | | | | | |
| 40_HKU488.crt.3 | NOHIT | 30 | | | | | | | | | | | | | | |
| 40_HKU488.crt.4 | NOHIT | 0 | | | | | | | | | | | | | | |
| 40_HKU488.crt.5 | NOHIT | 33 | | | | | | | | | | | | | | |
| 40_HKU488.crt.6 | NOHIT | 34 | | | | | | | | | | | | | | |
| 40_HKU488.crt.7 | NOHIT | 33 | | | | | | | | | | | | | | |
| 40_HKU488.crt.8 | NOHIT | 35 | | | | | | | | | | | | | | |
| 41_NGAS322.crt.1 | NOHIT | 41 | | | | | | | | | | | | | | |
| 41_NGAS322.crt.2 | NOHIT | 41 | | | | | | | | | | | | | | |
| 41_NGAS322.crt.3 | NOHIT | 41 | | | | | | | | | | | | | | |
| 41_NGAS322.crt.4 | NOHIT | 41 | | | | | | | | | | | | | | |
| 41_NGAS322.crt.5 | NOHIT | 41 | | | | | | | | | | | | | | |
| 41_NGAS322.crt.6 | NOHIT | 41 | | | | | | | | | | | | | | |
| 42_NGAS638.crt.1 | NOHIT | 29 | | | | | | | | | | | | | | |
| 42_NGAS638.crt.2 | NOHIT | 29 | | | | | | | | | | | | | | |
| 42_NGAS638.crt.3 | NOHIT | 29 | | | | | | | | | | | | | | |
| 42_NGAS638.crt.4 | NOHIT | 29 | | | | | | | | | | | | | | |
| 43_MEW427.crt.1 | NOHIT | 33 | | | | | | | | | | | | | | |
| 43_MEW427.crt.2 | NOHIT | 34 | | | | | | | | | | | | | | |
| 43_MEW427.crt.3 | NOHIT | 35 | | | | | | | | | | | | | | |
| 43_MEW427.crt.4 | NOHIT | 33 | | | | | | | | | | | | | | |
| 43_MEW427.crt.5 | NOHIT | 34 | | | | | | | | | | | | | | |
| 44_MEW123.crt.1 | NOHIT | 35 | | | | | | | | | | | | | | |
| 44_MEW123.crt.2 | NOHIT | 34 | | | | | | | | | | | | | | |
| 44_MEW123.crt.3 | NOHIT | 0 | | | | | | | | | | | | | | |
| 44_MEW123.crt.4 | NOHIT | 31 | | | | | | | | | | | | | | |
| 44_MEW123.crt.5 | NOHIT | 30 | | | | | | | | | | | | | | |
| 44_MEW123.crt.6 | NOHIT | 30 | | | | | | | | | | | | | | |
| 45_FDAARGOS_149.crt. | NOHIT | 35 | | | | | | | | | | | | | | |
| 45_FDAARGOS_149.crt. | NOHIT | 33 | | | | | | | | | | | | | | |
| 45_FDAARGOS_149.crt. | NOHIT | 34 | | | | | | | | | | | | | | |
| 45_FDAARGOS_149.crt. | NOHIT | 33 | | | | | | | | | | | | | | |
| 45_FDAARGOS_149.crt. | NOHIT | 0 | | | | | | | | | | | | | | |
| 45_FDAARGOS_149.crt. | NOHIT | 30 | | | | | | | | | | | | | | |
| 45_FDAARGOS_149.crt. | NOHIT | 30 | | | | | | | | | | | | | | |
| 45_FDAARGOS_149.crt. | NOHIT | 30 | | | | | | | | | | | | | | |
| 47_MGAS11027.crt.1 | NOHIT | 30 | | | | | | | | | | | | | | |
| 47_MGAS11027.crt.2 | NOHIT | 30 | | | | | | | | | | | | | | |
| 47_MGAS11027.crt.3 | NOHIT | 30 | | | | | | | | | | | | | | |
| 47_MGAS11027.crt.4 | NOHIT | 30 | | | | | | | | | | | | | | |
| 47_MGAS11027.crt.5 | NOHIT | 30 | | | | | | | | | | | | | | |
| 47_MGAS11027.crt.6 | NOHIT | 30 | | | | | | | | | | | | | | |
| 47_MGAS11027.crt.7 | NOHIT | 30 | | | | | | | | | | | | | | |
| 47_MGAS11027.crt.8 | NOHIT | 0 | | | | | | | | | | | | | | |
| 47_MGAS11027.crt.9 | NOHIT | 34 | | | | | | | | | | | | | | |
| 47_MGAS11027.crt.10 | NOHIT | 34 | | | | | | | | | | | | | | |
| 47_MGAS11027.crt.11 | NOHIT | 37 | | | | | | | | | | | | | | |
| 47_MGAS11027.crt.12 | NOHIT | 35 | | | | | | | | | | | | | | |
| 47_MGAS11027.crt.13 | NOHIT | 34 | | | | | | | | | | | | | | |
| 47_MGAS11027.crt.14 | NOHIT | 35 | | | | | | | | | | | | | | |
| 47_MGAS11027.crt.15 | NOHIT | 35 | | | | | | | | | | | | | | |
| 47_MGAS11027.crt.16 | NOHIT | 33 | | | | | | | | | | | | | | |
| 47_MGAS11027.crt.17 | NOHIT | 36 | | | | | | | | | | | | | | |
| 47_MGAS11027.crt.18 | NOHIT | 34 | | | | | | | | | | | | | | |
| 47_MGAS11027.crt.19 | NOHIT | 37 | | | | | | | | | | | | | | |
| 47_MGAS11027.crt.20 | NOHIT | 33 | | | | | | | | | | | | | | |
| 47_MGAS11027.crt.21 | NOHIT | 34 | | | | | | | | | | | | | | |
| 48_MGAS23530.crt.1 | NOHIT | 30 | | | | | | | | | | | | | | |
| 48_MGAS23530.crt.2 | NOHIT | 30 | | | | | | | | | | | | | | |
| 48_MGAS23530.crt.3 | NOHIT | 30 | | | | | | | | | | | | | | |
| 48_MGAS23530.crt.4 | NOHIT | 30 | | | | | | | | | | | | | | |
| 48_MGAS23530.crt.5 | NOHIT | 30 | | | | | | | | | | | | | | |
| 48_MGAS23530.crt.6 | NOHIT | 0 | | | | | | | | | | | | | | |
| 48_MGAS23530.crt.7 | NOHIT | 19 | | | | | | | | | | | | | | |
| 48_MGAS23530.crt.8 | NOHIT | 28 | | | | | | | | | | | | | | |
| 48_MGAS23530.crt.9 | NOHIT | 19 | | | | | | | | | | | | | | |
| 48_MGAS23530.crt.10 | NOHIT | 19 | | | | | | | | | | | | | | |
| 48_MGAS23530.crt.11 | NOHIT | 0 | | | | | | | | | | | | | | |
| 48_MGAS23530.crt.12 | NOHIT | 35 | | | | | | | | | | | | | | |
| 48_MGAS23530.crt.13 | NOHIT | 31 | | | | | | | | | | | | | | |
| 48_MGAS23530.crt.14 | NOHIT | 32 | | | | | | | | | | | | | | |
| 49_MGAS27061.crt.1 | NOHIT | 30 | | | | | | | | | | | | | | |
| 49_MGAS27061.crt.2 | NOHIT | 30 | | | | | | | | | | | | | | |
| 49_MGAS27061.crt.3 | NOHIT | 30 | | | | | | | | | | | | | | |
| 49_MGAS27061.crt.4 | NOHIT | 30 | | | | | | | | | | | | | | |
| 49_MGAS27061.crt.5 | NOHIT | 0 | | | | | | | | | | | | | | |
| 49_MGAS27061.crt.6 | NOHIT | 19 | | | | | | | | | | | | | | |
| 49_MGAS27061.crt.7 | NOHIT | 28 | | | | | | | | | | | | | | |
| 49_MGAS27061.crt.8 | NOHIT | 19 | | | | | | | | | | | | | | |
| 49_MGAS27061.crt.9 | NOHIT | 19 | | | | | | | | | | | | | | |
| 49_MGAS27061.crt.10 | NOHIT | 0 | | | | | | | | | | | | | | |
| 49_MGAS27061.crt.11 | NOHIT | 33 | | | | | | | | | | | | | | |
| 49_MGAS27061.crt.12 | NOHIT | 33 | | | | | | | | | | | | | | |
| 49_MGAS27061.crt.13 | NOHIT | 36 | | | | | | | | | | | | | | |
| 49_MGAS27061.crt.14 | NOHIT | 33 | | | | | | | | | | | | | | |
| 49_MGAS27061.crt.15 | NOHIT | 34 | | | | | | | | | | | | | | |
| 49_MGAS27061.crt.16 | NOHIT | 33 | | | | | | | | | | | | | | |
| 49_MGAS27061.crt.17 | NOHIT | 32 | | | | | | | | | | | | | | |
| 49_MGAS27061.crt.18 | NOHIT | 33 | | | | | | | | | | | | | | |
| 51_STAB09014.crt.1 | NOHIT | 30 | | | | | | | | | | | | | | |
| 51_STAB09014.crt.2 | NOHIT | 30 | | | | | | | | | | | | | | |
| 51_STAB09014.crt.3 | NOHIT | 30 | | | | | | | | | | | | | | |
| 51_STAB09014.crt.4 | NOHIT | 30 | | | | | | | | | | | | | | |
| 52_ATCC14918.crt.1 | NOHIT | 32 | | | | | | | | | | | | | | |
| 52_ATCC14918.crt.2 | NOHIT | 34 | | | | | | | | | | | | | | |
| 52_ATCC14918.crt.3 | NOHIT | 34 | | | | | | | | | | | | | | |
| 52_ATCC14918.crt.4 | NOHIT | 34 | | | | | | | | | | | | | | |
| 52_ATCC14918.crt.5 | NOHIT | 34 | | | | | | | | | | | | | | |
| 52_ATCC14918.crt.6 | NOHIT | 35 | | | | | | | | | | | | | | |
| 52_ATCC14918.crt.7 | NOHIT | 35 | | | | | | | | | | | | | | |
| 52_ATCC14918.crt.8 | NOHIT | 33 | | | | | | | | | | | | | | |
| 53_ATCC14919.crt.1 | NOHIT | 38 | | | | | | | | | | | | | | |
| 53_ATCC14919.crt.2 | NOHIT | 37 | | | | | | | | | | | | | | |
| 53_ATCC14919.crt.3 | NOHIT | 0 | | | | | | | | | | | | | | |
| 53_ATCC14919.crt.4 | NOHIT | 32 | | | | | | | | | | | | | | |
| 53_ATCC14919.crt.5 | NOHIT | 34 | | | | | | | | | | | | | | |
| 53_ATCC14919.crt.6 | NOHIT | 34 | | | | | | | | | | | | | | |
| 53_ATCC14919.crt.7 | NOHIT | 34 | | | | | | | | | | | | | | |
| 53_ATCC14919.crt.8 | NOHIT | 35 | | | | | | | | | | | | | | |
| 53_ATCC14919.crt.9 | NOHIT | 35 | | | | | | | | | | | | | | |
| 53_ATCC14919.crt.10 | NOHIT | 33 | | | | | | | | | | | | | | |
| 54_MGAS9429_2.crt.1 | NOHIT | 32 | | | | | | | | | | | | | | |
| 54_MGAS9429_2.crt.2 | NOHIT | 34 | | | | | | | | | | | | | | |
| 54_MGAS9429_2.crt.3 | NOHIT | 34 | | | | | | | | | | | | | | |
| 54_MGAS9429_2.crt.4 | NOHIT | 34 | | | | | | | | | | | | | | |
| 54_MGAS9429_2.crt.5 | NOHIT | 34 | | | | | | | | | | | | | | |
| 54_MGAS9429_2.crt.6 | NOHIT | 35 | | | | | | | | | | | | | | |
| 54_MGAS9429_2.crt.7 | NOHIT | 35 | | | | | | | | | | | | | | |
| 54_MGAS9429_2.crt.8 | NOHIT | 33 | | | | | | | | | | | | | | |
| 55_MGAS2096_2.crt.1 | NOHIT | 34 | | | | | | | | | | | | | | |
| 55_MGAS2096_2.crt.2 | NOHIT | 36 | | | | | | | | | | | | | | |
| 55_MGAS2096_2.crt.3 | NOHIT | 35 | | | | | | | | | | | | | | |
| 55_MGAS2096_2.crt.4 | NOHIT | 35 | | | | | | | | | | | | | | |
| 55_MGAS2096_2.crt.5 | NOHIT | 35 | | | | | | | | | | | | | | |
| 55_MGAS2096_2.crt.6 | NOHIT | 33 | | | | | | | | | | | | | | |
| 55_MGAS2096_2.crt.7 | NOHIT | 0 | | | | | | | | | | | | | | |
| 55_MGAS2096_2.crt.8 | NOHIT | 37 | | | | | | | | | | | | | | |
| 55_MGAS2096_2.crt.9 | NOHIT | 38 | | | | | | | | | | | | | | |
| 58_KIH06_201.crt.1 | NOHIT | 34 | | | | | | | | | | | | | | |
| 58_KIH06_201.crt.2 | NOHIT | 34 | | | | | | | | | | | | | | |
| 58_KIH06_201.crt.3 | NOHIT | 33 | | | | | | | | | | | | | | |
| 6_MGAS6180.crt.1 | NOHIT | 30 | | | | | | | | | | | | | | |
| 6_MGAS6180.crt.2 | NOHIT | 30 | | | | | | | | | | | | | | |
| 6_MGAS6180.crt.3 | NOHIT | 30 | | | | | | | | | | | | | | |
| 6_MGAS6180.crt.4 | NOHIT | 30 | | | | | | | | | | | | | | |
| 7_MGAS5005.crt.1 | NOHIT | 30 | | | | | | | | | | | | | | |
| 7_MGAS5005.crt.2 | NOHIT | 30 | | | | | | | | | | | | | | |
| 7_MGAS5005.crt.3 | NOHIT | 30 | | | | | | | | | | | | | | |
| 7_MGAS5005.crt.4 | NOHIT | 0 | | | | | | | | | | | | | | |
| 7_MGAS5005.crt.5 | NOHIT | 33 | | | | | | | | | | | | | | |
| 7_MGAS5005.crt.6 | NOHIT | 34 | | | | | | | | | | | | | | |
| 7_MGAS5005.crt.7 | NOHIT | 33 | | | | | | | | | | | | | | |
| 7_MGAS5005.crt.8 | NOHIT | 35 | | | | | | | | | | | | | | |
| 8_MGAS9429.crt.1 | NOHIT | 38 | | | | | | | | | | | | | | |
| 8_MGAS9429.crt.2 | NOHIT | 37 | | | | | | | | | | | | | | |
| 8_MGAS9429.crt.3 | NOHIT | 0 | | | | | | | | | | | | | | |
| 8_MGAS9429.crt.4 | NOHIT | 32 | | | | | | | | | | | | | | |
| 8_MGAS9429.crt.5 | NOHIT | 34 | | | | | | | | | | | | | | |
| 8_MGAS9429.crt.6 | NOHIT | 34 | | | | | | | | | | | | | | |
| 8_MGAS9429.crt.7 | NOHIT | 34 | | | | | | | | | | | | | | |
| 8_MGAS9429.crt.8 | NOHIT | 35 | | | | | | | | | | | | | | |
| 8_MGAS9429.crt.9 | NOHIT | 35 | | | | | | | | | | | | | | |
| 8_MGAS9429.crt.10 | NOHIT | 33 | | | | | | | | | | | | | | |
| 9_MGAS10270.crt.1 | NOHIT | 37 | | | | | | | | | | | | | | |
| 9_MGAS10270.crt.2 | NOHIT | 37 | | | | | | | | | | | | | | |
| 9_MGAS10270.crt.3 | NOHIT | 0 | | | | | | | | | | | | | | |
| 9_MGAS10270.crt.4 | NOHIT | 34 | | | | | | | | | | | | | | |
| 9_MGAS10270.crt.5 | NOHIT | 34 | | | | | | | | | | | | | | |
| 9_MGAS10270.crt.6 | NOHIT | 32 | | | | | | | | | | | | | | |

**Supplementary Table S4. BLASTN outputs of spacer sequences against 59 *S. pyogenes* genome sequences**

| query_name | subject_name | query_length | subject_length | score | E-value | identity | identity_per | gap | gap_per | query_start | query_end | coverage | subject_start | subject_end | subject_strand | subject_index |
|---|---|---|---|---|---|---|---|---|---|---|---|---|---|---|---|---|
| 10_MGAS2096.crt.1 | NOHIT | 38 | | | | | | | | | | | | | | |
| 10_MGAS2096.crt.2 | NOHIT | 37 | | | | | | | | | | | | | | |
| 10_MGAS2096.crt.3 | NOHIT | 0 | | | | | | | | | | | | | | |
| 10_MGAS2096.crt.4 | NOHIT | 33 | | | | | | | | | | | | | | |
| 10_MGAS2096.crt.5 | BNEKOFAL_00674 hypothetical protein | 35 | 285 | 65.8 | 2.00E-11 | 35 | 100% | 0 | 0% | 1 | 35 | 100% | 19 | 53 | + | 19 |
| 10_MGAS2096.crt.5 | FOAEMPCG_01040 hypothetical protein | 35 | 285 | 65.8 | 2.00E-11 | 35 | 100% | 0 | 0% | 1 | 35 | 100% | 19 | 53 | + | 19 |
| 10_MGAS2096.crt.5 | MCEIOEBL_00624 hypothetical protein | 35 | 285 | 65.8 | 2.00E-11 | 35 | 100% | 0 | 0% | 1 | 35 | 100% | 19 | 53 | + | 19 |
| 10_MGAS2096.crt.5 | PPKGMMPD_01012 hypothetical protein | 35 | 285 | 65.8 | 2.00E-11 | 35 | 100% | 0 | 0% | 1 | 35 | 100% | 19 | 53 | + | 19 |
| 10_MGAS2096.crt.5 | LLAAGEIN_01368 hypothetical protein | 35 | 285 | 65.8 | 2.00E-11 | 35 | 100% | 0 | 0% | 1 | 35 | 100% | 19 | 53 | + | 19 |
| 10_MGAS2096.crt.5 | AIKJPEBD_01573 hypothetical protein | 35 | 285 | 65.8 | 2.00E-11 | 35 | 100% | 0 | 0% | 1 | 35 | 100% | 19 | 53 | + | 19 |
| 10_MGAS2096.crt.5 | FPFNMMFM_00882 hypothetical protein | 35 | 285 | 65.8 | 2.00E-11 | 35 | 100% | 0 | 0% | 1 | 35 | 100% | 19 | 53 | + | 19 |
| 10_MGAS2096.crt.5 | IJFGMCCO_00995 hypothetical protein | 35 | 285 | 65.8 | 2.00E-11 | 35 | 100% | 0 | 0% | 1 | 35 | 100% | 19 | 53 | + | 19 |
| 10_MGAS2096.crt.5 | EJBPJIHF_01045 hypothetical protein | 35 | 285 | 65.8 | 2.00E-11 | 35 | 100% | 0 | 0% | 1 | 35 | 100% | 19 | 53 | + | 19 |
| 10_MGAS2096.crt.5 | LAKBNEPL_01011 hypothetical protein | 35 | 285 | 65.8 | 2.00E-11 | 35 | 100% | 0 | 0% | 1 | 35 | 100% | 19 | 53 | + | 19 |
| 10_MGAS2096.crt.5 | MMBCLDPD_00865 hypothetical protein | 35 | 285 | 65.8 | 2.00E-11 | 35 | 100% | 0 | 0% | 1 | 35 | 100% | 19 | 53 | + | 19 |
| 10_MGAS2096.crt.5 | PLPKHLCC_00893 hypothetical protein | 35 | 285 | 65.8 | 2.00E-11 | 35 | 100% | 0 | 0% | 1 | 35 | 100% | 19 | 53 | + | 19 |
| 10_MGAS2096.crt.5 | JNLHFPPN_01263 hypothetical protein | 35 | 285 | 65.8 | 2.00E-11 | 35 | 100% | 0 | 0% | 1 | 35 | 100% | 19 | 53 | + | 19 |
| 10_MGAS2096.crt.5 | ODIAAKBJ_01263 hypothetical protein | 35 | 285 | 65.8 | 2.00E-11 | 35 | 100% | 0 | 0% | 1 | 35 | 100% | 19 | 53 | + | 19 |
| 10_MGAS2096.crt.5 | ECKACGMC_00553 hypothetical protein | 35 | 285 | 65.8 | 2.00E-11 | 35 | 100% | 0 | 0% | 1 | 35 | 100% | 19 | 53 | + | 19 |
| 10_MGAS2096.crt.5 | DIPDIGDN_01262 hypothetical protein | 35 | 285 | 65.8 | 2.00E-11 | 35 | 100% | 0 | 0% | 1 | 35 | 100% | 19 | 53 | + | 19 |
| 10_MGAS2096.crt.5 | EFGBCCNL_00703 hypothetical protein | 35 | 285 | 65.8 | 2.00E-11 | 35 | 100% | 0 | 0% | 1 | 35 | 100% | 19 | 53 | + | 19 |
| 10_MGAS2096.crt.5 | JHELIIPC_00975 hypothetical protein | 35 | 285 | 65.8 | 2.00E-11 | 35 | 100% | 0 | 0% | 1 | 35 | 100% | 19 | 53 | + | 19 |
| 10_MGAS2096.crt.5 | GPBJDOFB_00997 hypothetical protein | 35 | 285 | 65.8 | 2.00E-11 | 35 | 100% | 0 | 0% | 1 | 35 | 100% | 19 | 53 | + | 19 |
| 10_MGAS2096.crt.5 | ACJECOPP_01021 hypothetical protein | 35 | 285 | 65.8 | 2.00E-11 | 35 | 100% | 0 | 0% | 1 | 35 | 100% | 19 | 53 | + | 19 |
| 10_MGAS2096.crt.5 | ICMALKPE_01033 hypothetical protein | 35 | 285 | 65.8 | 2.00E-11 | 35 | 100% | 0 | 0% | 1 | 35 | 100% | 19 | 53 | + | 19 |
| 10_MGAS2096.crt.5 | AJHAHHEL_01315 hypothetical protein | 35 | 285 | 65.8 | 2.00E-11 | 35 | 100% | 0 | 0% | 1 | 35 | 100% | 19 | 53 | + | 19 |
| 10_MGAS2096.crt.5 | DMLKMGFL_01490 hypothetical protein | 35 | 285 | 65.8 | 2.00E-11 | 35 | 100% | 0 | 0% | 1 | 35 | 100% | 19 | 53 | + | 19 |
| 10_MGAS2096.crt.5 | HEGDFFNC_00554 hypothetical protein | 35 | 285 | 65.8 | 2.00E-11 | 35 | 100% | 0 | 0% | 1 | 35 | 100% | 19 | 53 | + | 19 |
| 10_MGAS2096.crt.5 | IIJJCGIL_01024 hypothetical protein | 35 | 285 | 65.8 | 2.00E-11 | 35 | 100% | 0 | 0% | 1 | 35 | 100% | 19 | 53 | + | 19 |
| 10_MGAS2096.crt.5 | GLBJHDNN_01015 hypothetical protein | 35 | 285 | 65.8 | 2.00E-11 | 35 | 100% | 0 | 0% | 1 | 35 | 100% | 19 | 53 | + | 19 |
| 10_MGAS2096.crt.5 | HPLLMOOM_00975 hypothetical protein | 35 | 285 | 65.8 | 2.00E-11 | 35 | 100% | 0 | 0% | 1 | 35 | 100% | 19 | 53 | + | 19 |
| 10_MGAS2096.crt.5 | MCFNMICF_01027 hypothetical protein | 35 | 285 | 65.8 | 2.00E-11 | 35 | 100% | 0 | 0% | 1 | 35 | 100% | 19 | 53 | + | 19 |
| 10_MGAS2096.crt.5 | MONDJAPC_01581 hypothetical protein | 35 | 285 | 65.8 | 2.00E-11 | 35 | 100% | 0 | 0% | 1 | 35 | 100% | 19 | 53 | + | 19 |
| 10_MGAS2096.crt.6 | JEOJEKGO_01460 hypothetical protein | 35 | 306 | 65.8 | 2.00E-11 | 35 | 100% | 0 | 0% | 1 | 35 | 100% | 146 | 180 | + | 146 |
| 10_MGAS2096.crt.6 | DCKFNFNP_00568 hypothetical protein | 35 | 306 | 65.8 | 2.00E-11 | 35 | 100% | 0 | 0% | 1 | 35 | 100% | 146 | 180 | + | 146 |
| 10_MGAS2096.crt.6 | ECKACGMC_01462 hypothetical protein | 35 | 306 | 65.8 | 2.00E-11 | 35 | 100% | 0 | 0% | 1 | 35 | 100% | 146 | 180 | + | 146 |
| 10_MGAS2096.crt.6 | MDEAFGPO_00191 hypothetical protein | 35 | 306 | 65.8 | 2.00E-11 | 35 | 100% | 0 | 0% | 1 | 35 | 100% | 146 | 180 | + | 146 |
| 10_MGAS2096.crt.6 | HEGDFFNC_01370 hypothetical protein | 35 | 306 | 65.8 | 2.00E-11 | 35 | 100% | 0 | 0% | 1 | 35 | 100% | 146 | 180 | + | 146 |
| 10_MGAS2096.crt.6 | MONDJAPC_00627 hypothetical protein | 35 | 306 | 65.8 | 2.00E-11 | 35 | 100% | 0 | 0% | 1 | 35 | 100% | 146 | 180 | + | 146 |
| 10_MGAS2096.crt.7 | NOHIT | 35 | | | | | | | | | | | | | | |
| 10_MGAS2096.crt.8 | FOAEMPCG_00787 eukaryotic hypersensitive-induced | 36 | 828 | 67.6 | 6.00E-12 | 36 | 100% | 0 | 0% | 1 | 36 | 100% | 511 | 546 | + | 511 |
| 10_MGAS2096.crt.8 | LLAAGEIN_00659 eukaryotic hypersensitive-induced | 36 | 828 | 67.6 | 6.00E-12 | 36 | 100% | 0 | 0% | 1 | 36 | 100% | 511 | 546 | + | 511 |
| 10_MGAS2096.crt.8 | EFGBCCNL_00468 eukaryotic hypersensitive-induced | 36 | 828 | 67.6 | 6.00E-12 | 36 | 100% | 0 | 0% | 1 | 36 | 100% | 511 | 546 | + | 511 |
| 10_MGAS2096.crt.8 | JHELIIPC_01229 eukaryotic hypersensitive-induced re | 36 | 828 | 67.6 | 6.00E-12 | 36 | 100% | 0 | 0% | 1 | 36 | 100% | 511 | 546 | + | 511 |
| 10_MGAS2096.crt.8 | HPLLMOOM_01228 eukaryotic hypersensitive-induce | 36 | 828 | 67.6 | 6.00E-12 | 36 | 100% | 0 | 0% | 1 | 36 | 100% | 511 | 546 | + | 511 |
| 10_MGAS2096.crt.8 | MCFNMICF_00775 eukaryotic hypersensitive-induced | 36 | 828 | 67.6 | 6.00E-12 | 36 | 100% | 0 | 0% | 1 | 36 | 100% | 511 | 546 | + | 511 |
| 10_MGAS2096.crt.8 | MONDJAPC_00342 eukaryotic hypersensitive-induced | 36 | 828 | 67.6 | 6.00E-12 | 36 | 100% | 0 | 0% | 1 | 36 | 100% | 511 | 546 | + | 511 |
| 10_MGAS2096.crt.9 | FOAEMPCG_00795 hypothetical protein | 34 | 1731 | 63.9 | 7.00E-11 | 34 | 100% | 0 | 0% | 1 | 34 | 100% | 744 | 777 | + | 744 |
| 10_MGAS2096.crt.9 | LLAAGEIN_00651 hypothetical protein | 34 | 1731 | 63.9 | 7.00E-11 | 34 | 100% | 0 | 0% | 1 | 34 | 100% | 744 | 777 | + | 744 |
| 10_MGAS2096.crt.9 | EFGBCCNL_00460 hypothetical protein | 34 | 1731 | 63.9 | 7.00E-11 | 34 | 100% | 0 | 0% | 1 | 34 | 100% | 744 | 777 | + | 744 |
| 10_MGAS2096.crt.9 | JHELIIPC_01221 hypothetical protein | 34 | 1731 | 63.9 | 7.00E-11 | 34 | 100% | 0 | 0% | 1 | 34 | 100% | 744 | 777 | + | 744 |
| 10_MGAS2096.crt.9 | HPLLMOOM_01220 hypothetical protein | 34 | 1731 | 63.9 | 7.00E-11 | 34 | 100% | 0 | 0% | 1 | 34 | 100% | 744 | 777 | + | 744 |
| 10_MGAS2096.crt.9 | MCFNMICF_00783 hypothetical protein | 34 | 1731 | 63.9 | 7.00E-11 | 34 | 100% | 0 | 0% | 1 | 34 | 100% | 744 | 777 | + | 744 |
| 10_MGAS2096.crt.9 | MONDJAPC_00350 hypothetical protein | 34 | 1731 | 63.9 | 7.00E-11 | 34 | 100% | 0 | 0% | 1 | 34 | 100% | 744 | 777 | + | 744 |
| 11_MGAS10750.crt.1 | NOHIT | 33 | | | | | | | | | | | | | | |
| 11_MGAS10750.crt.2 | BNEKOFAL_00650 Transposase from transposon Tn9 | 34 | 1167 | 63.9 | 7.00E-11 | 34 | 100% | 0 | 0% | 1 | 34 | 100% | 1080 | 1113 | - | 1080 |
| 11_MGAS10750.crt.2 | FOAEMPCG_01447 Transposase from transposon Tn9 | 34 | 1167 | 63.9 | 7.00E-11 | 34 | 100% | 0 | 0% | 1 | 34 | 100% | 1080 | 1113 | - | 1080 |
| 11_MGAS10750.crt.2 | MCEIOEBL_00600 Transposase from transposon Tn91 | 34 | 1167 | 63.9 | 7.00E-11 | 34 | 100% | 0 | 0% | 1 | 34 | 100% | 1080 | 1113 | - | 1080 |
| 11_MGAS10750.crt.2 | LLAAGEIN_01392 Transposase from transposon Tn91 | 34 | 1167 | 63.9 | 7.00E-11 | 34 | 100% | 0 | 0% | 1 | 34 | 100% | 1080 | 1113 | - | 1080 |
| 11_MGAS10750.crt.2 | JNLHFPPN_01288 Transposase from transposon Tn91 | 34 | 1167 | 63.9 | 7.00E-11 | 34 | 100% | 0 | 0% | 1 | 34 | 100% | 1080 | 1113 | - | 1080 |
| 11_MGAS10750.crt.2 | ODIAAKBJ_01288 Transposase from transposon Tn91 | 34 | 1167 | 63.9 | 7.00E-11 | 34 | 100% | 0 | 0% | 1 | 34 | 100% | 1080 | 1113 | - | 1080 |
| 11_MGAS10750.crt.2 | ECKACGMC_00529 Transposase from transposon Tn | 34 | 1167 | 63.9 | 7.00E-11 | 34 | 100% | 0 | 0% | 1 | 34 | 100% | 1080 | 1113 | - | 1080 |
| 11_MGAS10750.crt.2 | DIPDIGDN_01287 Transposase from transposon Tn91 | 34 | 1167 | 63.9 | 7.00E-11 | 34 | 100% | 0 | 0% | 1 | 34 | 100% | 1080 | 1113 | - | 1080 |
| 11_MGAS10750.crt.2 | JHELIIPC_00570 Transposase from transposon Tn916 | 34 | 1167 | 63.9 | 7.00E-11 | 34 | 100% | 0 | 0% | 1 | 34 | 100% | 1080 | 1113 | - | 1080 |
| 11_MGAS10750.crt.2 | AJHAHHEL_01339 Transposase from transposon Tn9 | 34 | 1167 | 63.9 | 7.00E-11 | 34 | 100% | 0 | 0% | 1 | 34 | 100% | 1080 | 1113 | - | 1080 |
| 11_MGAS10750.crt.2 | HPLLMOOM_00568 Transposase from transposon Tn | 34 | 1167 | 63.9 | 7.00E-11 | 34 | 100% | 0 | 0% | 1 | 34 | 100% | 1080 | 1113 | - | 1080 |
| 11_MGAS10750.crt.2 | MCFNMICF_01433 Transposase from transposon Tn9 | 34 | 1167 | 63.9 | 7.00E-11 | 34 | 100% | 0 | 0% | 1 | 34 | 100% | 1080 | 1113 | - | 1080 |
| 11_MGAS10750.crt.2 | MONDJAPC_01605 Transposase from transposon Tn9 | 34 | 1167 | 63.9 | 7.00E-11 | 34 | 100% | 0 | 0% | 1 | 34 | 100% | 1080 | 1113 | - | 1080 |
| 11_MGAS10750.crt.3 | JEOJEKGO_01300 hypothetical protein | 35 | 513 | 65.8 | 2.00E-11 | 35 | 100% | 0 | 0% | 1 | 35 | 100% | 246 | 280 | + | 246 |
| 11_MGAS10750.crt.3 | MONDJAPC_01319 hypothetical protein | 35 | 513 | 65.8 | 2.00E-11 | 35 | 100% | 0 | 0% | 1 | 35 | 100% | 246 | 280 | + | 246 |
| 11_MGAS10750.crt.4 | NOHIT | 33 | | | | | | | | | | | | | | |
| 11_MGAS10750.crt.5 | NOHIT | 34 | | | | | | | | | | | | | | |
| 13_NZ131.crt.1 | NOHIT | 30 | | | | | | | | | | | | | | |
| 13_NZ131.crt.2 | NOHIT | 31 | | | | | | | | | | | | | | |
| 13_NZ131.crt.3 | NOHIT | 30 | | | | | | | | | | | | | | |
| 13_NZ131.crt.4 | NOHIT | 30 | | | | | | | | | | | | | | |
| 13_NZ131.crt.5 | NOHIT | 0 | | | | | | | | | | | | | | |
| 13_NZ131.crt.6 | NOHIT | 35 | | | | | | | | | | | | | | |
| 13_NZ131.crt.7 | NOHIT | 35 | | | | | | | | | | | | | | |
| 13_NZ131.crt.8 | MDEAFGPO_00918 hypothetical protein | 35 | 720 | 65.8 | 2.00E-11 | 35 | 100% | 0 | 0% | 1 | 35 | 100% | 399 | 433 | + | 399 |
| 13_NZ131.crt.9 | BNEKOFAL_00469 hypothetical protein | 35 | 183 | 65.8 | 2.00E-11 | 35 | 100% | 0 | 0% | 1 | 35 | 100% | 95 | 129 | + | 95 |
| 13_NZ131.crt.9 | MCEIOEBL_00418 hypothetical protein | 35 | 183 | 65.8 | 2.00E-11 | 35 | 100% | 0 | 0% | 1 | 35 | 100% | 95 | 129 | + | 95 |
| 13_NZ131.crt.9 | LLAAGEIN_01332 hypothetical protein | 35 | 183 | 65.8 | 2.00E-11 | 35 | 100% | 0 | 0% | 1 | 35 | 100% | 95 | 129 | + | 95 |
| 13_NZ131.crt.9 | DCKFNFNP_00808 hypothetical protein | 35 | 189 | 65.8 | 2.00E-11 | 35 | 100% | 0 | 0% | 1 | 35 | 100% | 95 | 129 | + | 95 |
| 13_NZ131.crt.9 | ECKACGMC_00589 hypothetical protein | 35 | 183 | 65.8 | 2.00E-11 | 35 | 100% | 0 | 0% | 1 | 35 | 100% | 95 | 129 | + | 95 |
| 13_NZ131.crt.9 | GPBJDOFB_01469 hypothetical protein | 35 | 183 | 65.8 | 2.00E-11 | 35 | 100% | 0 | 0% | 1 | 35 | 100% | 95 | 129 | + | 95 |
| 13_NZ131.crt.9 | AJHAHHEL_01279 hypothetical protein | 35 | 183 | 65.8 | 2.00E-11 | 35 | 100% | 0 | 0% | 1 | 35 | 100% | 95 | 129 | + | 95 |
| 13_NZ131.crt.9 | HEGDFFNC_00593 hypothetical protein | 35 | 183 | 65.8 | 2.00E-11 | 35 | 100% | 0 | 0% | 1 | 35 | 100% | 95 | 129 | + | 95 |
| 13_NZ131.crt.9 | NELECJBE_01577 hypothetical protein | 35 | 183 | 65.8 | 2.00E-11 | 35 | 100% | 0 | 0% | 1 | 35 | 100% | 95 | 129 | + | 95 |
| 13_NZ131.crt.10 | NOHIT | 33 | | | | | | | | | | | | | | |
| 15_MGAS15252.crt.1 | NOHIT | 37 | | | | | | | | | | | | | | |
| 15_MGAS15252.crt.2 | NOHIT | 37 | | | | | | | | | | | | | | |
| 16_MGAS1882.crt.1 | NOHIT | 30 | | | | | | | | | | | | | | |
| 16_MGAS1882.crt.2 | NOHIT | 30 | | | | | | | | | | | | | | |
| 16_MGAS1882.crt.3 | NOHIT | 30 | | | | | | | | | | | | | | |
| 17_M1_476.crt.1 | NOHIT | 30 | | | | | | | | | | | | | | |
| 17_M1_476.crt.2 | NOHIT | 30 | | | | | | | | | | | | | | |
| 17_M1_476.crt.3 | NOHIT | 30 | | | | | | | | | | | | | | |
| 17_M1_476.crt.4 | NOHIT | 0 | | | | | | | | | | | | | | |
| 17_M1_476.crt.5 | NOHIT | 33 | | | | | | | | | | | | | | |
| 17_M1_476.crt.6 | NOHIT | 34 | | | | | | | | | | | | | | |
| 17_M1_476.crt.7 | NOHIT | 33 | | | | | | | | | | | | | | |
| 17_M1_476.crt.8 | NOHIT | 35 | | | | | | | | | | | | | | |
| 18_A20.crt.1 | NOHIT | 30 | | | | | | | | | | | | | | |
| 18_A20.crt.2 | NOHIT | 30 | | | | | | | | | | | | | | |
| 18_A20.crt.3 | NOHIT | 30 | | | | | | | | | | | | | | |
| 18_A20.crt.4 | NOHIT | 0 | | | | | | | | | | | | | | |
| 18_A20.crt.5 | NOHIT | 33 | | | | | | | | | | | | | | |
| 18_A20.crt.6 | NOHIT | 34 | | | | | | | | | | | | | | |
| 18_A20.crt.7 | NOHIT | 33 | | | | | | | | | | | | | | |
| 18_A20.crt.8 | NOHIT | 35 | | | | | | | | | | | | | | |
| 1_SF370.crt.1 | NOHIT | 30 | | | | | | | | | | | | | | |
| 1_SF370.crt.2 | NOHIT | 30 | | | | | | | | | | | | | | |
| 1_SF370.crt.3 | NOHIT | 30 | | | | | | | | | | | | | | |
| 1_SF370.crt.4 | NOHIT | 30 | | | | | | | | | | | | | | |
| 1_SF370.crt.5 | NOHIT | 30 | | | | | | | | | | | | | | |
| 1_SF370.crt.6 | NOHIT | 30 | | | | | | | | | | | | | | |
| 1_SF370.crt.7 | NOHIT | 0 | | | | | | | | | | | | | | |
| 1_SF370.crt.8 | NOHIT | 33 | | | | | | | | | | | | | | |
| 1_SF370.crt.9 | NOHIT | 32 | | | | | | | | | | | | | | |
| 1_SF370.crt.10 | NOHIT | 34 | | | | | | | | | | | | | | |
| 20_H293.crt.1 | NOHIT | 30 | | | | | | | | | | | | | | |
| 20_H293.crt.2 | NOHIT | 30 | | | | | | | | | | | | | | |
| 20_H293.crt.3 | NOHIT | 30 | | | | | | | | | | | | | | |
| 20_H293.crt.4 | NOHIT | 30 | | | | | | | | | | | | | | |
| 20_H293.crt.5 | NOHIT | 30 | | | | | | | | | | | | | | |
| 20_H293.crt.6 | NOHIT | 0 | | | | | | | | | | | | | | |
| 20_H293.crt.7 | NOHIT | 19 | | | | | | | | | | | | | | |
| 20_H293.crt.8 | NOHIT | 28 | | | | | | | | | | | | | | |
| 20_H293.crt.9 | NOHIT | 19 | | | | | | | | | | | | | | |
| 20_H293.crt.10 | NOHIT | 19 | | | | | | | | | | | | | | |
| 20_H293.crt.11 | NOHIT | 0 | | | | | | | | | | | | | | |
| 20_H293.crt.12 | NOHIT | 34 | | | | | | | | | | | | | | |
| 20_H293.crt.13 | NOHIT | 34 | | | | | | | | | | | | | | |
| 20_H293.crt.14 | GGADKFFH_01023 putative DNA primase - phage as | 37 | 2385 | 63.9 | 9.00E-11 | 34 | 100% | 0 | 0% | 4 | 37 | 91.89% | 1638 | 1671 | + | 1638 |
| 20_H293.crt.14 | JNLHFPPN_01503 putative DNA primase - phage asso | 37 | 2385 | 63.9 | 9.00E-11 | 34 | 100% | 0 | 0% | 4 | 37 | 91.89% | 1638 | 1671 | + | 1638 |
| 20_H293.crt.14 | ODIAAKBJ_01503 putative DNA primase - phage asso | 37 | 2385 | 63.9 | 9.00E-11 | 34 | 100% | 0 | 0% | 4 | 37 | 91.89% | 1638 | 1671 | + | 1638 |
| 20_H293.crt.14 | DIPDIGDN_01504 putative DNA primase - phage asso | 37 | 2385 | 63.9 | 9.00E-11 | 34 | 100% | 0 | 0% | 4 | 37 | 91.89% | 1638 | 1671 | + | 1638 |
| 20_H293.crt.14 | MDEAFGPO_00463 putative DNA primase - phage as | 37 | 2385 | 63.9 | 9.00E-11 | 34 | 100% | 0 | 0% | 4 | 37 | 91.89% | 1638 | 1671 | + | 1638 |
| 20_H293.crt.14 | DMLKMGFL_00783 putative DNA primase - phage as | 37 | 2385 | 63.9 | 9.00E-11 | 34 | 100% | 0 | 0% | 4 | 37 | 91.89% | 1638 | 1671 | + | 1638 |
| 20_H293.crt.14 | HEGDFFNC_01101 putative DNA primase - phage ass | 37 | 2385 | 63.9 | 9.00E-11 | 34 | 100% | 0 | 0% | 4 | 37 | 91.89% | 1638 | 1671 | + | 1638 |
| 20_H293.crt.15 | NOHIT | 35 | | | | | | | | | | | | | | |
| 20_H293.crt.16 | MHKCMMMI_00605 hypothetical protein | 34 | 678 | 63.9 | 7.00E-11 | 34 | 100% | 0 | 0% | 1 | 34 | 100% | 593 | 626 | - | 593 |
| 20_H293.crt.16 | FOAEMPCG_01323 hypothetical protein | 34 | 678 | 63.9 | 7.00E-11 | 34 | 100% | 0 | 0% | 1 | 34 | 100% | 593 | 626 | - | 593 |
| 20_H293.crt.16 | MCEIOEBL_00630 hypothetical protein | 34 | 678 | 63.9 | 7.00E-11 | 34 | 100% | 0 | 0% | 1 | 34 | 100% | 593 | 626 | - | 593 |
| 20_H293.crt.16 | LAKBNEPL_01310 hypothetical protein | 34 | 678 | 63.9 | 7.00E-11 | 34 | 100% | 0 | 0% | 1 | 34 | 100% | 593 | 626 | - | 593 |
| 20_H293.crt.16 | PLPKHLCC_00644 hypothetical protein | 34 | 678 | 63.9 | 7.00E-11 | 34 | 100% | 0 | 0% | 1 | 34 | 100% | 593 | 626 | - | 593 |
| 20_H293.crt.16 | DCKFNFNP_01264 hypothetical protein | 34 | 678 | 63.9 | 7.00E-11 | 34 | 100% | 0 | 0% | 1 | 34 | 100% | 593 | 626 | - | 593 |
| 20_H293.crt.16 | JNLHFPPN_01257 hypothetical protein | 34 | 678 | 63.9 | 7.00E-11 | 34 | 100% | 0 | 0% | 1 | 34 | 100% | 593 | 626 | - | 593 |
| 20_H293.crt.16 | ODIAAKBJ_01257 hypothetical protein | 34 | 678 | 63.9 | 7.00E-11 | 34 | 100% | 0 | 0% | 1 | 34 | 100% | 593 | 626 | - | 593 |
| 20_H293.crt.16 | DIPDIGDN_01256 hypothetical protein | 34 | 678 | 63.9 | 7.00E-11 | 34 | 100% | 0 | 0% | 1 | 34 | 100% | 593 | 626 | - | 593 |
| 20_H293.crt.16 | JHELIIPC_00693 hypothetical protein | 34 | 678 | 63.9 | 7.00E-11 | 34 | 100% | 0 | 0% | 1 | 34 | 100% | 593 | 626 | - | 593 |
| 20_H293.crt.16 | IPBLJGMA_01319 hypothetical protein | 34 | 678 | 63.9 | 7.00E-11 | 34 | 100% | 0 | 0% | 1 | 34 | 100% | 593 | 626 | - | 593 |
| 20_H293.crt.16 | GLBJHDNN_01264 hypothetical protein | 34 | 678 | 63.9 | 7.00E-11 | 34 | 100% | 0 | 0% | 1 | 34 | 100% | 593 | 626 | - | 593 |
| 20_H293.crt.16 | HPLLMOOM_00691 hypothetical protein | 34 | 678 | 63.9 | 7.00E-11 | 34 | 100% | 0 | 0% | 1 | 34 | 100% | 593 | 626 | - | 593 |
| 20_H293.crt.16 | MCFNMICF_01310 hypothetical protein | 34 | 678 | 63.9 | 7.00E-11 | 34 | 100% | 0 | 0% | 1 | 34 | 100% | 593 | 626 | - | 593 |
| 20_H293.crt.16 | MONDJAPC_01575 hypothetical protein | 34 | 678 | 63.9 | 7.00E-11 | 34 | 100% | 0 | 0% | 1 | 34 | 100% | 593 | 626 | - | 593 |
| 20_H293.crt.17 | NOHIT | 35 | | | | | | | | | | | | | | |
| 20_H293.crt.18 | NOHIT | 33 | | | | | | | | | | | | | | |
| 20_H293.crt.19 | NOHIT | 34 | | | | | | | | | | | | | | |
| 20_H293.crt.20 | NOHIT | 37 | | | | | | | | | | | | | | |
| 20_H293.crt.21 | NOHIT | 33 | | | | | | | | | | | | | | |
| 20_H293.crt.22 | BGBHDJOA_00791 putative repressor protein - phage | 34 | 756 | 63.9 | 7.00E-11 | 34 | 100% | 0 | 0% | 1 | 34 | 100% | 687 | 720 | + | 687 |
| 20_H293.crt.22 | EJBPJIHF_01687 putative repressor protein - phage as | 34 | 759 | 63.9 | 7.00E-11 | 34 | 100% | 0 | 0% | 1 | 34 | 100% | 696 | 729 | + | 696 |
| 20_H293.crt.22 | GBCACICN_01644 putative repressor protein - phage | 34 | 759 | 63.9 | 7.00E-11 | 34 | 100% | 0 | 0% | 1 | 34 | 100% | 696 | 729 | + | 696 |
| 20_H293.crt.22 | GBCACICN_00736 putative repressor protein - phage | 34 | 756 | 63.9 | 7.00E-11 | 34 | 100% | 0 | 0% | 1 | 34 | 100% | 687 | 720 | + | 687 |
| 20_H293.crt.22 | EFGBCCNL_00740 putative repressor protein - phage | 34 | 759 | 63.9 | 7.00E-11 | 34 | 100% | 0 | 0% | 1 | 34 | 100% | 696 | 729 | + | 696 |
| 20_H293.crt.22 | DMLKMGFL_01526 putative repressor protein - phag | 34 | 759 | 63.9 | 7.00E-11 | 34 | 100% | 0 | 0% | 1 | 34 | 100% | 696 | 729 | + | 696 |
| 20_H293.crt.22 | IPNPMFKD_00585 putative repressor protein - phage | 34 | 750 | 63.9 | 7.00E-11 | 34 | 100% | 0 | 0% | 1 | 34 | 100% | 687 | 720 | + | 687 |
| 20_H293.crt.22 | IPBLJGMA_00554 putative repressor protein - phage a | 34 | 750 | 63.9 | 7.00E-11 | 34 | 100% | 0 | 0% | 1 | 34 | 100% | 687 | 720 | + | 687 |
| 20_H293.crt.22 | ELLIANMJ_00803 putative repressor protein - phage a | 34 | 756 | 63.9 | 7.00E-11 | 34 | 100% | 0 | 0% | 1 | 34 | 100% | 687 | 720 | + | 687 |
| 20_H293.crt.22 | ELLIANMJ_00546 putative repressor protein - phage a | 34 | 750 | 63.9 | 7.00E-11 | 34 | 100% | 0 | 0% | 1 | 34 | 100% | 687 | 720 | + | 687 |
| 20_H293.crt.23 | NOHIT | 33 | | | | | | | | | | | | | | |
| 20_H293.crt.24 | BNEKOFAL_00698 hypothetical protein | 34 | 717 | 63.9 | 7.00E-11 | 34 | 100% | 0 | 0% | 1 | 34 | 100% | 452 | 485 | - | 452 |
| 20_H293.crt.24 | MHKCMMMI_00623 hypothetical protein | 34 | 717 | 63.9 | 7.00E-11 | 34 | 100% | 0 | 0% | 1 | 34 | 100% | 452 | 485 | - | 452 |
| 20_H293.crt.24 | FOAEMPCG_01305 hypothetical protein | 34 | 717 | 63.9 | 7.00E-11 | 34 | 100% | 0 | 0% | 1 | 34 | 100% | 452 | 485 | - | 452 |
| 20_H293.crt.24 | MCEIOEBL_00648 hypothetical protein | 34 | 717 | 63.9 | 7.00E-11 | 34 | 100% | 0 | 0% | 1 | 34 | 100% | 452 | 485 | - | 452 |
| 20_H293.crt.24 | HHEKEGMA_00585 hypothetical protein | 34 | 717 | 63.9 | 7.00E-11 | 34 | 100% | 0 | 0% | 1 | 34 | 100% | 452 | 485 | - | 452 |
| 20_H293.crt.24 | CMOPCPHD_00644 hypothetical protein | 34 | 717 | 63.9 | 7.00E-11 | 34 | 100% | 0 | 0% | 1 | 34 | 100% | 452 | 485 | - | 452 |
| 20_H293.crt.24 | CMOPCPHD_00580 hypothetical protein | 34 | 717 | 63.9 | 7.00E-11 | 34 | 100% | 0 | 0% | 1 | 34 | 100% | 452 | 485 | - | 452 |
| 20_H293.crt.24 | LLAAGEIN_01345 hypothetical protein | 34 | 717 | 63.9 | 7.00E-11 | 34 | 100% | 0 | 0% | 1 | 34 | 100% | 452 | 485 | - | 452 |
| 20_H293.crt.24 | LAKBNEPL_01292 hypothetical protein | 34 | 717 | 63.9 | 7.00E-11 | 34 | 100% | 0 | 0% | 1 | 34 | 100% | 452 | 485 | - | 452 |
| 20_H293.crt.24 | PLPKHLCC_00662 hypothetical protein | 34 | 717 | 63.9 | 7.00E-11 | 34 | 100% | 0 | 0% | 1 | 34 | 100% | 452 | 485 | - | 452 |
| 20_H293.crt.24 | DCKFNFNP_01317 hypothetical protein | 34 | 708 | 63.9 | 7.00E-11 | 34 | 100% | 0 | 0% | 1 | 34 | 100% | 443 | 476 | - | 443 |
| 20_H293.crt.24 | DCKFNFNP_01246 hypothetical protein | 34 | 717 | 63.9 | 7.00E-11 | 34 | 100% | 0 | 0% | 1 | 34 | 100% | 452 | 485 | - | 452 |
| 20_H293.crt.24 | JNLHFPPN_01238 hypothetical protein | 34 | 717 | 63.9 | 7.00E-11 | 34 | 100% | 0 | 0% | 1 | 34 | 100% | 452 | 485 | - | 452 |
| 20_H293.crt.24 | ODIAAKBJ_01238 hypothetical protein | 34 | 717 | 63.9 | 7.00E-11 | 34 | 100% | 0 | 0% | 1 | 34 | 100% | 452 | 485 | - | 452 |
| 20_H293.crt.24 | ECKACGMC_00576 hypothetical protein | 34 | 717 | 63.9 | 7.00E-11 | 34 | 100% | 0 | 0% | 1 | 34 | 100% | 452 | 485 | - | 452 |
| 20_H293.crt.24 | DIPDIGDN_01237 hypothetical protein | 34 | 717 | 63.9 | 7.00E-11 | 34 | 100% | 0 | 0% | 1 | 34 | 100% | 452 | 485 | - | 452 |
| 20_H293.crt.24 | JHELIIPC_00711 hypothetical protein | 34 | 696 | 63.9 | 7.00E-11 | 34 | 100% | 0 | 0% | 1 | 34 | 100% | 431 | 464 | - | 431 |
| 20_H293.crt.24 | GPBJDOFB_01483 hypothetical protein | 34 | 717 | 63.9 | 7.00E-11 | 34 | 100% | 0 | 0% | 1 | 34 | 100% | 452 | 485 | - | 452 |
| 20_H293.crt.24 | AJHAHHEL_01292 hypothetical protein | 34 | 717 | 63.9 | 7.00E-11 | 34 | 100% | 0 | 0% | 1 | 34 | 100% | 452 | 485 | - | 452 |
| 20_H293.crt.24 | HEGDFFNC_00579 hypothetical protein | 34 | 717 | 63.9 | 7.00E-11 | 34 | 100% | 0 | 0% | 1 | 34 | 100% | 452 | 485 | - | 452 |
| 20_H293.crt.24 | KCAAAFFC_00607 hypothetical protein | 34 | 717 | 63.9 | 7.00E-11 | 34 | 100% | 0 | 0% | 1 | 34 | 100% | 452 | 485 | - | 452 |
| 20_H293.crt.24 | IPBLJGMA_01301 hypothetical protein | 34 | 717 | 63.9 | 7.00E-11 | 34 | 100% | 0 | 0% | 1 | 34 | 100% | 452 | 485 | - | 452 |
| 20_H293.crt.24 | GLBJHDNN_01246 hypothetical protein | 34 | 717 | 63.9 | 7.00E-11 | 34 | 100% | 0 | 0% | 1 | 34 | 100% | 452 | 485 | - | 452 |
| 20_H293.crt.24 | NELECJBE_01381 hypothetical protein | 34 | 717 | 63.9 | 7.00E-11 | 34 | 100% | 0 | 0% | 1 | 34 | 100% | 452 | 485 | - | 452 |
| 20_H293.crt.24 | HPLLMOOM_00709 hypothetical protein | 34 | 717 | 63.9 | 7.00E-11 | 34 | 100% | 0 | 0% | 1 | 34 | 100% | 452 | 485 | - | 452 |
| 20_H293.crt.24 | MCFNMICF_01292 hypothetical protein | 34 | 717 | 63.9 | 7.00E-11 | 34 | 100% | 0 | 0% | 1 | 34 | 100% | 452 | 485 | - | 452 |
| 20_H293.crt.24 | MONDJAPC_01557 hypothetical protein | 34 | 717 | 63.9 | 7.00E-11 | 34 | 100% | 0 | 0% | 1 | 34 | 100% | 452 | 485 | - | 452 |
| 20_H293.crt.24 | EEABGJJD_00588 hypothetical protein | 34 | 717 | 63.9 | 7.00E-11 | 34 | 100% | 0 | 0% | 1 | 34 | 100% | 452 | 485 | - | 452 |
| 22_STAB901.crt.1 | NOHIT | 30 | | | | | | | | | | | | | | |
| 22_STAB901.crt.2 | NOHIT | 30 | | | | | | | | | | | | | | |
| 22_STAB901.crt.3 | NOHIT | 30 | | | | | | | | | | | | | | |
| 22_STAB901.crt.4 | NOHIT | 30 | | | | | | | | | | | | | | |
| 22_STAB901.crt.5 | NOHIT | 0 | | | | | | | | | | | | | | |
| 22_STAB901.crt.6 | NOHIT | 29 | | | | | | | | | | | | | | |
| 22_STAB901.crt.7 | NOHIT | 31 | | | | | | | | | | | | | | |
| 22_STAB901.crt.8 | NOHIT | 32 | | | | | | | | | | | | | | |
| 25_7F7.crt.1 | NOHIT | 30 | | | | | | | | | | | | | | |
| 25_7F7.crt.2 | NOHIT | 30 | | | | | | | | | | | | | | |
| 25_7F7.crt.3 | NOHIT | 30 | | | | | | | | | | | | | | |
| 25_7F7.crt.4 | NOHIT | 30 | | | | | | | | | | | | | | |
| 25_7F7.crt.5 | NOHIT | 30 | | | | | | | | | | | | | | |
| 25_7F7.crt.6 | NOHIT | 30 | | | | | | | | | | | | | | |
| 25_7F7.crt.7 | NOHIT | 30 | | | | | | | | | | | | | | |
| 25_7F7.crt.8 | NOHIT | 30 | | | | | | | | | | | | | | |
| 26_HKU360.crt.1 | NOHIT | 38 | | | | | | | | | | | | | | |
| 26_HKU360.crt.2 | NOHIT | 37 | | | | | | | | | | | | | | |
| 26_HKU360.crt.3 | NOHIT | 0 | | | | | | | | | | | | | | |
| 26_HKU360.crt.4 | NOHIT | 32 | | | | | | | | | | | | | | |
| 26_HKU360.crt.5 | BNEKOFAL_00674 hypothetical protein | 34 | 285 | 63.9 | 7.00E-11 | 34 | 100% | 0 | 0% | 1 | 34 | 100% | 19 | 52 | + | 19 |
| 26_HKU360.crt.5 | FOAEMPCG_01040 hypothetical protein | 34 | 285 | 63.9 | 7.00E-11 | 34 | 100% | 0 | 0% | 1 | 34 | 100% | 19 | 52 | + | 19 |
| 26_HKU360.crt.5 | MCEIOEBL_00624 hypothetical protein | 34 | 285 | 63.9 | 7.00E-11 | 34 | 100% | 0 | 0% | 1 | 34 | 100% | 19 | 52 | + | 19 |
| 26_HKU360.crt.5 | PPKGMMPD_01012 hypothetical protein | 34 | 285 | 63.9 | 7.00E-11 | 34 | 100% | 0 | 0% | 1 | 34 | 100% | 19 | 52 | + | 19 |
| 26_HKU360.crt.5 | LLAAGEIN_01368 hypothetical protein | 34 | 285 | 63.9 | 7.00E-11 | 34 | 100% | 0 | 0% | 1 | 34 | 100% | 19 | 52 | + | 19 |
| 26_HKU360.crt.5 | AIKJPEBD_01573 hypothetical protein | 34 | 285 | 63.9 | 7.00E-11 | 34 | 100% | 0 | 0% | 1 | 34 | 100% | 19 | 52 | + | 19 |
| 26_HKU360.crt.5 | FPFNMMFM_00882 hypothetical protein | 34 | 285 | 63.9 | 7.00E-11 | 34 | 100% | 0 | 0% | 1 | 34 | 100% | 19 | 52 | + | 19 |
| 26_HKU360.crt.5 | IJFGMCCO_00995 hypothetical protein | 34 | 285 | 63.9 | 7.00E-11 | 34 | 100% | 0 | 0% | 1 | 34 | 100% | 19 | 52 | + | 19 |
| 26_HKU360.crt.5 | EJBPJIHF_01045 hypothetical protein | 34 | 285 | 63.9 | 7.00E-11 | 34 | 100% | 0 | 0% | 1 | 34 | 100% | 19 | 52 | + | 19 |
| 26_HKU360.crt.5 | LAKBNEPL_01011 hypothetical protein | 34 | 285 | 63.9 | 7.00E-11 | 34 | 100% | 0 | 0% | 1 | 34 | 100% | 19 | 52 | + | 19 |
| 26_HKU360.crt.5 | MMBCLDPD_00865 hypothetical protein | 34 | 285 | 63.9 | 7.00E-11 | 34 | 100% | 0 | 0% | 1 | 34 | 100% | 19 | 52 | + | 19 |
| 26_HKU360.crt.5 | PLPKHLCC_00893 hypothetical protein | 34 | 285 | 63.9 | 7.00E-11 | 34 | 100% | 0 | 0% | 1 | 34 | 100% | 19 | 52 | + | 19 |
| 26_HKU360.crt.5 | JNLHFPPN_01263 hypothetical protein | 34 | 285 | 63.9 | 7.00E-11 | 34 | 100% | 0 | 0% | 1 | 34 | 100% | 19 | 52 | + | 19 |
| 26_HKU360.crt.5 | ODIAAKBJ_01263 hypothetical protein | 34 | 285 | 63.9 | 7.00E-11 | 34 | 100% | 0 | 0% | 1 | 34 | 100% | 19 | 52 | + | 19 |
| 26_HKU360.crt.5 | ECKACGMC_00553 hypothetical protein | 34 | 285 | 63.9 | 7.00E-11 | 34 | 100% | 0 | 0% | 1 | 34 | 100% | 19 | 52 | + | 19 |
| 26_HKU360.crt.5 | DIPDIGDN_01262 hypothetical protein | 34 | 285 | 63.9 | 7.00E-11 | 34 | 100% | 0 | 0% | 1 | 34 | 100% | 19 | 52 | + | 19 |
| 26_HKU360.crt.5 | EFGBCCNL_00703 hypothetical protein | 34 | 285 | 63.9 | 7.00E-11 | 34 | 100% | 0 | 0% | 1 | 34 | 100% | 19 | 52 | + | 19 |
| 26_HKU360.crt.5 | JHELIIPC_00975 hypothetical protein | 34 | 285 | 63.9 | 7.00E-11 | 34 | 100% | 0 | 0% | 1 | 34 | 100% | 19 | 52 | + | 19 |
| 26_HKU360.crt.5 | GPBJDOFB_00997 hypothetical protein | 34 | 285 | 63.9 | 7.00E-11 | 34 | 100% | 0 | 0% | 1 | 34 | 100% | 19 | 52 | + | 19 |
| 26_HKU360.crt.5 | ACJECOPP_01021 hypothetical protein | 34 | 285 | 63.9 | 7.00E-11 | 34 | 100% | 0 | 0% | 1 | 34 | 100% | 19 | 52 | + | 19 |
| 26_HKU360.crt.5 | ICMALKPE_01033 hypothetical protein | 34 | 285 | 63.9 | 7.00E-11 | 34 | 100% | 0 | 0% | 1 | 34 | 100% | 19 | 52 | + | 19 |
| 26_HKU360.crt.5 | AJHAHHEL_01315 hypothetical protein | 34 | 285 | 63.9 | 7.00E-11 | 34 | 100% | 0 | 0% | 1 | 34 | 100% | 19 | 52 | + | 19 |
| 26_HKU360.crt.5 | DMLKMGFL_01490 hypothetical protein | 34 | 285 | 63.9 | 7.00E-11 | 34 | 100% | 0 | 0% | 1 | 34 | 100% | 19 | 52 | + | 19 |
| 26_HKU360.crt.5 | HEGDFFNC_00554 hypothetical protein | 34 | 285 | 63.9 | 7.00E-11 | 34 | 100% | 0 | 0% | 1 | 34 | 100% | 19 | 52 | + | 19 |
| 26_HKU360.crt.5 | IIJJCGIL_01024 hypothetical protein | 34 | 285 | 63.9 | 7.00E-11 | 34 | 100% | 0 | 0% | 1 | 34 | 100% | 19 | 52 | + | 19 |
| 26_HKU360.crt.5 | GLBJHDNN_01015 hypothetical protein | 34 | 285 | 63.9 | 7.00E-11 | 34 | 100% | 0 | 0% | 1 | 34 | 100% | 19 | 52 | + | 19 |
| 26_HKU360.crt.5 | HPLLMOOM_00975 hypothetical protein | 34 | 285 | 63.9 | 7.00E-11 | 34 | 100% | 0 | 0% | 1 | 34 | 100% | 19 | 52 | + | 19 |
| 26_HKU360.crt.5 | MCFNMICF_01027 hypothetical protein | 34 | 285 | 63.9 | 7.00E-11 | 34 | 100% | 0 | 0% | 1 | 34 | 100% | 19 | 52 | + | 19 |
| 26_HKU360.crt.5 | MONDJAPC_01581 hypothetical protein | 34 | 285 | 63.9 | 7.00E-11 | 34 | 100% | 0 | 0% | 1 | 34 | 100% | 19 | 52 | + | 19 |
| 26_HKU360.crt.6 | NOHIT | 34 | | | | | | | | | | | | | | |
| 26_HKU360.crt.7 | NOHIT | 33 | | | | | | | | | | | | | | |
| 27_10.crt.1 | NOHIT | 30 | | | | | | | | | | | | | | |
| 27_10.crt.2 | NOHIT | 30 | | | | | | | | | | | | | | |
| 27_10.crt.3 | NOHIT | 30 | | | | | | | | | | | | | | |
| 27_10.crt.4 | NOHIT | 30 | | | | | | | | | | | | | | |
| 27_10.crt.5 | NOHIT | 30 | | | | | | | | | | | | | | |
| 27_10.crt.6 | NOHIT | 30 | | | | | | | | | | | | | | |
| 27_10.crt.7 | NOHIT | 30 | | | | | | | | | | | | | | |
| 27_10.crt.8 | NOHIT | 30 | | | | | | | | | | | | | | |
| 27_10.crt.9 | NOHIT | 30 | | | | | | | | | | | | | | |
| 27_10.crt.10 | NOHIT | 30 | | | | | | | | | | | | | | |
| 27_10.crt.11 | NOHIT | 0 | | | | | | | | | | | | | | |
| 27_10.crt.12 | NOHIT | 29 | | | | | | | | | | | | | | |
| 27_10.crt.13 | NOHIT | 31 | | | | | | | | | | | | | | |
| 27_10.crt.14 | NOHIT | 32 | | | | | | | | | | | | | | |
| 28_MTB313.crt.1 | NOHIT | 29 | | | | | | | | | | | | | | |
| 28_MTB313.crt.2 | NOHIT | 31 | | | | | | | | | | | | | | |
| 28_MTB313.crt.3 | NOHIT | 29 | | | | | | | | | | | | | | |
| 28_MTB313.crt.4 | NOHIT | 0 | | | | | | | | | | | | | | |
| 28_MTB313.crt.5 | NOHIT | 33 | | | | | | | | | | | | | | |
| 28_MTB313.crt.6 | NOHIT | 34 | | | | | | | | | | | | | | |
| 28_MTB313.crt.7 | NOHIT | 33 | | | | | | | | | | | | | | |
| 28_MTB313.crt.8 | NOHIT | 35 | | | | | | | | | | | | | | |
| 29_MTB314.crt.1 | NOHIT | 30 | | | | | | | | | | | | | | |
| 29_MTB314.crt.2 | NOHIT | 30 | | | | | | | | | | | | | | |
| 29_MTB314.crt.3 | NOHIT | 30 | | | | | | | | | | | | | | |
| 29_MTB314.crt.4 | NOHIT | 0 | | | | | | | | | | | | | | |
| 29_MTB314.crt.5 | NOHIT | 33 | | | | | | | | | | | | | | |
| 29_MTB314.crt.6 | NOHIT | 33 | | | | | | | | | | | | | | |
| 29_MTB314.crt.7 | NOHIT | 33 | | | | | | | | | | | | | | |
| 29_MTB314.crt.8 | NOHIT | 35 | | | | | | | | | | | | | | |
| 31_AP1.crt.1 | NOHIT | 35 | | | | | | | | | | | | | | |
| 31_AP1.crt.2 | NOHIT | 33 | | | | | | | | | | | | | | |
| 31_AP1.crt.3 | NOHIT | 34 | | | | | | | | | | | | | | |
| 31_AP1.crt.4 | NOHIT | 33 | | | | | | | | | | | | | | |
| 35_NGAS596.crt.1 | NOHIT | 32 | | | | | | | | | | | | | | |
| 35_NGAS596.crt.2 | NOHIT | 33 | | | | | | | | | | | | | | |
| 35_NGAS596.crt.3 | MHKCMMMI_01167 hypothetical protein | 34 | 1863 | 63.9 | 7.00E-11 | 34 | 100% | 0 | 0% | 1 | 34 | 100% | 1146 | 1179 | + | 1146 |
| 35_NGAS596.crt.3 | FOAEMPCG_01399 hypothetical protein | 34 | 1863 | 63.9 | 7.00E-11 | 34 | 100% | 0 | 0% | 1 | 34 | 100% | 1146 | 1179 | + | 1146 |
| 35_NGAS596.crt.3 | AIKJPEBD_01597 hypothetical protein | 34 | 1863 | 63.9 | 7.00E-11 | 34 | 100% | 0 | 0% | 1 | 34 | 100% | 1146 | 1179 | + | 1146 |
| 35_NGAS596.crt.3 | FDNDANMJ_00819 hypothetical protein | 34 | 1863 | 63.9 | 7.00E-11 | 34 | 100% | 0 | 0% | 1 | 34 | 100% | 1146 | 1179 | + | 1146 |
| 35_NGAS596.crt.3 | EJBPJIHF_01021 hypothetical protein | 34 | 1863 | 63.9 | 7.00E-11 | 34 | 100% | 0 | 0% | 1 | 34 | 100% | 1146 | 1179 | + | 1146 |
| 35_NGAS596.crt.3 | MMBCLDPD_00889 hypothetical protein | 34 | 1863 | 63.9 | 7.00E-11 | 34 | 100% | 0 | 0% | 1 | 34 | 100% | 1146 | 1179 | + | 1146 |
| 35_NGAS596.crt.3 | ECKACGMC_00893 hypothetical protein | 34 | 1863 | 63.9 | 7.00E-11 | 34 | 100% | 0 | 0% | 1 | 34 | 100% | 1146 | 1179 | + | 1146 |
| 35_NGAS596.crt.3 | JHELIIPC_00617 hypothetical protein | 34 | 1863 | 63.9 | 7.00E-11 | 34 | 100% | 0 | 0% | 1 | 34 | 100% | 1146 | 1179 | + | 1146 |
| 35_NGAS596.crt.3 | ACJECOPP_00997 hypothetical protein | 34 | 1863 | 63.9 | 7.00E-11 | 34 | 100% | 0 | 0% | 1 | 34 | 100% | 1146 | 1179 | + | 1146 |
| 35_NGAS596.crt.3 | ICMALKPE_01009 hypothetical protein | 34 | 1863 | 63.9 | 7.00E-11 | 34 | 100% | 0 | 0% | 1 | 34 | 100% | 1146 | 1179 | + | 1146 |
| 35_NGAS596.crt.3 | IPNPMFKD_00893 hypothetical protein | 34 | 1863 | 63.9 | 7.00E-11 | 34 | 100% | 0 | 0% | 1 | 34 | 100% | 1146 | 1179 | + | 1146 |
| 35_NGAS596.crt.3 | IPBLJGMA_00850 hypothetical protein | 34 | 1863 | 63.9 | 7.00E-11 | 34 | 100% | 0 | 0% | 1 | 34 | 100% | 1146 | 1179 | + | 1146 |
| 35_NGAS596.crt.3 | IIJJCGIL_01000 hypothetical protein | 34 | 1863 | 63.9 | 7.00E-11 | 34 | 100% | 0 | 0% | 1 | 34 | 100% | 1146 | 1179 | + | 1146 |
| 35_NGAS596.crt.3 | HPLLMOOM_00615 hypothetical protein | 34 | 1863 | 63.9 | 7.00E-11 | 34 | 100% | 0 | 0% | 1 | 34 | 100% | 1146 | 1179 | + | 1146 |
| 35_NGAS596.crt.3 | MCFNMICF_01386 hypothetical protein | 34 | 1863 | 63.9 | 7.00E-11 | 34 | 100% | 0 | 0% | 1 | 34 | 100% | 1146 | 1179 | + | 1146 |
| 35_NGAS596.crt.4 | NOHIT | 33 | | | | | | | | | | | | | | |
| 35_NGAS596.crt.5 | NOHIT | 33 | | | | | | | | | | | | | | |
| 36_NGAS327.crt.1 | NOHIT | 30 | | | | | | | | | | | | | | |
| 36_NGAS327.crt.2 | NOHIT | 30 | | | | | | | | | | | | | | |
| 36_NGAS327.crt.3 | NOHIT | 30 | | | | | | | | | | | | | | |
| 36_NGAS327.crt.4 | NOHIT | 30 | | | | | | | | | | | | | | |
| 36_NGAS327.crt.5 | NOHIT | 30 | | | | | | | | | | | | | | |
| 36_NGAS327.crt.6 | NOHIT | 30 | | | | | | | | | | | | | | |
| 36_NGAS327.crt.7 | NOHIT | 30 | | | | | | | | | | | | | | |
| 36_NGAS327.crt.8 | NOHIT | 30 | | | | | | | | | | | | | | |
| 37_M28PF1.crt.1 | BNEKOFAL_00650 Transposase from transposon Tn9 | 35 | 1167 | 63.9 | 8.00E-11 | 34 | 100% | 0 | 0% | 2 | 35 | 97.14% | 1080 | 1113 | + | 1080 |
| 37_M28PF1.crt.1 | FOAEMPCG_01447 Transposase from transposon Tn9 | 35 | 1167 | 63.9 | 8.00E-11 | 34 | 100% | 0 | 0% | 2 | 35 | 97.14% | 1080 | 1113 | + | 1080 |
| 37_M28PF1.crt.1 | MCEIOEBL_00600 Transposase from transposon Tn91 | 35 | 1167 | 63.9 | 8.00E-11 | 34 | 100% | 0 | 0% | 2 | 35 | 97.14% | 1080 | 1113 | + | 1080 |
| 37_M28PF1.crt.1 | LLAAGEIN_01392 Transposase from transposon Tn91 | 35 | 1167 | 63.9 | 8.00E-11 | 34 | 100% | 0 | 0% | 2 | 35 | 97.14% | 1080 | 1113 | + | 1080 |
| 37_M28PF1.crt.1 | JNLHFPPN_01288 Transposase from transposon Tn91 | 35 | 1167 | 63.9 | 8.00E-11 | 34 | 100% | 0 | 0% | 2 | 35 | 97.14% | 1080 | 1113 | + | 1080 |
| 37_M28PF1.crt.1 | ODIAAKBJ_01288 Transposase from transposon Tn91 | 35 | 1167 | 63.9 | 8.00E-11 | 34 | 100% | 0 | 0% | 2 | 35 | 97.14% | 1080 | 1113 | + | 1080 |
| 37_M28PF1.crt.1 | ECKACGMC_00529 Transposase from transposon Tn | 35 | 1167 | 63.9 | 8.00E-11 | 34 | 100% | 0 | 0% | 2 | 35 | 97.14% | 1080 | 1113 | + | 1080 |
| 37_M28PF1.crt.1 | DIPDIGDN_01287 Transposase from transposon Tn91 | 35 | 1167 | 63.9 | 8.00E-11 | 34 | 100% | 0 | 0% | 2 | 35 | 97.14% | 1080 | 1113 | + | 1080 |
| 37_M28PF1.crt.1 | JHELIIPC_00570 Transposase from transposon Tn916 | 35 | 1167 | 63.9 | 8.00E-11 | 34 | 100% | 0 | 0% | 2 | 35 | 97.14% | 1080 | 1113 | + | 1080 |
| 37_M28PF1.crt.1 | AJHAHHEL_01339 Transposase from transposon Tn9 | 35 | 1167 | 63.9 | 8.00E-11 | 34 | 100% | 0 | 0% | 2 | 35 | 97.14% | 1080 | 1113 | + | 1080 |
| 37_M28PF1.crt.1 | HPLLMOOM_00568 Transposase from transposon Tn | 35 | 1167 | 63.9 | 8.00E-11 | 34 | 100% | 0 | 0% | 2 | 35 | 97.14% | 1080 | 1113 | + | 1080 |
| 37_M28PF1.crt.1 | MCFNMICF_01433 Transposase from transposon Tn9 | 35 | 1167 | 63.9 | 8.00E-11 | 34 | 100% | 0 | 0% | 2 | 35 | 97.14% | 1080 | 1113 | + | 1080 |
| 37_M28PF1.crt.1 | MONDJAPC_01605 Transposase from transposon Tn9 | 35 | 1167 | 63.9 | 8.00E-11 | 34 | 100% | 0 | 0% | 2 | 35 | 97.14% | 1080 | 1113 | + | 1080 |
| 37_M28PF1.crt.2 | NOHIT | 34 | | | | | | | | | | | | | | |
| 37_M28PF1.crt.3 | NOHIT | 0 | | | | | | | | | | | | | | |
| 37_M28PF1.crt.4 | NOHIT | 30 | | | | | | | | | | | | | | |
| 37_M28PF1.crt.5 | NOHIT | 30 | | | | | | | | | | | | | | |
| 37_M28PF1.crt.6 | NOHIT | 30 | | | | | | | | | | | | | | |
| 37_M28PF1.crt.7 | NOHIT | 30 | | | | | | | | | | | | | | |
| 38_5448.crt.1 | NOHIT | 35 | | | | | | | | | | | | | | |
| 38_5448.crt.2 | NOHIT | 33 | | | | | | | | | | | | | | |
| 38_5448.crt.3 | NOHIT | 34 | | | | | | | | | | | | | | |
| 38_5448.crt.4 | NOHIT | 33 | | | | | | | | | | | | | | |
| 38_5448.crt.5 | NOHIT | 0 | | | | | | | | | | | | | | |
| 38_5448.crt.6 | NOHIT | 30 | | | | | | | | | | | | | | |
| 38_5448.crt.7 | NOHIT | 30 | | | | | | | | | | | | | | |
| 38_5448.crt.8 | NOHIT | 30 | | | | | | | | | | | | | | |
| 39_STAB10015.crt.1 | NOHIT | 30 | | | | | | | | | | | | | | |
| 39_STAB10015.crt.2 | NOHIT | 30 | | | | | | | | | | | | | | |
| 39_STAB10015.crt.3 | NOHIT | 30 | | | | | | | | | | | | | | |
| 39_STAB10015.crt.4 | NOHIT | 30 | | | | | | | | | | | | | | |
| 39_STAB10015.crt.5 | NOHIT | 0 | | | | | | | | | | | | | | |
| 39_STAB10015.crt.6 | NOHIT | 34 | | | | | | | | | | | | | | |

| | | | | | | | | | | | | | | | | |
|---|---|---|---|---|---|---|---|---|---|---|---|---|---|---|---|---|
| 39_STAB10015.crt.7 | BNEKOFAL_00650 Transposase from transposon Tn9 | 35 | 1167 | 63.9 | 8.00E-11 | 34 | 100% | 0 | 0% | 1 | 34 | 97.14% | 1080 | 1113 | - | 1080 |
| 39_STAB10015.crt.7 | FOAEMPCG_01447 Transposase from transposon Tn9 | 35 | 1167 | 63.9 | 8.00E-11 | 34 | 100% | 0 | 0% | 1 | 34 | 97.14% | 1080 | 1113 | - | 1080 |
| 39_STAB10015.crt.7 | MCEIOEBL_00600 Transposase from transposon Tn91 | 35 | 1167 | 63.9 | 8.00E-11 | 34 | 100% | 0 | 0% | 1 | 34 | 97.14% | 1080 | 1113 | - | 1080 |
| 39_STAB10015.crt.7 | LLAAGEIN_01392 Transposase from transposon Tn91 | 35 | 1167 | 63.9 | 8.00E-11 | 34 | 100% | 0 | 0% | 1 | 34 | 97.14% | 1080 | 1113 | - | 1080 |
| 39_STAB10015.crt.7 | JNLHFPPN_01288 Transposase from transposon Tn91 | 35 | 1167 | 63.9 | 8.00E-11 | 34 | 100% | 0 | 0% | 1 | 34 | 97.14% | 1080 | 1113 | - | 1080 |
| 39_STAB10015.crt.7 | ODIAAKBJ_01288 Transposase from transposon Tn91 | 35 | 1167 | 63.9 | 8.00E-11 | 34 | 100% | 0 | 0% | 1 | 34 | 97.14% | 1080 | 1113 | - | 1080 |
| 39_STAB10015.crt.7 | ECKACGMC_00529 Transposase from transposon Tn9 | 35 | 1167 | 63.9 | 8.00E-11 | 34 | 100% | 0 | 0% | 1 | 34 | 97.14% | 1080 | 1113 | - | 1080 |
| 39_STAB10015.crt.7 | DIPDIGDN_01287 Transposase from transposon Tn91 | 35 | 1167 | 63.9 | 8.00E-11 | 34 | 100% | 0 | 0% | 1 | 34 | 97.14% | 1080 | 1113 | - | 1080 |
| 39_STAB10015.crt.7 | JHELIIPC_00570 Transposase from transposon Tn916 | 35 | 1167 | 63.9 | 8.00E-11 | 34 | 100% | 0 | 0% | 1 | 34 | 97.14% | 1080 | 1113 | - | 1080 |
| 39_STAB10015.crt.7 | AJHAHHEL_01339 Transposase from transposon Tn9 | 35 | 1167 | 63.9 | 8.00E-11 | 34 | 100% | 0 | 0% | 1 | 34 | 97.14% | 1080 | 1113 | - | 1080 |
| 39_STAB10015.crt.7 | HPLLMOOM_00568 Transposase from transposon Tn9 | 35 | 1167 | 63.9 | 8.00E-11 | 34 | 100% | 0 | 0% | 1 | 34 | 97.14% | 1080 | 1113 | - | 1080 |
| 39_STAB10015.crt.7 | MCFNMICF_01433 Transposase from transposon Tn9 | 35 | 1167 | 63.9 | 8.00E-11 | 34 | 100% | 0 | 0% | 1 | 34 | 97.14% | 1080 | 1113 | - | 1080 |
| 39_STAB10015.crt.7 | MONDJAPC_01605 Transposase from transposon Tn9 | 35 | 1167 | 63.9 | 8.00E-11 | 34 | 100% | 0 | 0% | 1 | 34 | 97.14% | 1080 | 1113 | - | 1080 |
| 40_HKU488.crt.1 | NOHIT | 30 | | | | | | | | | | | | | | |
| 40_HKU488.crt.2 | NOHIT | 30 | | | | | | | | | | | | | | |
| 40_HKU488.crt.3 | NOHIT | 30 | | | | | | | | | | | | | | |
| 40_HKU488.crt.4 | NOHIT | 0 | | | | | | | | | | | | | | |
| 40_HKU488.crt.5 | NOHIT | 33 | | | | | | | | | | | | | | |
| 40_HKU488.crt.6 | NOHIT | 34 | | | | | | | | | | | | | | |
| 40_HKU488.crt.7 | NOHIT | 33 | | | | | | | | | | | | | | |
| 40_HKU488.crt.8 | NOHIT | 35 | | | | | | | | | | | | | | |
| 41_NGAS322.crt.1 | NOHIT | 41 | | | | | | | | | | | | | | |
| 41_NGAS322.crt.2 | NOHIT | 41 | | | | | | | | | | | | | | |
| 41_NGAS322.crt.3 | NOHIT | 41 | | | | | | | | | | | | | | |
| 41_NGAS322.crt.4 | NOHIT | 41 | | | | | | | | | | | | | | |
| 41_NGAS322.crt.5 | NOHIT | 41 | | | | | | | | | | | | | | |
| 41_NGAS322.crt.6 | NOHIT | 41 | | | | | | | | | | | | | | |
| 42_NGAS638.crt.1 | NOHIT | 29 | | | | | | | | | | | | | | |
| 42_NGAS638.crt.2 | NOHIT | 29 | | | | | | | | | | | | | | |
| 42_NGAS638.crt.3 | NOHIT | 29 | | | | | | | | | | | | | | |
| 42_NGAS638.crt.4 | NOHIT | 29 | | | | | | | | | | | | | | |
| 43_MEW427.crt.1 | NOHIT | 33 | | | | | | | | | | | | | | |
| 43_MEW427.crt.2 | BNEKOFAL_00650 Transposase from transposon Tn9 | 34 | 1167 | 63.9 | 7.00E-11 | 34 | 100% | 0 | 0% | 1 | 34 | 100% | 1080 | 1113 | - | 1080 |
| 43_MEW427.crt.2 | FOAEMPCG_01447 Transposase from transposon Tn9 | 34 | 1167 | 63.9 | 7.00E-11 | 34 | 100% | 0 | 0% | 1 | 34 | 100% | 1080 | 1113 | - | 1080 |
| 43_MEW427.crt.2 | MCEIOEBL_00600 Transposase from transposon Tn91 | 34 | 1167 | 63.9 | 7.00E-11 | 34 | 100% | 0 | 0% | 1 | 34 | 100% | 1080 | 1113 | - | 1080 |
| 43_MEW427.crt.2 | LLAAGEIN_01392 Transposase from transposon Tn91 | 34 | 1167 | 63.9 | 7.00E-11 | 34 | 100% | 0 | 0% | 1 | 34 | 100% | 1080 | 1113 | - | 1080 |
| 43_MEW427.crt.2 | JNLHFPPN_01288 Transposase from transposon Tn91 | 34 | 1167 | 63.9 | 7.00E-11 | 34 | 100% | 0 | 0% | 1 | 34 | 100% | 1080 | 1113 | - | 1080 |
| 43_MEW427.crt.2 | ODIAAKBJ_01288 Transposase from transposon Tn91 | 34 | 1167 | 63.9 | 7.00E-11 | 34 | 100% | 0 | 0% | 1 | 34 | 100% | 1080 | 1113 | - | 1080 |
| 43_MEW427.crt.2 | ECKACGMC_00529 Transposase from transposon Tn9 | 34 | 1167 | 63.9 | 7.00E-11 | 34 | 100% | 0 | 0% | 1 | 34 | 100% | 1080 | 1113 | - | 1080 |
| 43_MEW427.crt.2 | DIPDIGDN_01287 Transposase from transposon Tn91 | 34 | 1167 | 63.9 | 7.00E-11 | 34 | 100% | 0 | 0% | 1 | 34 | 100% | 1080 | 1113 | - | 1080 |
| 43_MEW427.crt.2 | JHELIIPC_00570 Transposase from transposon Tn916 | 34 | 1167 | 63.9 | 7.00E-11 | 34 | 100% | 0 | 0% | 1 | 34 | 100% | 1080 | 1113 | - | 1080 |
| 43_MEW427.crt.2 | AJHAHHEL_01339 Transposase from transposon Tn9 | 34 | 1167 | 63.9 | 7.00E-11 | 34 | 100% | 0 | 0% | 1 | 34 | 100% | 1080 | 1113 | - | 1080 |
| 43_MEW427.crt.2 | HPLLMOOM_00568 Transposase from transposon Tn9 | 34 | 1167 | 63.9 | 7.00E-11 | 34 | 100% | 0 | 0% | 1 | 34 | 100% | 1080 | 1113 | - | 1080 |
| 43_MEW427.crt.2 | MCFNMICF_01433 Transposase from transposon Tn9 | 34 | 1167 | 63.9 | 7.00E-11 | 34 | 100% | 0 | 0% | 1 | 34 | 100% | 1080 | 1113 | - | 1080 |
| 43_MEW427.crt.2 | MONDJAPC_01605 Transposase from transposon Tn9 | 34 | 1167 | 63.9 | 7.00E-11 | 34 | 100% | 0 | 0% | 1 | 34 | 100% | 1080 | 1113 | - | 1080 |
| 43_MEW427.crt.3 | JEOJEKGO_01300 hypothetical protein | 35 | 513 | 65.8 | 2.00E-11 | 35 | 100% | 0 | 0% | 1 | 35 | 100% | 246 | 280 | + | 246 |
| 43_MEW427.crt.3 | MONDJAPC_01319 hypothetical protein | 35 | 513 | 65.8 | 2.00E-11 | 35 | 100% | 0 | 0% | 1 | 35 | 100% | 246 | 280 | + | 246 |
| 43_MEW427.crt.4 | NOHIT | 33 | | | | | | | | | | | | | | |
| 43_MEW427.crt.5 | NOHIT | 34 | | | | | | | | | | | | | | |
| 44_MEW123.crt.1 | BNEKOFAL_00650 Transposase from transposon Tn9 | 35 | 1167 | 63.9 | 8.00E-11 | 34 | 100% | 0 | 0% | 2 | 35 | 97.14% | 1080 | 1113 | + | 1080 |
| 44_MEW123.crt.1 | FOAEMPCG_01447 Transposase from transposon Tn9 | 35 | 1167 | 63.9 | 8.00E-11 | 34 | 100% | 0 | 0% | 2 | 35 | 97.14% | 1080 | 1113 | + | 1080 |
| 44_MEW123.crt.1 | MCEIOEBL_00600 Transposase from transposon Tn91 | 35 | 1167 | 63.9 | 8.00E-11 | 34 | 100% | 0 | 0% | 2 | 35 | 97.14% | 1080 | 1113 | + | 1080 |
| 44_MEW123.crt.1 | LLAAGEIN_01392 Transposase from transposon Tn91 | 35 | 1167 | 63.9 | 8.00E-11 | 34 | 100% | 0 | 0% | 2 | 35 | 97.14% | 1080 | 1113 | + | 1080 |
| 44_MEW123.crt.1 | JNLHFPPN_01288 Transposase from transposon Tn91 | 35 | 1167 | 63.9 | 8.00E-11 | 34 | 100% | 0 | 0% | 2 | 35 | 97.14% | 1080 | 1113 | + | 1080 |
| 44_MEW123.crt.1 | ODIAAKBJ_01288 Transposase from transposon Tn91 | 35 | 1167 | 63.9 | 8.00E-11 | 34 | 100% | 0 | 0% | 2 | 35 | 97.14% | 1080 | 1113 | + | 1080 |
| 44_MEW123.crt.1 | ECKACGMC_00529 Transposase from transposon Tn9 | 35 | 1167 | 63.9 | 8.00E-11 | 34 | 100% | 0 | 0% | 2 | 35 | 97.14% | 1080 | 1113 | + | 1080 |
| 44_MEW123.crt.1 | DIPDIGDN_01287 Transposase from transposon Tn91 | 35 | 1167 | 63.9 | 8.00E-11 | 34 | 100% | 0 | 0% | 2 | 35 | 97.14% | 1080 | 1113 | + | 1080 |
| 44_MEW123.crt.1 | JHELIIPC_00570 Transposase from transposon Tn916 | 35 | 1167 | 63.9 | 8.00E-11 | 34 | 100% | 0 | 0% | 2 | 35 | 97.14% | 1080 | 1113 | + | 1080 |
| 44_MEW123.crt.1 | AJHAHHEL_01339 Transposase from transposon Tn9 | 35 | 1167 | 63.9 | 8.00E-11 | 34 | 100% | 0 | 0% | 2 | 35 | 97.14% | 1080 | 1113 | + | 1080 |
| 44_MEW123.crt.1 | HPLLMOOM_00568 Transposase from transposon Tn9 | 35 | 1167 | 63.9 | 8.00E-11 | 34 | 100% | 0 | 0% | 2 | 35 | 97.14% | 1080 | 1113 | + | 1080 |
| 44_MEW123.crt.1 | MCFNMICF_01433 Transposase from transposon Tn9 | 35 | 1167 | 63.9 | 8.00E-11 | 34 | 100% | 0 | 0% | 2 | 35 | 97.14% | 1080 | 1113 | + | 1080 |
| 44_MEW123.crt.1 | MONDJAPC_01605 Transposase from transposon Tn9 | 35 | 1167 | 63.9 | 8.00E-11 | 34 | 100% | 0 | 0% | 2 | 35 | 97.14% | 1080 | 1113 | + | 1080 |
| 44_MEW123.crt.2 | NOHIT | 34 | | | | | | | | | | | | | | |
| 44_MEW123.crt.3 | NOHIT | 0 | | | | | | | | | | | | | | |
| 44_MEW123.crt.4 | NOHIT | 31 | | | | | | | | | | | | | | |
| 44_MEW123.crt.5 | NOHIT | 30 | | | | | | | | | | | | | | |
| 44_MEW123.crt.6 | NOHIT | 30 | | | | | | | | | | | | | | |
| 45_FDAARGOS_149.crt.1 | NOHIT | 35 | | | | | | | | | | | | | | |
| 45_FDAARGOS_149.crt.2 | NOHIT | 33 | | | | | | | | | | | | | | |
| 45_FDAARGOS_149.crt.3 | NOHIT | 34 | | | | | | | | | | | | | | |
| 45_FDAARGOS_149.crt.4 | NOHIT | 33 | | | | | | | | | | | | | | |
| 45_FDAARGOS_149.crt.5 | NOHIT | 0 | | | | | | | | | | | | | | |
| 45_FDAARGOS_149.crt.6 | NOHIT | 30 | | | | | | | | | | | | | | |
| 45_FDAARGOS_149.crt.7 | NOHIT | 30 | | | | | | | | | | | | | | |
| 45_FDAARGOS_149.crt.8 | NOHIT | 30 | | | | | | | | | | | | | | |
| 47_MGAS11027.crt.1 | NOHIT | 30 | | | | | | | | | | | | | | |
| 47_MGAS11027.crt.2 | NOHIT | 30 | | | | | | | | | | | | | | |
| 47_MGAS11027.crt.3 | NOHIT | 30 | | | | | | | | | | | | | | |
| 47_MGAS11027.crt.4 | NOHIT | 30 | | | | | | | | | | | | | | |
| 47_MGAS11027.crt.5 | NOHIT | 30 | | | | | | | | | | | | | | |
| 47_MGAS11027.crt.6 | NOHIT | 30 | | | | | | | | | | | | | | |
| 47_MGAS11027.crt.7 | NOHIT | 30 | | | | | | | | | | | | | | |
| 47_MGAS11027.crt.8 | NOHIT | 0 | | | | | | | | | | | | | | |
| 47_MGAS11027.crt.9 | NOHIT | 34 | | | | | | | | | | | | | | |
| 47_MGAS11027.crt.10 | NOHIT | 34 | | | | | | | | | | | | | | |
| 47_MGAS11027.crt.11 | GGADKFFH_01023 putative DNA primase - phage as | 37 | 2385 | 63.9 | 9.00E-11 | 34 | 100% | 0 | 0% | 4 | 37 | 91.89% | 1638 | 1671 | + | 1638 |
| 47_MGAS11027.crt.11 | JNLHFPPN_01503 putative DNA primase - phage asso | 37 | 2385 | 63.9 | 9.00E-11 | 34 | 100% | 0 | 0% | 4 | 37 | 91.89% | 1638 | 1671 | + | 1638 |
| 47_MGAS11027.crt.11 | ODIAAKBJ_01503 putative DNA primase - phage asso | 37 | 2385 | 63.9 | 9.00E-11 | 34 | 100% | 0 | 0% | 4 | 37 | 91.89% | 1638 | 1671 | + | 1638 |
| 47_MGAS11027.crt.11 | DIPDIGDN_01504 putative DNA primase - phage asso | 37 | 2385 | 63.9 | 9.00E-11 | 34 | 100% | 0 | 0% | 4 | 37 | 91.89% | 1638 | 1671 | + | 1638 |
| 47_MGAS11027.crt.11 | MDEAFGPO_00463 putative DNA primase - phage as | 37 | 2385 | 63.9 | 9.00E-11 | 34 | 100% | 0 | 0% | 4 | 37 | 91.89% | 1638 | 1671 | + | 1638 |
| 47_MGAS11027.crt.11 | DMLKMGFL_00783 putative DNA primase - phage as | 37 | 2385 | 63.9 | 9.00E-11 | 34 | 100% | 0 | 0% | 4 | 37 | 91.89% | 1638 | 1671 | + | 1638 |
| 47_MGAS11027.crt.11 | HEGDFFNC_01101 putative DNA primase - phage ass | 37 | 2385 | 63.9 | 9.00E-11 | 34 | 100% | 0 | 0% | 4 | 37 | 91.89% | 1638 | 1671 | + | 1638 |
| 47_MGAS11027.crt.12 | NOHIT | 35 | | | | | | | | | | | | | | |
| 47_MGAS11027.crt.13 | MHKCMMMI_00605 hypothetical protein | 34 | 678 | 63.9 | 7.00E-11 | 34 | 100% | 0 | 0% | 1 | 34 | 100% | 593 | 626 | - | 593 |
| 47_MGAS11027.crt.13 | FOAEMPCG_01323 hypothetical protein | 34 | 678 | 63.9 | 7.00E-11 | 34 | 100% | 0 | 0% | 1 | 34 | 100% | 593 | 626 | - | 593 |
| 47_MGAS11027.crt.13 | MCEIOEBL_00630 hypothetical protein | 34 | 678 | 63.9 | 7.00E-11 | 34 | 100% | 0 | 0% | 1 | 34 | 100% | 593 | 626 | - | 593 |
| 47_MGAS11027.crt.13 | LAKBNEPL_01310 hypothetical protein | 34 | 678 | 63.9 | 7.00E-11 | 34 | 100% | 0 | 0% | 1 | 34 | 100% | 593 | 626 | - | 593 |
| 47_MGAS11027.crt.13 | PLPKHLCC_00644 hypothetical protein | 34 | 678 | 63.9 | 7.00E-11 | 34 | 100% | 0 | 0% | 1 | 34 | 100% | 593 | 626 | - | 593 |
| 47_MGAS11027.crt.13 | DCKFNFNP_01264 hypothetical protein | 34 | 678 | 63.9 | 7.00E-11 | 34 | 100% | 0 | 0% | 1 | 34 | 100% | 593 | 626 | - | 593 |
| 47_MGAS11027.crt.13 | JNLHFPPN_01257 hypothetical protein | 34 | 678 | 63.9 | 7.00E-11 | 34 | 100% | 0 | 0% | 1 | 34 | 100% | 593 | 626 | - | 593 |
| 47_MGAS11027.crt.13 | ODIAAKBJ_01257 hypothetical protein | 34 | 678 | 63.9 | 7.00E-11 | 34 | 100% | 0 | 0% | 1 | 34 | 100% | 593 | 626 | - | 593 |
| 47_MGAS11027.crt.13 | DIPDIGDN_01256 hypothetical protein | 34 | 678 | 63.9 | 7.00E-11 | 34 | 100% | 0 | 0% | 1 | 34 | 100% | 593 | 626 | - | 593 |
| 47_MGAS11027.crt.13 | JHELIIPC_00693 hypothetical protein | 34 | 678 | 63.9 | 7.00E-11 | 34 | 100% | 0 | 0% | 1 | 34 | 100% | 593 | 626 | - | 593 |
| 47_MGAS11027.crt.13 | IPBLJGMA_01319 hypothetical protein | 34 | 678 | 63.9 | 7.00E-11 | 34 | 100% | 0 | 0% | 1 | 34 | 100% | 593 | 626 | - | 593 |
| 47_MGAS11027.crt.13 | GLBJHDNN_01264 hypothetical protein | 34 | 678 | 63.9 | 7.00E-11 | 34 | 100% | 0 | 0% | 1 | 34 | 100% | 593 | 626 | - | 593 |
| 47_MGAS11027.crt.13 | HPLLMOOM_00691 hypothetical protein | 34 | 678 | 63.9 | 7.00E-11 | 34 | 100% | 0 | 0% | 1 | 34 | 100% | 593 | 626 | - | 593 |
| 47_MGAS11027.crt.13 | MCFNMICF_01310 hypothetical protein | 34 | 678 | 63.9 | 7.00E-11 | 34 | 100% | 0 | 0% | 1 | 34 | 100% | 593 | 626 | - | 593 |
| 47_MGAS11027.crt.13 | MONDJAPC_01575 hypothetical protein | 34 | 678 | 63.9 | 7.00E-11 | 34 | 100% | 0 | 0% | 1 | 34 | 100% | 593 | 626 | - | 593 |
| 47_MGAS11027.crt.14 | NOHIT | 35 | | | | | | | | | | | | | | |
| 47_MGAS11027.crt.15 | NOHIT | 35 | | | | | | | | | | | | | | |
| 47_MGAS11027.crt.16 | NOHIT | 33 | | | | | | | | | | | | | | |
| 47_MGAS11027.crt.17 | BNEKOFAL_00851 hypothetical protein | 36 | 1782 | 67.6 | 6.00E-12 | 36 | 100% | 0 | 0% | 1 | 36 | 100% | 292 | 327 | - | 292 |
| 47_MGAS11027.crt.17 | MHKCMMMI_00742 hypothetical protein | 36 | 1782 | 67.6 | 6.00E-12 | 36 | 100% | 0 | 0% | 1 | 36 | 100% | 292 | 327 | - | 292 |
| 47_MGAS11027.crt.17 | FOAEMPCG_01185 hypothetical protein | 36 | 1782 | 67.6 | 6.00E-12 | 36 | 100% | 0 | 0% | 1 | 36 | 100% | 292 | 327 | - | 292 |
| 47_MGAS11027.crt.17 | MCEIOEBL_00803 hypothetical protein | 36 | 1782 | 67.6 | 6.00E-12 | 36 | 100% | 0 | 0% | 1 | 36 | 100% | 292 | 327 | - | 292 |
| 47_MGAS11027.crt.17 | LLAAGEIN_01209 hypothetical protein | 36 | 1782 | 67.6 | 6.00E-12 | 36 | 100% | 0 | 0% | 1 | 36 | 100% | 292 | 327 | - | 292 |
| 47_MGAS11027.crt.17 | AIKJPEBD_01431 hypothetical protein | 36 | 1782 | 67.6 | 6.00E-12 | 36 | 100% | 0 | 0% | 1 | 36 | 100% | 292 | 327 | - | 292 |
| 47_MGAS11027.crt.17 | FDNDANMJ_01220 hypothetical protein | 36 | 1782 | 67.6 | 6.00E-12 | 36 | 100% | 0 | 0% | 1 | 36 | 100% | 292 | 327 | - | 292 |
| 47_MGAS11027.crt.17 | JEOJEKGO_01273 hypothetical protein | 36 | 1782 | 67.6 | 6.00E-12 | 36 | 100% | 0 | 0% | 1 | 36 | 100% | 292 | 327 | - | 292 |
| 47_MGAS11027.crt.17 | EJBPJIHF_01240 hypothetical protein | 36 | 1869 | 67.6 | 6.00E-12 | 36 | 100% | 0 | 0% | 1 | 36 | 100% | 376 | 411 | - | 376 |
| 47_MGAS11027.crt.17 | MMBCLDPD_00722 hypothetical protein | 36 | 1869 | 67.6 | 6.00E-12 | 36 | 100% | 0 | 0% | 1 | 36 | 100% | 376 | 411 | - | 376 |
| 47_MGAS11027.crt.17 | DCKFNFNP_00615 hypothetical protein | 36 | 1782 | 67.6 | 6.00E-12 | 36 | 100% | 0 | 0% | 1 | 36 | 100% | 292 | 327 | - | 292 |
| 47_MGAS11027.crt.17 | JNLHFPPN_01086 hypothetical protein | 36 | 1782 | 67.6 | 6.00E-12 | 36 | 100% | 0 | 0% | 1 | 36 | 100% | 292 | 327 | - | 292 |
| 47_MGAS11027.crt.17 | ODIAAKBJ_01086 hypothetical protein | 36 | 1782 | 67.6 | 6.00E-12 | 36 | 100% | 0 | 0% | 1 | 36 | 100% | 292 | 327 | - | 292 |
| 47_MGAS11027.crt.17 | ECKACGMC_00725 hypothetical protein | 36 | 1785 | 67.6 | 6.00E-12 | 36 | 100% | 0 | 0% | 1 | 36 | 100% | 292 | 327 | - | 292 |
| 47_MGAS11027.crt.17 | DIPDIGDN_01085 hypothetical protein | 36 | 1782 | 67.6 | 6.00E-12 | 36 | 100% | 0 | 0% | 1 | 36 | 100% | 292 | 327 | - | 292 |
| 47_MGAS11027.crt.17 | JHELIIPC_00832 hypothetical protein | 36 | 1782 | 67.6 | 6.00E-12 | 36 | 100% | 0 | 0% | 1 | 36 | 100% | 292 | 327 | - | 292 |
| 47_MGAS11027.crt.17 | GPBJDOFB_01147 hypothetical protein | 36 | 1785 | 67.6 | 6.00E-12 | 36 | 100% | 0 | 0% | 1 | 36 | 100% | 292 | 327 | - | 292 |
| 47_MGAS11027.crt.17 | ACJECOPP_01164 hypothetical protein | 36 | 1869 | 67.6 | 6.00E-12 | 36 | 100% | 0 | 0% | 1 | 36 | 100% | 376 | 411 | - | 376 |
| 47_MGAS11027.crt.17 | ICMALKPE_01176 hypothetical protein | 36 | 1869 | 67.6 | 6.00E-12 | 36 | 100% | 0 | 0% | 1 | 36 | 100% | 376 | 411 | - | 376 |
| 47_MGAS11027.crt.17 | AJHAHHEL_01155 hypothetical protein | 36 | 1782 | 67.6 | 6.00E-12 | 36 | 100% | 0 | 0% | 1 | 36 | 100% | 292 | 327 | - | 292 |
| 47_MGAS11027.crt.17 | HEGDFFNC_00730 hypothetical protein | 36 | 1785 | 67.6 | 6.00E-12 | 36 | 100% | 0 | 0% | 1 | 36 | 100% | 292 | 327 | - | 292 |
| 47_MGAS11027.crt.17 | IPNPMFKD_01282 hypothetical protein | 36 | 1782 | 67.6 | 6.00E-12 | 36 | 100% | 0 | 0% | 1 | 36 | 100% | 292 | 327 | - | 292 |
| 47_MGAS11027.crt.17 | IIJJCGIL_01167 hypothetical protein | 36 | 1869 | 67.6 | 6.00E-12 | 36 | 100% | 0 | 0% | 1 | 36 | 100% | 376 | 411 | - | 376 |
| 47_MGAS11027.crt.17 | NELECJBE_01235 hypothetical protein | 36 | 1782 | 67.6 | 6.00E-12 | 36 | 100% | 0 | 0% | 1 | 36 | 100% | 292 | 327 | - | 292 |
| 47_MGAS11027.crt.17 | HPLLMOOM_00829 hypothetical protein | 36 | 1782 | 67.6 | 6.00E-12 | 36 | 100% | 0 | 0% | 1 | 36 | 100% | 292 | 327 | - | 292 |
| 47_MGAS11027.crt.17 | MCFNMICF_01172 hypothetical protein | 36 | 1782 | 67.6 | 6.00E-12 | 36 | 100% | 0 | 0% | 1 | 36 | 100% | 292 | 327 | - | 292 |
| 47_MGAS11027.crt.17 | MONDJAPC_01286 hypothetical protein | 36 | 1785 | 67.6 | 6.00E-12 | 36 | 100% | 0 | 0% | 1 | 36 | 100% | 292 | 327 | - | 292 |
| 47_MGAS11027.crt.17 | EEABGJJD_01210 hypothetical protein | 36 | 1785 | 67.6 | 6.00E-12 | 36 | 100% | 0 | 0% | 1 | 36 | 100% | 292 | 327 | - | 292 |
| 47_MGAS11027.crt.18 | NOHIT | 34 | | | | | | | | | | | | | | |
| 47_MGAS11027.crt.19 | NOHIT | 37 | | | | | | | | | | | | | | |
| 47_MGAS11027.crt.20 | NOHIT | 33 | | | | | | | | | | | | | | |
| 47_MGAS11027.crt.21 | BGBHDJOA_00791 putative repressor protein - phage | 34 | 756 | 63.9 | 7.00E-11 | 34 | 100% | 0 | 0% | 1 | 34 | 100% | 687 | 720 | + | 687 |
| 47_MGAS11027.crt.21 | EJBPJIHF_01687 putative repressor protein - phage as | 34 | 759 | 63.9 | 7.00E-11 | 34 | 100% | 0 | 0% | 1 | 34 | 100% | 696 | 729 | + | 696 |
| 47_MGAS11027.crt.21 | GBCACICN_01644 putative repressor protein - phage | 34 | 759 | 63.9 | 7.00E-11 | 34 | 100% | 0 | 0% | 1 | 34 | 100% | 696 | 729 | + | 696 |
| 47_MGAS11027.crt.21 | GBCACICN_00736 putative repressor protein - phage | 34 | 756 | 63.9 | 7.00E-11 | 34 | 100% | 0 | 0% | 1 | 34 | 100% | 687 | 720 | + | 687 |
| 47_MGAS11027.crt.21 | EFGBCCNL_00740 putative repressor protein - phage | 34 | 759 | 63.9 | 7.00E-11 | 34 | 100% | 0 | 0% | 1 | 34 | 100% | 696 | 729 | + | 696 |
| 47_MGAS11027.crt.21 | DMLKMGFL_01526 putative repressor protein - phag | 34 | 759 | 63.9 | 7.00E-11 | 34 | 100% | 0 | 0% | 1 | 34 | 100% | 696 | 729 | + | 696 |
| 47_MGAS11027.crt.21 | IPNPMFKD_00585 putative repressor protein - phage | 34 | 750 | 63.9 | 7.00E-11 | 34 | 100% | 0 | 0% | 1 | 34 | 100% | 687 | 720 | + | 687 |
| 47_MGAS11027.crt.21 | IPBLJGMA_00554 putative repressor protein - phage | 34 | 750 | 63.9 | 7.00E-11 | 34 | 100% | 0 | 0% | 1 | 34 | 100% | 687 | 720 | + | 687 |
| 47_MGAS11027.crt.21 | ELLIANMJ_00803 putative repressor protein - phage | 34 | 756 | 63.9 | 7.00E-11 | 34 | 100% | 0 | 0% | 1 | 34 | 100% | 687 | 720 | + | 687 |
| 47_MGAS11027.crt.21 | ELLIANMJ_00546 putative repressor protein - phage | 34 | 750 | 63.9 | 7.00E-11 | 34 | 100% | 0 | 0% | 1 | 34 | 100% | 687 | 720 | + | 687 |
| 48_MGAS23530.crt.1 | NOHIT | 30 | | | | | | | | | | | | | | |
| 48_MGAS23530.crt.2 | NOHIT | 30 | | | | | | | | | | | | | | |
| 48_MGAS23530.crt.3 | NOHIT | 30 | | | | | | | | | | | | | | |
| 48_MGAS23530.crt.4 | NOHIT | 30 | | | | | | | | | | | | | | |
| 48_MGAS23530.crt.5 | NOHIT | 30 | | | | | | | | | | | | | | |
| 48_MGAS23530.crt.6 | NOHIT | 0 | | | | | | | | | | | | | | |
| 48_MGAS23530.crt.7 | NOHIT | 19 | | | | | | | | | | | | | | |
| 48_MGAS23530.crt.8 | NOHIT | 28 | | | | | | | | | | | | | | |
| 48_MGAS23530.crt.9 | NOHIT | 19 | | | | | | | | | | | | | | |
| 48_MGAS23530.crt.10 | NOHIT | 19 | | | | | | | | | | | | | | |
| 48_MGAS23530.crt.11 | NOHIT | 0 | | | | | | | | | | | | | | |
| 48_MGAS23530.crt.12 | NOHIT | 35 | | | | | | | | | | | | | | |
| 48_MGAS23530.crt.13 | NOHIT | 31 | | | | | | | | | | | | | | |
| 48_MGAS23530.crt.14 | NOHIT | 32 | | | | | | | | | | | | | | |
| 49_MGAS27061.crt.1 | NOHIT | 30 | | | | | | | | | | | | | | |
| 49_MGAS27061.crt.2 | NOHIT | 30 | | | | | | | | | | | | | | |
| 49_MGAS27061.crt.3 | NOHIT | 30 | | | | | | | | | | | | | | |
| 49_MGAS27061.crt.4 | NOHIT | 30 | | | | | | | | | | | | | | |
| 49_MGAS27061.crt.5 | NOHIT | 0 | | | | | | | | | | | | | | |
| 49_MGAS27061.crt.6 | NOHIT | 19 | | | | | | | | | | | | | | |
| 49_MGAS27061.crt.7 | NOHIT | 28 | | | | | | | | | | | | | | |
| 49_MGAS27061.crt.8 | NOHIT | 19 | | | | | | | | | | | | | | |
| 49_MGAS27061.crt.9 | NOHIT | 19 | | | | | | | | | | | | | | |
| 49_MGAS27061.crt.10 | NOHIT | 0 | | | | | | | | | | | | | | |
| 49_MGAS27061.crt.11 | NOHIT | 33 | | | | | | | | | | | | | | |
| 49_MGAS27061.crt.12 | NOHIT | 33 | | | | | | | | | | | | | | |
| 49_MGAS27061.crt.13 | NOHIT | 36 | | | | | | | | | | | | | | |
| 49_MGAS27061.crt.14 | NOHIT | 33 | | | | | | | | | | | | | | |
| 49_MGAS27061.crt.15 | NOHIT | 34 | | | | | | | | | | | | | | |
| 49_MGAS27061.crt.16 | NOHIT | 33 | | | | | | | | | | | | | | |
| 49_MGAS27061.crt.17 | NOHIT | 32 | | | | | | | | | | | | | | |
| 49_MGAS27061.crt.18 | NOHIT | 33 | | | | | | | | | | | | | | |
| 51_STAB09014.crt.1 | NOHIT | 30 | | | | | | | | | | | | | | |
| 51_STAB09014.crt.2 | NOHIT | 30 | | | | | | | | | | | | | | |
| 51_STAB09014.crt.3 | NOHIT | 30 | | | | | | | | | | | | | | |
| 51_STAB09014.crt.4 | NOHIT | 30 | | | | | | | | | | | | | | |
| 52_ATCC14918.crt.1 | NOHIT | 32 | | | | | | | | | | | | | | |
| 52_ATCC14918.crt.2 | BNEKOFAL_00674 hypothetical protein | 34 | 285 | 63.9 | 7.00E-11 | 34 | 100% | 0 | 0% | 1 | 34 | 100% | 19 | 52 | + | 19 |
| 52_ATCC14918.crt.2 | FOAEMPCG_01040 hypothetical protein | 34 | 285 | 63.9 | 7.00E-11 | 34 | 100% | 0 | 0% | 1 | 34 | 100% | 19 | 52 | + | 19 |
| 52_ATCC14918.crt.2 | MCEIOEBL_00624 hypothetical protein | 34 | 285 | 63.9 | 7.00E-11 | 34 | 100% | 0 | 0% | 1 | 34 | 100% | 19 | 52 | + | 19 |
| 52_ATCC14918.crt.2 | PPKGMMPD_01012 hypothetical protein | 34 | 285 | 63.9 | 7.00E-11 | 34 | 100% | 0 | 0% | 1 | 34 | 100% | 19 | 52 | + | 19 |
| 52_ATCC14918.crt.2 | LLAAGEIN_01368 hypothetical protein | 34 | 285 | 63.9 | 7.00E-11 | 34 | 100% | 0 | 0% | 1 | 34 | 100% | 19 | 52 | + | 19 |
| 52_ATCC14918.crt.2 | AIKJPEBD_01573 hypothetical protein | 34 | 285 | 63.9 | 7.00E-11 | 34 | 100% | 0 | 0% | 1 | 34 | 100% | 19 | 52 | + | 19 |
| 52_ATCC14918.crt.2 | FPFNMMFM_00882 hypothetical protein | 34 | 285 | 63.9 | 7.00E-11 | 34 | 100% | 0 | 0% | 1 | 34 | 100% | 19 | 52 | + | 19 |
| 52_ATCC14918.crt.2 | IJFGMCCO_00995 hypothetical protein | 34 | 285 | 63.9 | 7.00E-11 | 34 | 100% | 0 | 0% | 1 | 34 | 100% | 19 | 52 | + | 19 |
| 52_ATCC14918.crt.2 | EJBPJIHF_01045 hypothetical protein | 34 | 285 | 63.9 | 7.00E-11 | 34 | 100% | 0 | 0% | 1 | 34 | 100% | 19 | 52 | + | 19 |
| 52_ATCC14918.crt.2 | LAKBNEPL_01011 hypothetical protein | 34 | 285 | 63.9 | 7.00E-11 | 34 | 100% | 0 | 0% | 1 | 34 | 100% | 19 | 52 | + | 19 |
| 52_ATCC14918.crt.2 | MMBCLDPD_00865 hypothetical protein | 34 | 285 | 63.9 | 7.00E-11 | 34 | 100% | 0 | 0% | 1 | 34 | 100% | 19 | 52 | + | 19 |
| 52_ATCC14918.crt.2 | PLPKHLCC_00893 hypothetical protein | 34 | 285 | 63.9 | 7.00E-11 | 34 | 100% | 0 | 0% | 1 | 34 | 100% | 19 | 52 | + | 19 |
| 52_ATCC14918.crt.2 | JNLHFPPN_01263 hypothetical protein | 34 | 285 | 63.9 | 7.00E-11 | 34 | 100% | 0 | 0% | 1 | 34 | 100% | 19 | 52 | + | 19 |
| 52_ATCC14918.crt.2 | ODIAAKBJ_01263 hypothetical protein | 34 | 285 | 63.9 | 7.00E-11 | 34 | 100% | 0 | 0% | 1 | 34 | 100% | 19 | 52 | + | 19 |
| 52_ATCC14918.crt.2 | ECKACGMC_00553 hypothetical protein | 34 | 285 | 63.9 | 7.00E-11 | 34 | 100% | 0 | 0% | 1 | 34 | 100% | 19 | 52 | + | 19 |
| 52_ATCC14918.crt.2 | DIPDIGDN_01262 hypothetical protein | 34 | 285 | 63.9 | 7.00E-11 | 34 | 100% | 0 | 0% | 1 | 34 | 100% | 19 | 52 | + | 19 |
| 52_ATCC14918.crt.2 | EFGBCCNL_00703 hypothetical protein | 34 | 285 | 63.9 | 7.00E-11 | 34 | 100% | 0 | 0% | 1 | 34 | 100% | 19 | 52 | + | 19 |
| 52_ATCC14918.crt.2 | JHELIIPC_00975 hypothetical protein | 34 | 285 | 63.9 | 7.00E-11 | 34 | 100% | 0 | 0% | 1 | 34 | 100% | 19 | 52 | + | 19 |
| 52_ATCC14918.crt.2 | GPBJDOFB_00997 hypothetical protein | 34 | 285 | 63.9 | 7.00E-11 | 34 | 100% | 0 | 0% | 1 | 34 | 100% | 19 | 52 | + | 19 |
| 52_ATCC14918.crt.2 | ACJECOPP_01021 hypothetical protein | 34 | 285 | 63.9 | 7.00E-11 | 34 | 100% | 0 | 0% | 1 | 34 | 100% | 19 | 52 | + | 19 |
| 52_ATCC14918.crt.2 | ICMALKPE_01033 hypothetical protein | 34 | 285 | 63.9 | 7.00E-11 | 34 | 100% | 0 | 0% | 1 | 34 | 100% | 19 | 52 | + | 19 |
| 52_ATCC14918.crt.2 | AJHAHHEL_01315 hypothetical protein | 34 | 285 | 63.9 | 7.00E-11 | 34 | 100% | 0 | 0% | 1 | 34 | 100% | 19 | 52 | + | 19 |
| 52_ATCC14918.crt.2 | DMLKMGFL_01490 hypothetical protein | 34 | 285 | 63.9 | 7.00E-11 | 34 | 100% | 0 | 0% | 1 | 34 | 100% | 19 | 52 | + | 19 |
| 52_ATCC14918.crt.2 | HEGDFFNC_00554 hypothetical protein | 34 | 285 | 63.9 | 7.00E-11 | 34 | 100% | 0 | 0% | 1 | 34 | 100% | 19 | 52 | + | 19 |
| 52_ATCC14918.crt.2 | IIJJCGIL_01024 hypothetical protein | 34 | 285 | 63.9 | 7.00E-11 | 34 | 100% | 0 | 0% | 1 | 34 | 100% | 19 | 52 | + | 19 |
| 52_ATCC14918.crt.2 | GLBJHDNN_01015 hypothetical protein | 34 | 285 | 63.9 | 7.00E-11 | 34 | 100% | 0 | 0% | 1 | 34 | 100% | 19 | 52 | + | 19 |
| 52_ATCC14918.crt.2 | HPLLMOOM_00975 hypothetical protein | 34 | 285 | 63.9 | 7.00E-11 | 34 | 100% | 0 | 0% | 1 | 34 | 100% | 19 | 52 | + | 19 |
| 52_ATCC14918.crt.2 | MCFNMICF_01027 hypothetical protein | 34 | 285 | 63.9 | 7.00E-11 | 34 | 100% | 0 | 0% | 1 | 34 | 100% | 19 | 52 | + | 19 |
| 52_ATCC14918.crt.2 | MONDJAPC_01581 hypothetical protein | 34 | 285 | 63.9 | 7.00E-11 | 34 | 100% | 0 | 0% | 1 | 34 | 100% | 19 | 52 | + | 19 |
| 52_ATCC14918.crt.3 | BNEKOFAL_01130 hypothetical protein | 34 | 186 | 63.9 | 7.00E-11 | 34 | 100% | 0 | 0% | 1 | 34 | 100% | 150 | 183 | + | 150 |
| 52_ATCC14918.crt.3 | FOAEMPCG_01422 hypothetical protein | 34 | 186 | 63.9 | 7.00E-11 | 34 | 100% | 0 | 0% | 1 | 34 | 100% | 150 | 183 | + | 150 |
| 52_ATCC14918.crt.3 | MCEIOEBL_01079 hypothetical protein | 34 | 186 | 63.9 | 7.00E-11 | 34 | 100% | 0 | 0% | 1 | 34 | 100% | 150 | 183 | + | 150 |
| 52_ATCC14918.crt.3 | LLAAGEIN_01062 hypothetical protein | 34 | 186 | 63.9 | 7.00E-11 | 34 | 100% | 0 | 0% | 1 | 34 | 100% | 150 | 183 | + | 150 |
| 52_ATCC14918.crt.3 | IJFGMCCO_00990 hypothetical protein | 34 | 186 | 63.9 | 7.00E-11 | 34 | 100% | 0 | 0% | 1 | 34 | 100% | 150 | 183 | + | 150 |
| 52_ATCC14918.crt.3 | MDEAFGPO_00646 hypothetical protein | 34 | 186 | 63.9 | 7.00E-11 | 34 | 100% | 0 | 0% | 1 | 34 | 100% | 150 | 183 | + | 150 |
| 52_ATCC14918.crt.3 | MDEAFGPO_00220 hypothetical protein | 34 | 186 | 63.9 | 7.00E-11 | 34 | 100% | 0 | 0% | 1 | 34 | 100% | 150 | 183 | + | 150 |
| 52_ATCC14918.crt.3 | EFGBCCNL_00701 hypothetical protein | 34 | 186 | 63.9 | 7.00E-11 | 34 | 100% | 0 | 0% | 1 | 34 | 100% | 150 | 183 | + | 150 |
| 52_ATCC14918.crt.3 | JHELIIPC_00594 hypothetical protein | 34 | 186 | 63.9 | 7.00E-11 | 34 | 100% | 0 | 0% | 1 | 34 | 100% | 150 | 183 | + | 150 |
| 52_ATCC14918.crt.3 | AJHAHHEL_01010 hypothetical protein | 34 | 186 | 63.9 | 7.00E-11 | 34 | 100% | 0 | 0% | 1 | 34 | 100% | 150 | 183 | + | 150 |
| 52_ATCC14918.crt.3 | DMLKMGFL_01488 hypothetical protein | 34 | 186 | 63.9 | 7.00E-11 | 34 | 100% | 0 | 0% | 1 | 34 | 100% | 150 | 183 | + | 150 |
| 52_ATCC14918.crt.3 | NELECJBE_01036 hypothetical protein | 34 | 186 | 63.9 | 7.00E-11 | 34 | 100% | 0 | 0% | 1 | 34 | 100% | 150 | 183 | + | 150 |
| 52_ATCC14918.crt.3 | HPLLMOOM_00592 hypothetical protein | 34 | 186 | 63.9 | 7.00E-11 | 34 | 100% | 0 | 0% | 1 | 34 | 100% | 150 | 183 | + | 150 |
| 52_ATCC14918.crt.3 | MCFNMICF_01409 hypothetical protein | 34 | 186 | 63.9 | 7.00E-11 | 34 | 100% | 0 | 0% | 1 | 34 | 100% | 150 | 183 | + | 150 |
| 52_ATCC14918.crt.4 | JEOJEKGO_01460 hypothetical protein | 34 | 306 | 63.9 | 7.00E-11 | 34 | 100% | 0 | 0% | 1 | 34 | 100% | 146 | 179 | + | 146 |
| 52_ATCC14918.crt.4 | DCKFNFNP_00568 hypothetical protein | 34 | 306 | 63.9 | 7.00E-11 | 34 | 100% | 0 | 0% | 1 | 34 | 100% | 146 | 179 | + | 146 |
| 52_ATCC14918.crt.4 | ECKACGMC_01462 hypothetical protein | 34 | 306 | 63.9 | 7.00E-11 | 34 | 100% | 0 | 0% | 1 | 34 | 100% | 146 | 179 | + | 146 |
| 52_ATCC14918.crt.4 | MDEAFGPO_00191 hypothetical protein | 34 | 306 | 63.9 | 7.00E-11 | 34 | 100% | 0 | 0% | 1 | 34 | 100% | 146 | 179 | + | 146 |
| 52_ATCC14918.crt.4 | HEGDFFNC_01370 hypothetical protein | 34 | 306 | 63.9 | 7.00E-11 | 34 | 100% | 0 | 0% | 1 | 34 | 100% | 146 | 179 | + | 146 |
| 52_ATCC14918.crt.4 | MONDJAPC_00627 hypothetical protein | 34 | 306 | 63.9 | 7.00E-11 | 34 | 100% | 0 | 0% | 1 | 34 | 100% | 146 | 179 | + | 146 |
| 52_ATCC14918.crt.5 | NOHIT | 34 | | | | | | | | | | | | | | |
| 52_ATCC14918.crt.6 | EJBPJIHF_01284 hypothetical protein | 35 | 381 | 65.8 | 2.00E-11 | 35 | 100% | 0 | 0% | 1 | 35 | 100% | 156 | 190 | - | 156 |
| 52_ATCC14918.crt.6 | MMBCLDPD_00678 hypothetical protein | 35 | 381 | 65.8 | 2.00E-11 | 35 | 100% | 0 | 0% | 1 | 35 | 100% | 156 | 190 | - | 156 |
| 52_ATCC14918.crt.6 | GGLKEJFF_01158 hypothetical protein | 35 | 381 | 65.8 | 2.00E-11 | 35 | 100% | 0 | 0% | 1 | 35 | 100% | 156 | 190 | - | 156 |
| 52_ATCC14918.crt.6 | GIKPJECN_01232 hypothetical protein | 35 | 381 | 65.8 | 2.00E-11 | 35 | 100% | 0 | 0% | 1 | 35 | 100% | 156 | 190 | - | 156 |
| 52_ATCC14918.crt.6 | GPBJDOFB_01192 hypothetical protein | 35 | 381 | 65.8 | 2.00E-11 | 35 | 100% | 0 | 0% | 1 | 35 | 100% | 156 | 190 | - | 156 |
| 52_ATCC14918.crt.6 | ACJECOPP_01208 hypothetical protein | 35 | 381 | 65.8 | 2.00E-11 | 35 | 100% | 0 | 0% | 1 | 35 | 100% | 156 | 190 | - | 156 |
| 52_ATCC14918.crt.6 | ICMALKPE_01219 hypothetical protein | 35 | 381 | 65.8 | 2.00E-11 | 35 | 100% | 0 | 0% | 1 | 35 | 100% | 156 | 190 | - | 156 |
| 52_ATCC14918.crt.6 | IIJJCGIL_01210 hypothetical protein | 35 | 381 | 65.8 | 2.00E-11 | 35 | 100% | 0 | 0% | 1 | 35 | 100% | 156 | 190 | - | 156 |
| 52_ATCC14918.crt.7 | FOAEMPCG_00787 eukaryotic hypersensitive-induced | 35 | 828 | 65.8 | 2.00E-11 | 35 | 100% | 0 | 0% | 1 | 35 | 100% | 511 | 545 | + | 511 |
| 52_ATCC14918.crt.7 | LLAAGEIN_00659 eukaryotic hypersensitive-induced | 35 | 828 | 65.8 | 2.00E-11 | 35 | 100% | 0 | 0% | 1 | 35 | 100% | 511 | 545 | + | 511 |
| 52_ATCC14918.crt.7 | EFGBCCNL_00468 eukaryotic hypersensitive-induced | 35 | 828 | 65.8 | 2.00E-11 | 35 | 100% | 0 | 0% | 1 | 35 | 100% | 511 | 545 | + | 511 |
| 52_ATCC14918.crt.7 | JHELIIPC_01229 eukaryotic hypersensitive-induced re | 35 | 828 | 65.8 | 2.00E-11 | 35 | 100% | 0 | 0% | 1 | 35 | 100% | 511 | 545 | + | 511 |
| 52_ATCC14918.crt.7 | HPLLMOOM_01228 eukaryotic hypersensitive-induce | 35 | 828 | 65.8 | 2.00E-11 | 35 | 100% | 0 | 0% | 1 | 35 | 100% | 511 | 545 | + | 511 |
| 52_ATCC14918.crt.7 | MCFNMICF_00775 eukaryotic hypersensitive-induced | 35 | 828 | 65.8 | 2.00E-11 | 35 | 100% | 0 | 0% | 1 | 35 | 100% | 511 | 545 | + | 511 |
| 52_ATCC14918.crt.7 | MONDJAPC_00342 eukaryotic hypersensitive-induced | 35 | 828 | 65.8 | 2.00E-11 | 35 | 100% | 0 | 0% | 1 | 35 | 100% | 511 | 545 | + | 511 |
| 52_ATCC14918.crt.8 | NOHIT | 33 | | | | | | | | | | | | | | |
| 53_ATCC14919.crt.1 | NOHIT | 38 | | | | | | | | | | | | | | |
| 53_ATCC14919.crt.2 | NOHIT | 37 | | | | | | | | | | | | | | |
| 53_ATCC14919.crt.3 | NOHIT | 0 | | | | | | | | | | | | | | |
| 53_ATCC14919.crt.4 | NOHIT | 32 | | | | | | | | | | | | | | |
| 53_ATCC14919.crt.5 | BNEKOFAL_00674 hypothetical protein | 34 | 285 | 63.9 | 7.00E-11 | 34 | 100% | 0 | 0% | 1 | 34 | 100% | 19 | 52 | + | 19 |
| 53_ATCC14919.crt.5 | FOAEMPCG_01040 hypothetical protein | 34 | 285 | 63.9 | 7.00E-11 | 34 | 100% | 0 | 0% | 1 | 34 | 100% | 19 | 52 | + | 19 |
| 53_ATCC14919.crt.5 | MCEIOEBL_00624 hypothetical protein | 34 | 285 | 63.9 | 7.00E-11 | 34 | 100% | 0 | 0% | 1 | 34 | 100% | 19 | 52 | + | 19 |
| 53_ATCC14919.crt.5 | PPKGMMPD_01012 hypothetical protein | 34 | 285 | 63.9 | 7.00E-11 | 34 | 100% | 0 | 0% | 1 | 34 | 100% | 19 | 52 | + | 19 |
| 53_ATCC14919.crt.5 | LLAAGEIN_01368 hypothetical protein | 34 | 285 | 63.9 | 7.00E-11 | 34 | 100% | 0 | 0% | 1 | 34 | 100% | 19 | 52 | + | 19 |
| 53_ATCC14919.crt.5 | AIKJPEBD_01573 hypothetical protein | 34 | 285 | 63.9 | 7.00E-11 | 34 | 100% | 0 | 0% | 1 | 34 | 100% | 19 | 52 | + | 19 |
| 53_ATCC14919.crt.5 | FPFNMMFM_00882 hypothetical protein | 34 | 285 | 63.9 | 7.00E-11 | 34 | 100% | 0 | 0% | 1 | 34 | 100% | 19 | 52 | + | 19 |
| 53_ATCC14919.crt.5 | IJFGMCCO_00995 hypothetical protein | 34 | 285 | 63.9 | 7.00E-11 | 34 | 100% | 0 | 0% | 1 | 34 | 100% | 19 | 52 | + | 19 |
| 53_ATCC14919.crt.5 | EJBPJIHF_01045 hypothetical protein | 34 | 285 | 63.9 | 7.00E-11 | 34 | 100% | 0 | 0% | 1 | 34 | 100% | 19 | 52 | + | 19 |
| 53_ATCC14919.crt.5 | LAKBNEPL_01011 hypothetical protein | 34 | 285 | 63.9 | 7.00E-11 | 34 | 100% | 0 | 0% | 1 | 34 | 100% | 19 | 52 | + | 19 |
| 53_ATCC14919.crt.5 | MMBCLDPD_00865 hypothetical protein | 34 | 285 | 63.9 | 7.00E-11 | 34 | 100% | 0 | 0% | 1 | 34 | 100% | 19 | 52 | + | 19 |
| 53_ATCC14919.crt.5 | PLPKHLCC_00893 hypothetical protein | 34 | 285 | 63.9 | 7.00E-11 | 34 | 100% | 0 | 0% | 1 | 34 | 100% | 19 | 52 | + | 19 |
| 53_ATCC14919.crt.5 | JNLHFPPN_01263 hypothetical protein | 34 | 285 | 63.9 | 7.00E-11 | 34 | 100% | 0 | 0% | 1 | 34 | 100% | 19 | 52 | + | 19 |
| 53_ATCC14919.crt.5 | ODIAAKBJ_01263 hypothetical protein | 34 | 285 | 63.9 | 7.00E-11 | 34 | 100% | 0 | 0% | 1 | 34 | 100% | 19 | 52 | + | 19 |
| 53_ATCC14919.crt.5 | ECKACGMC_00553 hypothetical protein | 34 | 285 | 63.9 | 7.00E-11 | 34 | 100% | 0 | 0% | 1 | 34 | 100% | 19 | 52 | + | 19 |
| 53_ATCC14919.crt.5 | DIPDIGDN_01262 hypothetical protein | 34 | 285 | 63.9 | 7.00E-11 | 34 | 100% | 0 | 0% | 1 | 34 | 100% | 19 | 52 | + | 19 |
| 53_ATCC14919.crt.5 | EFGBCCNL_00703 hypothetical protein | 34 | 285 | 63.9 | 7.00E-11 | 34 | 100% | 0 | 0% | 1 | 34 | 100% | 19 | 52 | + | 19 |
| 53_ATCC14919.crt.5 | JHELIIPC_00975 hypothetical protein | 34 | 285 | 63.9 | 7.00E-11 | 34 | 100% | 0 | 0% | 1 | 34 | 100% | 19 | 52 | + | 19 |
| 53_ATCC14919.crt.5 | GPBJDOFB_00997 hypothetical protein | 34 | 285 | 63.9 | 7.00E-11 | 34 | 100% | 0 | 0% | 1 | 34 | 100% | 19 | 52 | + | 19 |
| 53_ATCC14919.crt.5 | ACJECOPP_01021 hypothetical protein | 34 | 285 | 63.9 | 7.00E-11 | 34 | 100% | 0 | 0% | 1 | 34 | 100% | 19 | 52 | + | 19 |
| 53_ATCC14919.crt.5 | ICMALKPE_01033 hypothetical protein | 34 | 285 | 63.9 | 7.00E-11 | 34 | 100% | 0 | 0% | 1 | 34 | 100% | 19 | 52 | + | 19 |
| 53_ATCC14919.crt.5 | AJHAHHEL_01315 hypothetical protein | 34 | 285 | 63.9 | 7.00E-11 | 34 | 100% | 0 | 0% | 1 | 34 | 100% | 19 | 52 | + | 19 |
| 53_ATCC14919.crt.5 | DMLKMGFL_01490 hypothetical protein | 34 | 285 | 63.9 | 7.00E-11 | 34 | 100% | 0 | 0% | 1 | 34 | 100% | 19 | 52 | + | 19 |
| 53_ATCC14919.crt.5 | HEGDFFNC_00554 hypothetical protein | 34 | 285 | 63.9 | 7.00E-11 | 34 | 100% | 0 | 0% | 1 | 34 | 100% | 19 | 52 | + | 19 |
| 53_ATCC14919.crt.5 | IIJJCGIL_01024 hypothetical protein | 34 | 285 | 63.9 | 7.00E-11 | 34 | 100% | 0 | 0% | 1 | 34 | 100% | 19 | 52 | + | 19 |
| 53_ATCC14919.crt.5 | GLBJHDNN_01015 hypothetical protein | 34 | 285 | 63.9 | 7.00E-11 | 34 | 100% | 0 | 0% | 1 | 34 | 100% | 19 | 52 | + | 19 |
| 53_ATCC14919.crt.5 | HPLLMOOM_00975 hypothetical protein | 34 | 285 | 63.9 | 7.00E-11 | 34 | 100% | 0 | 0% | 1 | 34 | 100% | 19 | 52 | + | 19 |
| 53_ATCC14919.crt.5 | MCFNMICF_01027 hypothetical protein | 34 | 285 | 63.9 | 7.00E-11 | 34 | 100% | 0 | 0% | 1 | 34 | 100% | 19 | 52 | + | 19 |
| 53_ATCC14919.crt.5 | MONDJAPC_01581 hypothetical protein | 34 | 285 | 63.9 | 7.00E-11 | 34 | 100% | 0 | 0% | 1 | 34 | 100% | 19 | 52 | + | 19 |
| 53_ATCC14919.crt.6 | JEOJEKGO_01460 hypothetical protein | 34 | 306 | 63.9 | 7.00E-11 | 34 | 100% | 0 | 0% | 1 | 34 | 100% | 146 | 179 | + | 146 |
| 53_ATCC14919.crt.6 | DCKFNFNP_00568 hypothetical protein | 34 | 306 | 63.9 | 7.00E-11 | 34 | 100% | 0 | 0% | 1 | 34 | 100% | 146 | 179 | + | 146 |
| 53_ATCC14919.crt.6 | ECKACGMC_01462 hypothetical protein | 34 | 306 | 63.9 | 7.00E-11 | 34 | 100% | 0 | 0% | 1 | 34 | 100% | 146 | 179 | + | 146 |
| 53_ATCC14919.crt.6 | MDEAFGPO_00191 hypothetical protein | 34 | 306 | 63.9 | 7.00E-11 | 34 | 100% | 0 | 0% | 1 | 34 | 100% | 146 | 179 | + | 146 |
| 53_ATCC14919.crt.6 | HEGDFFNC_01370 hypothetical protein | 34 | 306 | 63.9 | 7.00E-11 | 34 | 100% | 0 | 0% | 1 | 34 | 100% | 146 | 179 | + | 146 |
| 53_ATCC14919.crt.6 | MONDJAPC_00627 hypothetical protein | 34 | 306 | 63.9 | 7.00E-11 | 34 | 100% | 0 | 0% | 1 | 34 | 100% | 146 | 179 | + | 146 |
| 53_ATCC14919.crt.7 | NOHIT | 34 | | | | | | | | | | | | | | |
| 53_ATCC14919.crt.8 | EJBPJIHF_01284 hypothetical protein | 35 | 381 | 65.8 | 2.00E-11 | 35 | 100% | 0 | 0% | 1 | 35 | 100% | 156 | 190 | - | 156 |
| 53_ATCC14919.crt.8 | MMBCLDPD_00678 hypothetical protein | 35 | 381 | 65.8 | 2.00E-11 | 35 | 100% | 0 | 0% | 1 | 35 | 100% | 156 | 190 | - | 156 |
| 53_ATCC14919.crt.8 | GGLKEJFF_01158 hypothetical protein | 35 | 381 | 65.8 | 2.00E-11 | 35 | 100% | 0 | 0% | 1 | 35 | 100% | 156 | 190 | - | 156 |
| 53_ATCC14919.crt.8 | GIKPJECN_01232 hypothetical protein | 35 | 381 | 65.8 | 2.00E-11 | 35 | 100% | 0 | 0% | 1 | 35 | 100% | 156 | 190 | - | 156 |
| 53_ATCC14919.crt.8 | GPBJDOFB_01192 hypothetical protein | 35 | 381 | 65.8 | 2.00E-11 | 35 | 100% | 0 | 0% | 1 | 35 | 100% | 156 | 190 | - | 156 |
| 53_ATCC14919.crt.8 | ACJECOPP_01208 hypothetical protein | 35 | 381 | 65.8 | 2.00E-11 | 35 | 100% | 0 | 0% | 1 | 35 | 100% | 156 | 190 | - | 156 |
| 53_ATCC14919.crt.8 | ICMALKPE_01219 hypothetical protein | 35 | 381 | 65.8 | 2.00E-11 | 35 | 100% | 0 | 0% | 1 | 35 | 100% | 156 | 190 | - | 156 |
| 53_ATCC14919.crt.8 | IIJJCGIL_01210 hypothetical protein | 35 | 381 | 65.8 | 2.00E-11 | 35 | 100% | 0 | 0% | 1 | 35 | 100% | 156 | 190 | - | 156 |
| 53_ATCC14919.crt.9 | FOAEMPCG_00787 eukaryotic hypersensitive-induced | 35 | 828 | 65.8 | 2.00E-11 | 35 | 100% | 0 | 0% | 1 | 35 | 100% | 511 | 545 | + | 511 |
| 53_ATCC14919.crt.9 | LLAAGEIN_00659 eukaryotic hypersensitive-induced | 35 | 828 | 65.8 | 2.00E-11 | 35 | 100% | 0 | 0% | 1 | 35 | 100% | 511 | 545 | + | 511 |
| 53_ATCC14919.crt.9 | EFGBCCNL_00468 eukaryotic hypersensitive-induced | 35 | 828 | 65.8 | 2.00E-11 | 35 | 100% | 0 | 0% | 1 | 35 | 100% | 511 | 545 | + | 511 |
| 53_ATCC14919.crt.9 | JHELIIPC_01229 eukaryotic hypersensitive-induced re | 35 | 828 | 65.8 | 2.00E-11 | 35 | 100% | 0 | 0% | 1 | 35 | 100% | 511 | 545 | + | 511 |
| 53_ATCC14919.crt.9 | HPLLMOOM_01228 eukaryotic hypersensitive-induce | 35 | 828 | 65.8 | 2.00E-11 | 35 | 100% | 0 | 0% | 1 | 35 | 100% | 511 | 545 | + | 511 |
| 53_ATCC14919.crt.9 | MCFNMICF_00775 eukaryotic hypersensitive-induced | 35 | 828 | 65.8 | 2.00E-11 | 35 | 100% | 0 | 0% | 1 | 35 | 100% | 511 | 545 | + | 511 |
| 53_ATCC14919.crt.9 | MONDJAPC_00342 eukaryotic hypersensitive-induced | 35 | 828 | 65.8 | 2.00E-11 | 35 | 100% | 0 | 0% | 1 | 35 | 100% | 511 | 545 | + | 511 |
| 53_ATCC14919.crt.10 | NOHIT | 33 | | | | | | | | | | | | | | |
| 54_MGAS9429_2.crt.1 | NOHIT | 32 | | | | | | | | | | | | | | |
| 54_MGAS9429_2.crt.2 | BNEKOFAL_00674 hypothetical protein | 34 | 285 | 63.9 | 7.00E-11 | 34 | 100% | 0 | 0% | 1 | 34 | 100% | 19 | 52 | + | 19 |
| 54_MGAS9429_2.crt.2 | FOAEMPCG_01040 hypothetical protein | 34 | 285 | 63.9 | 7.00E-11 | 34 | 100% | 0 | 0% | 1 | 34 | 100% | 19 | 52 | + | 19 |
| 54_MGAS9429_2.crt.2 | MCEIOEBL_00624 hypothetical protein | 34 | 285 | 63.9 | 7.00E-11 | 34 | 100% | 0 | 0% | 1 | 34 | 100% | 19 | 52 | + | 19 |
| 54_MGAS9429_2.crt.2 | PPKGMMPD_01012 hypothetical protein | 34 | 285 | 63.9 | 7.00E-11 | 34 | 100% | 0 | 0% | 1 | 34 | 100% | 19 | 52 | + | 19 |
| 54_MGAS9429_2.crt.2 | LLAAGEIN_01368 hypothetical protein | 34 | 285 | 63.9 | 7.00E-11 | 34 | 100% | 0 | 0% | 1 | 34 | 100% | 19 | 52 | + | 19 |
| 54_MGAS9429_2.crt.2 | AIKJPEBD_01573 hypothetical protein | 34 | 285 | 63.9 | 7.00E-11 | 34 | 100% | 0 | 0% | 1 | 34 | 100% | 19 | 52 | + | 19 |
| 54_MGAS9429_2.crt.2 | FPFNMMFM_00882 hypothetical protein | 34 | 285 | 63.9 | 7.00E-11 | 34 | 100% | 0 | 0% | 1 | 34 | 100% | 19 | 52 | + | 19 |
| 54_MGAS9429_2.crt.2 | IJFGMCCO_00995 hypothetical protein | 34 | 285 | 63.9 | 7.00E-11 | 34 | 100% | 0 | 0% | 1 | 34 | 100% | 19 | 52 | + | 19 |
| 54_MGAS9429_2.crt.2 | EJBPJIHF_01045 hypothetical protein | 34 | 285 | 63.9 | 7.00E-11 | 34 | 100% | 0 | 0% | 1 | 34 | 100% | 19 | 52 | + | 19 |
| 54_MGAS9429_2.crt.2 | LAKBNEPL_01011 hypothetical protein | 34 | 285 | 63.9 | 7.00E-11 | 34 | 100% | 0 | 0% | 1 | 34 | 100% | 19 | 52 | + | 19 |
| 54_MGAS9429_2.crt.2 | MMBCLDPD_00865 hypothetical protein | 34 | 285 | 63.9 | 7.00E-11 | 34 | 100% | 0 | 0% | 1 | 34 | 100% | 19 | 52 | + | 19 |
| 54_MGAS9429_2.crt.2 | PLPKHLCC_00893 hypothetical protein | 34 | 285 | 63.9 | 7.00E-11 | 34 | 100% | 0 | 0% | 1 | 34 | 100% | 19 | 52 | + | 19 |
| 54_MGAS9429_2.crt.2 | JNLHFPPN_01263 hypothetical protein | 34 | 285 | 63.9 | 7.00E-11 | 34 | 100% | 0 | 0% | 1 | 34 | 100% | 19 | 52 | + | 19 |
| 54_MGAS9429_2.crt.2 | ODIAAKBJ_01263 hypothetical protein | 34 | 285 | 63.9 | 7.00E-11 | 34 | 100% | 0 | 0% | 1 | 34 | 100% | 19 | 52 | + | 19 |
| 54_MGAS9429_2.crt.2 | ECKACGMC_00553 hypothetical protein | 34 | 285 | 63.9 | 7.00E-11 | 34 | 100% | 0 | 0% | 1 | 34 | 100% | 19 | 52 | + | 19 |
| 54_MGAS9429_2.crt.2 | DIPDIGDN_01262 hypothetical protein | 34 | 285 | 63.9 | 7.00E-11 | 34 | 100% | 0 | 0% | 1 | 34 | 100% | 19 | 52 | + | 19 |
| 54_MGAS9429_2.crt.2 | EFGBCCNL_00703 hypothetical protein | 34 | 285 | 63.9 | 7.00E-11 | 34 | 100% | 0 | 0% | 1 | 34 | 100% | 19 | 52 | + | 19 |
| 54_MGAS9429_2.crt.2 | JHELIIPC_00975 hypothetical protein | 34 | 285 | 63.9 | 7.00E-11 | 34 | 100% | 0 | 0% | 1 | 34 | 100% | 19 | 52 | + | 19 |
| 54_MGAS9429_2.crt.2 | GPBJDOFB_00997 hypothetical protein | 34 | 285 | 63.9 | 7.00E-11 | 34 | 100% | 0 | 0% | 1 | 34 | 100% | 19 | 52 | + | 19 |
| 54_MGAS9429_2.crt.2 | ACJECOPP_01021 hypothetical protein | 34 | 285 | 63.9 | 7.00E-11 | 34 | 100% | 0 | 0% | 1 | 34 | 100% | 19 | 52 | + | 19 |
| 54_MGAS9429_2.crt.2 | ICMALKPE_01033 hypothetical protein | 34 | 285 | 63.9 | 7.00E-11 | 34 | 100% | 0 | 0% | 1 | 34 | 100% | 19 | 52 | + | 19 |
| 54_MGAS9429_2.crt.2 | AJHAHHEL_01315 hypothetical protein | 34 | 285 | 63.9 | 7.00E-11 | 34 | 100% | 0 | 0% | 1 | 34 | 100% | 19 | 52 | + | 19 |
| 54_MGAS9429_2.crt.2 | DMLKMGFL_01490 hypothetical protein | 34 | 285 | 63.9 | 7.00E-11 | 34 | 100% | 0 | 0% | 1 | 34 | 100% | 19 | 52 | + | 19 |
| 54_MGAS9429_2.crt.2 | HEGDFFNC_00554 hypothetical protein | 34 | 285 | 63.9 | 7.00E-11 | 34 | 100% | 0 | 0% | 1 | 34 | 100% | 19 | 52 | + | 19 |
| 54_MGAS9429_2.crt.2 | IIJJCGIL_01024 hypothetical protein | 34 | 285 | 63.9 | 7.00E-11 | 34 | 100% | 0 | 0% | 1 | 34 | 100% | 19 | 52 | + | 19 |
| 54_MGAS9429_2.crt.2 | GLBJHDNN_01015 hypothetical protein | 34 | 285 | 63.9 | 7.00E-11 | 34 | 100% | 0 | 0% | 1 | 34 | 100% | 19 | 52 | + | 19 |
| 54_MGAS9429_2.crt.2 | HPLLMOOM_00975 hypothetical protein | 34 | 285 | 63.9 | 7.00E-11 | 34 | 100% | 0 | 0% | 1 | 34 | 100% | 19 | 52 | + | 19 |
| 54_MGAS9429_2.crt.2 | MCFNMICF_01027 hypothetical protein | 34 | 285 | 63.9 | 7.00E-11 | 34 | 100% | 0 | 0% | 1 | 34 | 100% | 19 | 52 | + | 19 |
| 54_MGAS9429_2.crt.2 | MONDJAPC_01581 hypothetical protein | 34 | 285 | 63.9 | 7.00E-11 | 34 | 100% | 0 | 0% | 1 | 34 | 100% | 19 | 52 | + | 19 |
| 54_MGAS9429_2.crt.3 | BNEKOFAL_01130 hypothetical protein | 34 | 186 | 63.9 | 7.00E-11 | 34 | 100% | 0 | 0% | 1 | 34 | 100% | 150 | 183 | + | 150 |
| 54_MGAS9429_2.crt.3 | FOAEMPCG_01422 hypothetical protein | 34 | 186 | 63.9 | 7.00E-11 | 34 | 100% | 0 | 0% | 1 | 34 | 100% | 150 | 183 | + | 150 |
| 54_MGAS9429_2.crt.3 | MCEIOEBL_01079 hypothetical protein | 34 | 186 | 63.9 | 7.00E-11 | 34 | 100% | 0 | 0% | 1 | 34 | 100% | 150 | 183 | + | 150 |
| 54_MGAS9429_2.crt.3 | LLAAGEIN_01062 hypothetical protein | 34 | 186 | 63.9 | 7.00E-11 | 34 | 100% | 0 | 0% | 1 | 34 | 100% | 150 | 183 | + | 150 |
| 54_MGAS9429_2.crt.3 | IJFGMCCO_00990 hypothetical protein | 34 | 186 | 63.9 | 7.00E-11 | 34 | 100% | 0 | 0% | 1 | 34 | 100% | 150 | 183 | + | 150 |
| 54_MGAS9429_2.crt.3 | MDEAFGPO_00646 hypothetical protein | 34 | 186 | 63.9 | 7.00E-11 | 34 | 100% | 0 | 0% | 1 | 34 | 100% | 150 | 183 | + | 150 |
| 54_MGAS9429_2.crt.3 | MDEAFGPO_00220 hypothetical protein | 34 | 186 | 63.9 | 7.00E-11 | 34 | 100% | 0 | 0% | 1 | 34 | 100% | 150 | 183 | + | 150 |
| 54_MGAS9429_2.crt.3 | EFGBCCNL_00701 hypothetical protein | 34 | 186 | 63.9 | 7.00E-11 | 34 | 100% | 0 | 0% | 1 | 34 | 100% | 150 | 183 | + | 150 |

| | | | | | | | | | | | | | | | |
|---|---|---|---|---|---|---|---|---|---|---|---|---|---|---|---|
| 54_MGAS9429_2.crt.3 | JHELIIPC_00594 hypothetical protein | 34 | 186 | 63.9 | 7.00E-11 | 34 | 100% | 0 | 0% | 1 | 34 | 100% | 150 | 183 + | 150 |
| 54_MGAS9429_2.crt.3 | AJHAHHEL_01010 hypothetical protein | 34 | 186 | 63.9 | 7.00E-11 | 34 | 100% | 0 | 0% | 1 | 34 | 100% | 150 | 183 + | 150 |
| 54_MGAS9429_2.crt.3 | DMLKMGFL_01488 hypothetical protein | 34 | 186 | 63.9 | 7.00E-11 | 34 | 100% | 0 | 0% | 1 | 34 | 100% | 150 | 183 + | 150 |
| 54_MGAS9429_2.crt.3 | NELECJBE_01036 hypothetical protein | 34 | 186 | 63.9 | 7.00E-11 | 34 | 100% | 0 | 0% | 1 | 34 | 100% | 150 | 183 + | 150 |
| 54_MGAS9429_2.crt.3 | HPLLMOOM_00592 hypothetical protein | 34 | 186 | 63.9 | 7.00E-11 | 34 | 100% | 0 | 0% | 1 | 34 | 100% | 150 | 183 + | 150 |
| 54_MGAS9429_2.crt.3 | MCFNMICF_01409 hypothetical protein | 34 | 186 | 63.9 | 7.00E-11 | 34 | 100% | 0 | 0% | 1 | 34 | 100% | 150 | 183 + | 150 |
| 54_MGAS9429_2.crt.4 | JEOJEKGO_01460 hypothetical protein | 34 | 306 | 63.9 | 7.00E-11 | 34 | 100% | 0 | 0% | 1 | 34 | 100% | 146 | 179 + | 146 |
| 54_MGAS9429_2.crt.4 | DCKFNFNP_00568 hypothetical protein | 34 | 306 | 63.9 | 7.00E-11 | 34 | 100% | 0 | 0% | 1 | 34 | 100% | 146 | 179 + | 146 |
| 54_MGAS9429_2.crt.4 | ECKACGMC_01462 hypothetical protein | 34 | 306 | 63.9 | 7.00E-11 | 34 | 100% | 0 | 0% | 1 | 34 | 100% | 146 | 179 + | 146 |
| 54_MGAS9429_2.crt.4 | MDEAFGPO_00191 hypothetical protein | 34 | 306 | 63.9 | 7.00E-11 | 34 | 100% | 0 | 0% | 1 | 34 | 100% | 146 | 179 + | 146 |
| 54_MGAS9429_2.crt.4 | HEGDFFNC_01370 hypothetical protein | 34 | 306 | 63.9 | 7.00E-11 | 34 | 100% | 0 | 0% | 1 | 34 | 100% | 146 | 179 + | 146 |
| 54_MGAS9429_2.crt.4 | MONDJAPC_00627 hypothetical protein | 34 | 306 | 63.9 | 7.00E-11 | 34 | 100% | 0 | 0% | 1 | 34 | 100% | 146 | 179 + | 146 |
| 54_MGAS9429_2.crt.5 | NOHIT | 34 | | | | | | | | | | | | | |
| 54_MGAS9429_2.crt.6 | EJBPJIHF_01284 hypothetical protein | 35 | 381 | 65.8 | 2.00E-11 | 35 | 100% | 0 | 0% | 1 | 35 | 100% | 156 | 190 - | 156 |
| 54_MGAS9429_2.crt.6 | MMBCLDPD_00678 hypothetical protein | 35 | 381 | 65.8 | 2.00E-11 | 35 | 100% | 0 | 0% | 1 | 35 | 100% | 156 | 190 - | 156 |
| 54_MGAS9429_2.crt.6 | GGLKEJFF_01158 hypothetical protein | 35 | 381 | 65.8 | 2.00E-11 | 35 | 100% | 0 | 0% | 1 | 35 | 100% | 156 | 190 - | 156 |
| 54_MGAS9429_2.crt.6 | GIKPJECN_01232 hypothetical protein | 35 | 381 | 65.8 | 2.00E-11 | 35 | 100% | 0 | 0% | 1 | 35 | 100% | 156 | 190 - | 156 |
| 54_MGAS9429_2.crt.6 | GPBJDOFB_01192 hypothetical protein | 35 | 381 | 65.8 | 2.00E-11 | 35 | 100% | 0 | 0% | 1 | 35 | 100% | 156 | 190 - | 156 |
| 54_MGAS9429_2.crt.6 | ACJECOPP_01208 hypothetical protein | 35 | 381 | 65.8 | 2.00E-11 | 35 | 100% | 0 | 0% | 1 | 35 | 100% | 156 | 190 - | 156 |
| 54_MGAS9429_2.crt.6 | ICMALKPE_01219 hypothetical protein | 35 | 381 | 65.8 | 2.00E-11 | 35 | 100% | 0 | 0% | 1 | 35 | 100% | 156 | 190 - | 156 |
| 54_MGAS9429_2.crt.6 | IIJJCGIL_01210 hypothetical protein | 35 | 381 | 65.8 | 2.00E-11 | 35 | 100% | 0 | 0% | 1 | 35 | 100% | 156 | 190 - | 156 |
| 54_MGAS9429_2.crt.7 | FOAEMPCG_00787 eukaryotic hypersensitive-induced | 35 | 828 | 65.8 | 2.00E-11 | 35 | 100% | 0 | 0% | 1 | 35 | 100% | 511 | 545 + | 511 |
| 54_MGAS9429_2.crt.7 | LLAAGEIN_00659 eukaryotic hypersensitive-induced | 35 | 828 | 65.8 | 2.00E-11 | 35 | 100% | 0 | 0% | 1 | 35 | 100% | 511 | 545 + | 511 |
| 54_MGAS9429_2.crt.7 | EFGBCCNL_00468 eukaryotic hypersensitive-induced | 35 | 828 | 65.8 | 2.00E-11 | 35 | 100% | 0 | 0% | 1 | 35 | 100% | 511 | 545 + | 511 |
| 54_MGAS9429_2.crt.7 | JHELIIPC_01229 eukaryotic hypersensitive-induced re | 35 | 828 | 65.8 | 2.00E-11 | 35 | 100% | 0 | 0% | 1 | 35 | 100% | 511 | 545 + | 511 |
| 54_MGAS9429_2.crt.7 | HPLLMOOM_01228 eukaryotic hypersensitive-induce | 35 | 828 | 65.8 | 2.00E-11 | 35 | 100% | 0 | 0% | 1 | 35 | 100% | 511 | 545 + | 511 |
| 54_MGAS9429_2.crt.7 | MCFNMICF_00775 eukaryotic hypersensitive-induced | 35 | 828 | 65.8 | 2.00E-11 | 35 | 100% | 0 | 0% | 1 | 35 | 100% | 511 | 545 + | 511 |
| 54_MGAS9429_2.crt.7 | MONDJAPC_00342 eukaryotic hypersensitive-induced | 35 | 828 | 65.8 | 2.00E-11 | 35 | 100% | 0 | 0% | 1 | 35 | 100% | 511 | 545 + | 511 |
| 54_MGAS9429_2.crt.8 | NOHIT | 33 | | | | | | | | | | | | | |
| 55_MGAS2096_2.crt.1 | FOAEMPCG_00795 hypothetical protein | 34 | 1731 | 63.9 | 7.00E-11 | 34 | 100% | 0 | 0% | 1 | 34 | 100% | 744 | 777 - | 744 |
| 55_MGAS2096_2.crt.1 | LLAAGEIN_00651 hypothetical protein | 34 | 1731 | 63.9 | 7.00E-11 | 34 | 100% | 0 | 0% | 1 | 34 | 100% | 744 | 777 - | 744 |
| 55_MGAS2096_2.crt.1 | EFGBCCNL_00460 hypothetical protein | 34 | 1731 | 63.9 | 7.00E-11 | 34 | 100% | 0 | 0% | 1 | 34 | 100% | 744 | 777 - | 744 |
| 55_MGAS2096_2.crt.1 | JHELIIPC_01221 hypothetical protein | 34 | 1731 | 63.9 | 7.00E-11 | 34 | 100% | 0 | 0% | 1 | 34 | 100% | 744 | 777 - | 744 |
| 55_MGAS2096_2.crt.1 | HPLLMOOM_01220 hypothetical protein | 34 | 1731 | 63.9 | 7.00E-11 | 34 | 100% | 0 | 0% | 1 | 34 | 100% | 744 | 777 - | 744 |
| 55_MGAS2096_2.crt.1 | MCFNMICF_00783 hypothetical protein | 34 | 1731 | 63.9 | 7.00E-11 | 34 | 100% | 0 | 0% | 1 | 34 | 100% | 744 | 777 - | 744 |
| 55_MGAS2096_2.crt.1 | MONDJAPC_00350 hypothetical protein | 34 | 1731 | 63.9 | 7.00E-11 | 34 | 100% | 0 | 0% | 1 | 34 | 100% | 744 | 777 - | 744 |
| 55_MGAS2096_2.crt.2 | FOAEMPCG_00787 eukaryotic hypersensitive-induced | 36 | 828 | 67.6 | 6.00E-12 | 36 | 100% | 0 | 0% | 1 | 36 | 100% | 511 | 546 - | 511 |
| 55_MGAS2096_2.crt.2 | LLAAGEIN_00659 eukaryotic hypersensitive-induced | 36 | 828 | 67.6 | 6.00E-12 | 36 | 100% | 0 | 0% | 1 | 36 | 100% | 511 | 546 - | 511 |
| 55_MGAS2096_2.crt.2 | EFGBCCNL_00468 eukaryotic hypersensitive-induced | 36 | 828 | 67.6 | 6.00E-12 | 36 | 100% | 0 | 0% | 1 | 36 | 100% | 511 | 546 - | 511 |
| 55_MGAS2096_2.crt.2 | JHELIIPC_01229 eukaryotic hypersensitive-induced re | 36 | 828 | 67.6 | 6.00E-12 | 36 | 100% | 0 | 0% | 1 | 36 | 100% | 511 | 546 - | 511 |
| 55_MGAS2096_2.crt.2 | HPLLMOOM_01228 eukaryotic hypersensitive-induce | 36 | 828 | 67.6 | 6.00E-12 | 36 | 100% | 0 | 0% | 1 | 36 | 100% | 511 | 546 - | 511 |
| 55_MGAS2096_2.crt.2 | MCFNMICF_00775 eukaryotic hypersensitive-induced | 36 | 828 | 67.6 | 6.00E-12 | 36 | 100% | 0 | 0% | 1 | 36 | 100% | 511 | 546 - | 511 |
| 55_MGAS2096_2.crt.2 | MONDJAPC_00342 eukaryotic hypersensitive-induced | 36 | 828 | 67.6 | 6.00E-12 | 36 | 100% | 0 | 0% | 1 | 36 | 100% | 511 | 546 - | 511 |
| 55_MGAS2096_2.crt.3 | NOHIT | 35 | | | | | | | | | | | | | |
| 55_MGAS2096_2.crt.4 | JEOJEKGO_01460 hypothetical protein | 35 | 306 | 65.8 | 2.00E-11 | 35 | 100% | 0 | 0% | 1 | 35 | 100% | 146 | 180 - | 146 |
| 55_MGAS2096_2.crt.4 | DCKFNFNP_00568 hypothetical protein | 35 | 306 | 65.8 | 2.00E-11 | 35 | 100% | 0 | 0% | 1 | 35 | 100% | 146 | 180 - | 146 |
| 55_MGAS2096_2.crt.4 | ECKACGMC_01462 hypothetical protein | 35 | 306 | 65.8 | 2.00E-11 | 35 | 100% | 0 | 0% | 1 | 35 | 100% | 146 | 180 - | 146 |
| 55_MGAS2096_2.crt.4 | MDEAFGPO_00191 hypothetical protein | 35 | 306 | 65.8 | 2.00E-11 | 35 | 100% | 0 | 0% | 1 | 35 | 100% | 146 | 180 - | 146 |
| 55_MGAS2096_2.crt.4 | HEGDFFNC_01370 hypothetical protein | 35 | 306 | 65.8 | 2.00E-11 | 35 | 100% | 0 | 0% | 1 | 35 | 100% | 146 | 180 - | 146 |
| 55_MGAS2096_2.crt.4 | MONDJAPC_00627 hypothetical protein | 35 | 306 | 65.8 | 2.00E-11 | 35 | 100% | 0 | 0% | 1 | 35 | 100% | 146 | 180 - | 146 |
| 55_MGAS2096_2.crt.5 | BNEKOFAL_00674 hypothetical protein | 35 | 285 | 65.8 | 2.00E-11 | 35 | 100% | 0 | 0% | 1 | 35 | 100% | 19 | 53 - | 19 |
| 55_MGAS2096_2.crt.5 | FOAEMPCG_01040 hypothetical protein | 35 | 285 | 65.8 | 2.00E-11 | 35 | 100% | 0 | 0% | 1 | 35 | 100% | 19 | 53 - | 19 |
| 55_MGAS2096_2.crt.5 | MCEIOEBL_00624 hypothetical protein | 35 | 285 | 65.8 | 2.00E-11 | 35 | 100% | 0 | 0% | 1 | 35 | 100% | 19 | 53 - | 19 |
| 55_MGAS2096_2.crt.5 | PPKGMMPD_01012 hypothetical protein | 35 | 285 | 65.8 | 2.00E-11 | 35 | 100% | 0 | 0% | 1 | 35 | 100% | 19 | 53 - | 19 |
| 55_MGAS2096_2.crt.5 | LLAAGEIN_01368 hypothetical protein | 35 | 285 | 65.8 | 2.00E-11 | 35 | 100% | 0 | 0% | 1 | 35 | 100% | 19 | 53 - | 19 |
| 55_MGAS2096_2.crt.5 | AIKJPEBD_01573 hypothetical protein | 35 | 285 | 65.8 | 2.00E-11 | 35 | 100% | 0 | 0% | 1 | 35 | 100% | 19 | 53 - | 19 |
| 55_MGAS2096_2.crt.5 | FPFNMMFM_00882 hypothetical protein | 35 | 285 | 65.8 | 2.00E-11 | 35 | 100% | 0 | 0% | 1 | 35 | 100% | 19 | 53 - | 19 |
| 55_MGAS2096_2.crt.5 | IJFGMCCO_00995 hypothetical protein | 35 | 285 | 65.8 | 2.00E-11 | 35 | 100% | 0 | 0% | 1 | 35 | 100% | 19 | 53 - | 19 |
| 55_MGAS2096_2.crt.5 | EJBPJIHF_01045 hypothetical protein | 35 | 285 | 65.8 | 2.00E-11 | 35 | 100% | 0 | 0% | 1 | 35 | 100% | 19 | 53 - | 19 |
| 55_MGAS2096_2.crt.5 | LAKBNEPL_01011 hypothetical protein | 35 | 285 | 65.8 | 2.00E-11 | 35 | 100% | 0 | 0% | 1 | 35 | 100% | 19 | 53 - | 19 |
| 55_MGAS2096_2.crt.5 | MMBCLDPD_00865 hypothetical protein | 35 | 285 | 65.8 | 2.00E-11 | 35 | 100% | 0 | 0% | 1 | 35 | 100% | 19 | 53 - | 19 |
| 55_MGAS2096_2.crt.5 | PLPKHLCC_00893 hypothetical protein | 35 | 285 | 65.8 | 2.00E-11 | 35 | 100% | 0 | 0% | 1 | 35 | 100% | 19 | 53 - | 19 |
| 55_MGAS2096_2.crt.5 | JNLHFPPN_01263 hypothetical protein | 35 | 285 | 65.8 | 2.00E-11 | 35 | 100% | 0 | 0% | 1 | 35 | 100% | 19 | 53 - | 19 |
| 55_MGAS2096_2.crt.5 | ODIAAKBJ_01263 hypothetical protein | 35 | 285 | 65.8 | 2.00E-11 | 35 | 100% | 0 | 0% | 1 | 35 | 100% | 19 | 53 - | 19 |
| 55_MGAS2096_2.crt.5 | ECKACGMC_00553 hypothetical protein | 35 | 285 | 65.8 | 2.00E-11 | 35 | 100% | 0 | 0% | 1 | 35 | 100% | 19 | 53 - | 19 |
| 55_MGAS2096_2.crt.5 | DIPDIGDN_01262 hypothetical protein | 35 | 285 | 65.8 | 2.00E-11 | 35 | 100% | 0 | 0% | 1 | 35 | 100% | 19 | 53 - | 19 |
| 55_MGAS2096_2.crt.5 | EFGBCCNL_00703 hypothetical protein | 35 | 285 | 65.8 | 2.00E-11 | 35 | 100% | 0 | 0% | 1 | 35 | 100% | 19 | 53 - | 19 |
| 55_MGAS2096_2.crt.5 | JHELIIPC_00975 hypothetical protein | 35 | 285 | 65.8 | 2.00E-11 | 35 | 100% | 0 | 0% | 1 | 35 | 100% | 19 | 53 - | 19 |
| 55_MGAS2096_2.crt.5 | GPBJDOFB_00997 hypothetical protein | 35 | 285 | 65.8 | 2.00E-11 | 35 | 100% | 0 | 0% | 1 | 35 | 100% | 19 | 53 - | 19 |
| 55_MGAS2096_2.crt.5 | ACJECOPP_01021 hypothetical protein | 35 | 285 | 65.8 | 2.00E-11 | 35 | 100% | 0 | 0% | 1 | 35 | 100% | 19 | 53 - | 19 |
| 55_MGAS2096_2.crt.5 | ICMALKPE_01033 hypothetical protein | 35 | 285 | 65.8 | 2.00E-11 | 35 | 100% | 0 | 0% | 1 | 35 | 100% | 19 | 53 - | 19 |
| 55_MGAS2096_2.crt.5 | AJHAHHEL_01315 hypothetical protein | 35 | 285 | 65.8 | 2.00E-11 | 35 | 100% | 0 | 0% | 1 | 35 | 100% | 19 | 53 - | 19 |
| 55_MGAS2096_2.crt.5 | DMLKMGFL_01490 hypothetical protein | 35 | 285 | 65.8 | 2.00E-11 | 35 | 100% | 0 | 0% | 1 | 35 | 100% | 19 | 53 - | 19 |
| 55_MGAS2096_2.crt.5 | HEGDFFNC_00554 hypothetical protein | 35 | 285 | 65.8 | 2.00E-11 | 35 | 100% | 0 | 0% | 1 | 35 | 100% | 19 | 53 - | 19 |
| 55_MGAS2096_2.crt.5 | IIJJCGIL_01024 hypothetical protein | 35 | 285 | 65.8 | 2.00E-11 | 35 | 100% | 0 | 0% | 1 | 35 | 100% | 19 | 53 - | 19 |
| 55_MGAS2096_2.crt.5 | GLBJHDNN_01015 hypothetical protein | 35 | 285 | 65.8 | 2.00E-11 | 35 | 100% | 0 | 0% | 1 | 35 | 100% | 19 | 53 - | 19 |
| 55_MGAS2096_2.crt.5 | HPLLMOOM_00975 hypothetical protein | 35 | 285 | 65.8 | 2.00E-11 | 35 | 100% | 0 | 0% | 1 | 35 | 100% | 19 | 53 - | 19 |
| 55_MGAS2096_2.crt.5 | MCFNMICF_01027 hypothetical protein | 35 | 285 | 65.8 | 2.00E-11 | 35 | 100% | 0 | 0% | 1 | 35 | 100% | 19 | 53 - | 19 |
| 55_MGAS2096_2.crt.5 | MONDJAPC_01581 hypothetical protein | 35 | 285 | 65.8 | 2.00E-11 | 35 | 100% | 0 | 0% | 1 | 35 | 100% | 19 | 53 - | 19 |
| 55_MGAS2096_2.crt.6 | NOHIT | 33 | | | | | | | | | | | | | |
| 55_MGAS2096_2.crt.7 | NOHIT | 0 | | | | | | | | | | | | | |
| 55_MGAS2096_2.crt.8 | NOHIT | 37 | | | | | | | | | | | | | |
| 55_MGAS2096_2.crt.9 | NOHIT | 38 | | | | | | | | | | | | | |
| 58_KIH06_201.crt.1 | NOHIT | 34 | | | | | | | | | | | | | |
| 58_KIH06_201.crt.2 | BNEKOFAL_00650 Transposase from transposon Tn9 | 34 | 1167 | 63.9 | 7.00E-11 | 34 | 100% | 0 | 0% | 1 | 34 | 100% | 1080 | 1113 + | 1080 |
| 58_KIH06_201.crt.2 | FOAEMPCG_01447 Transposase from transposon Tn9 | 34 | 1167 | 63.9 | 7.00E-11 | 34 | 100% | 0 | 0% | 1 | 34 | 100% | 1080 | 1113 + | 1080 |
| 58_KIH06_201.crt.2 | MCEIOEBL_00600 Transposase from transposon Tn91 | 34 | 1167 | 63.9 | 7.00E-11 | 34 | 100% | 0 | 0% | 1 | 34 | 100% | 1080 | 1113 + | 1080 |
| 58_KIH06_201.crt.2 | LLAAGEIN_01392 Transposase from transposon Tn91 | 34 | 1167 | 63.9 | 7.00E-11 | 34 | 100% | 0 | 0% | 1 | 34 | 100% | 1080 | 1113 + | 1080 |
| 58_KIH06_201.crt.2 | JNLHFPPN_01288 Transposase from transposon Tn91 | 34 | 1167 | 63.9 | 7.00E-11 | 34 | 100% | 0 | 0% | 1 | 34 | 100% | 1080 | 1113 + | 1080 |
| 58_KIH06_201.crt.2 | ODIAAKBJ_01288 Transposase from transposon Tn91 | 34 | 1167 | 63.9 | 7.00E-11 | 34 | 100% | 0 | 0% | 1 | 34 | 100% | 1080 | 1113 + | 1080 |
| 58_KIH06_201.crt.2 | ECKACGMC_00529 Transposase from transposon Tn9 | 34 | 1167 | 63.9 | 7.00E-11 | 34 | 100% | 0 | 0% | 1 | 34 | 100% | 1080 | 1113 + | 1080 |
| 58_KIH06_201.crt.2 | DIPDIGDN_01287 Transposase from transposon Tn91 | 34 | 1167 | 63.9 | 7.00E-11 | 34 | 100% | 0 | 0% | 1 | 34 | 100% | 1080 | 1113 + | 1080 |
| 58_KIH06_201.crt.2 | JHELIIPC_00570 Transposase from transposon Tn916 | 34 | 1167 | 63.9 | 7.00E-11 | 34 | 100% | 0 | 0% | 1 | 34 | 100% | 1080 | 1113 + | 1080 |
| 58_KIH06_201.crt.2 | AJHAHHEL_01339 Transposase from transposon Tn9 | 34 | 1167 | 63.9 | 7.00E-11 | 34 | 100% | 0 | 0% | 1 | 34 | 100% | 1080 | 1113 + | 1080 |
| 58_KIH06_201.crt.2 | HPLLMOOM_00568 Transposase from transposon Tn | 34 | 1167 | 63.9 | 7.00E-11 | 34 | 100% | 0 | 0% | 1 | 34 | 100% | 1080 | 1113 + | 1080 |
| 58_KIH06_201.crt.2 | MCFNMICF_01433 Transposase from transposon Tn9 | 34 | 1167 | 63.9 | 7.00E-11 | 34 | 100% | 0 | 0% | 1 | 34 | 100% | 1080 | 1113 + | 1080 |
| 58_KIH06_201.crt.2 | MONDJAPC_01605 Transposase from transposon Tn9 | 34 | 1167 | 63.9 | 7.00E-11 | 34 | 100% | 0 | 0% | 1 | 34 | 100% | 1080 | 1113 + | 1080 |
| 58_KIH06_201.crt.3 | NOHIT | 33 | | | | | | | | | | | | | |
| 6_MGAS6180.crt.1 | NOHIT | 30 | | | | | | | | | | | | | |
| 6_MGAS6180.crt.2 | NOHIT | 30 | | | | | | | | | | | | | |
| 6_MGAS6180.crt.3 | NOHIT | 30 | | | | | | | | | | | | | |
| 6_MGAS6180.crt.4 | NOHIT | 30 | | | | | | | | | | | | | |
| 7_MGAS5005.crt.1 | NOHIT | 30 | | | | | | | | | | | | | |
| 7_MGAS5005.crt.2 | NOHIT | 30 | | | | | | | | | | | | | |
| 7_MGAS5005.crt.3 | NOHIT | 30 | | | | | | | | | | | | | |
| 7_MGAS5005.crt.4 | NOHIT | 0 | | | | | | | | | | | | | |
| 7_MGAS5005.crt.5 | NOHIT | 33 | | | | | | | | | | | | | |
| 7_MGAS5005.crt.6 | NOHIT | 34 | | | | | | | | | | | | | |
| 7_MGAS5005.crt.7 | NOHIT | 33 | | | | | | | | | | | | | |
| 7_MGAS5005.crt.8 | NOHIT | 35 | | | | | | | | | | | | | |
| 8_MGAS9429.crt.1 | NOHIT | 38 | | | | | | | | | | | | | |
| 8_MGAS9429.crt.2 | NOHIT | 37 | | | | | | | | | | | | | |
| 8_MGAS9429.crt.3 | NOHIT | 0 | | | | | | | | | | | | | |
| 8_MGAS9429.crt.4 | NOHIT | 32 | | | | | | | | | | | | | |
| 8_MGAS9429.crt.5 | BNEKOFAL_00674 hypothetical protein | 34 | 285 | 63.9 | 7.00E-11 | 34 | 100% | 0 | 0% | 1 | 34 | 100% | 19 | 52 + | 19 |
| 8_MGAS9429.crt.5 | FOAEMPCG_01040 hypothetical protein | 34 | 285 | 63.9 | 7.00E-11 | 34 | 100% | 0 | 0% | 1 | 34 | 100% | 19 | 52 + | 19 |
| 8_MGAS9429.crt.5 | MCEIOEBL_00624 hypothetical protein | 34 | 285 | 63.9 | 7.00E-11 | 34 | 100% | 0 | 0% | 1 | 34 | 100% | 19 | 52 + | 19 |
| 8_MGAS9429.crt.5 | PPKGMMPD_01012 hypothetical protein | 34 | 285 | 63.9 | 7.00E-11 | 34 | 100% | 0 | 0% | 1 | 34 | 100% | 19 | 52 + | 19 |
| 8_MGAS9429.crt.5 | LLAAGEIN_01368 hypothetical protein | 34 | 285 | 63.9 | 7.00E-11 | 34 | 100% | 0 | 0% | 1 | 34 | 100% | 19 | 52 + | 19 |
| 8_MGAS9429.crt.5 | AIKJPEBD_01573 hypothetical protein | 34 | 285 | 63.9 | 7.00E-11 | 34 | 100% | 0 | 0% | 1 | 34 | 100% | 19 | 52 + | 19 |
| 8_MGAS9429.crt.5 | FPFNMMFM_00882 hypothetical protein | 34 | 285 | 63.9 | 7.00E-11 | 34 | 100% | 0 | 0% | 1 | 34 | 100% | 19 | 52 + | 19 |
| 8_MGAS9429.crt.5 | IJFGMCCO_00995 hypothetical protein | 34 | 285 | 63.9 | 7.00E-11 | 34 | 100% | 0 | 0% | 1 | 34 | 100% | 19 | 52 + | 19 |
| 8_MGAS9429.crt.5 | EJBPJIHF_01045 hypothetical protein | 34 | 285 | 63.9 | 7.00E-11 | 34 | 100% | 0 | 0% | 1 | 34 | 100% | 19 | 52 + | 19 |
| 8_MGAS9429.crt.5 | LAKBNEPL_01011 hypothetical protein | 34 | 285 | 63.9 | 7.00E-11 | 34 | 100% | 0 | 0% | 1 | 34 | 100% | 19 | 52 + | 19 |
| 8_MGAS9429.crt.5 | MMBCLDPD_00865 hypothetical protein | 34 | 285 | 63.9 | 7.00E-11 | 34 | 100% | 0 | 0% | 1 | 34 | 100% | 19 | 52 + | 19 |
| 8_MGAS9429.crt.5 | PLPKHLCC_00893 hypothetical protein | 34 | 285 | 63.9 | 7.00E-11 | 34 | 100% | 0 | 0% | 1 | 34 | 100% | 19 | 52 + | 19 |
| 8_MGAS9429.crt.5 | JNLHFPPN_01263 hypothetical protein | 34 | 285 | 63.9 | 7.00E-11 | 34 | 100% | 0 | 0% | 1 | 34 | 100% | 19 | 52 + | 19 |
| 8_MGAS9429.crt.5 | ODIAAKBJ_01263 hypothetical protein | 34 | 285 | 63.9 | 7.00E-11 | 34 | 100% | 0 | 0% | 1 | 34 | 100% | 19 | 52 + | 19 |
| 8_MGAS9429.crt.5 | ECKACGMC_00553 hypothetical protein | 34 | 285 | 63.9 | 7.00E-11 | 34 | 100% | 0 | 0% | 1 | 34 | 100% | 19 | 52 + | 19 |
| 8_MGAS9429.crt.5 | DIPDIGDN_01262 hypothetical protein | 34 | 285 | 63.9 | 7.00E-11 | 34 | 100% | 0 | 0% | 1 | 34 | 100% | 19 | 52 + | 19 |
| 8_MGAS9429.crt.5 | EFGBCCNL_00703 hypothetical protein | 34 | 285 | 63.9 | 7.00E-11 | 34 | 100% | 0 | 0% | 1 | 34 | 100% | 19 | 52 + | 19 |
| 8_MGAS9429.crt.5 | JHELIIPC_00975 hypothetical protein | 34 | 285 | 63.9 | 7.00E-11 | 34 | 100% | 0 | 0% | 1 | 34 | 100% | 19 | 52 + | 19 |
| 8_MGAS9429.crt.5 | GPBJDOFB_00997 hypothetical protein | 34 | 285 | 63.9 | 7.00E-11 | 34 | 100% | 0 | 0% | 1 | 34 | 100% | 19 | 52 + | 19 |
| 8_MGAS9429.crt.5 | ACJECOPP_01021 hypothetical protein | 34 | 285 | 63.9 | 7.00E-11 | 34 | 100% | 0 | 0% | 1 | 34 | 100% | 19 | 52 + | 19 |
| 8_MGAS9429.crt.5 | ICMALKPE_01033 hypothetical protein | 34 | 285 | 63.9 | 7.00E-11 | 34 | 100% | 0 | 0% | 1 | 34 | 100% | 19 | 52 + | 19 |
| 8_MGAS9429.crt.5 | AJHAHHEL_01315 hypothetical protein | 34 | 285 | 63.9 | 7.00E-11 | 34 | 100% | 0 | 0% | 1 | 34 | 100% | 19 | 52 + | 19 |
| 8_MGAS9429.crt.5 | DMLKMGFL_01490 hypothetical protein | 34 | 285 | 63.9 | 7.00E-11 | 34 | 100% | 0 | 0% | 1 | 34 | 100% | 19 | 52 + | 19 |
| 8_MGAS9429.crt.5 | HEGDFFNC_00554 hypothetical protein | 34 | 285 | 63.9 | 7.00E-11 | 34 | 100% | 0 | 0% | 1 | 34 | 100% | 19 | 52 + | 19 |
| 8_MGAS9429.crt.5 | IIJJCGIL_01024 hypothetical protein | 34 | 285 | 63.9 | 7.00E-11 | 34 | 100% | 0 | 0% | 1 | 34 | 100% | 19 | 52 + | 19 |
| 8_MGAS9429.crt.5 | GLBJHDNN_01015 hypothetical protein | 34 | 285 | 63.9 | 7.00E-11 | 34 | 100% | 0 | 0% | 1 | 34 | 100% | 19 | 52 + | 19 |
| 8_MGAS9429.crt.5 | HPLLMOOM_00975 hypothetical protein | 34 | 285 | 63.9 | 7.00E-11 | 34 | 100% | 0 | 0% | 1 | 34 | 100% | 19 | 52 + | 19 |
| 8_MGAS9429.crt.5 | MCFNMICF_01027 hypothetical protein | 34 | 285 | 63.9 | 7.00E-11 | 34 | 100% | 0 | 0% | 1 | 34 | 100% | 19 | 52 + | 19 |
| 8_MGAS9429.crt.5 | MONDJAPC_01581 hypothetical protein | 34 | 285 | 63.9 | 7.00E-11 | 34 | 100% | 0 | 0% | 1 | 34 | 100% | 19 | 52 + | 19 |
| 8_MGAS9429.crt.6 | JEOJEKGO_01460 hypothetical protein | 34 | 306 | 63.9 | 7.00E-11 | 34 | 100% | 0 | 0% | 1 | 34 | 100% | 146 | 179 + | 146 |
| 8_MGAS9429.crt.6 | DCKFNFNP_00568 hypothetical protein | 34 | 306 | 63.9 | 7.00E-11 | 34 | 100% | 0 | 0% | 1 | 34 | 100% | 146 | 179 + | 146 |
| 8_MGAS9429.crt.6 | ECKACGMC_01462 hypothetical protein | 34 | 306 | 63.9 | 7.00E-11 | 34 | 100% | 0 | 0% | 1 | 34 | 100% | 146 | 179 + | 146 |
| 8_MGAS9429.crt.6 | MDEAFGPO_00191 hypothetical protein | 34 | 306 | 63.9 | 7.00E-11 | 34 | 100% | 0 | 0% | 1 | 34 | 100% | 146 | 179 + | 146 |
| 8_MGAS9429.crt.6 | HEGDFFNC_01370 hypothetical protein | 34 | 306 | 63.9 | 7.00E-11 | 34 | 100% | 0 | 0% | 1 | 34 | 100% | 146 | 179 + | 146 |
| 8_MGAS9429.crt.6 | MONDJAPC_00627 hypothetical protein | 34 | 306 | 63.9 | 7.00E-11 | 34 | 100% | 0 | 0% | 1 | 34 | 100% | 146 | 179 + | 146 |
| 8_MGAS9429.crt.7 | NOHIT | 34 | | | | | | | | | | | | | |
| 8_MGAS9429.crt.8 | EJBPJIHF_01284 hypothetical protein | 35 | 381 | 65.8 | 2.00E-11 | 35 | 100% | 0 | 0% | 1 | 35 | 100% | 156 | 190 - | 156 |
| 8_MGAS9429.crt.8 | MMBCLDPD_00678 hypothetical protein | 35 | 381 | 65.8 | 2.00E-11 | 35 | 100% | 0 | 0% | 1 | 35 | 100% | 156 | 190 - | 156 |
| 8_MGAS9429.crt.8 | GGLKEJFF_01158 hypothetical protein | 35 | 381 | 65.8 | 2.00E-11 | 35 | 100% | 0 | 0% | 1 | 35 | 100% | 156 | 190 - | 156 |
| 8_MGAS9429.crt.8 | GIKPJECN_01232 hypothetical protein | 35 | 381 | 65.8 | 2.00E-11 | 35 | 100% | 0 | 0% | 1 | 35 | 100% | 156 | 190 - | 156 |
| 8_MGAS9429.crt.8 | GPBJDOFB_01192 hypothetical protein | 35 | 381 | 65.8 | 2.00E-11 | 35 | 100% | 0 | 0% | 1 | 35 | 100% | 156 | 190 - | 156 |
| 8_MGAS9429.crt.8 | ACJECOPP_01208 hypothetical protein | 35 | 381 | 65.8 | 2.00E-11 | 35 | 100% | 0 | 0% | 1 | 35 | 100% | 156 | 190 - | 156 |
| 8_MGAS9429.crt.8 | ICMALKPE_01219 hypothetical protein | 35 | 381 | 65.8 | 2.00E-11 | 35 | 100% | 0 | 0% | 1 | 35 | 100% | 156 | 190 - | 156 |
| 8_MGAS9429.crt.8 | IIJJCGIL_01210 hypothetical protein | 35 | 381 | 65.8 | 2.00E-11 | 35 | 100% | 0 | 0% | 1 | 35 | 100% | 156 | 190 - | 156 |
| 8_MGAS9429.crt.9 | FOAEMPCG_00787 eukaryotic hypersensitive-induced | 35 | 828 | 65.8 | 2.00E-11 | 35 | 100% | 0 | 0% | 1 | 35 | 100% | 511 | 545 + | 511 |
| 8_MGAS9429.crt.9 | LLAAGEIN_00659 eukaryotic hypersensitive-induced | 35 | 828 | 65.8 | 2.00E-11 | 35 | 100% | 0 | 0% | 1 | 35 | 100% | 511 | 545 + | 511 |
| 8_MGAS9429.crt.9 | EFGBCCNL_00468 eukaryotic hypersensitive-induced | 35 | 828 | 65.8 | 2.00E-11 | 35 | 100% | 0 | 0% | 1 | 35 | 100% | 511 | 545 + | 511 |
| 8_MGAS9429.crt.9 | JHELIIPC_01229 eukaryotic hypersensitive-induced re | 35 | 828 | 65.8 | 2.00E-11 | 35 | 100% | 0 | 0% | 1 | 35 | 100% | 511 | 545 + | 511 |
| 8_MGAS9429.crt.9 | HPLLMOOM_01228 eukaryotic hypersensitive-induce | 35 | 828 | 65.8 | 2.00E-11 | 35 | 100% | 0 | 0% | 1 | 35 | 100% | 511 | 545 + | 511 |
| 8_MGAS9429.crt.9 | MCFNMICF_00775 eukaryotic hypersensitive-induced | 35 | 828 | 65.8 | 2.00E-11 | 35 | 100% | 0 | 0% | 1 | 35 | 100% | 511 | 545 + | 511 |
| 8_MGAS9429.crt.9 | MONDJAPC_00342 eukaryotic hypersensitive-induced | 35 | 828 | 65.8 | 2.00E-11 | 35 | 100% | 0 | 0% | 1 | 35 | 100% | 511 | 545 + | 511 |
| 8_MGAS9429.crt.10 | NOHIT | 33 | | | | | | | | | | | | | |
| 9_MGAS10270.crt.1 | NOHIT | 37 | | | | | | | | | | | | | |
| 9_MGAS10270.crt.2 | NOHIT | 37 | | | | | | | | | | | | | |
| 9_MGAS10270.crt.3 | NOHIT | 0 | | | | | | | | | | | | | |
| 9_MGAS10270.crt.4 | MHKCMMMI_00745 hypothetical protein | 34 | 273 | 63.9 | 7.00E-11 | 34 | 100% | 0 | 0% | 1 | 34 | 100% | 230 | 263 + | 230 |
| 9_MGAS10270.crt.4 | FOAEMPCG_01014 hypothetical protein | 34 | 273 | 63.9 | 7.00E-11 | 34 | 100% | 0 | 0% | 1 | 34 | 100% | 230 | 263 + | 230 |
| 9_MGAS10270.crt.4 | FOAEMPCG_00816 hypothetical protein | 34 | 273 | 63.9 | 7.00E-11 | 34 | 100% | 0 | 0% | 1 | 34 | 100% | 230 | 263 + | 230 |
| 9_MGAS10270.crt.4 | CMOPCPHD_00651 hypothetical protein | 34 | 273 | 63.9 | 7.00E-11 | 34 | 100% | 0 | 0% | 1 | 34 | 100% | 230 | 263 + | 230 |
| 9_MGAS10270.crt.4 | LLAAGEIN_01206 hypothetical protein | 34 | 273 | 63.9 | 7.00E-11 | 34 | 100% | 0 | 0% | 1 | 34 | 100% | 230 | 263 + | 230 |
| 9_MGAS10270.crt.4 | LLAAGEIN_00632 hypothetical protein | 34 | 273 | 63.9 | 7.00E-11 | 34 | 100% | 0 | 0% | 1 | 34 | 100% | 230 | 263 + | 230 |
| 9_MGAS10270.crt.4 | FDNDANMJ_01217 hypothetical protein | 34 | 273 | 63.9 | 7.00E-11 | 34 | 100% | 0 | 0% | 1 | 34 | 100% | 230 | 263 + | 230 |
| 9_MGAS10270.crt.4 | JEOJEKGO_01405 hypothetical protein | 34 | 273 | 63.9 | 7.00E-11 | 34 | 100% | 0 | 0% | 1 | 34 | 100% | 230 | 263 + | 230 |
| 9_MGAS10270.crt.4 | DCKFNFNP_01239 hypothetical protein | 34 | 273 | 63.9 | 7.00E-11 | 34 | 100% | 0 | 0% | 1 | 34 | 100% | 230 | 263 + | 230 |
| 9_MGAS10270.crt.4 | DCKFNFNP_00618 hypothetical protein | 34 | 273 | 63.9 | 7.00E-11 | 34 | 100% | 0 | 0% | 1 | 34 | 100% | 230 | 263 + | 230 |
| 9_MGAS10270.crt.4 | JNLHFPPN_01474 hypothetical protein | 34 | 273 | 63.9 | 7.00E-11 | 34 | 100% | 0 | 0% | 1 | 34 | 100% | 230 | 263 + | 230 |
| 9_MGAS10270.crt.4 | ODIAAKBJ_01474 hypothetical protein | 34 | 273 | 63.9 | 7.00E-11 | 34 | 100% | 0 | 0% | 1 | 34 | 100% | 230 | 263 + | 230 |
| 9_MGAS10270.crt.4 | ECKACGMC_01122 hypothetical protein | 34 | 273 | 63.9 | 7.00E-11 | 34 | 100% | 0 | 0% | 1 | 34 | 100% | 230 | 263 + | 230 |
| 9_MGAS10270.crt.4 | DIPDIGDN_01475 hypothetical protein | 34 | 273 | 63.9 | 7.00E-11 | 34 | 100% | 0 | 0% | 1 | 34 | 100% | 230 | 263 + | 230 |
| 9_MGAS10270.crt.4 | GBCACICN_01185 hypothetical protein | 34 | 273 | 63.9 | 7.00E-11 | 34 | 100% | 0 | 0% | 1 | 34 | 100% | 230 | 263 + | 230 |
| 9_MGAS10270.crt.4 | MDEAFGPO_00493 hypothetical protein | 34 | 273 | 63.9 | 7.00E-11 | 34 | 100% | 0 | 0% | 1 | 34 | 100% | 230 | 263 + | 230 |
| 9_MGAS10270.crt.4 | EFGBCCNL_00440 hypothetical protein | 34 | 273 | 63.9 | 7.00E-11 | 34 | 100% | 0 | 0% | 1 | 34 | 100% | 230 | 263 + | 230 |
| 9_MGAS10270.crt.4 | GPBJDOFB_01477 hypothetical protein | 34 | 273 | 63.9 | 7.00E-11 | 34 | 100% | 0 | 0% | 1 | 34 | 100% | 230 | 263 + | 230 |
| 9_MGAS10270.crt.4 | AJHAHHEL_01152 hypothetical protein | 34 | 273 | 63.9 | 7.00E-11 | 34 | 100% | 0 | 0% | 1 | 34 | 100% | 230 | 263 + | 230 |
| 9_MGAS10270.crt.4 | IPNPMFKD_01279 hypothetical protein | 34 | 273 | 63.9 | 7.00E-11 | 34 | 100% | 0 | 0% | 1 | 34 | 100% | 230 | 263 + | 230 |
| 9_MGAS10270.crt.4 | HPLLMOOM_01199 hypothetical protein | 34 | 273 | 63.9 | 7.00E-11 | 34 | 100% | 0 | 0% | 1 | 34 | 100% | 230 | 263 + | 230 |
| 9_MGAS10270.crt.4 | MCFNMICF_00803 hypothetical protein | 34 | 273 | 63.9 | 7.00E-11 | 34 | 100% | 0 | 0% | 1 | 34 | 100% | 230 | 263 + | 230 |
| 9_MGAS10270.crt.4 | MONDJAPC_00369 hypothetical protein | 34 | 273 | 63.9 | 7.00E-11 | 34 | 100% | 0 | 0% | 1 | 34 | 100% | 230 | 263 + | 230 |
| 9_MGAS10270.crt.5 | BNEKOFAL_01159 putative repressor protein | 34 | 789 | 63.9 | 7.00E-11 | 34 | 100% | 0 | 0% | 1 | 34 | 100% | 400 | 433 - | 400 |
| 9_MGAS10270.crt.5 | MHKCMMMI_01476 putative repressor protein | 34 | 789 | 63.9 | 7.00E-11 | 34 | 100% | 0 | 0% | 1 | 34 | 100% | 400 | 433 - | 400 |
| 9_MGAS10270.crt.5 | FOAEMPCG_01065 putative repressor protein | 34 | 789 | 63.9 | 7.00E-11 | 34 | 100% | 0 | 0% | 1 | 34 | 100% | 400 | 433 - | 400 |
| 9_MGAS10270.crt.5 | MCEIOEBL_01108 putative repressor protein | 34 | 789 | 63.9 | 7.00E-11 | 34 | 100% | 0 | 0% | 1 | 34 | 100% | 400 | 433 - | 400 |
| 9_MGAS10270.crt.5 | LLAAGEIN_01091 putative repressor protein | 34 | 789 | 63.9 | 7.00E-11 | 34 | 100% | 0 | 0% | 1 | 34 | 100% | 400 | 433 - | 400 |
| 9_MGAS10270.crt.5 | JEOJEKGO_01459 putative repressor protein | 34 | 765 | 63.9 | 7.00E-11 | 34 | 100% | 0 | 0% | 1 | 34 | 100% | 376 | 409 - | 376 |
| 9_MGAS10270.crt.5 | DCKFNFNP_00569 putative repressor protein | 34 | 765 | 63.9 | 7.00E-11 | 34 | 100% | 0 | 0% | 1 | 34 | 100% | 376 | 409 - | 376 |
| 9_MGAS10270.crt.5 | ECKACGMC_01461 putative repressor protein | 34 | 765 | 63.9 | 7.00E-11 | 34 | 100% | 0 | 0% | 1 | 34 | 100% | 376 | 409 - | 376 |
| 9_MGAS10270.crt.5 | DDKNKGAM_00756 putative repressor protein | 34 | 789 | 63.9 | 7.00E-11 | 34 | 100% | 0 | 0% | 1 | 34 | 100% | 400 | 433 - | 400 |
| 9_MGAS10270.crt.5 | MDEAFGPO_00192 putative repressor protein | 34 | 765 | 63.9 | 7.00E-11 | 34 | 100% | 0 | 0% | 1 | 34 | 100% | 376 | 409 - | 376 |
| 9_MGAS10270.crt.5 | EFGBCCNL_00296 putative repressor protein | 34 | 789 | 63.9 | 7.00E-11 | 34 | 100% | 0 | 0% | 1 | 34 | 100% | 400 | 433 - | 400 |
| 9_MGAS10270.crt.5 | DEMOPPLM_00869 putative repressor protein | 34 | 789 | 63.9 | 7.00E-11 | 34 | 100% | 0 | 0% | 1 | 34 | 100% | 400 | 433 - | 400 |
| 9_MGAS10270.crt.5 | JHELIIPC_00950 putative repressor protein | 34 | 789 | 63.9 | 7.00E-11 | 34 | 100% | 0 | 0% | 1 | 34 | 100% | 400 | 433 - | 400 |
| 9_MGAS10270.crt.5 | IFBBIBNC_01136 putative repressor protein | 34 | 789 | 63.9 | 7.00E-11 | 34 | 100% | 0 | 0% | 1 | 34 | 100% | 400 | 433 - | 400 |
| 9_MGAS10270.crt.5 | HEGDFFNC_01369 putative repressor protein | 34 | 765 | 63.9 | 7.00E-11 | 34 | 100% | 0 | 0% | 1 | 34 | 100% | 376 | 409 - | 376 |
| 9_MGAS10270.crt.5 | NELECJBE_01048 putative repressor protein | 34 | 789 | 63.9 | 7.00E-11 | 34 | 100% | 0 | 0% | 1 | 34 | 100% | 400 | 433 - | 400 |
| 9_MGAS10270.crt.5 | HPLLMOOM_00950 putative repressor protein | 34 | 789 | 63.9 | 7.00E-11 | 34 | 100% | 0 | 0% | 1 | 34 | 100% | 400 | 433 - | 400 |
| 9_MGAS10270.crt.5 | MCFNMICF_01052 putative repressor protein | 34 | 789 | 63.9 | 7.00E-11 | 34 | 100% | 0 | 0% | 1 | 34 | 100% | 400 | 433 - | 400 |
| 9_MGAS10270.crt.5 | MONDJAPC_00628 putative repressor protein | 34 | 765 | 63.9 | 7.00E-11 | 34 | 100% | 0 | 0% | 1 | 34 | 100% | 376 | 409 - | 376 |
| 9_MGAS10270.crt.6 | NOHIT | 32 | | | | | | | | | | | | | |

**Supplementary Table S5. BLASTN outputs of spacer sequences against predicted RM systems in 59 *S. pyogenes* strains**

| query_name | subject_name | query_length | subject_length | score | E-value | identity | identity_per | gap | gap_per | query_start | query_end | coverage | subject_start | subject_end | subject_strand | subject_index |
|---|---|---|---|---|---|---|---|---|---|---|---|---|---|---|---|---|
| 1_SF370.crt.10 | M.Spy19615ORF703P: WARNING1, ID: 99.18%, HSP/Length: 486/486, Contig name: A06_1, Position: 1008224..1008709 | 34 | 486 | 58.4 | 2.00E-11 | 34 | 97% | 1 | 3% | 1 | 34 | 100% | 137 | 171 | + | 137 |
| 1_SF370.crt.10 | M.Spy19615ORF703P: WARNING1, ID: 99.18%, HSP/Length: 486/486, Contig name: A09_1, Position: 1046472..1046957 | 34 | 486 | 58.4 | 2.00E-11 | 34 | 97% | 1 | 3% | 1 | 34 | 100% | 137 | 171 | + | 137 |
| 1_SF370.crt.10 | M.Spy19615ORF703P: WARNING1, ID: 99.18%, HSP/Length: 486/486, Contig name: CP013672.1, Position: 1031405..1031890 | 34 | 486 | 58.4 | 2.00E-11 | 34 | 97% | 1 | 3% | 1 | 34 | 100% | 137 | 171 | + | 137 |
| 17_M1_476.crt.8 | M.Spy19615ORF703P: WARNING1, ID: 99.18%, HSP/Length: 486/486, Contig name: A06_1, Position: 1008224..1008709 | 35 | 486 | 60.2 | 7.00E-12 | 34 | 97% | 0 | 0% | 1 | 35 | 100% | 137 | 171 | + | 137 |
| 17_M1_476.crt.8 | M.Spy19615ORF703P: WARNING1, ID: 99.18%, HSP/Length: 486/486, Contig name: A09_1, Position: 1046472..1046957 | 35 | 486 | 60.2 | 7.00E-12 | 34 | 97% | 0 | 0% | 1 | 35 | 100% | 137 | 171 | + | 137 |
| 17_M1_476.crt.8 | M.Spy19615ORF703P: WARNING1, ID: 99.18%, HSP/Length: 486/486, Contig name: CP013672.1, Position: 1031405..1031890 | 35 | 486 | 60.2 | 7.00E-12 | 34 | 97% | 0 | 0% | 1 | 35 | 100% | 137 | 171 | + | 137 |
| 18_A20.crt.8 | M.Spy19615ORF703P: WARNING1, ID: 99.18%, HSP/Length: 486/486, Contig name: A06_1, Position: 1008224..1008709 | 35 | 486 | 60.2 | 7.00E-12 | 34 | 97% | 0 | 0% | 1 | 35 | 100% | 137 | 171 | + | 137 |
| 18_A20.crt.8 | M.Spy19615ORF703P: WARNING1, ID: 99.18%, HSP/Length: 486/486, Contig name: A09_1, Position: 1046472..1046957 | 35 | 486 | 60.2 | 7.00E-12 | 34 | 97% | 0 | 0% | 1 | 35 | 100% | 137 | 171 | + | 137 |
| 18_A20.crt.8 | M.Spy19615ORF703P: WARNING1, ID: 99.18%, HSP/Length: 486/486, Contig name: CP013672.1, Position: 1031405..1031890 | 35 | 486 | 60.2 | 7.00E-12 | 34 | 97% | 0 | 0% | 1 | 35 | 100% | 137 | 171 | + | 137 |
| 28_MTB313.crt.8 | M.Spy19615ORF703P: WARNING1, ID: 99.18%, HSP/Length: 486/486, Contig name: A06_1, Position: 1008224..1008709 | 35 | 486 | 60.2 | 7.00E-12 | 34 | 97% | 0 | 0% | 1 | 35 | 100% | 137 | 171 | + | 137 |
| 28_MTB313.crt.8 | M.Spy19615ORF703P: WARNING1, ID: 99.18%, HSP/Length: 486/486, Contig name: A09_1, Position: 1046472..1046957 | 35 | 486 | 60.2 | 7.00E-12 | 34 | 97% | 0 | 0% | 1 | 35 | 100% | 137 | 171 | + | 137 |
| 28_MTB313.crt.8 | M.Spy19615ORF703P: WARNING1, ID: 99.18%, HSP/Length: 486/486, Contig name: CP013672.1, Position: 1031405..1031890 | 35 | 486 | 60.2 | 7.00E-12 | 34 | 97% | 0 | 0% | 1 | 35 | 100% | 137 | 171 | + | 137 |
| 29_MTB314.crt.8 | M.Spy19615ORF703P: WARNING1, ID: 99.18%, HSP/Length: 486/486, Contig name: A06_1, Position: 1008224..1008709 | 35 | 486 | 60.2 | 7.00E-12 | 34 | 97% | 0 | 0% | 1 | 35 | 100% | 137 | 171 | + | 137 |
| 29_MTB314.crt.8 | M.Spy19615ORF703P: WARNING1, ID: 99.18%, HSP/Length: 486/486, Contig name: A09_1, Position: 1046472..1046957 | 35 | 486 | 60.2 | 7.00E-12 | 34 | 97% | 0 | 0% | 1 | 35 | 100% | 137 | 171 | + | 137 |
| 29_MTB314.crt.8 | M.Spy19615ORF703P: WARNING1, ID: 99.18%, HSP/Length: 486/486, Contig name: CP013672.1, Position: 1031405..1031890 | 35 | 486 | 60.2 | 7.00E-12 | 34 | 97% | 0 | 0% | 1 | 35 | 100% | 137 | 171 | + | 137 |
| 31_AP1.crt.1 | M.Spy19615ORF703P: WARNING1, ID: 99.18%, HSP/Length: 486/486, Contig name: A06_1, Position: 1008224..1008709 | 35 | 486 | 60.2 | 7.00E-12 | 34 | 97% | 0 | 0% | 1 | 35 | 100% | 137 | 171 | - | 137 |
| 31_AP1.crt.1 | M.Spy19615ORF703P: WARNING1, ID: 99.18%, HSP/Length: 486/486, Contig name: A09_1, Position: 1046472..1046957 | 35 | 486 | 60.2 | 7.00E-12 | 34 | 97% | 0 | 0% | 1 | 35 | 100% | 137 | 171 | - | 137 |
| 31_AP1.crt.1 | M.Spy19615ORF703P: WARNING1, ID: 99.18%, HSP/Length: 486/486, Contig name: CP013672.1, Position: 1031405..1031890 | 35 | 486 | 60.2 | 7.00E-12 | 34 | 97% | 0 | 0% | 1 | 35 | 100% | 137 | 171 | - | 137 |
| 38_5448.crt.1 | M.Spy19615ORF703P: WARNING1, ID: 99.18%, HSP/Length: 486/486, Contig name: A06_1, Position: 1008224..1008709 | 35 | 486 | 60.2 | 7.00E-12 | 34 | 97% | 0 | 0% | 1 | 35 | 100% | 137 | 171 | - | 137 |
| 38_5448.crt.1 | M.Spy19615ORF703P: WARNING1, ID: 99.18%, HSP/Length: 486/486, Contig name: A09_1, Position: 1046472..1046957 | 35 | 486 | 60.2 | 7.00E-12 | 34 | 97% | 0 | 0% | 1 | 35 | 100% | 137 | 171 | - | 137 |
| 38_5448.crt.1 | M.Spy19615ORF703P: WARNING1, ID: 99.18%, HSP/Length: 486/486, Contig name: CP013672.1, Position: 1031405..1031890 | 35 | 486 | 60.2 | 7.00E-12 | 34 | 97% | 0 | 0% | 1 | 35 | 100% | 137 | 171 | - | 137 |
| 40_HKU488.crt.8 | M.Spy19615ORF703P: WARNING1, ID: 99.18%, HSP/Length: 486/486, Contig name: A06_1, Position: 1008224..1008709 | 35 | 486 | 60.2 | 7.00E-12 | 34 | 97% | 0 | 0% | 1 | 35 | 100% | 137 | 171 | + | 137 |
| 40_HKU488.crt.8 | M.Spy19615ORF703P: WARNING1, ID: 99.18%, HSP/Length: 486/486, Contig name: A09_1, Position: 1046472..1046957 | 35 | 486 | 60.2 | 7.00E-12 | 34 | 97% | 0 | 0% | 1 | 35 | 100% | 137 | 171 | + | 137 |
| 40_HKU488.crt.8 | M.Spy19615ORF703P: WARNING1, ID: 99.18%, HSP/Length: 486/486, Contig name: CP013672.1, Position: 1031405..1031890 | 35 | 486 | 60.2 | 7.00E-12 | 34 | 97% | 0 | 0% | 1 | 35 | 100% | 137 | 171 | + | 137 |
| 45_FDAARGOS_149.crt.1 | M.Spy19615ORF703P: WARNING1, ID: 99.18%, HSP/Length: 486/486, Contig name: A06_1, Position: 1008224..1008709 | 35 | 486 | 60.2 | 7.00E-12 | 34 | 97% | 0 | 0% | 1 | 35 | 100% | 137 | 171 | - | 137 |
| 45_FDAARGOS_149.crt.1 | M.Spy19615ORF703P: WARNING1, ID: 99.18%, HSP/Length: 486/486, Contig name: A09_1, Position: 1046472..1046957 | 35 | 486 | 60.2 | 7.00E-12 | 34 | 97% | 0 | 0% | 1 | 35 | 100% | 137 | 171 | - | 137 |
| 45_FDAARGOS_149.crt.1 | M.Spy19615ORF703P: WARNING1, ID: 99.18%, HSP/Length: 486/486, Contig name: CP013672.1, Position: 1031405..1031890 | 35 | 486 | 60.2 | 7.00E-12 | 34 | 97% | 0 | 0% | 1 | 35 | 100% | 137 | 171 | - | 137 |
| 7_MGAS5005.crt.8 | M.Spy19615ORF703P: WARNING1, ID: 99.18%, HSP/Length: 486/486, Contig name: A06_1, Position: 1008224..1008709 | 35 | 486 | 60.2 | 7.00E-12 | 34 | 97% | 0 | 0% | 1 | 35 | 100% | 137 | 171 | + | 137 |
| 7_MGAS5005.crt.8 | M.Spy19615ORF703P: WARNING1, ID: 99.18%, HSP/Length: 486/486, Contig name: A09_1, Position: 1046472..1046957 | 35 | 486 | 60.2 | 7.00E-12 | 34 | 97% | 0 | 0% | 1 | 35 | 100% | 137 | 171 | + | 137 |
| 7_MGAS5005.crt.8 | M.Spy19615ORF703P: WARNING1, ID: 99.18%, HSP/Length: 486/486, Contig name: CP013672.1, Position: 1031405..1031890 | 35 | 486 | 60.2 | 7.00E-12 | 34 | 97% | 0 | 0% | 1 | 35 | 100% | 137 | 171 | + | 137 |
| 1_SF370.crt.10 | M.SpyM23NDORF616P: PERFECT MATCH, ID: 100.00%, HSP/Length: 444/444, Contig name: CP008695.1, Position: 180965..181408 | 34 | 444 | 58.4 | 2.00E-11 | 34 | 97% | 1 | 3% | 1 | 34 | 100% | 137 | 171 | + | 137 |
| 1_SF370.crt.10 | M.SpyM23NDORF616P: PERFECT MATCH, ID: 100.00%, HSP/Length: 444/444, Contig name: CP008695.1, Position: 585419..585862 | 34 | 444 | 58.4 | 2.00E-11 | 34 | 97% | 1 | 3% | 1 | 34 | 100% | 137 | 171 | + | 137 |
| 1_SF370.crt.10 | M.SpyM23NDORF616P: WARNING2, ID: 100.00%, HSP/Length: 419/444, Contig name: CP006366.1, Position: 1483939..1484357 | 34 | 444 | 58.4 | 2.00E-11 | 34 | 97% | 1 | 3% | 1 | 34 | 100% | 137 | 171 | + | 137 |
| 1_SF370.crt.10 | M.SpyM23NDORF616P: WARNING2, ID: 100.00%, HSP/Length: 419/444, Contig name: CP009612.1, Position: 1222659..1223077 | 34 | 444 | 58.4 | 2.00E-11 | 34 | 97% | 1 | 3% | 1 | 34 | 100% | 137 | 171 | + | 137 |
| 17_M1_476.crt.8 | M.SpyM23NDORF616P: PERFECT MATCH, ID: 100.00%, HSP/Length: 444/444, Contig name: CP008695.1, Position: 180965..181408 | 35 | 444 | 60.2 | 7.00E-12 | 34 | 97% | 0 | 0% | 1 | 35 | 100% | 137 | 171 | + | 137 |
| 17_M1_476.crt.8 | M.SpyM23NDORF616P: PERFECT MATCH, ID: 100.00%, HSP/Length: 444/444, Contig name: CP008695.1, Position: 585419..585862 | 35 | 444 | 60.2 | 7.00E-12 | 34 | 97% | 0 | 0% | 1 | 35 | 100% | 137 | 171 | + | 137 |
| 17_M1_476.crt.8 | M.SpyM23NDORF616P: WARNING2, ID: 100.00%, HSP/Length: 419/444, Contig name: CP006366.1, Position: 1483939..1484357 | 35 | 444 | 60.2 | 7.00E-12 | 34 | 97% | 0 | 0% | 1 | 35 | 100% | 137 | 171 | + | 137 |
| 17_M1_476.crt.8 | M.SpyM23NDORF616P: WARNING2, ID: 100.00%, HSP/Length: 419/444, Contig name: CP009612.1, Position: 1222659..1223077 | 35 | 444 | 60.2 | 7.00E-12 | 34 | 97% | 0 | 0% | 1 | 35 | 100% | 137 | 171 | + | 137 |
| 18_A20.crt.8 | M.SpyM23NDORF616P: PERFECT MATCH, ID: 100.00%, HSP/Length: 444/444, Contig name: CP008695.1, Position: 180965..181408 | 35 | 444 | 60.2 | 7.00E-12 | 34 | 97% | 0 | 0% | 1 | 35 | 100% | 137 | 171 | + | 137 |
| 18_A20.crt.8 | M.SpyM23NDORF616P: PERFECT MATCH, ID: 100.00%, HSP/Length: 444/444, Contig name: CP008695.1, Position: 585419..585862 | 35 | 444 | 60.2 | 7.00E-12 | 34 | 97% | 0 | 0% | 1 | 35 | 100% | 137 | 171 | + | 137 |
| 18_A20.crt.8 | M.SpyM23NDORF616P: WARNING2, ID: 100.00%, HSP/Length: 419/444, Contig name: CP006366.1, Position: 1483939..1484357 | 35 | 444 | 60.2 | 7.00E-12 | 34 | 97% | 0 | 0% | 1 | 35 | 100% | 137 | 171 | + | 137 |
| 18_A20.crt.8 | M.SpyM23NDORF616P: WARNING2, ID: 100.00%, HSP/Length: 419/444, Contig name: CP009612.1, Position: 1222659..1223077 | 35 | 444 | 60.2 | 7.00E-12 | 34 | 97% | 0 | 0% | 1 | 35 | 100% | 137 | 171 | + | 137 |
| 28_MTB313.crt.8 | M.SpyM23NDORF616P: PERFECT MATCH, ID: 100.00%, HSP/Length: 444/444, Contig name: CP008695.1, Position: 180965..181408 | 35 | 444 | 60.2 | 7.00E-12 | 34 | 97% | 0 | 0% | 1 | 35 | 100% | 137 | 171 | + | 137 |
| 28_MTB313.crt.8 | M.SpyM23NDORF616P: PERFECT MATCH, ID: 100.00%, HSP/Length: 444/444, Contig name: CP008695.1, Position: 585419..585862 | 35 | 444 | 60.2 | 7.00E-12 | 34 | 97% | 0 | 0% | 1 | 35 | 100% | 137 | 171 | + | 137 |
| 28_MTB313.crt.8 | M.SpyM23NDORF616P: WARNING2, ID: 100.00%, HSP/Length: 419/444, Contig name: CP006366.1, Position: 1483939..1484357 | 35 | 444 | 60.2 | 7.00E-12 | 34 | 97% | 0 | 0% | 1 | 35 | 100% | 137 | 171 | + | 137 |
| 28_MTB313.crt.8 | M.SpyM23NDORF616P: WARNING2, ID: 100.00%, HSP/Length: 419/444, Contig name: CP009612.1, Position: 1222659..1223077 | 35 | 444 | 60.2 | 7.00E-12 | 34 | 97% | 0 | 0% | 1 | 35 | 100% | 137 | 171 | + | 137 |
| 29_MTB314.crt.8 | M.SpyM23NDORF616P: PERFECT MATCH, ID: 100.00%, HSP/Length: 444/444, Contig name: CP008695.1, Position: 180965..181408 | 35 | 444 | 60.2 | 7.00E-12 | 34 | 97% | 0 | 0% | 1 | 35 | 100% | 137 | 171 | + | 137 |
| 29_MTB314.crt.8 | M.SpyM23NDORF616P: PERFECT MATCH, ID: 100.00%, HSP/Length: 444/444, Contig name: CP008695.1, Position: 585419..585862 | 35 | 444 | 60.2 | 7.00E-12 | 34 | 97% | 0 | 0% | 1 | 35 | 100% | 137 | 171 | + | 137 |
| 29_MTB314.crt.8 | M.SpyM23NDORF616P: WARNING2, ID: 100.00%, HSP/Length: 419/444, Contig name: CP006366.1, Position: 1483939..1484357 | 35 | 444 | 60.2 | 7.00E-12 | 34 | 97% | 0 | 0% | 1 | 35 | 100% | 137 | 171 | + | 137 |
| 29_MTB314.crt.8 | M.SpyM23NDORF616P: WARNING2, ID: 100.00%, HSP/Length: 419/444, Contig name: CP009612.1, Position: 1222659..1223077 | 35 | 444 | 60.2 | 7.00E-12 | 34 | 97% | 0 | 0% | 1 | 35 | 100% | 137 | 171 | + | 137 |
| 31_AP1.crt.1 | M.SpyM23NDORF616P: PERFECT MATCH, ID: 100.00%, HSP/Length: 444/444, Contig name: CP008695.1, Position: 180965..181408 | 35 | 444 | 60.2 | 7.00E-12 | 34 | 97% | 0 | 0% | 1 | 35 | 100% | 137 | 171 | - | 137 |
| 31_AP1.crt.1 | M.SpyM23NDORF616P: PERFECT MATCH, ID: 100.00%, HSP/Length: 444/444, Contig name: CP008695.1, Position: 585419..585862 | 35 | 444 | 60.2 | 7.00E-12 | 34 | 97% | 0 | 0% | 1 | 35 | 100% | 137 | 171 | - | 137 |
| 31_AP1.crt.1 | M.SpyM23NDORF616P: WARNING2, ID: 100.00%, HSP/Length: 419/444, Contig name: CP006366.1, Position: 1483939..1484357 | 35 | 444 | 60.2 | 7.00E-12 | 34 | 97% | 0 | 0% | 1 | 35 | 100% | 137 | 171 | - | 137 |
| 31_AP1.crt.1 | M.SpyM23NDORF616P: WARNING2, ID: 100.00%, HSP/Length: 419/444, Contig name: CP009612.1, Position: 1222659..1223077 | 35 | 444 | 60.2 | 7.00E-12 | 34 | 97% | 0 | 0% | 1 | 35 | 100% | 137 | 171 | - | 137 |
| 38_5448.crt.1 | M.SpyM23NDORF616P: PERFECT MATCH, ID: 100.00%, HSP/Length: 444/444, Contig name: CP008695.1, Position: 180965..181408 | 35 | 444 | 60.2 | 7.00E-12 | 34 | 97% | 0 | 0% | 1 | 35 | 100% | 137 | 171 | - | 137 |
| 38_5448.crt.1 | M.SpyM23NDORF616P: PERFECT MATCH, ID: 100.00%, HSP/Length: 444/444, Contig name: CP008695.1, Position: 585419..585862 | 35 | 444 | 60.2 | 7.00E-12 | 34 | 97% | 0 | 0% | 1 | 35 | 100% | 137 | 171 | - | 137 |
| 38_5448.crt.1 | M.SpyM23NDORF616P: WARNING2, ID: 100.00%, HSP/Length: 419/444, Contig name: CP006366.1, Position: 1483939..1484357 | 35 | 444 | 60.2 | 7.00E-12 | 34 | 97% | 0 | 0% | 1 | 35 | 100% | 137 | 171 | - | 137 |
| 38_5448.crt.1 | M.SpyM23NDORF616P: WARNING2, ID: 100.00%, HSP/Length: 419/444, Contig name: CP009612.1, Position: 1222659..1223077 | 35 | 444 | 60.2 | 7.00E-12 | 34 | 97% | 0 | 0% | 1 | 35 | 100% | 137 | 171 | - | 137 |
| 40_HKU488.crt.8 | M.SpyM23NDORF616P: PERFECT MATCH, ID: 100.00%, HSP/Length: 444/444, Contig name: CP008695.1, Position: 180965..181408 | 35 | 444 | 60.2 | 7.00E-12 | 34 | 97% | 0 | 0% | 1 | 35 | 100% | 137 | 171 | + | 137 |
| 40_HKU488.crt.8 | M.SpyM23NDORF616P: PERFECT MATCH, ID: 100.00%, HSP/Length: 444/444, Contig name: CP008695.1, Position: 585419..585862 | 35 | 444 | 60.2 | 7.00E-12 | 34 | 97% | 0 | 0% | 1 | 35 | 100% | 137 | 171 | + | 137 |
| 40_HKU488.crt.8 | M.SpyM23NDORF616P: WARNING2, ID: 100.00%, HSP/Length: 419/444, Contig name: CP006366.1, Position: 1483939..1484357 | 35 | 444 | 60.2 | 7.00E-12 | 34 | 97% | 0 | 0% | 1 | 35 | 100% | 137 | 171 | + | 137 |
| 40_HKU488.crt.8 | M.SpyM23NDORF616P: WARNING2, ID: 100.00%, HSP/Length: 419/444, Contig name: CP009612.1, Position: 1222659..1223077 | 35 | 444 | 60.2 | 7.00E-12 | 34 | 97% | 0 | 0% | 1 | 35 | 100% | 137 | 171 | + | 137 |
| 45_FDAARGOS_149.crt.1 | M.SpyM23NDORF616P: PERFECT MATCH, ID: 100.00%, HSP/Length: 444/444, Contig name: CP008695.1, Position: 180965..181408 | 35 | 444 | 60.2 | 7.00E-12 | 34 | 97% | 0 | 0% | 1 | 35 | 100% | 137 | 171 | - | 137 |
| 45_FDAARGOS_149.crt.1 | M.SpyM23NDORF616P: PERFECT MATCH, ID: 100.00%, HSP/Length: 444/444, Contig name: CP008695.1, Position: 585419..585862 | 35 | 444 | 60.2 | 7.00E-12 | 34 | 97% | 0 | 0% | 1 | 35 | 100% | 137 | 171 | - | 137 |
| 45_FDAARGOS_149.crt.1 | M.SpyM23NDORF616P: WARNING2, ID: 100.00%, HSP/Length: 419/444, Contig name: CP006366.1, Position: 1483939..1484357 | 35 | 444 | 60.2 | 7.00E-12 | 34 | 97% | 0 | 0% | 1 | 35 | 100% | 137 | 171 | - | 137 |
| 45_FDAARGOS_149.crt.1 | M.SpyM23NDORF616P: WARNING2, ID: 100.00%, HSP/Length: 419/444, Contig name: CP009612.1, Position: 1222659..1223077 | 35 | 444 | 60.2 | 7.00E-12 | 34 | 97% | 0 | 0% | 1 | 35 | 100% | 137 | 171 | - | 137 |
| 7_MGAS5005.crt.8 | M.SpyM23NDORF616P: PERFECT MATCH, ID: 100.00%, HSP/Length: 444/444, Contig name: CP008695.1, Position: 180965..181408 | 35 | 444 | 60.2 | 7.00E-12 | 34 | 97% | 0 | 0% | 1 | 35 | 100% | 137 | 171 | + | 137 |
| 7_MGAS5005.crt.8 | M.SpyM23NDORF616P: PERFECT MATCH, ID: 100.00%, HSP/Length: 444/444, Contig name: CP008695.1, Position: 585419..585862 | 35 | 444 | 60.2 | 7.00E-12 | 34 | 97% | 0 | 0% | 1 | 35 | 100% | 137 | 171 | + | 137 |
| 7_MGAS5005.crt.8 | M.SpyM23NDORF616P: WARNING2, ID: 100.00%, HSP/Length: 419/444, Contig name: CP006366.1, Position: 1483939..1484357 | 35 | 444 | 60.2 | 7.00E-12 | 34 | 97% | 0 | 0% | 1 | 35 | 100% | 137 | 171 | + | 137 |
| 7_MGAS5005.crt.8 | M.SpyM23NDORF616P: WARNING2, ID: 100.00%, HSP/Length: 419/444, Contig name: CP009612.1, Position: 1222659..1223077 | 35 | 444 | 60.2 | 7.00E-12 | 34 | 97% | 0 | 0% | 1 | 35 | 100% | 137 | 171 | + | 137 |
| 1_SF370.crt.10 | M.SpyHSC5ORF7630P: PERFECT MATCH, ID: 100.00%, HSP/Length: 483/483, Contig name: AE009949.1, Position: 590014..590496 | 34 | 483 | 58.4 | 2.00E-11 | 34 | 97% | 1 | 3% | 1 | 34 | 100% | 137 | 171 | + | 137 |
| 1_SF370.crt.10 | M.SpyHSC5ORF7630P: PERFECT MATCH, ID: 100.00%, HSP/Length: 483/483, Contig name: CP000260.1, Position: 543421..543903 | 34 | 483 | 58.4 | 2.00E-11 | 34 | 97% | 1 | 3% | 1 | 34 | 100% | 137 | 171 | + | 137 |
| 1_SF370.crt.10 | M.SpyHSC5ORF7630P: PERFECT MATCH, ID: 100.00%, HSP/Length: 483/483, Contig name: CP000262.1, Position: 560598..561080 | 34 | 483 | 58.4 | 2.00E-11 | 34 | 97% | 1 | 3% | 1 | 34 | 100% | 137 | 171 | + | 137 |
| 1_SF370.crt.10 | M.SpyHSC5ORF7630P: WARNING1, ID: 99.59%, HSP/Length: 483/483, Contig name: CP013838.1, Position: 540664..541146 | 34 | 483 | 58.4 | 2.00E-11 | 34 | 97% | 1 | 3% | 1 | 34 | 100% | 137 | 171 | + | 137 |
| 17_M1_476.crt.8 | M.SpyHSC5ORF7630P: PERFECT MATCH, ID: 100.00%, HSP/Length: 483/483, Contig name: AE009949.1, Position: 590014..590496 | 35 | 483 | 60.2 | 7.00E-12 | 34 | 97% | 0 | 0% | 1 | 35 | 100% | 137 | 171 | + | 137 |
| 17_M1_476.crt.8 | M.SpyHSC5ORF7630P: PERFECT MATCH, ID: 100.00%, HSP/Length: 483/483, Contig name: CP000260.1, Position: 543421..543903 | 35 | 483 | 60.2 | 7.00E-12 | 34 | 97% | 0 | 0% | 1 | 35 | 100% | 137 | 171 | + | 137 |
| 17_M1_476.crt.8 | M.SpyHSC5ORF7630P: PERFECT MATCH, ID: 100.00%, HSP/Length: 483/483, Contig name: CP000262.1, Position: 560598..561080 | 35 | 483 | 60.2 | 7.00E-12 | 34 | 97% | 0 | 0% | 1 | 35 | 100% | 137 | 171 | + | 137 |
| 17_M1_476.crt.8 | M.SpyHSC5ORF7630P: WARNING1, ID: 99.59%, HSP/Length: 483/483, Contig name: CP013838.1, Position: 540664..541146 | 35 | 483 | 60.2 | 7.00E-12 | 34 | 97% | 0 | 0% | 1 | 35 | 100% | 137 | 171 | + | 137 |
| 18_A20.crt.8 | M.SpyHSC5ORF7630P: PERFECT MATCH, ID: 100.00%, HSP/Length: 483/483, Contig name: AE009949.1, Position: 590014..590496 | 35 | 483 | 60.2 | 7.00E-12 | 34 | 97% | 0 | 0% | 1 | 35 | 100% | 137 | 171 | + | 137 |
| 18_A20.crt.8 | M.SpyHSC5ORF7630P: PERFECT MATCH, ID: 100.00%, HSP/Length: 483/483, Contig name: CP000260.1, Position: 543421..543903 | 35 | 483 | 60.2 | 7.00E-12 | 34 | 97% | 0 | 0% | 1 | 35 | 100% | 137 | 171 | + | 137 |
| 18_A20.crt.8 | M.SpyHSC5ORF7630P: PERFECT MATCH, ID: 100.00%, HSP/Length: 483/483, Contig name: CP000262.1, Position: 560598..561080 | 35 | 483 | 60.2 | 7.00E-12 | 34 | 97% | 0 | 0% | 1 | 35 | 100% | 137 | 171 | + | 137 |
| 18_A20.crt.8 | M.SpyHSC5ORF7630P: WARNING1, ID: 99.59%, HSP/Length: 483/483, Contig name: CP013838.1, Position: 540664..541146 | 35 | 483 | 60.2 | 7.00E-12 | 34 | 97% | 0 | 0% | 1 | 35 | 100% | 137 | 171 | + | 137 |
| 28_MTB313.crt.8 | M.SpyHSC5ORF7630P: PERFECT MATCH, ID: 100.00%, HSP/Length: 483/483, Contig name: AE009949.1, Position: 590014..590496 | 35 | 483 | 60.2 | 7.00E-12 | 34 | 97% | 0 | 0% | 1 | 35 | 100% | 137 | 171 | + | 137 |
| 28_MTB313.crt.8 | M.SpyHSC5ORF7630P: PERFECT MATCH, ID: 100.00%, HSP/Length: 483/483, Contig name: CP000260.1, Position: 543421..543903 | 35 | 483 | 60.2 | 7.00E-12 | 34 | 97% | 0 | 0% | 1 | 35 | 100% | 137 | 171 | + | 137 |
| 28_MTB313.crt.8 | M.SpyHSC5ORF7630P: PERFECT MATCH, ID: 100.00%, HSP/Length: 483/483, Contig name: CP000262.1, Position: 560598..561080 | 35 | 483 | 60.2 | 7.00E-12 | 34 | 97% | 0 | 0% | 1 | 35 | 100% | 137 | 171 | + | 137 |
| 28_MTB313.crt.8 | M.SpyHSC5ORF7630P: WARNING1, ID: 99.59%, HSP/Length: 483/483, Contig name: CP013838.1, Position: 540664..541146 | 35 | 483 | 60.2 | 7.00E-12 | 34 | 97% | 0 | 0% | 1 | 35 | 100% | 137 | 171 | + | 137 |
| 29_MTB314.crt.8 | M.SpyHSC5ORF7630P: PERFECT MATCH, ID: 100.00%, HSP/Length: 483/483, Contig name: AE009949.1, Position: 590014..590496 | 35 | 483 | 60.2 | 7.00E-12 | 34 | 97% | 0 | 0% | 1 | 35 | 100% | 137 | 171 | + | 137 |
| 29_MTB314.crt.8 | M.SpyHSC5ORF7630P: PERFECT MATCH, ID: 100.00%, HSP/Length: 483/483, Contig name: CP000260.1, Position: 543421..543903 | 35 | 483 | 60.2 | 7.00E-12 | 34 | 97% | 0 | 0% | 1 | 35 | 100% | 137 | 171 | + | 137 |
| 29_MTB314.crt.8 | M.SpyHSC5ORF7630P: PERFECT MATCH, ID: 100.00%, HSP/Length: 483/483, Contig name: CP000262.1, Position: 560598..561080 | 35 | 483 | 60.2 | 7.00E-12 | 34 | 97% | 0 | 0% | 1 | 35 | 100% | 137 | 171 | + | 137 |
| 29_MTB314.crt.8 | M.SpyHSC5ORF7630P: WARNING1, ID: 99.59%, HSP/Length: 483/483, Contig name: CP013838.1, Position: 540664..541146 | 35 | 483 | 60.2 | 7.00E-12 | 34 | 97% | 0 | 0% | 1 | 35 | 100% | 137 | 171 | + | 137 |
| 31_AP1.crt.1 | M.SpyHSC5ORF7630P: PERFECT MATCH, ID: 100.00%, HSP/Length: 483/483, Contig name: AE009949.1, Position: 590014..590496 | 35 | 483 | 60.2 | 7.00E-12 | 34 | 97% | 0 | 0% | 1 | 35 | 100% | 137 | 171 | - | 137 |
| 31_AP1.crt.1 | M.SpyHSC5ORF7630P: PERFECT MATCH, ID: 100.00%, HSP/Length: 483/483, Contig name: CP000260.1, Position: 543421..543903 | 35 | 483 | 60.2 | 7.00E-12 | 34 | 97% | 0 | 0% | 1 | 35 | 100% | 137 | 171 | - | 137 |
| 31_AP1.crt.1 | M.SpyHSC5ORF7630P: PERFECT MATCH, ID: 100.00%, HSP/Length: 483/483, Contig name: CP000262.1, Position: 560598..561080 | 35 | 483 | 60.2 | 7.00E-12 | 34 | 97% | 0 | 0% | 1 | 35 | 100% | 137 | 171 | - | 137 |
| 31_AP1.crt.1 | M.SpyHSC5ORF7630P: WARNING1, ID: 99.59%, HSP/Length: 483/483, Contig name: CP013838.1, Position: 540664..541146 | 35 | 483 | 60.2 | 7.00E-12 | 34 | 97% | 0 | 0% | 1 | 35 | 100% | 137 | 171 | - | 137 |
| 38_5448.crt.1 | M.SpyHSC5ORF7630P: PERFECT MATCH, ID: 100.00%, HSP/Length: 483/483, Contig name: AE009949.1, Position: 590014..590496 | 35 | 483 | 60.2 | 7.00E-12 | 34 | 97% | 0 | 0% | 1 | 35 | 100% | 137 | 171 | - | 137 |
| 38_5448.crt.1 | M.SpyHSC5ORF7630P: PERFECT MATCH, ID: 100.00%, HSP/Length: 483/483, Contig name: CP000260.1, Position: 543421..543903 | 35 | 483 | 60.2 | 7.00E-12 | 34 | 97% | 0 | 0% | 1 | 35 | 100% | 137 | 171 | - | 137 |
| 38_5448.crt.1 | M.SpyHSC5ORF7630P: PERFECT MATCH, ID: 100.00%, HSP/Length: 483/483, Contig name: CP000262.1, Position: 560598..561080 | 35 | 483 | 60.2 | 7.00E-12 | 34 | 97% | 0 | 0% | 1 | 35 | 100% | 137 | 171 | - | 137 |
| 38_5448.crt.1 | M.SpyHSC5ORF7630P: WARNING1, ID: 99.59%, HSP/Length: 483/483, Contig name: CP013838.1, Position: 540664..541146 | 35 | 483 | 60.2 | 7.00E-12 | 34 | 97% | 0 | 0% | 1 | 35 | 100% | 137 | 171 | - | 137 |
| 40_HKU488.crt.8 | M.SpyHSC5ORF7630P: PERFECT MATCH, ID: 100.00%, HSP/Length: 483/483, Contig name: AE009949.1, Position: 590014..590496 | 35 | 483 | 60.2 | 7.00E-12 | 34 | 97% | 0 | 0% | 1 | 35 | 100% | 137 | 171 | + | 137 |
| 40_HKU488.crt.8 | M.SpyHSC5ORF7630P: PERFECT MATCH, ID: 100.00%, HSP/Length: 483/483, Contig name: CP000260.1, Position: 543421..543903 | 35 | 483 | 60.2 | 7.00E-12 | 34 | 97% | 0 | 0% | 1 | 35 | 100% | 137 | 171 | + | 137 |
| 40_HKU488.crt.8 | M.SpyHSC5ORF7630P: PERFECT MATCH, ID: 100.00%, HSP/Length: 483/483, Contig name: CP000262.1, Position: 560598..561080 | 35 | 483 | 60.2 | 7.00E-12 | 34 | 97% | 0 | 0% | 1 | 35 | 100% | 137 | 171 | + | 137 |
| 40_HKU488.crt.8 | M.SpyHSC5ORF7630P: WARNING1, ID: 99.59%, HSP/Length: 483/483, Contig name: CP013838.1, Position: 540664..541146 | 35 | 483 | 60.2 | 7.00E-12 | 34 | 97% | 0 | 0% | 1 | 35 | 100% | 137 | 171 | + | 137 |
| 45_FDAARGOS_149.crt.1 | M.SpyHSC5ORF7630P: PERFECT MATCH, ID: 100.00%, HSP/Length: 483/483, Contig name: AE009949.1, Position: 590014..590496 | 35 | 483 | 60.2 | 7.00E-12 | 34 | 97% | 0 | 0% | 1 | 35 | 100% | 137 | 171 | - | 137 |
| 45_FDAARGOS_149.crt.1 | M.SpyHSC5ORF7630P: PERFECT MATCH, ID: 100.00%, HSP/Length: 483/483, Contig name: CP000260.1, Position: 543421..543903 | 35 | 483 | 60.2 | 7.00E-12 | 34 | 97% | 0 | 0% | 1 | 35 | 100% | 137 | 171 | - | 137 |
| 45_FDAARGOS_149.crt.1 | M.SpyHSC5ORF7630P: PERFECT MATCH, ID: 100.00%, HSP/Length: 483/483, Contig name: CP000262.1, Position: 560598..561080 | 35 | 483 | 60.2 | 7.00E-12 | 34 | 97% | 0 | 0% | 1 | 35 | 100% | 137 | 171 | - | 137 |
| 45_FDAARGOS_149.crt.1 | M.SpyHSC5ORF7630P: WARNING1, ID: 99.59%, HSP/Length: 483/483, Contig name: CP013838.1, Position: 540664..541146 | 35 | 483 | 60.2 | 7.00E-12 | 34 | 97% | 0 | 0% | 1 | 35 | 100% | 137 | 171 | - | 137 |
| 7_MGAS5005.crt.8 | M.SpyHSC5ORF7630P: PERFECT MATCH, ID: 100.00%, HSP/Length: 483/483, Contig name: AE009949.1, Position: 590014..590496 | 35 | 483 | 60.2 | 7.00E-12 | 34 | 97% | 0 | 0% | 1 | 35 | 100% | 137 | 171 | + | 137 |
| 7_MGAS5005.crt.8 | M.SpyHSC5ORF7630P: PERFECT MATCH, ID: 100.00%, HSP/Length: 483/483, Contig name: CP000260.1, Position: 543421..543903 | 35 | 483 | 60.2 | 7.00E-12 | 34 | 97% | 0 | 0% | 1 | 35 | 100% | 137 | 171 | + | 137 |
| 7_MGAS5005.crt.8 | M.SpyHSC5ORF7630P: PERFECT MATCH, ID: 100.00%, HSP/Length: 483/483, Contig name: CP000262.1, Position: 560598..561080 | 35 | 483 | 60.2 | 7.00E-12 | 34 | 97% | 0 | 0% | 1 | 35 | 100% | 137 | 171 | + | 137 |
| 7_MGAS5005.crt.8 | M.SpyHSC5ORF7630P: WARNING1, ID: 99.59%, HSP/Length: 483/483, Contig name: CP013838.1, Position: 540664..541146 | 35 | 483 | 60.2 | 7.00E-12 | 34 | 97% | 0 | 0% | 1 | 35 | 100% | 137 | 171 | + | 137 |
| 1_SF370.crt.10 | M.Spy743ORF6920P: PERFECT MATCH, ID: 100.00%, HSP/Length: 483/483, Contig name: A08_1, Position: 1415220..1415702 | 34 | 483 | 58.4 | 2.00E-11 | 34 | 97% | 1 | 3% | 1 | 34 | 100% | 137 | 171 | + | 137 |
| 1_SF370.crt.10 | M.Spy743ORF6920P: PERFECT MATCH, ID: 100.00%, HSP/Length: 483/483, Contig name: AM295007.1, Position: 1297556..1298038 | 34 | 483 | 58.4 | 2.00E-11 | 34 | 97% | 1 | 3% | 1 | 34 | 100% | 137 | 171 | + | 137 |
| 1_SF370.crt.10 | M.Spy743ORF6920P: PERFECT MATCH, ID: 100.00%, HSP/Length: 483/483, Contig name: CP007560.1, Position: 1282454..1282936 | 34 | 483 | 58.4 | 2.00E-11 | 34 | 97% | 1 | 3% | 1 | 34 | 100% | 137 | 171 | + | 137 |
| 1_SF370.crt.10 | M.Spy743ORF6920P: PERFECT MATCH, ID: 100.00%, HSP/Length: 483/483, Contig name: CP007560.1, Position: 725011..725493 | 34 | 483 | 58.4 | 2.00E-11 | 34 | 97% | 1 | 3% | 1 | 34 | 100% | 137 | 171 | + | 137 |
| 17_M1_476.crt.8 | M.Spy743ORF6920P: PERFECT MATCH, ID: 100.00%, HSP/Length: 483/483, Contig name: A08_1, Position: 1415220..1415702 | 35 | 483 | 60.2 | 7.00E-12 | 34 | 97% | 0 | 0% | 1 | 35 | 100% | 137 | 171 | + | 137 |
| 17_M1_476.crt.8 | M.Spy743ORF6920P: PERFECT MATCH, ID: 100.00%, HSP/Length: 483/483, Contig name: AM295007.1, Position: 1297556..1298038 | 35 | 483 | 60.2 | 7.00E-12 | 34 | 97% | 0 | 0% | 1 | 35 | 100% | 137 | 171 | + | 137 |
| 17_M1_476.crt.8 | M.Spy743ORF6920P: PERFECT MATCH, ID: 100.00%, HSP/Length: 483/483, Contig name: CP007560.1, Position: 1282454..1282936 | 35 | 483 | 60.2 | 7.00E-12 | 34 | 97% | 0 | 0% | 1 | 35 | 100% | 137 | 171 | + | 137 |
| 17_M1_476.crt.8 | M.Spy743ORF6920P: PERFECT MATCH, ID: 100.00%, HSP/Length: 483/483, Contig name: CP007560.1, Position: 725011..725493 | 35 | 483 | 60.2 | 7.00E-12 | 34 | 97% | 0 | 0% | 1 | 35 | 100% | 137 | 171 | + | 137 |
| 18_A20.crt.8 | M.Spy743ORF6920P: PERFECT MATCH, ID: 100.00%, HSP/Length: 483/483, Contig name: A08_1, Position: 1415220..1415702 | 35 | 483 | 60.2 | 7.00E-12 | 34 | 97% | 0 | 0% | 1 | 35 | 100% | 137 | 171 | + | 137 |
| 18_A20.crt.8 | M.Spy743ORF6920P: PERFECT MATCH, ID: 100.00%, HSP/Length: 483/483, Contig name: AM295007.1, Position: 1297556..1298038 | 35 | 483 | 60.2 | 7.00E-12 | 34 | 97% | 0 | 0% | 1 | 35 | 100% | 137 | 171 | + | 137 |
| 18_A20.crt.8 | M.Spy743ORF6920P: PERFECT MATCH, ID: 100.00%, HSP/Length: 483/483, Contig name: CP007560.1, Position: 1282454..1282936 | 35 | 483 | 60.2 | 7.00E-12 | 34 | 97% | 0 | 0% | 1 | 35 | 100% | 137 | 171 | + | 137 |
| 18_A20.crt.8 | M.Spy743ORF6920P: PERFECT MATCH, ID: 100.00%, HSP/Length: 483/483, Contig name: CP007560.1, Position: 725011..725493 | 35 | 483 | 60.2 | 7.00E-12 | 34 | 97% | 0 | 0% | 1 | 35 | 100% | 137 | 171 | + | 137 |
| 28_MTB313.crt.8 | M.Spy743ORF6920P: PERFECT MATCH, ID: 100.00%, HSP/Length: 483/483, Contig name: A08_1, Position: 1415220..1415702 | 35 | 483 | 60.2 | 7.00E-12 | 34 | 97% | 0 | 0% | 1 | 35 | 100% | 137 | 171 | + | 137 |
| 28_MTB313.crt.8 | M.Spy743ORF6920P: PERFECT MATCH, ID: 100.00%, HSP/Length: 483/483, Contig name: AM295007.1, Position: 1297556..1298038 | 35 | 483 | 60.2 | 7.00E-12 | 34 | 97% | 0 | 0% | 1 | 35 | 100% | 137 | 171 | + | 137 |
| 28_MTB313.crt.8 | M.Spy743ORF6920P: PERFECT MATCH, ID: 100.00%, HSP/Length: 483/483, Contig name: CP007560.1, Position: 1282454..1282936 | 35 | 483 | 60.2 | 7.00E-12 | 34 | 97% | 0 | 0% | 1 | 35 | 100% | 137 | 171 | + | 137 |
| 28_MTB313.crt.8 | M.Spy743ORF6920P: PERFECT MATCH, ID: 100.00%, HSP/Length: 483/483, Contig name: CP007560.1, Position: 725011..725493 | 35 | 483 | 60.2 | 7.00E-12 | 34 | 97% | 0 | 0% | 1 | 35 | 100% | 137 | 171 | + | 137 |
| 29_MTB314.crt.8 | M.Spy743ORF6920P: PERFECT MATCH, ID: 100.00%, HSP/Length: 483/483, Contig name: A08_1, Position: 1415220..1415702 | 35 | 483 | 60.2 | 7.00E-12 | 34 | 97% | 0 | 0% | 1 | 35 | 100% | 137 | 171 | + | 137 |
| 29_MTB314.crt.8 | M.Spy743ORF6920P: PERFECT MATCH, ID: 100.00%, HSP/Length: 483/483, Contig name: AM295007.1, Position: 1297556..1298038 | 35 | 483 | 60.2 | 7.00E-12 | 34 | 97% | 0 | 0% | 1 | 35 | 100% | 137 | 171 | + | 137 |
| 29_MTB314.crt.8 | M.Spy743ORF6920P: PERFECT MATCH, ID: 100.00%, HSP/Length: 483/483, Contig name: CP007560.1, Position: 1282454..1282936 | 35 | 483 | 60.2 | 7.00E-12 | 34 | 97% | 0 | 0% | 1 | 35 | 100% | 137 | 171 | + | 137 |
| 29_MTB314.crt.8 | M.Spy743ORF6920P: PERFECT MATCH, ID: 100.00%, HSP/Length: 483/483, Contig name: CP007560.1, Position: 725011..725493 | 35 | 483 | 60.2 | 7.00E-12 | 34 | 97% | 0 | 0% | 1 | 35 | 100% | 137 | 171 | + | 137 |
| 31_AP1.crt.1 | M.Spy743ORF6920P: PERFECT MATCH, ID: 100.00%, HSP/Length: 483/483, Contig name: A08_1, Position: 1415220..1415702 | 35 | 483 | 60.2 | 7.00E-12 | 34 | 97% | 0 | 0% | 1 | 35 | 100% | 137 | 171 | - | 137 |
| 31_AP1.crt.1 | M.Spy743ORF6920P: PERFECT MATCH, ID: 100.00%, HSP/Length: 483/483, Contig name: AM295007.1, Position: 1297556..1298038 | 35 | 483 | 60.2 | 7.00E-12 | 34 | 97% | 0 | 0% | 1 | 35 | 100% | 137 | 171 | - | 137 |
| 31_AP1.crt.1 | M.Spy743ORF6920P: PERFECT MATCH, ID: 100.00%, HSP/Length: 483/483, Contig name: CP007560.1, Position: 1282454..1282936 | 35 | 483 | 60.2 | 7.00E-12 | 34 | 97% | 0 | 0% | 1 | 35 | 100% | 137 | 171 | - | 137 |
| 31_AP1.crt.1 | M.Spy743ORF6920P: PERFECT MATCH, ID: 100.00%, HSP/Length: 483/483, Contig name: CP007560.1, Position: 725011..725493 | 35 | 483 | 60.2 | 7.00E-12 | 34 | 97% | 0 | 0% | 1 | 35 | 100% | 137 | 171 | - | 137 |
| 38_5448.crt.1 | M.Spy743ORF6920P: PERFECT MATCH, ID: 100.00%, HSP/Length: 483/483, Contig name: A08_1, Position: 1415220..1415702 | 35 | 483 | 60.2 | 7.00E-12 | 34 | 97% | 0 | 0% | 1 | 35 | 100% | 137 | 171 | - | 137 |
| 38_5448.crt.1 | M.Spy743ORF6920P: PERFECT MATCH, ID: 100.00%, HSP/Length: 483/483, Contig name: AM295007.1, Position: 1297556..1298038 | 35 | 483 | 60.2 | 7.00E-12 | 34 | 97% | 0 | 0% | 1 | 35 | 100% | 137 | 171 | - | 137 |
| 38_5448.crt.1 | M.Spy743ORF6920P: PERFECT MATCH, ID: 100.00%, HSP/Length: 483/483, Contig name: CP007560.1, Position: 1282454..1282936 | 35 | 483 | 60.2 | 7.00E-12 | 34 | 97% | 0 | 0% | 1 | 35 | 100% | 137 | 171 | - | 137 |
| 38_5448.crt.1 | M.Spy743ORF6920P: PERFECT MATCH, ID: 100.00%, HSP/Length: 483/483, Contig name: CP007560.1, Position: 725011..725493 | 35 | 483 | 60.2 | 7.00E-12 | 34 | 97% | 0 | 0% | 1 | 35 | 100% | 137 | 171 | - | 137 |
| 40_HKU488.crt.8 | M.Spy743ORF6920P: PERFECT MATCH, ID: 100.00%, HSP/Length: 483/483, Contig name: A08_1, Position: 1415220..1415702 | 35 | 483 | 60.2 | 7.00E-12 | 34 | 97% | 0 | 0% | 1 | 35 | 100% | 137 | 171 | + | 137 |
| 40_HKU488.crt.8 | M.Spy743ORF6920P: PERFECT MATCH, ID: 100.00%, HSP/Length: 483/483, Contig name: AM295007.1, Position: 1297556..1298038 | 35 | 483 | 60.2 | 7.00E-12 | 34 | 97% | 0 | 0% | 1 | 35 | 100% | 137 | 171 | + | 137 |
| 40_HKU488.crt.8 | M.Spy743ORF6920P: PERFECT MATCH, ID: 100.00%, HSP/Length: 483/483, Contig name: CP007560.1, Position: 1282454..1282936 | 35 | 483 | 60.2 | 7.00E-12 | 34 | 97% | 0 | 0% | 1 | 35 | 100% | 137 | 171 | + | 137 |
| 40_HKU488.crt.8 | M.Spy743ORF6920P: PERFECT MATCH, ID: 100.00%, HSP/Length: 483/483, Contig name: CP007560.1, Position: 725011..725493 | 35 | 483 | 60.2 | 7.00E-12 | 34 | 97% | 0 | 0% | 1 | 35 | 100% | 137 | 171 | + | 137 |
| 45_FDAARGOS_149.crt.1 | M.Spy743ORF6920P: PERFECT MATCH, ID: 100.00%, HSP/Length: 483/483, Contig name: A08_1, Position: 1415220..1415702 | 35 | 483 | 60.2 | 7.00E-12 | 34 | 97% | 0 | 0% | 1 | 35 | 100% | 137 | 171 | - | 137 |
| 45_FDAARGOS_149.crt.1 | M.Spy743ORF6920P: PERFECT MATCH, ID: 100.00%, HSP/Length: 483/483, Contig name: AM295007.1, Position: 1297556..1298038 | 35 | 483 | 60.2 | 7.00E-12 | 34 | 97% | 0 | 0% | 1 | 35 | 100% | 137 | 171 | - | 137 |
| 45_FDAARGOS_149.crt.1 | M.Spy743ORF6920P: PERFECT MATCH, ID: 100.00%, HSP/Length: 483/483, Contig name: CP007560.1, Position: 1282454..1282936 | 35 | 483 | 60.2 | 7.00E-12 | 34 | 97% | 0 | 0% | 1 | 35 | 100% | 137 | 171 | - | 137 |
| 45_FDAARGOS_149.crt.1 | M.Spy743ORF6920P: PERFECT MATCH, ID: 100.00%, HSP/Length: 483/483, Contig name: CP007560.1, Position: 725011..725493 | 35 | 483 | 60.2 | 7.00E-12 | 34 | 97% | 0 | 0% | 1 | 35 | 100% | 137 | 171 | - | 137 |
| 7_MGAS5005.crt.8 | M.Spy743ORF6920P: PERFECT MATCH, ID: 100.00%, HSP/Length: 483/483, Contig name: A08_1, Position: 1415220..1415702 | 35 | 483 | 60.2 | 7.00E-12 | 34 | 97% | 0 | 0% | 1 | 35 | 100% | 137 | 171 | + | 137 |
| 7_MGAS5005.crt.8 | M.Spy743ORF6920P: PERFECT MATCH, ID: 100.00%, HSP/Length: 483/483, Contig name: AM295007.1, Position: 1297556..1298038 | 35 | 483 | 60.2 | 7.00E-12 | 34 | 97% | 0 | 0% | 1 | 35 | 100% | 137 | 171 | + | 137 |
| 7_MGAS5005.crt.8 | M.Spy743ORF6920P: PERFECT MATCH, ID: 100.00%, HSP/Length: 483/483, Contig name: CP007560.1, Position: 1282454..1282936 | 35 | 483 | 60.2 | 7.00E-12 | 34 | 97% | 0 | 0% | 1 | 35 | 100% | 137 | 171 | + | 137 |
| 7_MGAS5005.crt.8 | M.Spy743ORF6920P: PERFECT MATCH, ID: 100.00%, HSP/Length: 483/483, Contig name: CP007560.1, Position: 725011..725493 | 35 | 483 | 60.2 | 7.00E-12 | 34 | 97% | 0 | 0% | 1 | 35 | 100% | 137 | 171 | + | 137 |
| 1_SF370.crt.3 | M2.SpySP1ORFDP: PERFECT MATCH, ID: 100.00%, HSP/Length: 789/789, Contig name: CP006366.1, Position: 1483121..1483909 | 30 | 789 | 56.5 | 7.00E-11 | 30 | 100% | 0 | 0% | 1 | 30 | 100% | 97 | 126 | - | 97 |
| 1_SF370.crt.3 | M2.SpySP1ORFDP: PERFECT MATCH, ID: 100.00%, HSP/Length: 789/789, Contig name: CP009612.1, Position: 1221841..1222629 | 30 | 789 | 56.5 | 7.00E-11 | 30 | 100% | 0 | 0% | 1 | 30 | 100% | 97 | 126 | - | 97 |
| 1_SF370.crt.3 | M2.SpySP1ORFDP: WARNING1, ID: 99.75%, HSP/Length: 789/789, Contig name: A06_1, Position: 340718..341506 | 30 | 789 | 56.5 | 7.00E-11 | 30 | 100% | 0 | 0% | 1 | 30 | 100% | 97 | 126 | - | 97 |
| 1_SF370.crt.3 | M2.SpySP1ORFDP: WARNING1, ID: 99.75%, HSP/Length: 789/789, Contig name: A09_1, Position: 379858..380646 | 30 | 789 | 56.5 | 7.00E-11 | 30 | 100% | 0 | 0% | 1 | 30 | 100% | 97 | 126 | - | 97 |
| 17_M1_476.crt.2 | M2.SpySP1ORFDP: PERFECT MATCH, ID: 100.00%, HSP/Length: 789/789, Contig name: CP006366.1, Position: 1483121..1483909 | 30 | 789 | 56.5 | 7.00E-11 | 30 | 100% | 0 | 0% | 1 | 30 | 100% | 97 | 126 | - | 97 |
| 17_M1_476.crt.2 | M2.SpySP1ORFDP: PERFECT MATCH, ID: 100.00%, HSP/Length: 789/789, Contig name: CP009612.1, Position: 1221841..1222629 | 30 | 789 | 56.5 | 7.00E-11 | 30 | 100% | 0 | 0% | 1 | 30 | 100% | 97 | 126 | - | 97 |
| 17_M1_476.crt.2 | M2.SpySP1ORFDP: WARNING1, ID: 99.75%, HSP/Length: 789/789, Contig name: A06_1, Position: 340718..341506 | 30 | 789 | 56.5 | 7.00E-11 | 30 | 100% | 0 | 0% | 1 | 30 | 100% | 97 | 126 | - | 97 |
| 17_M1_476.crt.2 | M2.SpySP1ORFDP: WARNING1, ID: 99.75%, HSP/Length: 789/789, Contig name: A09_1, Position: 379858..380646 | 30 | 789 | 56.5 | 7.00E-11 | 30 | 100% | 0 | 0% | 1 | 30 | 100% | 97 | 126 | - | 97 |
| 18_A20.crt.2 | M2.SpySP1ORFDP: PERFECT MATCH, ID: 100.00%, HSP/Length: 789/789, Contig name: CP006366.1, Position: 1483121..1483909 | 30 | 789 | 56.5 | 7.00E-11 | 30 | 100% | 0 | 0% | 1 | 30 | 100% | 97 | 126 | - | 97 |
| 18_A20.crt.2 | M2.SpySP1ORFDP: PERFECT MATCH, ID: 100.00%, HSP/Length: 789/789, Contig name: CP009612.1, Position: 1221841..1222629 | 30 | 789 | 56.5 | 7.00E-11 | 30 | 100% | 0 | 0% | 1 | 30 | 100% | 97 | 126 | - | 97 |
| 18_A20.crt.2 | M2.SpySP1ORFDP: WARNING1, ID: 99.75%, HSP/Length: 789/789, Contig name: A06_1, Position: 340718..341506 | 30 | 789 | 56.5 | 7.00E-11 | 30 | 100% | 0 | 0% | 1 | 30 | 100% | 97 | 126 | - | 97 |
| 18_A20.crt.2 | M2.SpySP1ORFDP: WARNING1, ID: 99.75%, HSP/Length: 789/789, Contig name: A09_1, Position: 379858..380646 | 30 | 789 | 56.5 | 7.00E-11 | 30 | 100% | 0 | 0% | 1 | 30 | 100% | 97 | 126 | - | 97 |
| 29_MTB314.crt.2 | M2.SpySP1ORFDP: PERFECT MATCH, ID: 100.00%, HSP/Length: 789/789, Contig name: CP006366.1, Position: 1483121..1483909 | 30 | 789 | 56.5 | 7.00E-11 | 30 | 100% | 0 | 0% | 1 | 30 | 100% | 97 | 126 | - | 97 |
| 29_MTB314.crt.2 | M2.SpySP1ORFDP: PERFECT MATCH, ID: 100.00%, HSP/Length: 789/789, Contig name: CP009612.1, Position: 1221841..1222629 | 30 | 789 | 56.5 | 7.00E-11 | 30 | 100% | 0 | 0% | 1 | 30 | 100% | 97 | 126 | - | 97 |
| 29_MTB314.crt.2 | M2.SpySP1ORFDP: WARNING1, ID: 99.75%, HSP/Length: 789/789, Contig name: A06_1, Position: 340718..341506 | 30 | 789 | 56.5 | 7.00E-11 | 30 | 100% | 0 | 0% | 1 | 30 | 100% | 97 | 126 | - | 97 |
| 29_MTB314.crt.2 | M2.SpySP1ORFDP: WARNING1, ID: 99.75%, HSP/Length: 789/789, Contig name: A09_1, Position: 379858..380646 | 30 | 789 | 56.5 | 7.00E-11 | 30 | 100% | 0 | 0% | 1 | 30 | 100% | 97 | 126 | - | 97 |
| 38_5448.crt.7 | M2.SpySP1ORFDP: PERFECT MATCH, ID: 100.00%, HSP/Length: 789/789, Contig name: CP006366.1, Position: 1483121..1483909 | 30 | 789 | 56.5 | 7.00E-11 | 30 | 100% | 0 | 0% | 1 | 30 | 100% | 97 | 126 | + | 97 |
| 38_5448.crt.7 | M2.SpySP1ORFDP: PERFECT MATCH, ID: 100.00%, HSP/Length: 789/789, Contig name: CP009612.1, Position: 1221841..1222629 | 30 | 789 | 56.5 | 7.00E-11 | 30 | 100% | 0 | 0% | 1 | 30 | 100% | 97 | 126 | + | 97 |
| 38_5448.crt.7 | M2.SpySP1ORFDP: WARNING1, ID: 99.75%, HSP/Length: 789/789, Contig name: A06_1, Position: 340718..341506 | 30 | 789 | 56.5 | 7.00E-11 | 30 | 100% | 0 | 0% | 1 | 30 | 100% | 97 | 126 | + | 97 |
| 38_5448.crt.7 | M2.SpySP1ORFDP: WARNING1, ID: 99.75%, HSP/Length: 789/789, Contig name: A09_1, Position: 379858..380646 | 30 | 789 | 56.5 | 7.00E-11 | 30 | 100% | 0 | 0% | 1 | 30 | 100% | 97 | 126 | + | 97 |
| 40_HKU488.crt.2 | M2.SpySP1ORFDP: PERFECT MATCH, ID: 100.00%, HSP/Length: 789/789, Contig name: CP006366.1, Position: 1483121..1483909 | 30 | 789 | 56.5 | 7.00E-11 | 30 | 100% | 0 | 0% | 1 | 30 | 100% | 97 | 126 | - | 97 |
| 40_HKU488.crt.2 | M2.SpySP1ORFDP: PERFECT MATCH, ID: 100.00%, HSP/Length: 789/789, Contig name: CP009612.1, Position: 1221841..1222629 | 30 | 789 | 56.5 | 7.00E-11 | 30 | 100% | 0 | 0% | 1 | 30 | 100% | 97 | 126 | - | 97 |
| 40_HKU488.crt.2 | M2.SpySP1ORFDP: WARNING1, ID: 99.75%, HSP/Length: 789/789, Contig name: A06_1, Position: 340718..341506 | 30 | 789 | 56.5 | 7.00E-11 | 30 | 100% | 0 | 0% | 1 | 30 | 100% | 97 | 126 | - | 97 |
| 40_HKU488.crt.2 | M2.SpySP1ORFDP: WARNING1, ID: 99.75%, HSP/Length: 789/789, Contig name: A09_1, Position: 379858..380646 | 30 | 789 | 56.5 | 7.00E-11 | 30 | 100% | 0 | 0% | 1 | 30 | 100% | 97 | 126 | - | 97 |
| 45_FDAARGOS_149.crt.7 | M2.SpySP1ORFDP: PERFECT MATCH, ID: 100.00%, HSP/Length: 789/789, Contig name: CP006366.1, Position: 1483121..1483909 | 30 | 789 | 56.5 | 7.00E-11 | 30 | 100% | 0 | 0% | 1 | 30 | 100% | 97 | 126 | + | 97 |
| 45_FDAARGOS_149.crt.7 | M2.SpySP1ORFDP: PERFECT MATCH, ID: 100.00%, HSP/Length: 789/789, Contig name: CP009612.1, Position: 1221841..1222629 | 30 | 789 | 56.5 | 7.00E-11 | 30 | 100% | 0 | 0% | 1 | 30 | 100% | 97 | 126 | + | 97 |
| 45_FDAARGOS_149.crt.7 | M2.SpySP1ORFDP: WARNING1, ID: 99.75%, HSP/Length: 789/789, Contig name: A06_1, Position: 340718..341506 | 30 | 789 | 56.5 | 7.00E-11 | 30 | 100% | 0 | 0% | 1 | 30 | 100% | 97 | 126 | + | 97 |
| 45_FDAARGOS_149.crt.7 | M2.SpySP1ORFDP: WARNING1, ID: 99.75%, HSP/Length: 789/789, Contig name: A09_1, Position: 379858..380646 | 30 | 789 | 56.5 | 7.00E-11 | 30 | 100% | 0 | 0% | 1 | 30 | 100% | 97 | 126 | + | 97 |
| 7_MGAS5005.crt.2 | M2.SpySP1ORFDP: PERFECT MATCH, ID: 100.00%, HSP/Length: 789/789, Contig name: CP006366.1, Position: 1483121..1483909 | 30 | 789 | 56.5 | 7.00E-11 | 30 | 100% | 0 | 0% | 1 | 30 | 100% | 97 | 126 | - | 97 |
| 7_MGAS5005.crt.2 | M2.SpySP1ORFDP: PERFECT MATCH, ID: 100.00%, HSP/Length: 789/789, Contig name: CP009612.1, Position: 1221841..1222629 | 30 | 789 | 56.5 | 7.00E-11 | 30 | 100% | 0 | 0% | 1 | 30 | 100% | 97 | 126 | - | 97 |
| 7_MGAS5005.crt.2 | M2.SpySP1ORFDP: WARNING1, ID: 99.75%, HSP/Length: 789/789, Contig name: A06_1, Position: 340718..341506 | 30 | 789 | 56.5 | 7.00E-11 | 30 | 100% | 0 | 0% | 1 | 30 | 100% | 97 | 126 | - | 97 |
| 7_MGAS5005.crt.2 | M2.SpySP1ORFDP: WARNING1, ID: 99.75%, HSP/Length: 789/789, Contig name: A09_1, Position: 379858..380646 | 30 | 789 | 56.5 | 7.00E-11 | 30 | 100% | 0 | 0% | 1 | 30 | 100% | 97 | 126 | - | 97 |
| 1_SF370.crt.3 | M.SpyHSC5ORF7625P: PERFECT MATCH, ID: 100.00%, HSP/Length: 750/750, Contig name: AE009949.1, Position: 590501..591250 | 30 | 750 | 56.5 | 7.00E-11 | 30 | 100% | 0 | 0% | 1 | 30 | 100% | 58 | 87 | - | 58 |
| 1_SF370.crt.3 | M.SpyHSC5ORF7625P: PERFECT MATCH, ID: 100.00%, HSP/Length: 750/750, Contig name: CP000262.1, Position: 561085..561834 | 30 | 750 | 56.5 | 7.00E-11 | 30 | 100% | 0 | 0% | 1 | 30 | 100% | 58 | 87 | - | 58 |
| 1_SF370.crt.3 | M.SpyHSC5ORF7625P: WARNING1, ID: 99.87%, HSP/Length: 750/750, Contig name: CP000260.1, Position: 543908..544657 | 30 | 750 | 56.5 | 7.00E-11 | 30 | 100% | 0 | 0% | 1 | 30 | 100% | 58 | 87 | - | 58 |
| 17_M1_476.crt.2 | M.SpyHSC5ORF7625P: PERFECT MATCH, ID: 100.00%, HSP/Length: 750/750, Contig name: AE009949.1, Position: 590501..591250 | 30 | 750 | 56.5 | 7.00E-11 | 30 | 100% | 0 | 0% | 1 | 30 | 100% | 58 | 87 | - | 58 |
| 17_M1_476.crt.2 | M.SpyHSC5ORF7625P: PERFECT MATCH, ID: 100.00%, HSP/Length: 750/750, Contig name: CP000262.1, Position: 561085..561834 | 30 | 750 | 56.5 | 7.00E-11 | 30 | 100% | 0 | 0% | 1 | 30 | 100% | 58 | 87 | - | 58 |
| 17_M1_476.crt.2 | M.SpyHSC5ORF7625P: WARNING1, ID: 99.87%, HSP/Length: 750/750, Contig name: CP000260.1, Position: 543908..544657 | 30 | 750 | 56.5 | 7.00E-11 | 30 | 100% | 0 | 0% | 1 | 30 | 100% | 58 | 87 | - | 58 |
| 18_A20.crt.2 | M.SpyHSC5ORF7625P: PERFECT MATCH, ID: 100.00%, HSP/Length: 750/750, Contig name: AE009949.1, Position: 590501..591250 | 30 | 750 | 56.5 | 7.00E-11 | 30 | 100% | 0 | 0% | 1 | 30 | 100% | 58 | 87 | - | 58 |
| 18_A20.crt.2 | M.SpyHSC5ORF7625P: PERFECT MATCH, ID: 100.00%, HSP/Length: 750/750, Contig name: CP000262.1, Position: 561085..561834 | 30 | 750 | 56.5 | 7.00E-11 | 30 | 100% | 0 | 0% | 1 | 30 | 100% | 58 | 87 | - | 58 |
| 18_A20.crt.2 | M.SpyHSC5ORF7625P: WARNING1, ID: 99.87%, HSP/Length: 750/750, Contig name: CP000260.1, Position: 543908..544657 | 30 | 750 | 56.5 | 7.00E-11 | 30 | 100% | 0 | 0% | 1 | 30 | 100% | 58 | 87 | - | 58 |
| 29_MTB314.crt.2 | M.SpyHSC5ORF7625P: PERFECT MATCH, ID: 100.00%, HSP/Length: 750/750, Contig name: AE009949.1, Position: 590501..591250 | 30 | 750 | 56.5 | 7.00E-11 | 30 | 100% | 0 | 0% | 1 | 30 | 100% | 58 | 87 | - | 58 |
| 29_MTB314.crt.2 | M.SpyHSC5ORF7625P: PERFECT MATCH, ID: 100.00%, HSP/Length: 750/750, Contig name: CP000262.1, Position: 561085..561834 | 30 | 750 | 56.5 | 7.00E-11 | 30 | 100% | 0 | 0% | 1 | 30 | 100% | 58 | 87 | - | 58 |
| 29_MTB314.crt.2 | M.SpyHSC5ORF7625P: WARNING1, ID: 99.87%, HSP/Length: 750/750, Contig name: CP000260.1, Position: 543908..544657 | 30 | 750 | 56.5 | 7.00E-11 | 30 | 100% | 0 | 0% | 1 | 30 | 100% | 58 | 87 | - | 58 |
| 38_5448.crt.7 | M.SpyHSC5ORF7625P: PERFECT MATCH, ID: 100.00%, HSP/Length: 750/750, Contig name: AE009949.1, Position: 590501..591250 | 30 | 750 | 56.5 | 7.00E-11 | 30 | 100% | 0 | 0% | 1 | 30 | 100% | 58 | 87 | + | 58 |
| 38_5448.crt.7 | M.SpyHSC5ORF7625P: PERFECT MATCH, ID: 100.00%, HSP/Length: 750/750, Contig name: CP000262.1, Position: 561085..561834 | 30 | 750 | 56.5 | 7.00E-11 | 30 | 100% | 0 | 0% | 1 | 30 | 100% | 58 | 87 | + | 58 |
| 38_5448.crt.7 | M.SpyHSC5ORF7625P: WARNING1, ID: 99.87%, HSP/Length: 750/750, Contig name: CP000260.1, Position: 543908..544657 | 30 | 750 | 56.5 | 7.00E-11 | 30 | 100% | 0 | 0% | 1 | 30 | 100% | 58 | 87 | + | 58 |
| 40_HKU488.crt.2 | M.SpyHSC5ORF7625P: PERFECT MATCH, ID: 100.00%, HSP/Length: 750/750, Contig name: AE009949.1, Position: 590501..591250 | 30 | 750 | 56.5 | 7.00E-11 | 30 | 100% | 0 | 0% | 1 | 30 | 100% | 58 | 87 | - | 58 |
| 40_HKU488.crt.2 | M.SpyHSC5ORF7625P: PERFECT MATCH, ID: 100.00%, HSP/Length: 750/750, Contig name: CP000262.1, Position: 561085..561834 | 30 | 750 | 56.5 | 7.00E-11 | 30 | 100% | 0 | 0% | 1 | 30 | 100% | 58 | 87 | - | 58 |
| 40_HKU488.crt.2 | M.SpyHSC5ORF7625P: WARNING1, ID: 99.87%, HSP/Length: 750/750, Contig name: CP000260.1, Position: 543908..544657 | 30 | 750 | 56.5 | 7.00E-11 | 30 | 100% | 0 | 0% | 1 | 30 | 100% | 58 | 87 | - | 58 |
| 45_FDAARGOS_149.crt.7 | M.SpyHSC5ORF7625P: PERFECT MATCH, ID: 100.00%, HSP/Length: 750/750, Contig name: AE009949.1, Position: 590501..591250 | 30 | 750 | 56.5 | 7.00E-11 | 30 | 100% | 0 | 0% | 1 | 30 | 100% | 58 | 87 | + | 58 |
| 45_FDAARGOS_149.crt.7 | M.SpyHSC5ORF7625P: PERFECT MATCH, ID: 100.00%, HSP/Length: 750/750, Contig name: CP000262.1, Position: 561085..561834 | 30 | 750 | 56.5 | 7.00E-11 | 30 | 100% | 0 | 0% | 1 | 30 | 100% | 58 | 87 | + | 58 |
| 45_FDAARGOS_149.crt.7 | M.SpyHSC5ORF7625P: WARNING1, ID: 99.87%, HSP/Length: 750/750, Contig name: CP000260.1, Position: 543908..544657 | 30 | 750 | 56.5 | 7.00E-11 | 30 | 100% | 0 | 0% | 1 | 30 | 100% | 58 | 87 | + | 58 |
| 7_MGAS5005.crt.2 | M.SpyHSC5ORF7625P: PERFECT MATCH, ID: 100.00%, HSP/Length: 750/750, Contig name: AE009949.1, Position: 590501..591250 | 30 | 750 | 56.5 | 7.00E-11 | 30 | 100% | 0 | 0% | 1 | 30 | 100% | 58 | 87 | - | 58 |
| 7_MGAS5005.crt.2 | M.SpyHSC5ORF7625P: PERFECT MATCH, ID: 100.00%, HSP/Length: 750/750, Contig name: CP000262.1, Position: 561085..561834 | 30 | 750 | 56.5 | 7.00E-11 | 30 | 100% | 0 | 0% | 1 | 30 | 100% | 58 | 87 | - | 58 |
| 7_MGAS5005.crt.2 | M.SpyHSC5ORF7625P: WARNING1, ID: 99.87%, HSP/Length: 750/750, Contig name: CP000260.1, Position: 543908..544657 | 30 | 750 | 56.5 | 7.00E-11 | 30 | 100% | 0 | 0% | 1 | 30 | 100% | 58 | 87 | - | 58 |

Supplementary Table S6. BLASTN outputs of spacer sequences against prophage region sequences in 59 *S. pyogenes* strains

| query_name | subject_name | query_length | subject_length | score | E-value | identity | identity_per | gap | gap_per | query_start | query_end | coverage | subject_start | subject_end | subject_strand | subject_index |
|---|---|---|---|---|---|---|---|---|---|---|---|---|---|---|---|---|
| 10_MGAS2096.crt.1 | NOHIT | 38 | | | | | | | | | | | | | | |
| 10_MGAS2096.crt.2 | NOHIT | 37 | | | | | | | | | | | | | | |
| 10_MGAS2096.crt.3 | NOHIT | 0 | | | | | | | | | | | | | | |
| 10_MGAS2096.crt.4 | NOHIT | 33 | | | | | | | | | | | | | | |
| 10_MGAS2096.crt.5 | CP000056.1 Streptococcus pyogenes MGAS6180, complete genome | 35 | 35292 | 65.8 | 1.00E-12 | 35 | 100% | 0 | 0% | 1 | 35 | 100% | 29394 | 29428 | - | 29394 |
| 10_MGAS2096.crt.5 | A09_1 | 35 | 30199 | 65.8 | 1.00E-12 | 35 | 100% | 0 | 0% | 1 | 35 | 100% | 8103 | 8137 | + | 8103 |
| 10_MGAS2096.crt.5 | A07_1 | 35 | 37110 | 65.8 | 1.00E-12 | 35 | 100% | 0 | 0% | 1 | 35 | 100% | 23710 | 23744 | - | 23710 |
| 10_MGAS2096.crt.5 | A06_1 | 35 | 30200 | 65.8 | 1.00E-12 | 35 | 100% | 0 | 0% | 1 | 35 | 100% | 8103 | 8137 | + | 8103 |
| 10_MGAS2096.crt.5 | CP011069.1 Streptococcus pyogenes strain STAB09014, complete genome | 35 | 35292 | 65.8 | 1.00E-12 | 35 | 100% | 0 | 0% | 1 | 35 | 100% | 29394 | 29428 | - | 29394 |
| 10_MGAS2096.crt.5 | BA000034.2 Streptococcus pyogenes SSI-1 DNA, complete genome | 35 | 37117 | 65.8 | 1.00E-12 | 35 | 100% | 0 | 0% | 1 | 35 | 100% | 13367 | 13401 | + | 13367 |
| 10_MGAS2096.crt.5 | CP013672.1 Streptococcus pyogenes strain AP53, complete genome | 35 | 29027 | 65.8 | 1.00E-12 | 35 | 100% | 0 | 0% | 1 | 35 | 100% | 21483 | 21517 | - | 21483 |
| 10_MGAS2096.crt.5 | CP014139.1 Streptococcus pyogenes strain MEW123, complete genome | 35 | 34850 | 65.8 | 1.00E-12 | 35 | 100% | 0 | 0% | 1 | 35 | 100% | 15233 | 15267 | + | 15233 |
| 10_MGAS2096.crt.5 | CP010450.1 Streptococcus pyogenes strain NGAS638, complete genome | 35 | 35488 | 65.8 | 1.00E-12 | 35 | 100% | 0 | 0% | 1 | 35 | 100% | 29070 | 29104 | - | 29070 |
| 10_MGAS2096.crt.5 | AE014074.1 Streptococcus pyogenes MGAS315, complete genome | 35 | 36875 | 65.8 | 1.00E-12 | 35 | 100% | 0 | 0% | 1 | 35 | 100% | 23486 | 23520 | - | 23486 |
| 10_MGAS2096.crt.5 | CP011068.1 Streptococcus pyogenes strain STAB10015, complete genome | 35 | 35292 | 65.8 | 1.00E-12 | 35 | 100% | 0 | 0% | 1 | 35 | 100% | 29394 | 29428 | - | 29394 |
| 10_MGAS2096.crt.5 | CP011535.2 Streptococcus pyogenes strain M28PF1, complete genome | 35 | 35292 | 65.8 | 1.00E-12 | 35 | 100% | 0 | 0% | 1 | 35 | 100% | 5865 | 5899 | + | 5865 |
| 10_MGAS2096.crt.5 | CP011415.1 Streptococcus pyogenes strain D471, complete genome | 35 | 30199 | 65.8 | 1.00E-12 | 35 | 100% | 0 | 0% | 1 | 35 | 100% | 22063 | 22097 | - | 22063 |
| 10_MGAS2096.crt.5 | CP011414.1 Streptococcus pyogenes JRS4, complete genome | 35 | 30199 | 65.8 | 1.00E-12 | 35 | 100% | 0 | 0% | 1 | 35 | 100% | 22063 | 22097 | - | 22063 |
| 10_MGAS2096.crt.5 | CP007537.1 Streptococcus pyogenes strain AP1, complete genome | 35 | 29027 | 65.8 | 1.00E-12 | 35 | 100% | 0 | 0% | 1 | 35 | 100% | 7511 | 7545 | + | 7511 |
| 10_MGAS2096.crt.5 | AP012335.1 Streptococcus pyogenes JRS4 DNA, complete genome | 35 | 30198 | 65.8 | 1.00E-12 | 35 | 100% | 0 | 0% | 1 | 35 | 100% | 22063 | 22097 | - | 22063 |
| 10_MGAS2096.crt.5 | AE009949.1 Streptococcus pyogenes MGAS8232, complete genome | 35 | 30227 | 65.8 | 1.00E-12 | 35 | 100% | 0 | 0% | 1 | 35 | 100% | 22059 | 22093 | - | 22059 |
| 10_MGAS2096.crt.5 | CP007041.1 Streptococcus pyogenes STAB902, complete genome | 35 | 36899 | 65.8 | 1.00E-12 | 35 | 100% | 0 | 0% | 1 | 35 | 100% | 13366 | 13400 | + | 13366 |
| 10_MGAS2096.crt.5 | CP006366.1 Streptococcus pyogenes HSC5, complete genome | 35 | 37520 | 65.8 | 1.00E-12 | 35 | 100% | 0 | 0% | 1 | 35 | 100% | 31113 | 31147 | - | 31113 |
| 10_MGAS2096.crt.5 | CP003068.1 Streptococcus pyogenes Alab49, complete genome | 35 | 29027 | 65.8 | 1.00E-12 | 35 | 100% | 0 | 0% | 1 | 35 | 100% | 21483 | 21517 | - | 21483 |
| 10_MGAS2096.crt.5 | AM295007.1 Streptococcus pyogenes Manfredo complete genome | 35 | 28976 | 65.8 | 1.00E-12 | 35 | 100% | 0 | 0% | 1 | 35 | 100% | 7465 | 7499 | + | 7465 |
| 10_MGAS2096.crt.6 | CP010449.1 Streptococcus pyogenes strain NGAS322, complete genome | 35 | 38059 | 65.8 | 1.00E-12 | 35 | 100% | 0 | 0% | 1 | 35 | 100% | 36361 | 36395 | + | 36361 |
| 10_MGAS2096.crt.6 | CP007560.1 Streptococcus pyogenes strain NGAS743, complete genome | 35 | 33384 | 65.8 | 1.00E-12 | 35 | 100% | 0 | 0% | 1 | 35 | 100% | 1665 | 1699 | - | 1665 |
| 10_MGAS2096.crt.6 | AE009949.1 Streptococcus pyogenes MGAS8232, complete genome | 35 | 39804 | 65.8 | 1.00E-12 | 35 | 100% | 0 | 0% | 1 | 35 | 100% | 1665 | 1699 | - | 1665 |
| 10_MGAS2096.crt.6 | AM295007.1 Streptococcus pyogenes Manfredo complete genome | 35 | 39447 | 65.8 | 1.00E-12 | 35 | 100% | 0 | 0% | 1 | 35 | 100% | 37749 | 37783 | + | 37749 |
| 10_MGAS2096.crt.6 | A07_1 | 35 | 37110 | 60.2 | 6.00E-11 | 34 | 97% | 0 | 0% | 1 | 35 | 100% | 35742 | 35776 | + | 35742 |
| 10_MGAS2096.crt.6 | BA000034.2 Streptococcus pyogenes SSI-1 DNA, complete genome | 35 | 37117 | 60.2 | 6.00E-11 | 34 | 97% | 0 | 0% | 1 | 35 | 100% | 1335 | 1369 | - | 1335 |
| 10_MGAS2096.crt.6 | AE014074.1 Streptococcus pyogenes MGAS315, complete genome | 35 | 36875 | 60.2 | 6.00E-11 | 34 | 97% | 0 | 0% | 1 | 35 | 100% | 35507 | 35541 | + | 35507 |
| 10_MGAS2096.crt.6 | CP007041.1 Streptococcus pyogenes STAB902, complete genome | 35 | 36899 | 60.2 | 6.00E-11 | 34 | 97% | 0 | 0% | 1 | 35 | 100% | 1335 | 1369 | - | 1335 |
| 10_MGAS2096.crt.7 | CP008695.1 Streptococcus pyogenes strain M23ND, complete genome | 35 | 37509 | 60.2 | 6.00E-11 | 34 | 97% | 0 | 0% | 1 | 35 | 100% | 36269 | 36303 | + | 36269 |
| 10_MGAS2096.crt.8 | A07_1 | 36 | 31011 | 67.6 | 3.00E-13 | 36 | 100% | 0 | 0% | 1 | 36 | 100% | 8161 | 8196 | + | 8161 |
| 10_MGAS2096.crt.8 | BA000034.2 Streptococcus pyogenes SSI-1 DNA, complete genome | 36 | 31011 | 67.6 | 3.00E-13 | 36 | 100% | 0 | 0% | 1 | 36 | 100% | 22816 | 22851 | - | 22816 |
| 10_MGAS2096.crt.8 | CP008926.1 Streptococcus pyogenes strain ATCC 19615, complete genome | 36 | 31521 | 67.6 | 3.00E-13 | 36 | 100% | 0 | 0% | 1 | 36 | 100% | 22816 | 22851 | - | 22816 |
| 10_MGAS2096.crt.9 | A07_1 | 34 | 31011 | 63.9 | 4.00E-12 | 34 | 100% | 0 | 0% | 1 | 34 | 100% | 12261 | 12294 | + | 12261 |
| 10_MGAS2096.crt.9 | BA000034.2 Streptococcus pyogenes SSI-1 DNA, complete genome | 34 | 31011 | 63.9 | 4.00E-12 | 34 | 100% | 0 | 0% | 1 | 34 | 100% | 18718 | 18751 | - | 18718 |
| 10_MGAS2096.crt.9 | CP008926.1 Streptococcus pyogenes strain ATCC 19615, complete genome | 34 | 31521 | 63.9 | 4.00E-12 | 34 | 100% | 0 | 0% | 1 | 34 | 100% | 18718 | 18751 | - | 18718 |
| 11_MGAS10750.crt.1 | AE004092.2 Streptococcus pyogenes M1 GAS, complete genome | 33 | 29920 | 62.1 | 1.00E-11 | 33 | 100% | 0 | 0% | 1 | 33 | 100% | 22970 | 23002 | - | 22970 |
| 11_MGAS10750.crt.1 | CP006366.1 Streptococcus pyogenes HSC5, complete genome | 33 | 39477 | 62.1 | 1.00E-11 | 33 | 100% | 0 | 0% | 1 | 33 | 100% | 27280 | 27312 | - | 27280 |
| 11_MGAS10750.crt.2 | NOHIT | 34 | | | | | | | | | | | | | | |
| 11_MGAS10750.crt.3 | CP010449.1 Streptococcus pyogenes strain NGAS322, complete genome | 35 | 34489 | 65.8 | 1.00E-12 | 35 | 100% | 0 | 0% | 1 | 35 | 100% | 26203 | 26237 | - | 26203 |
| 11_MGAS10750.crt.3 | AE009949.1 Streptococcus pyogenes MGAS8232, complete genome | 35 | 31741 | 65.8 | 1.00E-12 | 35 | 100% | 0 | 0% | 1 | 35 | 100% | 23469 | 23503 | - | 23469 |
| 11_MGAS10750.crt.4 | NOHIT | 33 | | | | | | | | | | | | | | |
| 11_MGAS10750.crt.5 | NOHIT | 34 | | | | | | | | | | | | | | |
| 13_NZ131.crt.1 | NOHIT | 30 | | | | | | | | | | | | | | |
| 13_NZ131.crt.2 | NOHIT | 31 | | | | | | | | | | | | | | |
| 13_NZ131.crt.3 | NOHIT | 30 | | | | | | | | | | | | | | |
| 13_NZ131.crt.4 | NOHIT | 30 | | | | | | | | | | | | | | |
| 13_NZ131.crt.5 | NOHIT | 0 | | | | | | | | | | | | | | |
| 13_NZ131.crt.6 | NOHIT | 35 | | | | | | | | | | | | | | |
| 13_NZ131.crt.7 | NOHIT | 35 | | | | | | | | | | | | | | |
| 13_NZ131.crt.8 | CP008695.1 Streptococcus pyogenes strain M23ND, complete genome | 35 | 27969 | 65.8 | 1.00E-12 | 35 | 100% | 0 | 0% | 1 | 35 | 100% | 8216 | 8250 | - | 8216 |
| 13_NZ131.crt.8 | CP000017.2 Streptococcus pyogenes MGAS5005, complete genome | 35 | 29805 | 60.2 | 6.00E-11 | 34 | 97% | 0 | 0% | 1 | 35 | 100% | 8604 | 8638 | - | 8604 |
| 13_NZ131.crt.8 | CP012045.1 Streptococcus pyogenes strain HKU488, complete genome | 35 | 29804 | 60.2 | 6.00E-11 | 34 | 97% | 0 | 0% | 1 | 35 | 100% | 8604 | 8638 | - | 8604 |
| 13_NZ131.crt.8 | CP008776.1 Streptococcus pyogenes strain 5448, complete genome | 35 | 29805 | 60.2 | 6.00E-11 | 34 | 97% | 0 | 0% | 1 | 35 | 100% | 21168 | 21202 | + | 21168 |
| 13_NZ131.crt.8 | CP007537.1 Streptococcus pyogenes strain AP1, complete genome | 35 | 29748 | 60.2 | 6.00E-11 | 34 | 97% | 0 | 0% | 1 | 35 | 100% | 18221 | 18255 | + | 18221 |
| 13_NZ131.crt.8 | AE009949.1 Streptococcus pyogenes MGAS8232, complete genome | 35 | 31741 | 60.2 | 6.00E-11 | 34 | 97% | 0 | 0% | 1 | 35 | 100% | 12413 | 12447 | - | 12413 |
| 13_NZ131.crt.8 | AP014585.1 Streptococcus pyogenes DNA, complete genome, strain: MTB314 | 35 | 18078 | 60.2 | 6.00E-11 | 34 | 97% | 0 | 0% | 1 | 35 | 100% | 2031 | 2065 | - | 2031 |
| 13_NZ131.crt.8 | AE004092.2 Streptococcus pyogenes M1 GAS, complete genome | 35 | 29920 | 60.2 | 6.00E-11 | 34 | 97% | 0 | 0% | 1 | 35 | 100% | 11605 | 11639 | - | 11605 |
| 13_NZ131.crt.8 | CP006366.1 Streptococcus pyogenes HSC5, complete genome | 35 | 32692 | 60.2 | 6.00E-11 | 34 | 97% | 0 | 0% | 1 | 35 | 100% | 11491 | 11525 | - | 11491 |
| 13_NZ131.crt.8 | CP003901.1 Streptococcus pyogenes A20, complete genome | 35 | 29805 | 60.2 | 6.00E-11 | 34 | 97% | 0 | 0% | 1 | 35 | 100% | 8604 | 8638 | - | 8604 |
| 13_NZ131.crt.8 | AP012491.2 Streptococcus pyogenes M1 476 DNA, complete genome | 35 | 29805 | 60.2 | 6.00E-11 | 34 | 97% | 0 | 0% | 1 | 35 | 100% | 8604 | 8638 | - | 8604 |
| 13_NZ131.crt.9 | NOHIT | 35 | | | | | | | | | | | | | | |
| 13_NZ131.crt.10 | CP000003.1 Streptococcus pyogenes MGAS10394, complete genome | 33 | 45489 | 62.1 | 1.00E-11 | 33 | 100% | 0 | 0% | 1 | 33 | 100% | 4084 | 4116 | + | 4084 |
| 13_NZ131.crt.10 | A09_1 | 33 | 22019 | 62.1 | 1.00E-11 | 33 | 100% | 0 | 0% | 1 | 33 | 100% | 17904 | 17936 | - | 17904 |
| 13_NZ131.crt.10 | A06_1 | 33 | 22019 | 62.1 | 1.00E-11 | 33 | 100% | 0 | 0% | 1 | 33 | 100% | 17904 | 17936 | - | 17904 |
| 13_NZ131.crt.10 | CP013838.1 Streptococcus pyogenes strain MGAS11027, complete genome | 33 | 35828 | 62.1 | 1.00E-11 | 33 | 100% | 0 | 0% | 1 | 33 | 100% | 31699 | 31731 | - | 31699 |
| 13_NZ131.crt.10 | CP013838.1 Streptococcus pyogenes strain MGAS11027, complete genome | 33 | 51647 | 62.1 | 1.00E-11 | 33 | 100% | 0 | 0% | 1 | 33 | 100% | 33171 | 33203 | - | 33171 |
| 15_MGAS15252.crt.1 | NOHIT | 37 | | | | | | | | | | | | | | |
| 15_MGAS15252.crt.2 | NOHIT | 37 | | | | | | | | | | | | | | |
| 16_MGAS1882.crt.1 | NOHIT | 30 | | | | | | | | | | | | | | |
| 16_MGAS1882.crt.2 | NOHIT | 30 | | | | | | | | | | | | | | |
| 16_MGAS1882.crt.3 | NOHIT | 30 | | | | | | | | | | | | | | |
| 17_M1_476.crt.1 | NOHIT | 30 | | | | | | | | | | | | | | |
| 17_M1_476.crt.2 | NOHIT | 30 | | | | | | | | | | | | | | |
| 17_M1_476.crt.3 | NOHIT | 30 | | | | | | | | | | | | | | |
| 17_M1_476.crt.4 | NOHIT | 0 | | | | | | | | | | | | | | |
| 17_M1_476.crt.5 | CP007561.1 Streptococcus pyogenes strain NGAS596, complete genome | 33 | 40047 | 62.1 | 1.00E-11 | 33 | 100% | 0 | 0% | 1 | 33 | 100% | 4156 | 4188 | - | 4156 |
| 17_M1_476.crt.5 | CP008695.1 Streptococcus pyogenes strain M23ND, complete genome | 33 | 37509 | 62.1 | 1.00E-11 | 33 | 100% | 0 | 0% | 1 | 33 | 100% | 34682 | 34714 | + | 34682 |
| 17_M1_476.crt.5 | AM295007.1 Streptococcus pyogenes Manfredo complete genome | 33 | 37666 | 62.1 | 1.00E-11 | 33 | 100% | 0 | 0% | 1 | 33 | 100% | 4183 | 4215 | - | 4183 |
| 17_M1_476.crt.6 | NOHIT | 34 | | | | | | | | | | | | | | |
| 17_M1_476.crt.7 | CP008695.1 Streptococcus pyogenes strain M23ND, complete genome | 33 | 37458 | 62.1 | 1.00E-11 | 33 | 100% | 0 | 0% | 1 | 33 | 100% | 21052 | 21084 | + | 21052 |
| 17_M1_476.crt.8 | A08_1 | 35 | 36269 | 60.2 | 6.00E-11 | 34 | 97% | 0 | 0% | 1 | 35 | 100% | 28369 | 28403 | - | 28369 |
| 17_M1_476.crt.8 | CP013838.1 Streptococcus pyogenes strain MGAS11027, complete genome | 35 | 51647 | 60.2 | 6.00E-11 | 34 | 97% | 0 | 0% | 1 | 35 | 100% | 13944 | 13978 | + | 13944 |
| 17_M1_476.crt.8 | CP007560.1 Streptococcus pyogenes strain NGAS743, complete genome | 35 | 38050 | 60.2 | 6.00E-11 | 34 | 97% | 0 | 0% | 1 | 35 | 100% | 27864 | 27898 | - | 27864 |
| 17_M1_476.crt.8 | CP007560.1 Streptococcus pyogenes strain NGAS743, complete genome | 35 | 39059 | 60.2 | 6.00E-11 | 34 | 97% | 0 | 0% | 1 | 35 | 100% | 14542 | 14576 | + | 14542 |
| 17_M1_476.crt.8 | AE009949.1 Streptococcus pyogenes MGAS8232, complete genome | 35 | 39804 | 60.2 | 6.00E-11 | 34 | 97% | 0 | 0% | 1 | 35 | 100% | 12058 | 12092 | + | 12058 |
| 17_M1_476.crt.8 | CP009612.1 Streptococcus pyogenes strain HKU360, complete genome | 35 | 38759 | 60.2 | 6.00E-11 | 34 | 97% | 0 | 0% | 1 | 35 | 100% | 26510 | 26544 | - | 26510 |
| 17_M1_476.crt.8 | CP008695.1 Streptococcus pyogenes strain M23ND, complete genome | 35 | 37458 | 60.2 | 6.00E-11 | 34 | 97% | 0 | 0% | 1 | 35 | 100% | 12800 | 12834 | + | 12800 |
| 17_M1_476.crt.8 | CP006366.1 Streptococcus pyogenes HSC5, complete genome | 35 | 39477 | 60.2 | 6.00E-11 | 34 | 97% | 0 | 0% | 1 | 35 | 100% | 24623 | 24657 | - | 24623 |
| 17_M1_476.crt.8 | AM295007.1 Streptococcus pyogenes Manfredo complete genome | 35 | 39447 | 60.2 | 6.00E-11 | 34 | 97% | 0 | 0% | 1 | 35 | 100% | 27341 | 27375 | - | 27341 |
| 18_A20.crt.1 | NOHIT | 30 | | | | | | | | | | | | | | |
| 18_A20.crt.2 | NOHIT | 30 | | | | | | | | | | | | | | |
| 18_A20.crt.3 | NOHIT | 30 | | | | | | | | | | | | | | |
| 18_A20.crt.4 | NOHIT | 0 | | | | | | | | | | | | | | |
| 18_A20.crt.5 | CP007561.1 Streptococcus pyogenes strain NGAS596, complete genome | 33 | 40047 | 62.1 | 1.00E-11 | 33 | 100% | 0 | 0% | 1 | 33 | 100% | 4156 | 4188 | - | 4156 |
| 18_A20.crt.5 | CP008695.1 Streptococcus pyogenes strain M23ND, complete genome | 33 | 37509 | 62.1 | 1.00E-11 | 33 | 100% | 0 | 0% | 1 | 33 | 100% | 34682 | 34714 | + | 34682 |
| 18_A20.crt.5 | AM295007.1 Streptococcus pyogenes Manfredo complete genome | 33 | 37666 | 62.1 | 1.00E-11 | 33 | 100% | 0 | 0% | 1 | 33 | 100% | 4183 | 4215 | - | 4183 |
| 18_A20.crt.6 | NOHIT | 34 | | | | | | | | | | | | | | |
| 18_A20.crt.7 | CP008695.1 Streptococcus pyogenes strain M23ND, complete genome | 33 | 37458 | 62.1 | 1.00E-11 | 33 | 100% | 0 | 0% | 1 | 33 | 100% | 21052 | 21084 | + | 21052 |
| 18_A20.crt.8 | A08_1 | 35 | 36269 | 60.2 | 6.00E-11 | 34 | 97% | 0 | 0% | 1 | 35 | 100% | 28369 | 28403 | - | 28369 |
| 18_A20.crt.8 | CP013838.1 Streptococcus pyogenes strain MGAS11027, complete genome | 35 | 51647 | 60.2 | 6.00E-11 | 34 | 97% | 0 | 0% | 1 | 35 | 100% | 13944 | 13978 | + | 13944 |
| 18_A20.crt.8 | CP007560.1 Streptococcus pyogenes strain NGAS743, complete genome | 35 | 38050 | 60.2 | 6.00E-11 | 34 | 97% | 0 | 0% | 1 | 35 | 100% | 27864 | 27898 | - | 27864 |
| 18_A20.crt.8 | CP007560.1 Streptococcus pyogenes strain NGAS743, complete genome | 35 | 39059 | 60.2 | 6.00E-11 | 34 | 97% | 0 | 0% | 1 | 35 | 100% | 14542 | 14576 | + | 14542 |
| 18_A20.crt.8 | AE009949.1 Streptococcus pyogenes MGAS8232, complete genome | 35 | 39804 | 60.2 | 6.00E-11 | 34 | 97% | 0 | 0% | 1 | 35 | 100% | 12058 | 12092 | + | 12058 |
| 18_A20.crt.8 | CP009612.1 Streptococcus pyogenes strain HKU360, complete genome | 35 | 38759 | 60.2 | 6.00E-11 | 34 | 97% | 0 | 0% | 1 | 35 | 100% | 26510 | 26544 | - | 26510 |
| 18_A20.crt.8 | CP008695.1 Streptococcus pyogenes strain M23ND, complete genome | 35 | 37458 | 60.2 | 6.00E-11 | 34 | 97% | 0 | 0% | 1 | 35 | 100% | 12800 | 12834 | + | 12800 |
| 18_A20.crt.8 | CP006366.1 Streptococcus pyogenes HSC5, complete genome | 35 | 39477 | 60.2 | 6.00E-11 | 34 | 97% | 0 | 0% | 1 | 35 | 100% | 24623 | 24657 | - | 24623 |
| 18_A20.crt.8 | AM295007.1 Streptococcus pyogenes Manfredo complete genome | 35 | 39447 | 60.2 | 6.00E-11 | 34 | 97% | 0 | 0% | 1 | 35 | 100% | 27341 | 27375 | - | 27341 |
| 1_SF370.crt.1 | NOHIT | 30 | | | | | | | | | | | | | | |
| 1_SF370.crt.2 | NOHIT | 30 | | | | | | | | | | | | | | |
| 1_SF370.crt.3 | NOHIT | 30 | | | | | | | | | | | | | | |
| 1_SF370.crt.4 | NOHIT | 30 | | | | | | | | | | | | | | |
| 1_SF370.crt.5 | NOHIT | 30 | | | | | | | | | | | | | | |
| 1_SF370.crt.6 | NOHIT | 30 | | | | | | | | | | | | | | |
| 1_SF370.crt.7 | NOHIT | 0 | | | | | | | | | | | | | | |
| 1_SF370.crt.8 | NOHIT | 33 | | | | | | | | | | | | | | |
| 1_SF370.crt.9 | CP008695.1 Streptococcus pyogenes strain M23ND, complete genome | 32 | 37458 | 60.2 | 5.00E-11 | 32 | 100% | 0 | 0% | 1 | 32 | 100% | 21052 | 21083 | + | 21052 |
| 1_SF370.crt.10 | NOHIT | 34 | | | | | | | | | | | | | | |
| 20_H293.crt.1 | NOHIT | 30 | | | | | | | | | | | | | | |
| 20_H293.crt.2 | NOHIT | 30 | | | | | | | | | | | | | | |
| 20_H293.crt.3 | NOHIT | 30 | | | | | | | | | | | | | | |
| 20_H293.crt.4 | NOHIT | 30 | | | | | | | | | | | | | | |
| 20_H293.crt.5 | NOHIT | 30 | | | | | | | | | | | | | | |
| 20_H293.crt.6 | NOHIT | 0 | | | | | | | | | | | | | | |
| 20_H293.crt.7 | NOHIT | 19 | | | | | | | | | | | | | | |
| 20_H293.crt.8 | NOHIT | 28 | | | | | | | | | | | | | | |
| 20_H293.crt.9 | NOHIT | 19 | | | | | | | | | | | | | | |
| 20_H293.crt.10 | NOHIT | 19 | | | | | | | | | | | | | | |
| 20_H293.crt.11 | NOHIT | 0 | | | | | | | | | | | | | | |
| 20_H293.crt.12 | CP000003.1 Streptococcus pyogenes MGAS10394, complete genome | 34 | 35956 | 62.1 | 1.00E-11 | 33 | 100% | 0 | 0% | 1 | 33 | 97.06% | 32035 | 32067 | + | 32035 |
| 20_H293.crt.12 | A09_1 | 34 | 35957 | 62.1 | 1.00E-11 | 33 | 100% | 0 | 0% | 1 | 33 | 97.06% | 32036 | 32068 | + | 32036 |
| 20_H293.crt.12 | A07_1 | 34 | 36064 | 62.1 | 1.00E-11 | 33 | 100% | 0 | 0% | 1 | 33 | 97.06% | 3998 | 4030 | - | 3998 |
| 20_H293.crt.12 | A07_1 | 34 | 31011 | 62.1 | 1.00E-11 | 33 | 100% | 0 | 0% | 1 | 33 | 97.06% | 27079 | 27111 | + | 27079 |
| 20_H293.crt.12 | BA000034.2 Streptococcus pyogenes SSI-1 DNA, complete genome | 34 | 31011 | 62.1 | 1.00E-11 | 33 | 100% | 0 | 0% | 1 | 33 | 97.06% | 3901 | 3933 | - | 3901 |
| 20_H293.crt.12 | BA000034.2 Streptococcus pyogenes SSI-1 DNA, complete genome | 34 | 36064 | 62.1 | 1.00E-11 | 33 | 100% | 0 | 0% | 1 | 33 | 97.06% | 32035 | 32067 | + | 32035 |
| 20_H293.crt.12 | AE014074.1 Streptococcus pyogenes MGAS315, complete genome | 34 | 36064 | 62.1 | 1.00E-11 | 33 | 100% | 0 | 0% | 1 | 33 | 97.06% | 3998 | 4030 | - | 3998 |
| 20_H293.crt.12 | CP008926.1 Streptococcus pyogenes strain ATCC 19615, complete genome | 34 | 52827 | 62.1 | 1.00E-11 | 33 | 100% | 0 | 0% | 1 | 33 | 97.06% | 34829 | 34861 | - | 34829 |
| 20_H293.crt.12 | CP007041.1 Streptococcus pyogenes STAB902, complete genome | 34 | 32470 | 62.1 | 1.00E-11 | 33 | 100% | 0 | 0% | 1 | 33 | 97.06% | 32035 | 32067 | + | 32035 |
| 20_H293.crt.12 | CP000829.1 Streptococcus pyogenes NZ131, complete genome | 34 | 52188 | 62.1 | 1.00E-11 | 33 | 100% | 0 | 0% | 1 | 33 | 97.06% | 34175 | 34207 | - | 34175 |
| 20_H293.crt.13 | A09_1 | 34 | 34696 | 63.9 | 4.00E-12 | 34 | 100% | 0 | 0% | 1 | 34 | 100% | 1215 | 1248 | + | 1215 |
| 20_H293.crt.13 | A08_1 | 34 | 31416 | 63.9 | 4.00E-12 | 34 | 100% | 0 | 0% | 1 | 34 | 100% | 1215 | 1248 | + | 1215 |
| 20_H293.crt.13 | A07_1 | 34 | 28192 | 63.9 | 4.00E-12 | 34 | 100% | 0 | 0% | 1 | 34 | 100% | 26945 | 26978 | - | 26945 |
| 20_H293.crt.13 | A06_1 | 34 | 34711 | 63.9 | 4.00E-12 | 34 | 100% | 0 | 0% | 1 | 34 | 100% | 1215 | 1248 | + | 1215 |
| 20_H293.crt.13 | BA000034.2 Streptococcus pyogenes SSI-1 DNA, complete genome | 34 | 28192 | 63.9 | 4.00E-12 | 34 | 100% | 0 | 0% | 1 | 34 | 100% | 1215 | 1248 | + | 1215 |
| 20_H293.crt.13 | CP013672.1 Streptococcus pyogenes strain AP53, complete genome | 34 | 24636 | 63.9 | 4.00E-12 | 34 | 100% | 0 | 0% | 1 | 34 | 100% | 23389 | 23422 | - | 23389 |
| 20_H293.crt.13 | CP014027.1 Streptococcus pyogenes strain FDAARGOS_149, complete genome | 34 | 28254 | 63.9 | 4.00E-12 | 34 | 100% | 0 | 0% | 1 | 34 | 100% | 1215 | 1248 | + | 1215 |
| 20_H293.crt.13 | CP010449.1 Streptococcus pyogenes strain NGAS322, complete genome | 34 | 34489 | 63.9 | 4.00E-12 | 34 | 100% | 0 | 0% | 1 | 34 | 100% | 33242 | 33275 | - | 33242 |
| 20_H293.crt.13 | AE014074.1 Streptococcus pyogenes MGAS315, complete genome | 34 | 28192 | 63.9 | 4.00E-12 | 34 | 100% | 0 | 0% | 1 | 34 | 100% | 26945 | 26978 | - | 26945 |
| 20_H293.crt.13 | CP007560.1 Streptococcus pyogenes strain NGAS743, complete genome | 34 | 38050 | 63.9 | 4.00E-12 | 34 | 100% | 0 | 0% | 1 | 34 | 100% | 36803 | 36836 | - | 36803 |
| 20_H293.crt.13 | CP011415.1 Streptococcus pyogenes strain D471, complete genome | 34 | 33148 | 63.9 | 4.00E-12 | 34 | 100% | 0 | 0% | 1 | 34 | 100% | 31901 | 31934 | - | 31901 |
| 20_H293.crt.13 | CP011414.1 Streptococcus pyogenes JRS4, complete genome | 34 | 33148 | 63.9 | 4.00E-12 | 34 | 100% | 0 | 0% | 1 | 34 | 100% | 31901 | 31934 | - | 31901 |
| 20_H293.crt.13 | CP007537.1 Streptococcus pyogenes strain AP1, complete genome | 34 | 29748 | 63.9 | 4.00E-12 | 34 | 100% | 0 | 0% | 1 | 34 | 100% | 1215 | 1248 | + | 1215 |
| 20_H293.crt.13 | AP012335.1 Streptococcus pyogenes JRS4 DNA, complete genome | 34 | 33148 | 63.9 | 4.00E-12 | 34 | 100% | 0 | 0% | 1 | 34 | 100% | 31901 | 31934 | - | 31901 |
| 20_H293.crt.13 | AE009949.1 Streptococcus pyogenes MGAS8232, complete genome | 34 | 31741 | 63.9 | 4.00E-12 | 34 | 100% | 0 | 0% | 1 | 34 | 100% | 30494 | 30527 | - | 30494 |
| 20_H293.crt.13 | CP009612.1 Streptococcus pyogenes strain HKU360, complete genome | 34 | 38759 | 63.9 | 4.00E-12 | 34 | 100% | 0 | 0% | 1 | 34 | 100% | 37512 | 37545 | - | 37512 |
| 20_H293.crt.13 | CP007041.1 Streptococcus pyogenes STAB902, complete genome | 34 | 28192 | 63.9 | 4.00E-12 | 34 | 100% | 0 | 0% | 1 | 34 | 100% | 1215 | 1248 | + | 1215 |
| 20_H293.crt.13 | AE004092.2 Streptococcus pyogenes M1 GAS, complete genome | 34 | 29920 | 63.9 | 4.00E-12 | 34 | 100% | 0 | 0% | 1 | 34 | 100% | 28673 | 28706 | - | 28673 |
| 20_H293.crt.13 | CP003068.1 Streptococcus pyogenes Alab49, complete genome | 34 | 24636 | 63.9 | 4.00E-12 | 34 | 100% | 0 | 0% | 1 | 34 | 100% | 23389 | 23422 | - | 23389 |
| 20_H293.crt.13 | CP000262.1 Streptococcus pyogenes MGAS10750, complete genome | 34 | 31381 | 63.9 | 4.00E-12 | 34 | 100% | 0 | 0% | 1 | 34 | 100% | 30134 | 30167 | - | 30134 |
| 20_H293.crt.13 | CP011415.1 Streptococcus pyogenes strain D471, complete genome | 34 | 39110 | 62.1 | 1.00E-11 | 33 | 100% | 0 | 0% | 2 | 34 | 97.06% | 34851 | 34883 | - | 34851 |
| 20_H293.crt.13 | CP011414.1 Streptococcus pyogenes JRS4, complete genome | 34 | 39110 | 62.1 | 1.00E-11 | 33 | 100% | 0 | 0% | 2 | 34 | 97.06% | 34851 | 34883 | - | 34851 |
| 20_H293.crt.13 | AP012335.1 Streptococcus pyogenes JRS4 DNA, complete genome | 34 | 39111 | 62.1 | 1.00E-11 | 33 | 100% | 0 | 0% | 2 | 34 | 97.06% | 34852 | 34884 | - | 34852 |
| 20_H293.crt.14 | CP007561.1 Streptococcus pyogenes strain NGAS596, complete genome | 37 | 40047 | 63.9 | 5.00E-12 | 34 | 100% | 0 | 0% | 4 | 37 | 91.89% | 26778 | 26811 | - | 26778 |
| 20_H293.crt.14 | CP011415.1 Streptococcus pyogenes strain D471, complete genome | 37 | 39110 | 63.9 | 5.00E-12 | 34 | 100% | 0 | 0% | 4 | 37 | 91.89% | 25212 | 25245 | - | 25212 |
| 20_H293.crt.14 | CP011414.1 Streptococcus pyogenes JRS4, complete genome | 37 | 39110 | 63.9 | 5.00E-12 | 34 | 100% | 0 | 0% | 4 | 37 | 91.89% | 25212 | 25245 | - | 25212 |
| 20_H293.crt.14 | AP012335.1 Streptococcus pyogenes JRS4 DNA, complete genome | 37 | 39111 | 63.9 | 5.00E-12 | 34 | 100% | 0 | 0% | 4 | 37 | 91.89% | 25212 | 25245 | - | 25212 |
| 20_H293.crt.14 | CP008695.1 Streptococcus pyogenes strain M23ND, complete genome | 37 | 37509 | 63.9 | 5.00E-12 | 34 | 100% | 0 | 0% | 4 | 37 | 91.89% | 12117 | 12150 | + | 12117 |
| 20_H293.crt.14 | CP000829.1 Streptococcus pyogenes NZ131, complete genome | 37 | 33890 | 63.9 | 5.00E-12 | 34 | 100% | 0 | 0% | 4 | 37 | 91.89% | 11020 | 11053 | + | 11020 |
| 20_H293.crt.14 | AM295007.1 Streptococcus pyogenes Manfredo complete genome | 37 | 37666 | 63.9 | 5.00E-12 | 34 | 100% | 0 | 0% | 4 | 37 | 91.89% | 26614 | 26647 | - | 26614 |
| 20_H293.crt.15 | NOHIT | 35 | | | | | | | | | | | | | | |
| 20_H293.crt.16 | CP000260.1 Streptococcus pyogenes MGAS10270, complete genome | 34 | 39966 | 63.9 | 4.00E-12 | 34 | 100% | 0 | 0% | 1 | 34 | 100% | 25392 | 25425 | + | 25392 |
| 20_H293.crt.16 | CP000056.1 Streptococcus pyogenes MGAS6180, complete genome | 34 | 36896 | 63.9 | 4.00E-12 | 34 | 100% | 0 | 0% | 1 | 34 | 100% | 25878 | 25911 | + | 25878 |
| 20_H293.crt.16 | A09_1 | 34 | 30199 | 63.9 | 4.00E-12 | 34 | 100% | 0 | 0% | 1 | 34 | 100% | 12024 | 12057 | - | 12024 |
| 20_H293.crt.16 | A08_1 | 34 | 32841 | 63.9 | 4.00E-12 | 34 | 100% | 0 | 0% | 1 | 34 | 100% | 14542 | 14575 | - | 14542 |
| 20_H293.crt.16 | A07_1 | 34 | 39948 | 63.9 | 4.00E-12 | 34 | 100% | 0 | 0% | 1 | 34 | 100% | 25359 | 25392 | + | 25359 |
| 20_H293.crt.16 | A06_1 | 34 | 30200 | 63.9 | 4.00E-12 | 34 | 100% | 0 | 0% | 1 | 34 | 100% | 12025 | 12058 | - | 12025 |
| 20_H293.crt.16 | BA000034.2 Streptococcus pyogenes SSI-1 DNA, complete genome | 34 | 39933 | 63.9 | 4.00E-12 | 34 | 100% | 0 | 0% | 1 | 34 | 100% | 14542 | 14575 | - | 14542 |
| 20_H293.crt.16 | AE014074.1 Streptococcus pyogenes MGAS315, complete genome | 34 | 39933 | 63.9 | 4.00E-12 | 34 | 100% | 0 | 0% | 1 | 34 | 100% | 25359 | 25392 | + | 25359 |
| 20_H293.crt.16 | CP011068.1 Streptococcus pyogenes strain STAB10015, complete genome | 34 | 36896 | 63.9 | 4.00E-12 | 34 | 100% | 0 | 0% | 1 | 34 | 100% | 25878 | 25911 | + | 25878 |
| 20_H293.crt.16 | CP011535.2 Streptococcus pyogenes strain M28PF1, complete genome | 34 | 32154 | 63.9 | 4.00E-12 | 34 | 100% | 0 | 0% | 1 | 34 | 100% | 6245 | 6278 | - | 6245 |
| 20_H293.crt.16 | CP007560.1 Streptococcus pyogenes strain NGAS743, complete genome | 34 | 39618 | 63.9 | 4.00E-12 | 34 | 100% | 0 | 0% | 1 | 34 | 100% | 22005 | 22038 | + | 22005 |
| 20_H293.crt.16 | CP011415.1 Streptococcus pyogenes strain D471, complete genome | 34 | 30199 | 63.9 | 4.00E-12 | 34 | 100% | 0 | 0% | 1 | 34 | 100% | 18142 | 18175 | + | 18142 |
| 20_H293.crt.16 | CP011414.1 Streptococcus pyogenes JRS4, complete genome | 34 | 30199 | 63.9 | 4.00E-12 | 34 | 100% | 0 | 0% | 1 | 34 | 100% | 18142 | 18175 | + | 18142 |
| 20_H293.crt.16 | AP012335.1 Streptococcus pyogenes JRS4 DNA, complete genome | 34 | 30198 | 63.9 | 4.00E-12 | 34 | 100% | 0 | 0% | 1 | 34 | 100% | 18142 | 18175 | + | 18142 |
| 20_H293.crt.16 | AE009949.1 Streptococcus pyogenes MGAS8232, complete genome | 34 | 30227 | 63.9 | 4.00E-12 | 34 | 100% | 0 | 0% | 1 | 34 | 100% | 18138 | 18171 | + | 18138 |
| 20_H293.crt.16 | CP007041.1 Streptococcus pyogenes STAB902, complete genome | 34 | 32577 | 63.9 | 4.00E-12 | 34 | 100% | 0 | 0% | 1 | 34 | 100% | 14542 | 14575 | - | 14542 |
| 20_H293.crt.17 | NOHIT | 35 | | | | | | | | | | | | | | |
| 20_H293.crt.18 | NOHIT | 33 | | | | | | | | | | | | | | |
| 20_H293.crt.19 | NOHIT | 34 | | | | | | | | | | | | | | |
| 20_H293.crt.20 | NOHIT | 37 | | | | | | | | | | | | | | |
| 20_H293.crt.21 | NOHIT | 33 | | | | | | | | | | | | | | |
| 20_H293.crt.22 | CP000259.1 Streptococcus pyogenes MGAS9429, complete genome | 34 | 37329 | 63.9 | 4.00E-12 | 34 | 100% | 0 | 0% | 1 | 34 | 100% | 2030 | 2063 | - | 2030 |
| 20_H293.crt.22 | CP000259.1 Streptococcus pyogenes MGAS9429, complete genome | 34 | 38133 | 63.9 | 4.00E-12 | 34 | 100% | 0 | 0% | 1 | 34 | 100% | 1826 | 1859 | - | 1826 |
| 20_H293.crt.22 | GAS003 | 34 | 37328 | 63.9 | 4.00E-12 | 34 | 100% | 0 | 0% | 1 | 34 | 100% | 2030 | 2063 | - | 2030 |
| 20_H293.crt.22 | CP009612.1 Streptococcus pyogenes strain HKU360, complete genome | 34 | 37343 | 63.9 | 4.00E-12 | 34 | 100% | 0 | 0% | 1 | 34 | 100% | 2030 | 2063 | - | 2030 |
| 20_H293.crt.23 | NOHIT | 33 | | | | | | | | | | | | | | |
| 20_H293.crt.24 | CP000260.1 Streptococcus pyogenes MGAS10270, complete genome | 34 | 39966 | 63.9 | 4.00E-12 | 34 | 100% | 0 | 0% | 1 | 34 | 100% | 10775 | 10808 | + | 10775 |
| 20_H293.crt.24 | CP000056.1 Streptococcus pyogenes MGAS6180, complete genome | 34 | 36896 | 63.9 | 4.00E-12 | 34 | 100% | 0 | 0% | 1 | 34 | 100% | 11261 | 11294 | + | 11261 |
| 20_H293.crt.24 | CP000003.1 Streptococcus pyogenes MGAS10394, complete genome | 34 | 15429 | 63.9 | 4.00E-12 | 34 | 100% | 0 | 0% | 1 | 34 | 100% | 3489 | 3522 | + | 3489 |
| 20_H293.crt.24 | A09_1 | 34 | 30199 | 63.9 | 4.00E-12 | 34 | 100% | 0 | 0% | 1 | 34 | 100% | 26678 | 26711 | - | 26678 |
| 20_H293.crt.24 | A08_1 | 34 | 32841 | 63.9 | 4.00E-12 | 34 | 100% | 0 | 0% | 1 | 34 | 100% | 29158 | 29191 | - | 29158 |
| 20_H293.crt.24 | A07_1 | 34 | 39948 | 63.9 | 4.00E-12 | 34 | 100% | 0 | 0% | 1 | 34 | 100% | 10742 | 10775 | + | 10742 |
| 20_H293.crt.24 | A06_1 | 34 | 30200 | 63.9 | 4.00E-12 | 34 | 100% | 0 | 0% | 1 | 34 | 100% | 26679 | 26712 | - | 26679 |
| 20_H293.crt.24 | GAS004 | 34 | 34136 | 63.9 | 4.00E-12 | 34 | 100% | 0 | 0% | 1 | 34 | 100% | 30675 | 30708 | - | 30675 |
| 20_H293.crt.24 | BA000034.2 Streptococcus pyogenes SSI-1 DNA, complete genome | 34 | 39933 | 63.9 | 4.00E-12 | 34 | 100% | 0 | 0% | 1 | 34 | 100% | 29159 | 29192 | - | 29159 |
| 20_H293.crt.24 | CP013838.1 Streptococcus pyogenes strain MGAS11027, complete genome | 34 | 35828 | 63.9 | 4.00E-12 | 34 | 100% | 0 | 0% | 1 | 34 | 100% | 28789 | 28822 | - | 28789 |
| 20_H293.crt.24 | CP013838.1 Streptococcus pyogenes strain MGAS11027, complete genome | 34 | 51647 | 63.9 | 4.00E-12 | 34 | 100% | 0 | 0% | 1 | 34 | 100% | 30261 | 30294 | - | 30261 |
| 20_H293.crt.24 | CP013672.1 Streptococcus pyogenes strain AP53, complete genome | 34 | 29027 | 63.9 | 4.00E-12 | 34 | 100% | 0 | 0% | 1 | 34 | 100% | 3381 | 3414 | + | 3381 |
| 20_H293.crt.24 | AE014074.1 Streptococcus pyogenes MGAS315, complete genome | 34 | 39933 | 63.9 | 4.00E-12 | 34 | 100% | 0 | 0% | 1 | 34 | 100% | 10742 | 10775 | + | 10742 |
| 20_H293.crt.24 | CP011068.1 Streptococcus pyogenes strain STAB10015, complete genome | 34 | 36896 | 63.9 | 4.00E-12 | 34 | 100% | 0 | 0% | 1 | 34 | 100% | 11261 | 11294 | + | 11261 |
| 20_H293.crt.24 | CP011535.2 Streptococcus pyogenes strain M28PF1, complete genome | 34 | 32154 | 63.9 | 4.00E-12 | 34 | 100% | 0 | 0% | 1 | 34 | 100% | 20862 | 20895 | - | 20862 |
| 20_H293.crt.24 | CP007560.1 Streptococcus pyogenes strain NGAS743, complete genome | 34 | 38050 | 63.9 | 4.00E-12 | 34 | 100% | 0 | 0% | 1 | 34 | 100% | 9472 | 9505 | + | 9472 |
| 20_H293.crt.24 | CP007560.1 Streptococcus pyogenes strain NGAS743, complete genome | 34 | 39618 | 63.9 | 4.00E-12 | 34 | 100% | 0 | 0% | 1 | 34 | 100% | 7387 | 7420 | + | 7387 |
| 20_H293.crt.24 | CP011415.1 Streptococcus pyogenes strain D471, complete genome | 34 | 30199 | 63.9 | 4.00E-12 | 34 | 100% | 0 | 0% | 1 | 34 | 100% | 3489 | 3522 | + | 3489 |
| 20_H293.crt.24 | CP011414.1 Streptococcus pyogenes JRS4, complete genome | 34 | 30199 | 63.9 | 4.00E-12 | 34 | 100% | 0 | 0% | 1 | 34 | 100% | 3489 | 3522 | + | 3489 |
| 20_H293.crt.24 | CP007537.1 Streptococcus pyogenes strain AP1, complete genome | 34 | 29027 | 63.9 | 4.00E-12 | 34 | 100% | 0 | 0% | 1 | 34 | 100% | 25614 | 25647 | - | 25614 |
| 20_H293.crt.24 | AP012335.1 Streptococcus pyogenes JRS4 DNA, complete genome | 34 | 30198 | 63.9 | 4.00E-12 | 34 | 100% | 0 | 0% | 1 | 34 | 100% | 3489 | 3522 | + | 3489 |
| 20_H293.crt.24 | AE009949.1 Streptococcus pyogenes MGAS8232, complete genome | 34 | 30227 | 63.9 | 4.00E-12 | 34 | 100% | 0 | 0% | 1 | 34 | 100% | 3484 | 3517 | + | 3484 |
| 20_H293.crt.24 | CP007041.1 Streptococcus pyogenes STAB902, complete genome | 34 | 32577 | 63.9 | 4.00E-12 | 34 | 100% | 0 | 0% | 1 | 34 | 100% | 29160 | 29193 | - | 29160 |
| 20_H293.crt.24 | AE004092.2 Streptococcus pyogenes M1 GAS, complete genome | 34 | 39657 | 63.9 | 4.00E-12 | 34 | 100% | 0 | 0% | 1 | 34 | 100% | 30675 | 30708 | - | 30675 |
| 20_H293.crt.24 | CP006366.1 Streptococcus pyogenes HSC5, complete genome | 34 | 39477 | 63.9 | 4.00E-12 | 34 | 100% | 0 | 0% | 1 | 34 | 100% | 6858 | 6891 | + | 6858 |
| 20_H293.crt.24 | CP003068.1 Streptococcus pyogenes Alab49, complete genome | 34 | 29027 | 63.9 | 4.00E-12 | 34 | 100% | 0 | 0% | 1 | 34 | 100% | 3381 | 3414 | + | 3381 |
| 20_H293.crt.24 | AM295007.1 Streptococcus pyogenes Manfredo complete genome | 34 | 28976 | 63.9 | 4.00E-12 | 34 | 100% | 0 | 0% | 1 | 34 | 100% | 25563 | 25596 | - | 25563 |
| 20_H293.crt.24 | CP000261.1 Streptococcus pyogenes MGAS2096, complete genome | 34 | 40056 | 63.9 | 4.00E-12 | 34 | 100% | 0 | 0% | 1 | 34 | 100% | 30675 | 30708 | - | 30675 |
| 22_STAB901.crt.1 | NOHIT | 30 | | | | | | | | | | | | | | |
| 22_STAB901.crt.2 | NOHIT | 30 | | | | | | | | | | | | | | |
| 22_STAB901.crt.3 | NOHIT | 30 | | | | | | | | | | | | | | |
| 22_STAB901.crt.4 | NOHIT | 30 | | | | | | | | | | | | | | |
| 22_STAB901.crt.5 | NOHIT | 0 | | | | | | | | | | | | | | |
| 22_STAB901.crt.6 | NOHIT | 29 | | | | | | | | | | | | | | |
| 22_STAB901.crt.7 | NOHIT | 31 | | | | | | | | | | | | | | |
| 22_STAB901.crt.8 | A07_1 | 32 | 31011 | 60.2 | 5.00E-11 | 32 | 100% | 0 | 0% | 1 | 32 | 100% | 7277 | 7308 | + | 7277 |
| 22_STAB901.crt.8 | BA000034.2 Streptococcus pyogenes SSI-1 DNA, complete genome | 32 | 31011 | 60.2 | 5.00E-11 | 32 | 100% | 0 | 0% | 1 | 32 | 100% | 23704 | 23735 | - | 23704 |
| 22_STAB901.crt.8 | CP008926.1 Streptococcus pyogenes strain ATCC 19615, complete genome | 32 | 31521 | 60.2 | 5.00E-11 | 32 | 100% | 0 | 0% | 1 | 32 | 100% | 23705 | 23736 | - | 23705 |
| 25_7F7.crt.1 | NOHIT | 30 | | | | | | | | | | | | | | |
| 25_7F7.crt.2 | NOHIT | 30 | | | | | | | | | | | | | | |
| 25_7F7.crt.3 | NOHIT | 30 | | | | | | | | | | | | | | |
| 25_7F7.crt.4 | NOHIT | 30 | | | | | | | | | | | | | | |
| 25_7F7.crt.5 | NOHIT | 30 | | | | | | | | | | | | | | |
| 25_7F7.crt.6 | NOHIT | 30 | | | | | | | | | | | | | | |
| 25_7F7.crt.7 | NOHIT | 30 | | | | | | | | | | | | | | |
| 25_7F7.crt.8 | NOHIT | 30 | | | | | | | | | | | | | | |
| 26_HKU360.crt.1 | NOHIT | 38 | | | | | | | | | | | | | | |
| 26_HKU360.crt.2 | NOHIT | 37 | | | | | | | | | | | | | | |
| 26_HKU360.crt.3 | NOHIT | 0 | | | | | | | | | | | | | | |
| 26_HKU360.crt.4 | NOHIT | 32 | | | | | | | | | | | | | | |
| 26_HKU360.crt.5 | CP000056.1 Streptococcus pyogenes MGAS6180, complete genome | 34 | 35292 | 63.9 | 4.00E-12 | 34 | 100% | 0 | 0% | 1 | 34 | 100% | 29395 | 29428 | - | 29395 |
| 26_HKU360.crt.5 | A09_1 | 34 | 30199 | 63.9 | 4.00E-12 | 34 | 100% | 0 | 0% | 1 | 34 | 100% | 8103 | 8136 | + | 8103 |
| 26_HKU360.crt.5 | A07_1 | 34 | 37110 | 63.9 | 4.00E-12 | 34 | 100% | 0 | 0% | 1 | 34 | 100% | 23711 | 23744 | - | 23711 |
| 26_HKU360.crt.5 | A06_1 | 34 | 30200 | 63.9 | 4.00E-12 | 34 | 100% | 0 | 0% | 1 | 34 | 100% | 8103 | 8136 | + | 8103 |
| 26_HKU360.crt.5 | CP011069.1 Streptococcus pyogenes strain STAB09014, complete genome | 34 | 35292 | 63.9 | 4.00E-12 | 34 | 100% | 0 | 0% | 1 | 34 | 100% | 29395 | 29428 | - | 29395 |
| 26_HKU360.crt.5 | BA000034.2 Streptococcus pyogenes SSI-1 DNA, complete genome | 34 | 37117 | 63.9 | 4.00E-12 | 34 | 100% | 0 | 0% | 1 | 34 | 100% | 13367 | 13400 | + | 13367 |
| 26_HKU360.crt.5 | CP013672.1 Streptococcus pyogenes strain AP53, complete genome | 34 | 29027 | 63.9 | 4.00E-12 | 34 | 100% | 0 | 0% | 1 | 34 | 100% | 21484 | 21517 | - | 21484 |
| 26_HKU360.crt.5 | CP014139.1 Streptococcus pyogenes strain MEW123, complete genome | 34 | 34850 | 63.9 | 4.00E-12 | 34 | 100% | 0 | 0% | 1 | 34 | 100% | 15233 | 15266 | + | 15233 |
| 26_HKU360.crt.5 | CP010450.1 Streptococcus pyogenes strain NGAS638, complete genome | 34 | 35488 | 63.9 | 4.00E-12 | 34 | 100% | 0 | 0% | 1 | 34 | 100% | 29071 | 29104 | - | 29071 |
| 26_HKU360.crt.5 | AE014074.1 Streptococcus pyogenes MGAS315, complete genome | 34 | 36875 | 63.9 | 4.00E-12 | 34 | 100% | 0 | 0% | 1 | 34 | 100% | 23487 | 23520 | - | 23487 |
| 26_HKU360.crt.5 | CP011068.1 Streptococcus pyogenes strain STAB10015, complete genome | 34 | 35292 | 63.9 | 4.00E-12 | 34 | 100% | 0 | 0% | 1 | 34 | 100% | 29395 | 29428 | - | 29395 |
| 26_HKU360.crt.5 | CP011535.2 Streptococcus pyogenes strain M28PF1, complete genome | 34 | 35292 | 63.9 | 4.00E-12 | 34 | 100% | 0 | 0% | 1 | 34 | 100% | 5865 | 5898 | + | 5865 |
| 26_HKU360.crt.5 | CP011415.1 Streptococcus pyogenes strain D471, complete genome | 34 | 30199 | 63.9 | 4.00E-12 | 34 | 100% | 0 | 0% | 1 | 34 | 100% | 22064 | 22097 | - | 22064 |
| 26_HKU360.crt.5 | CP011414.1 Streptococcus pyogenes JRS4, complete genome | 34 | 30199 | 63.9 | 4.00E-12 | 34 | 100% | 0 | 0% | 1 | 34 | 100% | 22064 | 22097 | - | 22064 |
| 26_HKU360.crt.5 | CP007537.1 Streptococcus pyogenes strain AP1, complete genome | 34 | 29027 | 63.9 | 4.00E-12 | 34 | 100% | 0 | 0% | 1 | 34 | 100% | 7511 | 7544 | + | 7511 |
| 26_HKU360.crt.5 | AP012335.1 Streptococcus pyogenes JRS4 DNA, complete genome | 34 | 30198 | 63.9 | 4.00E-12 | 34 | 100% | 0 | 0% | 1 | 34 | 100% | 22064 | 22097 | - | 22064 |
| 26_HKU360.crt.5 | AE009949.1 Streptococcus pyogenes MGAS8232, complete genome | 34 | 30227 | 63.9 | 4.00E-12 | 34 | 100% | 0 | 0% | 1 | 34 | 100% | 22060 | 22093 | - | 22060 |
| 26_HKU360.crt.5 | CP007041.1 Streptococcus pyogenes STAB902, complete genome | 34 | 36899 | 63.9 | 4.00E-12 | 34 | 100% | 0 | 0% | 1 | 34 | 100% | 13366 | 13399 | + | 13366 |
| 26_HKU360.crt.5 | CP006366.1 Streptococcus pyogenes HSC5, complete genome | 34 | 37520 | 63.9 | 4.00E-12 | 34 | 100% | 0 | 0% | 1 | 34 | 100% | 31114 | 31147 | - | 31114 |
| 26_HKU360.crt.5 | CP003068.1 Streptococcus pyogenes Alab49, complete genome | 34 | 29027 | 63.9 | 4.00E-12 | 34 | 100% | 0 | 0% | 1 | 34 | 100% | 21484 | 21517 | - | 21484 |
| 26_HKU360.crt.5 | AM295007.1 Streptococcus pyogenes Manfredo complete genome | 34 | 28976 | 63.9 | 4.00E-12 | 34 | 100% | 0 | 0% | 1 | 34 | 100% | 7465 | 7498 | + | 7465 |
| 26_HKU360.crt.6 | NOHIT | 34 | | | | | | | | | | | | | | |
| 26_HKU360.crt.7 | A07_1 | 33 | 31011 | 62.1 | 1.00E-11 | 33 | 100% | 0 | 0% | 1 | 33 | 100% | 12261 | 12293 | + | 12261 |
| 26_HKU360.crt.7 | BA000034.2 Streptococcus pyogenes SSI-1 DNA, complete genome | 33 | 31011 | 62.1 | 1.00E-11 | 33 | 100% | 0 | 0% | 1 | 33 | 100% | 18719 | 18751 | - | 18719 |
| 26_HKU360.crt.7 | CP008926.1 Streptococcus pyogenes strain ATCC 19615, complete genome | 33 | 31521 | 62.1 | 1.00E-11 | 33 | 100% | 0 | 0% | 1 | 33 | 100% | 18719 | 18751 | - | 18719 |
| 27_10.crt.1 | NOHIT | 30 | | | | | | | | | | | | | | |
| 27_10.crt.2 | NOHIT | 30 | | | | | | | | | | | | | | |
| 27_10.crt.3 | NOHIT | 30 | | | | | | | | | | | | | | |
| 27_10.crt.4 | NOHIT | 30 | | | | | | | | | | | | | | |
| 27_10.crt.5 | NOHIT | 30 | | | | | | | | | | | | | | |
| 27_10.crt.6 | NOHIT | 30 | | | | | | | | | | | | | | |
| 27_10.crt.7 | NOHIT | 30 | | | | | | | | | | | | | | |
| 27_10.crt.8 | NOHIT | 30 | | | | | | | | | | | | | | |
| 27_10.crt.9 | NOHIT | 30 | | | | | | | | | | | | | | |
| 27_10.crt.10 | NOHIT | 30 | | | | | | | | | | | | | | |
| 27_10.crt.11 | NOHIT | 0 | | | | | | | | | | | | | | |
| 27_10.crt.12 | NOHIT | 29 | | | | | | | | | | | | | | |
| 27_10.crt.13 | NOHIT | 31 | | | | | | | | | | | | | | |
| 27_10.crt.14 | A07_1 | 32 | 31011 | 60.2 | 5.00E-11 | 32 | 100% | 0 | 0% | 1 | 32 | 100% | 7277 | 7308 | + | 7277 |
| 27_10.crt.14 | BA000034.2 Streptococcus pyogenes SSI-1 DNA, complete genome | 32 | 31011 | 60.2 | 5.00E-11 | 32 | 100% | 0 | 0% | 1 | 32 | 100% | 23704 | 23735 | - | 23704 |
| 27_10.crt.14 | CP008926.1 Streptococcus pyogenes strain ATCC 19615, complete genome | 32 | 31521 | 60.2 | 5.00E-11 | 32 | 100% | 0 | 0% | 1 | 32 | 100% | 23705 | 23736 | - | 23705 |
| 28_MTB313.crt.1 | NOHIT | 29 | | | | | | | | | | | | | | |
| 28_MTB313.crt.2 | NOHIT | 31 | | | | | | | | | | | | | | |
| 28_MTB313.crt.3 | NOHIT | 29 | | | | | | | | | | | | | | |
| 28_MTB313.crt.4 | NOHIT | 0 | | | | | | | | | | | | | | |
| 28_MTB313.crt.5 | CP007561.1 Streptococcus pyogenes strain NGAS596, complete genome | 33 | 40047 | 62.1 | 1.00E-11 | 33 | 100% | 0 | 0% | 1 | 33 | 100% | 4156 | 4188 | - | 4156 |
| 28_MTB313.crt.5 | CP008695.1 Streptococcus pyogenes strain M23ND, complete genome | 33 | 37509 | 62.1 | 1.00E-11 | 33 | 100% | 0 | 0% | 1 | 33 | 100% | 34682 | 34714 | + | 34682 |
| 28_MTB313.crt.5 | AM295007.1 Streptococcus pyogenes Manfredo complete genome | 33 | 37666 | 62.1 | 1.00E-11 | 33 | 100% | 0 | 0% | 1 | 33 | 100% | 4183 | 4215 | - | 4183 |
| 28_MTB313.crt.6 | NOHIT | 34 | | | | | | | | | | | | | | |
| 28_MTB313.crt.7 | CP008695.1 Streptococcus pyogenes strain M23ND, complete genome | 33 | 37458 | 62.1 | 1.00E-11 | 33 | 100% | 0 | 0% | 1 | 33 | 100% | 21052 | 21084 | + | 21052 |
| 28_MTB313.crt.8 | A08_1 | 35 | 36269 | 60.2 | 6.00E-11 | 34 | 97% | 0 | 0% | 1 | 35 | 100% | 28369 | 28403 | - | 28369 |
| 28_MTB313.crt.8 | CP013838.1 Streptococcus pyogenes strain MGAS11027, complete genome | 35 | 51647 | 60.2 | 6.00E-11 | 34 | 97% | 0 | 0% | 1 | 35 | 100% | 13944 | 13978 | + | 13944 |
| 28_MTB313.crt.8 | CP007560.1 Streptococcus pyogenes strain NGAS743, complete genome | 35 | 38050 | 60.2 | 6.00E-11 | 34 | 97% | 0 | 0% | 1 | 35 | 100% | 27864 | 27898 | - | 27864 |
| 28_MTB313.crt.8 | CP007560.1 Streptococcus pyogenes strain NGAS743, complete genome | 35 | 39059 | 60.2 | 6.00E-11 | 34 | 97% | 0 | 0% | 1 | 35 | 100% | 14542 | 14576 | + | 14542 |
| 28_MTB313.crt.8 | AE009949.1 Streptococcus pyogenes MGAS8232, complete genome | 35 | 39804 | 60.2 | 6.00E-11 | 34 | 97% | 0 | 0% | 1 | 35 | 100% | 12058 | 12092 | + | 12058 |
| 28_MTB313.crt.8 | CP009612.1 Streptococcus pyogenes strain HKU360, complete genome | 35 | 38759 | 60.2 | 6.00E-11 | 34 | 97% | 0 | 0% | 1 | 35 | 100% | 26510 | 26544 | - | 26510 |
| 28_MTB313.crt.8 | CP008695.1 Streptococcus pyogenes strain M23ND, complete genome | 35 | 37458 | 60.2 | 6.00E-11 | 34 | 97% | 0 | 0% | 1 | 35 | 100% | 12800 | 12834 | + | 12800 |
| 28_MTB313.crt.8 | CP006366.1 Streptococcus pyogenes HSC5, complete genome | 35 | 39477 | 60.2 | 6.00E-11 | 34 | 97% | 0 | 0% | 1 | 35 | 100% | 24623 | 24657 | - | 24623 |
| 28_MTB313.crt.8 | AM295007.1 Streptococcus pyogenes Manfredo complete genome | 35 | 39447 | 60.2 | 6.00E-11 | 34 | 97% | 0 | 0% | 1 | 35 | 100% | 27341 | 27375 | - | 27341 |
| 29_MTB314.crt.1 | NOHIT | 30 | | | | | | | | | | | | | | |
| 29_MTB314.crt.2 | NOHIT | 30 | | | | | | | | | | | | | | |
| 29_MTB314.crt.3 | NOHIT | 30 | | | | | | | | | | | | | | |
| 29_MTB314.crt.4 | NOHIT | 0 | | | | | | | | | | | | | | |
| 29_MTB314.crt.5 | CP007561.1 Streptococcus pyogenes strain NGAS596, complete genome | 33 | 40047 | 62.1 | 1.00E-11 | 33 | 100% | 0 | 0% | 1 | 33 | 100% | 4156 | 4188 | - | 4156 |
| 29_MTB314.crt.5 | CP008695.1 Streptococcus pyogenes strain M23ND, complete genome | 33 | 37509 | 62.1 | 1.00E-11 | 33 | 100% | 0 | 0% | 1 | 33 | 100% | 34682 | 34714 | + | 34682 |
| 29_MTB314.crt.5 | AM295007.1 Streptococcus pyogenes Manfredo complete genome | 33 | 37666 | 62.1 | 1.00E-11 | 33 | 100% | 0 | 0% | 1 | 33 | 100% | 4183 | 4215 | - | 4183 |
| 29_MTB314.crt.6 | NOHIT | 33 | | | | | | | | | | | | | | |
| 29_MTB314.crt.7 | CP008695.1 Streptococcus pyogenes strain M23ND, complete genome | 33 | 37458 | 62.1 | 1.00E-11 | 33 | 100% | 0 | 0% | 1 | 33 | 100% | 21052 | 21084 | + | 21052 |
| 29_MTB314.crt.8 | A08_1 | 35 | 36269 | 60.2 | 6.00E-11 | 34 | 97% | 0 | 0% | 1 | 35 | 100% | 28369 | 28403 | - | 28369 |
| 29_MTB314.crt.8 | CP013838.1 Streptococcus pyogenes strain MGAS11027, complete genome | 35 | 51647 | 60.2 | 6.00E-11 | 34 | 97% | 0 | 0% | 1 | 35 | 100% | 13944 | 13978 | + | 13944 |
| 29_MTB314.crt.8 | CP007560.1 Streptococcus pyogenes strain NGAS743, complete genome | 35 | 38050 | 60.2 | 6.00E-11 | 34 | 97% | 0 | 0% | 1 | 35 | 100% | 27864 | 27898 | - | 27864 |
| 29_MTB314.crt.8 | CP007560.1 Streptococcus pyogenes strain NGAS743, complete genome | 35 | 39059 | 60.2 | 6.00E-11 | 34 | 97% | 0 | 0% | 1 | 35 | 100% | 14542 | 14576 | + | 14542 |
| 29_MTB314.crt.8 | AE009949.1 Streptococcus pyogenes MGAS8232, complete genome | 35 | 39804 | 60.2 | 6.00E-11 | 34 | 97% | 0 | 0% | 1 | 35 | 100% | 12058 | 12092 | + | 12058 |
| 29_MTB314.crt.8 | CP009612.1 Streptococcus pyogenes strain HKU360, complete genome | 35 | 38759 | 60.2 | 6.00E-11 | 34 | 97% | 0 | 0% | 1 | 35 | 100% | 26510 | 26544 | - | 26510 |
| 29_MTB314.crt.8 | CP008695.1 Streptococcus pyogenes strain M23ND, complete genome | 35 | 37458 | 60.2 | 6.00E-11 | 34 | 97% | 0 | 0% | 1 | 35 | 100% | 12800 | 12834 | + | 12800 |
| 29_MTB314.crt.8 | CP006366.1 Streptococcus pyogenes HSC5, complete genome | 35 | 39477 | 60.2 | 6.00E-11 | 34 | 97% | 0 | 0% | 1 | 35 | 100% | 24623 | 24657 | - | 24623 |
| 29_MTB314.crt.8 | AM295007.1 Streptococcus pyogenes Manfredo complete genome | 35 | 39447 | 60.2 | 6.00E-11 | 34 | 97% | 0 | 0% | 1 | 35 | 100% | 27341 | 27375 | - | 27341 |
| 31_AP1.crt.1 | A08_1 | 35 | 36269 | 60.2 | 6.00E-11 | 34 | 97% | 0 | 0% | 1 | 35 | 100% | 28369 | 28403 | + | 28369 |
| 31_AP1.crt.1 | CP013838.1 Streptococcus pyogenes strain MGAS11027, complete genome | 35 | 51647 | 60.2 | 6.00E-11 | 34 | 97% | 0 | 0% | 1 | 35 | 100% | 13944 | 13978 | - | 13944 |
| 31_AP1.crt.1 | CP007560.1 Streptococcus pyogenes strain NGAS743, complete genome | 35 | 38050 | 60.2 | 6.00E-11 | 34 | 97% | 0 | 0% | 1 | 35 | 100% | 27864 | 27898 | + | 27864 |
| 31_AP1.crt.1 | CP007560.1 Streptococcus pyogenes strain NGAS743, complete genome | 35 | 39059 | 60.2 | 6.00E-11 | 34 | 97% | 0 | 0% | 1 | 35 | 100% | 14542 | 14576 | - | 14542 |
| 31_AP1.crt.1 | AE009949.1 Streptococcus pyogenes MGAS8232, complete genome | 35 | 39804 | 60.2 | 6.00E-11 | 34 | 97% | 0 | 0% | 1 | 35 | 100% | 12058 | 12092 | - | 12058 |
| 31_AP1.crt.1 | CP009612.1 Streptococcus pyogenes strain HKU360, complete genome | 35 | 38759 | 60.2 | 6.00E-11 | 34 | 97% | 0 | 0% | 1 | 35 | 100% | 26510 | 26544 | + | 26510 |
| 31_AP1.crt.1 | CP008695.1 Streptococcus pyogenes strain M23ND, complete genome | 35 | 37458 | 60.2 | 6.00E-11 | 34 | 97% | 0 | 0% | 1 | 35 | 100% | 12800 | 12834 | - | 12800 |
| 31_AP1.crt.1 | CP006366.1 Streptococcus pyogenes HSC5, complete genome | 35 | 39477 | 60.2 | 6.00E-11 | 34 | 97% | 0 | 0% | 1 | 35 | 100% | 24623 | 24657 | + | 24623 |
| 31_AP1.crt.1 | AM295007.1 Streptococcus pyogenes Manfredo complete genome | 35 | 39447 | 60.2 | 6.00E-11 | 34 | 97% | 0 | 0% | 1 | 35 | 100% | 27341 | 27375 | + | 27341 |
| 31_AP1.crt.2 | CP008695.1 Streptococcus pyogenes strain M23ND, complete genome | 33 | 37458 | 62.1 | 1.00E-11 | 33 | 100% | 0 | 0% | 1 | 33 | 100% | 21052 | 21084 | - | 21052 |
| 31_AP1.crt.3 | NOHIT | 34 | | | | | | | | | | | | | | |
| 31_AP1.crt.4 | CP007561.1 Streptococcus pyogenes strain NGAS596, complete genome | 33 | 40047 | 62.1 | 1.00E-11 | 33 | 100% | 0 | 0% | 1 | 33 | 100% | 4156 | 4188 | + | 4156 |
| 31_AP1.crt.4 | CP008695.1 Streptococcus pyogenes strain M23ND, complete genome | 33 | 37509 | 62.1 | 1.00E-11 | 33 | 100% | 0 | 0% | 1 | 33 | 100% | 34682 | 34714 | - | 34682 |
| 31_AP1.crt.4 | AM295007.1 Streptococcus pyogenes Manfredo complete genome | 33 | 37666 | 62.1 | 1.00E-11 | 33 | 100% | 0 | 0% | 1 | 33 | 100% | 4183 | 4215 | + | 4183 |
| 35_NGAS596.crt.1 | CP000003.1 Streptococcus pyogenes MGAS10394, complete genome | 32 | 45489 | 60.2 | 5.00E-11 | 32 | 100% | 0 | 0% | 1 | 32 | 100% | 16996 | 17027 | - | 16996 |
| 35_NGAS596.crt.1 | CP011415.1 Streptococcus pyogenes strain D471, complete genome | 32 | 39110 | 60.2 | 5.00E-11 | 32 | 100% | 0 | 0% | 1 | 32 | 100% | 36057 | 36088 | - | 36057 |
| 35_NGAS596.crt.1 | CP011414.1 Streptococcus pyogenes JRS4, complete genome | 32 | 39110 | 60.2 | 5.00E-11 | 32 | 100% | 0 | 0% | 1 | 32 | 100% | 36057 | 36088 | - | 36057 |
| 35_NGAS596.crt.1 | AP012335.1 Streptococcus pyogenes JRS4 DNA, complete genome | 32 | 39111 | 60.2 | 5.00E-11 | 32 | 100% | 0 | 0% | 1 | 32 | 100% | 36058 | 36089 | - | 36058 |
| 35_NGAS596.crt.1 | CP006366.1 Streptococcus pyogenes HSC5, complete genome | 32 | 39477 | 60.2 | 5.00E-11 | 32 | 100% | 0 | 0% | 1 | 32 | 100% | 36425 | 36456 | - | 36425 |

| | | | | | | | |
|---|---|---|---|---|---|---|---|
| 35_NGAS596.crt.2 | A08_1 | 33 | 36269 | 62.1 | 1.00E-11 | 33 | [illegible] |
| 35_NGAS596.crt.2 | CP010449.1 Streptococcus pyogenes strain NGAS322, complete genome | 33 | 38059 | 62.1 | 1.00E-11 | 33 | [illegible] |
| 35_NGAS596.crt.2 | CP007560.1 Streptococcus pyogenes strain NGAS743, complete genome | 33 | 38050 | 62.1 | 1.00E-11 | 33 | [illegible] |
| 35_NGAS596.crt.2 | AE009949.1 Streptococcus pyogenes MGAS8232, complete genome | 33 | 39804 | 62.1 | 1.00E-11 | 33 | [illegible] |
| 35_NGAS596.crt.2 | CP008695.1 Streptococcus pyogenes strain M23ND, complete genome | 33 | 27969 | 62.1 | 1.00E-11 | 33 | [illegible] |
| 35_NGAS596.crt.2 | AE004092.2 Streptococcus pyogenes M1 GAS, complete genome | 33 | 40272 | 62.1 | 1.00E-11 | 33 | [illegible] |
| 35_NGAS596.crt.2 | CP006366.1 Streptococcus pyogenes HSC5, complete genome | 33 | 39477 | 62.1 | 1.00E-11 | 33 | [illegible] |
| 35_NGAS596.crt.2 | AM295007.1 Streptococcus pyogenes Manfredo complete genome | 33 | 39447 | 62.1 | 1.00E-11 | 33 | [illegible] |
| 35_NGAS596.crt.2 | CP000017.2 Streptococcus pyogenes MGAS5005, complete genome | 33 | 29805 | 60.2 | 5.00E-11 | 32 | [illegible] |
| 35_NGAS596.crt.2 | CP012045.1 Streptococcus pyogenes strain HKU488, complete genome | 33 | 39642 | 60.2 | 5.00E-11 | 32 | [illegible] |
| 35_NGAS596.crt.2 | CP012045.1 Streptococcus pyogenes strain HKU488, complete genome | 33 | 29804 | 60.2 | 5.00E-11 | 32 | [illegible] |
| 35_NGAS596.crt.2 | CP008776.1 Streptococcus pyogenes strain 5448, complete genome | 33 | 29805 | 60.2 | 5.00E-11 | 32 | [illegible] |
| 35_NGAS596.crt.2 | CP007537.1 Streptococcus pyogenes strain AP1, complete genome | 33 | 29748 | 60.2 | 5.00E-11 | 32 | [illegible] |
| 35_NGAS596.crt.2 | AP014585.1 Streptococcus pyogenes DNA, complete genome, strain: MTB314 | 33 | 18078 | 60.2 | 5.00E-11 | 32 | [illegible] |
| 35_NGAS596.crt.2 | CP009612.1 Streptococcus pyogenes strain HKU360, complete genome | 33 | 39642 | 60.2 | 5.00E-11 | 32 | [illegible] |
| 35_NGAS596.crt.2 | AE004092.2 Streptococcus pyogenes M1 GAS, complete genome | 33 | 29920 | 60.2 | 5.00E-11 | 32 | [illegible] |
| 35_NGAS596.crt.2 | CP006366.1 Streptococcus pyogenes HSC5, complete genome | 33 | 32692 | 60.2 | 5.00E-11 | 32 | [illegible] |
| 35_NGAS596.crt.2 | CP003901.1 Streptococcus pyogenes A20, complete genome | 33 | 29805 | 60.2 | 5.00E-11 | 32 | [illegible] |
| 35_NGAS596.crt.2 | AP012491.2 Streptococcus pyogenes M1 476 DNA, complete genome | 33 | 29805 | 60.2 | 5.00E-11 | 32 | [illegible] |
| 35_NGAS596.crt.3 | NOHIT | 34 | | | | | |
| 35_NGAS596.crt.4 | CP008695.1 Streptococcus pyogenes strain M23ND, complete genome | 33 | 37458 | 62.1 | 1.00E-11 | 33 | [illegible] |
| 35_NGAS596.crt.5 | A08_1 | 33 | 31416 | 62.1 | 1.00E-11 | 33 | [illegible] |
| 35_NGAS596.crt.5 | A07_1 | 33 | 31011 | 62.1 | 1.00E-11 | 33 | [illegible] |
| 35_NGAS596.crt.5 | BA000034.2 Streptococcus pyogenes SSI-1 DNA, complete genome | 33 | 31011 | 62.1 | 1.00E-11 | 33 | [illegible] |
| 35_NGAS596.crt.5 | CP013838.1 Streptococcus pyogenes strain MGAS11027, complete genome | 33 | 35828 | 62.1 | 1.00E-11 | 33 | [illegible] |
| 35_NGAS596.crt.5 | CP013672.1 Streptococcus pyogenes strain AP53, complete genome | 33 | 24636 | 62.1 | 1.00E-11 | 33 | [illegible] |
| 35_NGAS596.crt.5 | CP014138.1 Streptococcus pyogenes strain MEW427, complete genome | 33 | 22381 | 62.1 | 1.00E-11 | 33 | [illegible] |
| 35_NGAS596.crt.5 | CP010449.1 Streptococcus pyogenes strain NGAS322, complete genome | 33 | 34489 | 62.1 | 1.00E-11 | 33 | [illegible] |
| 35_NGAS596.crt.5 | CP012045.1 Streptococcus pyogenes strain HKU488, complete genome | 33 | 39642 | 62.1 | 1.00E-11 | 33 | [illegible] |
| 35_NGAS596.crt.5 | CP007560.1 Streptococcus pyogenes strain NGAS743, complete genome | 33 | 39618 | 62.1 | 1.00E-11 | 33 | [illegible] |
| 35_NGAS596.crt.5 | CP007560.1 Streptococcus pyogenes strain NGAS743, complete genome | 33 | 33384 | 62.1 | 1.00E-11 | 33 | [illegible] |
| 35_NGAS596.crt.5 | CP011415.1 Streptococcus pyogenes strain D471, complete genome | 33 | 39110 | 62.1 | 1.00E-11 | 33 | [illegible] |
| 35_NGAS596.crt.5 | CP011414.1 Streptococcus pyogenes JRS4, complete genome | 33 | 39110 | 62.1 | 1.00E-11 | 33 | [illegible] |
| 35_NGAS596.crt.5 | AP012335.1 Streptococcus pyogenes JRS4 DNA, complete genome | 33 | 39111 | 62.1 | 1.00E-11 | 33 | [illegible] |
| 35_NGAS596.crt.5 | CP009612.1 Streptococcus pyogenes strain HKU360, complete genome | 33 | 39642 | 62.1 | 1.00E-11 | 33 | [illegible] |
| 35_NGAS596.crt.5 | CP003068.1 Streptococcus pyogenes Alab49, complete genome | 33 | 24636 | 62.1 | 1.00E-11 | 33 | [illegible] |
| 35_NGAS596.crt.5 | AM295007.1 Streptococcus pyogenes Manfredo complete genome | 33 | 37666 | 62.1 | 1.00E-11 | 33 | [illegible] |

[illegible]

| | | | | | | | | | | | | | | | |
|---|---|---|---|---|---|---|---|---|---|---|---|---|---|---|---|
| 49_MGAS27061.crt.16 | GAS003 | 33 | 37328 | 62.1 | 1.00E-11 | 33 | 100% | 0 | 0% | 1 | 33 | 100% | 2031 | 2063 - | 2031 |
| 49_MGAS27061.crt.16 | CP009612.1 Streptococcus pyogenes strain HKU360, complete genome | 33 | 37343 | 62.1 | 1.00E-11 | 33 | 100% | 0 | 0% | 1 | 33 | 100% | 2031 | 2063 - | 2031 |
| 49_MGAS27061.crt.17 | NOHIT | 32 | | | | | | | | | | | | | |
| 49_MGAS27061.crt.18 | CP000260.1 Streptococcus pyogenes MGAS10270, complete genome | 33 | 39966 | 62.1 | 1.00E-11 | 33 | 100% | 0 | 0% | 1 | 33 | 100% | 10775 | 10807 + | 10775 |
| 49_MGAS27061.crt.18 | CP000056.1 Streptococcus pyogenes MGAS6180, complete genome | 33 | 36896 | 62.1 | 1.00E-11 | 33 | 100% | 0 | 0% | 1 | 33 | 100% | 11261 | 11293 + | 11261 |
| 49_MGAS27061.crt.18 | CP000003.1 Streptococcus pyogenes MGAS10394, complete genome | 33 | 15429 | 62.1 | 1.00E-11 | 33 | 100% | 0 | 0% | 1 | 33 | 100% | 3489 | 3521 + | 3489 |
| 49_MGAS27061.crt.18 | A09_1 | 33 | 30199 | 62.1 | 1.00E-11 | 33 | 100% | 0 | 0% | 1 | 33 | 100% | 26679 | 26711 - | 26679 |
| 49_MGAS27061.crt.18 | A08_1 | 33 | 32841 | 62.1 | 1.00E-11 | 33 | 100% | 0 | 0% | 1 | 33 | 100% | 29159 | 29191 - | 29159 |
| 49_MGAS27061.crt.18 | A07_1 | 33 | 39948 | 62.1 | 1.00E-11 | 33 | 100% | 0 | 0% | 1 | 33 | 100% | 10742 | 10774 + | 10742 |
| 49_MGAS27061.crt.18 | A06_1 | 33 | 30200 | 62.1 | 1.00E-11 | 33 | 100% | 0 | 0% | 1 | 33 | 100% | 26680 | 26712 - | 26680 |
| 49_MGAS27061.crt.18 | GAS004 | 33 | 34136 | 62.1 | 1.00E-11 | 33 | 100% | 0 | 0% | 1 | 33 | 100% | 30676 | 30708 - | 30676 |
| 49_MGAS27061.crt.18 | BA000034.2 Streptococcus pyogenes SSI-1 DNA, complete genome | 33 | 39933 | 62.1 | 1.00E-11 | 33 | 100% | 0 | 0% | 1 | 33 | 100% | 29160 | 29192 - | 29160 |
| 49_MGAS27061.crt.18 | CP013838.1 Streptococcus pyogenes strain MGAS11027, complete genome | 33 | 35828 | 62.1 | 1.00E-11 | 33 | 100% | 0 | 0% | 1 | 33 | 100% | 28790 | 28822 - | 28790 |
| 49_MGAS27061.crt.18 | CP013838.1 Streptococcus pyogenes strain MGAS11027, complete genome | 33 | 51647 | 62.1 | 1.00E-11 | 33 | 100% | 0 | 0% | 1 | 33 | 100% | 30262 | 30294 - | 30262 |
| 49_MGAS27061.crt.18 | CP013672.1 Streptococcus pyogenes strain AP53, complete genome | 33 | 29027 | 62.1 | 1.00E-11 | 33 | 100% | 0 | 0% | 1 | 33 | 100% | 3381 | 3413 + | 3381 |
| 49_MGAS27061.crt.18 | AE014074.1 Streptococcus pyogenes MGAS315, complete genome | 33 | 39933 | 62.1 | 1.00E-11 | 33 | 100% | 0 | 0% | 1 | 33 | 100% | 10742 | 10774 + | 10742 |
| 49_MGAS27061.crt.18 | CP011068.1 Streptococcus pyogenes strain STAB10015, complete genome | 33 | 36896 | 62.1 | 1.00E-11 | 33 | 100% | 0 | 0% | 1 | 33 | 100% | 11261 | 11293 + | 11261 |
| 49_MGAS27061.crt.18 | CP011535.2 Streptococcus pyogenes strain M28PF1, complete genome | 33 | 32154 | 62.1 | 1.00E-11 | 33 | 100% | 0 | 0% | 1 | 33 | 100% | 20863 | 20895 - | 20863 |
| 49_MGAS27061.crt.18 | CP007560.1 Streptococcus pyogenes strain NGAS743, complete genome | 33 | 38050 | 62.1 | 1.00E-11 | 33 | 100% | 0 | 0% | 1 | 33 | 100% | 9472 | 9504 + | 9472 |
| 49_MGAS27061.crt.18 | CP007560.1 Streptococcus pyogenes strain NGAS743, complete genome | 33 | 39618 | 62.1 | 1.00E-11 | 33 | 100% | 0 | 0% | 1 | 33 | 100% | 7387 | 7419 + | 7387 |
| 49_MGAS27061.crt.18 | CP011415.1 Streptococcus pyogenes strain D471, complete genome | 33 | 30199 | 62.1 | 1.00E-11 | 33 | 100% | 0 | 0% | 1 | 33 | 100% | 3489 | 3521 + | 3489 |
| 49_MGAS27061.crt.18 | CP011414.1 Streptococcus pyogenes JRS4, complete genome | 33 | 30199 | 62.1 | 1.00E-11 | 33 | 100% | 0 | 0% | 1 | 33 | 100% | 3489 | 3521 + | 3489 |
| 49_MGAS27061.crt.18 | CP007537.1 Streptococcus pyogenes strain AP1, complete genome | 33 | 29027 | 62.1 | 1.00E-11 | 33 | 100% | 0 | 0% | 1 | 33 | 100% | 25615 | 25647 - | 25615 |
| 49_MGAS27061.crt.18 | AP012335.1 Streptococcus pyogenes JRS4 DNA, complete genome | 33 | 30198 | 62.1 | 1.00E-11 | 33 | 100% | 0 | 0% | 1 | 33 | 100% | 3489 | 3521 + | 3489 |
| 49_MGAS27061.crt.18 | AE009949.1 Streptococcus pyogenes MGAS8232, complete genome | 33 | 30227 | 62.1 | 1.00E-11 | 33 | 100% | 0 | 0% | 1 | 33 | 100% | 3484 | 3516 + | 3484 |
| 49_MGAS27061.crt.18 | CP007041.1 Streptococcus pyogenes STAB902, complete genome | 33 | 32577 | 62.1 | 1.00E-11 | 33 | 100% | 0 | 0% | 1 | 33 | 100% | 29161 | 29193 - | 29161 |
| 49_MGAS27061.crt.18 | AE004092.2 Streptococcus pyogenes M1 GAS, complete genome | 33 | 39657 | 62.1 | 1.00E-11 | 33 | 100% | 0 | 0% | 1 | 33 | 100% | 30676 | 30708 - | 30676 |
| 49_MGAS27061.crt.18 | CP006366.1 Streptococcus pyogenes HSC5, complete genome | 33 | 39477 | 62.1 | 1.00E-11 | 33 | 100% | 0 | 0% | 1 | 33 | 100% | 6858 | 6890 + | 6858 |
| 49_MGAS27061.crt.18 | CP003068.1 Streptococcus pyogenes Alab49, complete genome | 33 | 29027 | 62.1 | 1.00E-11 | 33 | 100% | 0 | 0% | 1 | 33 | 100% | 3381 | 3413 + | 3381 |
| 49_MGAS27061.crt.18 | AM295007.1 Streptococcus pyogenes Manfredo complete genome | 33 | 28976 | 62.1 | 1.00E-11 | 33 | 100% | 0 | 0% | 1 | 33 | 100% | 25564 | 25596 - | 25564 |
| 49_MGAS27061.crt.18 | CP000261.1 Streptococcus pyogenes MGAS2096, complete genome | 33 | 40056 | 62.1 | 1.00E-11 | 33 | 100% | 0 | 0% | 1 | 33 | 100% | 30676 | 30708 - | 30676 |
| 51_STAB09014.crt.1 | NOHIT | 30 | | | | | | | | | | | | | |
| 51_STAB09014.crt.2 | NOHIT | 30 | | | | | | | | | | | | | |
| 51_STAB09014.crt.3 | NOHIT | 30 | | | | | | | | | | | | | |
| 51_STAB09014.crt.4 | NOHIT | 30 | | | | | | | | | | | | | |
| 52_ATCC14918.crt.1 | NOHIT | 32 | | | | | | | | | | | | | |
| 52_ATCC14918.crt.2 | CP000056.1 Streptococcus pyogenes MGAS6180, complete genome | 34 | 35292 | 63.9 | 4.00E-12 | 34 | 100% | 0 | 0% | 1 | 34 | 100% | 29395 | 29428 - | 29395 |
| 52_ATCC14918.crt.2 | A09_1 | 34 | 30199 | 63.9 | 4.00E-12 | 34 | 100% | 0 | 0% | 1 | 34 | 100% | 8103 | 8136 + | 8103 |
| 52_ATCC14918.crt.2 | A07_1 | 34 | 37110 | 63.9 | 4.00E-12 | 34 | 100% | 0 | 0% | 1 | 34 | 100% | 23711 | 23744 - | 23711 |
| 52_ATCC14918.crt.2 | A06_1 | 34 | 30200 | 63.9 | 4.00E-12 | 34 | 100% | 0 | 0% | 1 | 34 | 100% | 8103 | 8136 + | 8103 |
| 52_ATCC14918.crt.2 | CP011069.1 Streptococcus pyogenes strain STAB09014, complete genome | 34 | 35292 | 63.9 | 4.00E-12 | 34 | 100% | 0 | 0% | 1 | 34 | 100% | 29395 | 29428 - | 29395 |
| 52_ATCC14918.crt.2 | BA000034.2 Streptococcus pyogenes SSI-1 DNA, complete genome | 34 | 37117 | 63.9 | 4.00E-12 | 34 | 100% | 0 | 0% | 1 | 34 | 100% | 13367 | 13400 + | 13367 |
| 52_ATCC14918.crt.2 | CP013672.1 Streptococcus pyogenes strain AP53, complete genome | 34 | 29027 | 63.9 | 4.00E-12 | 34 | 100% | 0 | 0% | 1 | 34 | 100% | 21484 | 21517 - | 21484 |
| 52_ATCC14918.crt.2 | CP014139.1 Streptococcus pyogenes strain MEW123, complete genome | 34 | 34850 | 63.9 | 4.00E-12 | 34 | 100% | 0 | 0% | 1 | 34 | 100% | 15233 | 15266 + | 15233 |
| 52_ATCC14918.crt.2 | CP010450.1 Streptococcus pyogenes strain NGAS638, complete genome | 34 | 35488 | 63.9 | 4.00E-12 | 34 | 100% | 0 | 0% | 1 | 34 | 100% | 29071 | 29104 - | 29071 |
| 52_ATCC14918.crt.2 | AE014074.1 Streptococcus pyogenes MGAS315, complete genome | 34 | 36875 | 63.9 | 4.00E-12 | 34 | 100% | 0 | 0% | 1 | 34 | 100% | 23487 | 23520 - | 23487 |
| 52_ATCC14918.crt.2 | CP011068.1 Streptococcus pyogenes strain STAB10015, complete genome | 34 | 35292 | 63.9 | 4.00E-12 | 34 | 100% | 0 | 0% | 1 | 34 | 100% | 29395 | 29428 - | 29395 |
| 52_ATCC14918.crt.2 | CP011535.2 Streptococcus pyogenes strain M28PF1, complete genome | 34 | 35292 | 63.9 | 4.00E-12 | 34 | 100% | 0 | 0% | 1 | 34 | 100% | 5865 | 5898 + | 5865 |
| 52_ATCC14918.crt.2 | CP011415.1 Streptococcus pyogenes strain D471, complete genome | 34 | 30199 | 63.9 | 4.00E-12 | 34 | 100% | 0 | 0% | 1 | 34 | 100% | 22064 | 22097 - | 22064 |
| 52_ATCC14918.crt.2 | CP011414.1 Streptococcus pyogenes JRS4, complete genome | 34 | 30199 | 63.9 | 4.00E-12 | 34 | 100% | 0 | 0% | 1 | 34 | 100% | 22064 | 22097 - | 22064 |
| 52_ATCC14918.crt.2 | CP007537.1 Streptococcus pyogenes strain AP1, complete genome | 34 | 29027 | 63.9 | 4.00E-12 | 34 | 100% | 0 | 0% | 1 | 34 | 100% | 7511 | 7544 + | 7511 |
| 52_ATCC14918.crt.2 | AP012335.1 Streptococcus pyogenes JRS4 DNA, complete genome | 34 | 30198 | 63.9 | 4.00E-12 | 34 | 100% | 0 | 0% | 1 | 34 | 100% | 22064 | 22097 - | 22064 |
| 52_ATCC14918.crt.2 | AE009949.1 Streptococcus pyogenes MGAS8232, complete genome | 34 | 30227 | 63.9 | 4.00E-12 | 34 | 100% | 0 | 0% | 1 | 34 | 100% | 22060 | 22093 - | 22060 |
| 52_ATCC14918.crt.2 | CP007041.1 Streptococcus pyogenes STAB902, complete genome | 34 | 36899 | 63.9 | 4.00E-12 | 34 | 100% | 0 | 0% | 1 | 34 | 100% | 13366 | 13399 + | 13366 |
| 52_ATCC14918.crt.2 | CP006366.1 Streptococcus pyogenes HSC5, complete genome | 34 | 37520 | 63.9 | 4.00E-12 | 34 | 100% | 0 | 0% | 1 | 34 | 100% | 31114 | 31147 - | 31114 |
| 52_ATCC14918.crt.2 | CP003068.1 Streptococcus pyogenes Alab49, complete genome | 34 | 29027 | 63.9 | 4.00E-12 | 34 | 100% | 0 | 0% | 1 | 34 | 100% | 21484 | 21517 - | 21484 |
| 52_ATCC14918.crt.2 | AM295007.1 Streptococcus pyogenes Manfredo complete genome | 34 | 28976 | 63.9 | 4.00E-12 | 34 | 100% | 0 | 0% | 1 | 34 | 100% | 7465 | 7498 + | 7465 |
| 52_ATCC14918.crt.3 | A07_1 | 34 | 14000 | 63.9 | 4.00E-12 | 34 | 100% | 0 | 0% | 1 | 34 | 100% | 3933 | 3966 - | 3933 |
| 52_ATCC14918.crt.3 | BA000034.2 Streptococcus pyogenes SSI-1 DNA, complete genome | 34 | 14000 | 63.9 | 4.00E-12 | 34 | 100% | 0 | 0% | 1 | 34 | 100% | 10035 | 10068 + | 10035 |
| 52_ATCC14918.crt.3 | CP010450.1 Streptococcus pyogenes strain NGAS638, complete genome | 34 | 35488 | 63.9 | 4.00E-12 | 34 | 100% | 0 | 0% | 1 | 34 | 100% | 27731 | 27764 - | 27731 |
| 52_ATCC14918.crt.3 | AE014074.1 Streptococcus pyogenes MGAS315, complete genome | 34 | 14000 | 63.9 | 4.00E-12 | 34 | 100% | 0 | 0% | 1 | 34 | 100% | 3933 | 3966 - | 3933 |
| 52_ATCC14918.crt.3 | CP008695.1 Streptococcus pyogenes strain M23ND, complete genome | 34 | 37458 | 63.9 | 4.00E-12 | 34 | 100% | 0 | 0% | 1 | 34 | 100% | 13553 | 13586 + | 13553 |
| 52_ATCC14918.crt.3 | CP007041.1 Streptococcus pyogenes STAB902, complete genome | 34 | 14001 | 63.9 | 4.00E-12 | 34 | 100% | 0 | 0% | 1 | 34 | 100% | 10036 | 10069 + | 10036 |
| 52_ATCC14918.crt.3 | CP003068.1 Streptococcus pyogenes Alab49, complete genome | 34 | 35543 | 63.9 | 4.00E-12 | 34 | 100% | 0 | 0% | 1 | 34 | 100% | 30353 | 30386 - | 30353 |
| 52_ATCC14918.crt.4 | CP010449.1 Streptococcus pyogenes strain NGAS322, complete genome | 34 | 38059 | 63.9 | 4.00E-12 | 34 | 100% | 0 | 0% | 1 | 34 | 100% | 36361 | 36394 + | 36361 |
| 52_ATCC14918.crt.4 | CP007560.1 Streptococcus pyogenes strain NGAS743, complete genome | 34 | 33384 | 63.9 | 4.00E-12 | 34 | 100% | 0 | 0% | 1 | 34 | 100% | 1666 | 1699 - | 1666 |
| 52_ATCC14918.crt.4 | AE009949.1 Streptococcus pyogenes MGAS8232, complete genome | 34 | 39804 | 63.9 | 4.00E-12 | 34 | 100% | 0 | 0% | 1 | 34 | 100% | 1666 | 1699 - | 1666 |
| 52_ATCC14918.crt.4 | AM295007.1 Streptococcus pyogenes Manfredo complete genome | 34 | 39447 | 63.9 | 4.00E-12 | 34 | 100% | 0 | 0% | 1 | 34 | 100% | 37749 | 37782 + | 37749 |
| 52_ATCC14918.crt.5 | NOHIT | 34 | | | | | | | | | | | | | |
| 52_ATCC14918.crt.6 | CP000017.2 Streptococcus pyogenes MGAS5005, complete genome | 35 | 29805 | 65.8 | 1.00E-12 | 35 | 100% | 0 | 0% | 1 | 35 | 100% | 28036 | 28070 - | 28036 |
| 52_ATCC14918.crt.6 | CP012045.1 Streptococcus pyogenes strain HKU488, complete genome | 35 | 29804 | 65.8 | 1.00E-12 | 35 | 100% | 0 | 0% | 1 | 35 | 100% | 28035 | 28069 - | 28035 |
| 52_ATCC14918.crt.6 | CP008776.1 Streptococcus pyogenes strain 5448, complete genome | 35 | 29805 | 65.8 | 1.00E-12 | 35 | 100% | 0 | 0% | 1 | 35 | 100% | 1736 | 1770 + | 1736 |
| 52_ATCC14918.crt.6 | AP014585.1 Streptococcus pyogenes DNA, complete genome, strain: MTB314 | 35 | 18078 | 65.8 | 1.00E-12 | 35 | 100% | 0 | 0% | 1 | 35 | 100% | 16309 | 16343 - | 16309 |
| 52_ATCC14918.crt.6 | CP006366.1 Streptococcus pyogenes HSC5, complete genome | 35 | 32692 | 65.8 | 1.00E-12 | 35 | 100% | 0 | 0% | 1 | 35 | 100% | 30923 | 30957 - | 30923 |
| 52_ATCC14918.crt.6 | CP003901.1 Streptococcus pyogenes A20, complete genome | 35 | 29805 | 65.8 | 1.00E-12 | 35 | 100% | 0 | 0% | 1 | 35 | 100% | 28036 | 28070 - | 28036 |
| 52_ATCC14918.crt.6 | AP012491.2 Streptococcus pyogenes M1 476 DNA, complete genome | 35 | 29805 | 65.8 | 1.00E-12 | 35 | 100% | 0 | 0% | 1 | 35 | 100% | 28036 | 28070 - | 28036 |
| 52_ATCC14918.crt.7 | A07_1 | 35 | 31011 | 65.8 | 1.00E-12 | 35 | 100% | 0 | 0% | 1 | 35 | 100% | 8161 | 8195 + | 8161 |
| 52_ATCC14918.crt.7 | BA000034.2 Streptococcus pyogenes SSI-1 DNA, complete genome | 35 | 31011 | 65.8 | 1.00E-12 | 35 | 100% | 0 | 0% | 1 | 35 | 100% | 22817 | 22851 - | 22817 |
| 52_ATCC14918.crt.7 | CP008926.1 Streptococcus pyogenes strain ATCC 19615, complete genome | 35 | 31521 | 65.8 | 1.00E-12 | 35 | 100% | 0 | 0% | 1 | 35 | 100% | 22817 | 22851 - | 22817 |
| 52_ATCC14918.crt.8 | A07_1 | 33 | 31011 | 62.1 | 1.00E-11 | 33 | 100% | 0 | 0% | 1 | 33 | 100% | 12261 | 12293 + | 12261 |
| 52_ATCC14918.crt.8 | BA000034.2 Streptococcus pyogenes SSI-1 DNA, complete genome | 33 | 31011 | 62.1 | 1.00E-11 | 33 | 100% | 0 | 0% | 1 | 33 | 100% | 18719 | 18751 - | 18719 |
| 52_ATCC14918.crt.8 | CP008926.1 Streptococcus pyogenes strain ATCC 19615, complete genome | 33 | 31521 | 62.1 | 1.00E-11 | 33 | 100% | 0 | 0% | 1 | 33 | 100% | 18719 | 18751 - | 18719 |
| 53_ATCC14919.crt.1 | NOHIT | 38 | | | | | | | | | | | | | |
| 53_ATCC14919.crt.2 | NOHIT | 37 | | | | | | | | | | | | | |
| 53_ATCC14919.crt.3 | NOHIT | 0 | | | | | | | | | | | | | |
| 53_ATCC14919.crt.4 | NOHIT | 32 | | | | | | | | | | | | | |
| 53_ATCC14919.crt.5 | CP000056.1 Streptococcus pyogenes MGAS6180, complete genome | 34 | 35292 | 63.9 | 4.00E-12 | 34 | 100% | 0 | 0% | 1 | 34 | 100% | 29395 | 29428 - | 29395 |
| 53_ATCC14919.crt.5 | A09_1 | 34 | 30199 | 63.9 | 4.00E-12 | 34 | 100% | 0 | 0% | 1 | 34 | 100% | 8103 | 8136 + | 8103 |
| 53_ATCC14919.crt.5 | A07_1 | 34 | 37110 | 63.9 | 4.00E-12 | 34 | 100% | 0 | 0% | 1 | 34 | 100% | 23711 | 23744 - | 23711 |
| 53_ATCC14919.crt.5 | A06_1 | 34 | 30200 | 63.9 | 4.00E-12 | 34 | 100% | 0 | 0% | 1 | 34 | 100% | 8103 | 8136 + | 8103 |
| 53_ATCC14919.crt.5 | CP011069.1 Streptococcus pyogenes strain STAB09014, complete genome | 34 | 35292 | 63.9 | 4.00E-12 | 34 | 100% | 0 | 0% | 1 | 34 | 100% | 29395 | 29428 - | 29395 |
| 53_ATCC14919.crt.5 | BA000034.2 Streptococcus pyogenes SSI-1 DNA, complete genome | 34 | 37117 | 63.9 | 4.00E-12 | 34 | 100% | 0 | 0% | 1 | 34 | 100% | 13367 | 13400 + | 13367 |
| 53_ATCC14919.crt.5 | CP013672.1 Streptococcus pyogenes strain AP53, complete genome | 34 | 29027 | 63.9 | 4.00E-12 | 34 | 100% | 0 | 0% | 1 | 34 | 100% | 21484 | 21517 - | 21484 |
| 53_ATCC14919.crt.5 | CP014139.1 Streptococcus pyogenes strain MEW123, complete genome | 34 | 34850 | 63.9 | 4.00E-12 | 34 | 100% | 0 | 0% | 1 | 34 | 100% | 15233 | 15266 + | 15233 |
| 53_ATCC14919.crt.5 | CP010450.1 Streptococcus pyogenes strain NGAS638, complete genome | 34 | 35488 | 63.9 | 4.00E-12 | 34 | 100% | 0 | 0% | 1 | 34 | 100% | 29071 | 29104 - | 29071 |
| 53_ATCC14919.crt.5 | AE014074.1 Streptococcus pyogenes MGAS315, complete genome | 34 | 36875 | 63.9 | 4.00E-12 | 34 | 100% | 0 | 0% | 1 | 34 | 100% | 23487 | 23520 - | 23487 |
| 53_ATCC14919.crt.5 | CP011068.1 Streptococcus pyogenes strain STAB10015, complete genome | 34 | 35292 | 63.9 | 4.00E-12 | 34 | 100% | 0 | 0% | 1 | 34 | 100% | 29395 | 29428 - | 29395 |
| 53_ATCC14919.crt.5 | CP011535.2 Streptococcus pyogenes strain M28PF1, complete genome | 34 | 35292 | 63.9 | 4.00E-12 | 34 | 100% | 0 | 0% | 1 | 34 | 100% | 5865 | 5898 + | 5865 |
| 53_ATCC14919.crt.5 | CP011415.1 Streptococcus pyogenes strain D471, complete genome | 34 | 30199 | 63.9 | 4.00E-12 | 34 | 100% | 0 | 0% | 1 | 34 | 100% | 22064 | 22097 - | 22064 |
| 53_ATCC14919.crt.5 | CP011414.1 Streptococcus pyogenes JRS4, complete genome | 34 | 30199 | 63.9 | 4.00E-12 | 34 | 100% | 0 | 0% | 1 | 34 | 100% | 22064 | 22097 - | 22064 |
| 53_ATCC14919.crt.5 | CP007537.1 Streptococcus pyogenes strain AP1, complete genome | 34 | 29027 | 63.9 | 4.00E-12 | 34 | 100% | 0 | 0% | 1 | 34 | 100% | 7511 | 7544 + | 7511 |
| 53_ATCC14919.crt.5 | AP012335.1 Streptococcus pyogenes JRS4 DNA, complete genome | 34 | 30198 | 63.9 | 4.00E-12 | 34 | 100% | 0 | 0% | 1 | 34 | 100% | 22064 | 22097 - | 22064 |
| 53_ATCC14919.crt.5 | AE009949.1 Streptococcus pyogenes MGAS8232, complete genome | 34 | 30227 | 63.9 | 4.00E-12 | 34 | 100% | 0 | 0% | 1 | 34 | 100% | 22060 | 22093 - | 22060 |
| 53_ATCC14919.crt.5 | CP007041.1 Streptococcus pyogenes STAB902, complete genome | 34 | 36899 | 63.9 | 4.00E-12 | 34 | 100% | 0 | 0% | 1 | 34 | 100% | 13366 | 13399 + | 13366 |
| 53_ATCC14919.crt.5 | CP006366.1 Streptococcus pyogenes HSC5, complete genome | 34 | 37520 | 63.9 | 4.00E-12 | 34 | 100% | 0 | 0% | 1 | 34 | 100% | 31114 | 31147 - | 31114 |
| 53_ATCC14919.crt.5 | CP003068.1 Streptococcus pyogenes Alab49, complete genome | 34 | 29027 | 63.9 | 4.00E-12 | 34 | 100% | 0 | 0% | 1 | 34 | 100% | 21484 | 21517 - | 21484 |
| 53_ATCC14919.crt.5 | AM295007.1 Streptococcus pyogenes Manfredo complete genome | 34 | 28976 | 63.9 | 4.00E-12 | 34 | 100% | 0 | 0% | 1 | 34 | 100% | 7465 | 7498 + | 7465 |
| 53_ATCC14919.crt.6 | CP010449.1 Streptococcus pyogenes strain NGAS322, complete genome | 34 | 38059 | 63.9 | 4.00E-12 | 34 | 100% | 0 | 0% | 1 | 34 | 100% | 36361 | 36394 + | 36361 |
| 53_ATCC14919.crt.6 | CP007560.1 Streptococcus pyogenes strain NGAS743, complete genome | 34 | 33384 | 63.9 | 4.00E-12 | 34 | 100% | 0 | 0% | 1 | 34 | 100% | 1666 | 1699 - | 1666 |
| 53_ATCC14919.crt.6 | AE009949.1 Streptococcus pyogenes MGAS8232, complete genome | 34 | 39804 | 63.9 | 4.00E-12 | 34 | 100% | 0 | 0% | 1 | 34 | 100% | 1666 | 1699 - | 1666 |
| 53_ATCC14919.crt.6 | AM295007.1 Streptococcus pyogenes Manfredo complete genome | 34 | 39447 | 63.9 | 4.00E-12 | 34 | 100% | 0 | 0% | 1 | 34 | 100% | 37749 | 37782 + | 37749 |
| 53_ATCC14919.crt.7 | NOHIT | 34 | | | | | | | | | | | | | |
| 53_ATCC14919.crt.8 | CP000017.2 Streptococcus pyogenes MGAS5005, complete genome | 35 | 29805 | 65.8 | 1.00E-12 | 35 | 100% | 0 | 0% | 1 | 35 | 100% | 28036 | 28070 - | 28036 |
| 53_ATCC14919.crt.8 | CP012045.1 Streptococcus pyogenes strain HKU488, complete genome | 35 | 29804 | 65.8 | 1.00E-12 | 35 | 100% | 0 | 0% | 1 | 35 | 100% | 28035 | 28069 - | 28035 |
| 53_ATCC14919.crt.8 | CP008776.1 Streptococcus pyogenes strain 5448, complete genome | 35 | 29805 | 65.8 | 1.00E-12 | 35 | 100% | 0 | 0% | 1 | 35 | 100% | 1736 | 1770 + | 1736 |
| 53_ATCC14919.crt.8 | AP014585.1 Streptococcus pyogenes DNA, complete genome, strain: MTB314 | 35 | 18078 | 65.8 | 1.00E-12 | 35 | 100% | 0 | 0% | 1 | 35 | 100% | 16309 | 16343 - | 16309 |
| 53_ATCC14919.crt.8 | CP006366.1 Streptococcus pyogenes HSC5, complete genome | 35 | 32692 | 65.8 | 1.00E-12 | 35 | 100% | 0 | 0% | 1 | 35 | 100% | 30923 | 30957 - | 30923 |
| 53_ATCC14919.crt.8 | CP003901.1 Streptococcus pyogenes A20, complete genome | 35 | 29805 | 65.8 | 1.00E-12 | 35 | 100% | 0 | 0% | 1 | 35 | 100% | 28036 | 28070 - | 28036 |
| 53_ATCC14919.crt.8 | AP012491.2 Streptococcus pyogenes M1 476 DNA, complete genome | 35 | 29805 | 65.8 | 1.00E-12 | 35 | 100% | 0 | 0% | 1 | 35 | 100% | 28036 | 28070 - | 28036 |
| 53_ATCC14919.crt.9 | A07_1 | 35 | 31011 | 65.8 | 1.00E-12 | 35 | 100% | 0 | 0% | 1 | 35 | 100% | 8161 | 8195 + | 8161 |
| 53_ATCC14919.crt.9 | BA000034.2 Streptococcus pyogenes SSI-1 DNA, complete genome | 35 | 31011 | 65.8 | 1.00E-12 | 35 | 100% | 0 | 0% | 1 | 35 | 100% | 22817 | 22851 - | 22817 |
| 53_ATCC14919.crt.9 | CP008926.1 Streptococcus pyogenes strain ATCC 19615, complete genome | 35 | 31521 | 65.8 | 1.00E-12 | 35 | 100% | 0 | 0% | 1 | 35 | 100% | 22817 | 22851 - | 22817 |
| 53_ATCC14919.crt.10 | A07_1 | 33 | 31011 | 62.1 | 1.00E-11 | 33 | 100% | 0 | 0% | 1 | 33 | 100% | 12261 | 12293 + | 12261 |
| 53_ATCC14919.crt.10 | BA000034.2 Streptococcus pyogenes SSI-1 DNA, complete genome | 33 | 31011 | 62.1 | 1.00E-11 | 33 | 100% | 0 | 0% | 1 | 33 | 100% | 18719 | 18751 - | 18719 |
| 53_ATCC14919.crt.10 | CP008926.1 Streptococcus pyogenes strain ATCC 19615, complete genome | 33 | 31521 | 62.1 | 1.00E-11 | 33 | 100% | 0 | 0% | 1 | 33 | 100% | 18719 | 18751 - | 18719 |
| 54_MGAS9429_2.crt.1 | NOHIT | 32 | | | | | | | | | | | | | |
| 54_MGAS9429_2.crt.2 | CP000056.1 Streptococcus pyogenes MGAS6180, complete genome | 34 | 35292 | 63.9 | 4.00E-12 | 34 | 100% | 0 | 0% | 1 | 34 | 100% | 29395 | 29428 - | 29395 |
| 54_MGAS9429_2.crt.2 | A09_1 | 34 | 30199 | 63.9 | 4.00E-12 | 34 | 100% | 0 | 0% | 1 | 34 | 100% | 8103 | 8136 + | 8103 |
| 54_MGAS9429_2.crt.2 | A07_1 | 34 | 37110 | 63.9 | 4.00E-12 | 34 | 100% | 0 | 0% | 1 | 34 | 100% | 23711 | 23744 - | 23711 |
| 54_MGAS9429_2.crt.2 | A06_1 | 34 | 30200 | 63.9 | 4.00E-12 | 34 | 100% | 0 | 0% | 1 | 34 | 100% | 8103 | 8136 + | 8103 |
| 54_MGAS9429_2.crt.2 | CP011069.1 Streptococcus pyogenes strain STAB09014, complete genome | 34 | 35292 | 63.9 | 4.00E-12 | 34 | 100% | 0 | 0% | 1 | 34 | 100% | 29395 | 29428 - | 29395 |
| 54_MGAS9429_2.crt.2 | BA000034.2 Streptococcus pyogenes SSI-1 DNA, complete genome | 34 | 37117 | 63.9 | 4.00E-12 | 34 | 100% | 0 | 0% | 1 | 34 | 100% | 13367 | 13400 + | 13367 |
| 54_MGAS9429_2.crt.2 | CP013672.1 Streptococcus pyogenes strain AP53, complete genome | 34 | 29027 | 63.9 | 4.00E-12 | 34 | 100% | 0 | 0% | 1 | 34 | 100% | 21484 | 21517 - | 21484 |
| 54_MGAS9429_2.crt.2 | CP014139.1 Streptococcus pyogenes strain MEW123, complete genome | 34 | 34850 | 63.9 | 4.00E-12 | 34 | 100% | 0 | 0% | 1 | 34 | 100% | 15233 | 15266 + | 15233 |
| 54_MGAS9429_2.crt.2 | CP010450.1 Streptococcus pyogenes strain NGAS638, complete genome | 34 | 35488 | 63.9 | 4.00E-12 | 34 | 100% | 0 | 0% | 1 | 34 | 100% | 29071 | 29104 - | 29071 |
| 54_MGAS9429_2.crt.2 | AE014074.1 Streptococcus pyogenes MGAS315, complete genome | 34 | 36875 | 63.9 | 4.00E-12 | 34 | 100% | 0 | 0% | 1 | 34 | 100% | 23487 | 23520 - | 23487 |
| 54_MGAS9429_2.crt.2 | CP011068.1 Streptococcus pyogenes strain STAB10015, complete genome | 34 | 35292 | 63.9 | 4.00E-12 | 34 | 100% | 0 | 0% | 1 | 34 | 100% | 29395 | 29428 - | 29395 |
| 54_MGAS9429_2.crt.2 | CP011535.2 Streptococcus pyogenes strain M28PF1, complete genome | 34 | 35292 | 63.9 | 4.00E-12 | 34 | 100% | 0 | 0% | 1 | 34 | 100% | 5865 | 5898 + | 5865 |
| 54_MGAS9429_2.crt.2 | CP011415.1 Streptococcus pyogenes strain D471, complete genome | 34 | 30199 | 63.9 | 4.00E-12 | 34 | 100% | 0 | 0% | 1 | 34 | 100% | 22064 | 22097 - | 22064 |
| 54_MGAS9429_2.crt.2 | CP011414.1 Streptococcus pyogenes JRS4, complete genome | 34 | 30199 | 63.9 | 4.00E-12 | 34 | 100% | 0 | 0% | 1 | 34 | 100% | 22064 | 22097 - | 22064 |
| 54_MGAS9429_2.crt.2 | CP007537.1 Streptococcus pyogenes strain AP1, complete genome | 34 | 29027 | 63.9 | 4.00E-12 | 34 | 100% | 0 | 0% | 1 | 34 | 100% | 7511 | 7544 + | 7511 |
| 54_MGAS9429_2.crt.2 | AP012335.1 Streptococcus pyogenes JRS4 DNA, complete genome | 34 | 30198 | 63.9 | 4.00E-12 | 34 | 100% | 0 | 0% | 1 | 34 | 100% | 22064 | 22097 - | 22064 |
| 54_MGAS9429_2.crt.2 | AE009949.1 Streptococcus pyogenes MGAS8232, complete genome | 34 | 30227 | 63.9 | 4.00E-12 | 34 | 100% | 0 | 0% | 1 | 34 | 100% | 22060 | 22093 - | 22060 |
| 54_MGAS9429_2.crt.2 | CP007041.1 Streptococcus pyogenes STAB902, complete genome | 34 | 36899 | 63.9 | 4.00E-12 | 34 | 100% | 0 | 0% | 1 | 34 | 100% | 13366 | 13399 + | 13366 |
| 54_MGAS9429_2.crt.2 | CP006366.1 Streptococcus pyogenes HSC5, complete genome | 34 | 37520 | 63.9 | 4.00E-12 | 34 | 100% | 0 | 0% | 1 | 34 | 100% | 31114 | 31147 - | 31114 |
| 54_MGAS9429_2.crt.2 | CP003068.1 Streptococcus pyogenes Alab49, complete genome | 34 | 29027 | 63.9 | 4.00E-12 | 34 | 100% | 0 | 0% | 1 | 34 | 100% | 21484 | 21517 - | 21484 |
| 54_MGAS9429_2.crt.2 | AM295007.1 Streptococcus pyogenes Manfredo complete genome | 34 | 28976 | 63.9 | 4.00E-12 | 34 | 100% | 0 | 0% | 1 | 34 | 100% | 7465 | 7498 + | 7465 |
| 54_MGAS9429_2.crt.3 | A07_1 | 34 | 14000 | 63.9 | 4.00E-12 | 34 | 100% | 0 | 0% | 1 | 34 | 100% | 3933 | 3966 - | 3933 |
| 54_MGAS9429_2.crt.3 | BA000034.2 Streptococcus pyogenes SSI-1 DNA, complete genome | 34 | 14000 | 63.9 | 4.00E-12 | 34 | 100% | 0 | 0% | 1 | 34 | 100% | 10035 | 10068 + | 10035 |
| 54_MGAS9429_2.crt.3 | CP010450.1 Streptococcus pyogenes strain NGAS638, complete genome | 34 | 35488 | 63.9 | 4.00E-12 | 34 | 100% | 0 | 0% | 1 | 34 | 100% | 27731 | 27764 - | 27731 |
| 54_MGAS9429_2.crt.3 | AE014074.1 Streptococcus pyogenes MGAS315, complete genome | 34 | 14000 | 63.9 | 4.00E-12 | 34 | 100% | 0 | 0% | 1 | 34 | 100% | 3933 | 3966 - | 3933 |
| 54_MGAS9429_2.crt.3 | CP008695.1 Streptococcus pyogenes strain M23ND, complete genome | 34 | 37458 | 63.9 | 4.00E-12 | 34 | 100% | 0 | 0% | 1 | 34 | 100% | 13553 | 13586 + | 13553 |
| 54_MGAS9429_2.crt.3 | CP007041.1 Streptococcus pyogenes STAB902, complete genome | 34 | 14001 | 63.9 | 4.00E-12 | 34 | 100% | 0 | 0% | 1 | 34 | 100% | 10036 | 10069 + | 10036 |
| 54_MGAS9429_2.crt.3 | CP003068.1 Streptococcus pyogenes Alab49, complete genome | 34 | 35543 | 63.9 | 4.00E-12 | 34 | 100% | 0 | 0% | 1 | 34 | 100% | 30353 | 30386 - | 30353 |
| 54_MGAS9429_2.crt.4 | CP010449.1 Streptococcus pyogenes strain NGAS322, complete genome | 34 | 38059 | 63.9 | 4.00E-12 | 34 | 100% | 0 | 0% | 1 | 34 | 100% | 36361 | 36394 + | 36361 |
| 54_MGAS9429_2.crt.4 | CP007560.1 Streptococcus pyogenes strain NGAS743, complete genome | 34 | 33384 | 63.9 | 4.00E-12 | 34 | 100% | 0 | 0% | 1 | 34 | 100% | 1666 | 1699 - | 1666 |
| 54_MGAS9429_2.crt.4 | AE009949.1 Streptococcus pyogenes MGAS8232, complete genome | 34 | 39804 | 63.9 | 4.00E-12 | 34 | 100% | 0 | 0% | 1 | 34 | 100% | 1666 | 1699 - | 1666 |
| 54_MGAS9429_2.crt.4 | AM295007.1 Streptococcus pyogenes Manfredo complete genome | 34 | 39447 | 63.9 | 4.00E-12 | 34 | 100% | 0 | 0% | 1 | 34 | 100% | 37749 | 37782 + | 37749 |
| 54_MGAS9429_2.crt.5 | NOHIT | 34 | | | | | | | | | | | | | |
| 54_MGAS9429_2.crt.6 | CP000017.2 Streptococcus pyogenes MGAS5005, complete genome | 35 | 29805 | 65.8 | 1.00E-12 | 35 | 100% | 0 | 0% | 1 | 35 | 100% | 28036 | 28070 - | 28036 |
| 54_MGAS9429_2.crt.6 | CP012045.1 Streptococcus pyogenes strain HKU488, complete genome | 35 | 29804 | 65.8 | 1.00E-12 | 35 | 100% | 0 | 0% | 1 | 35 | 100% | 28035 | 28069 - | 28035 |
| 54_MGAS9429_2.crt.6 | CP008776.1 Streptococcus pyogenes strain 5448, complete genome | 35 | 29805 | 65.8 | 1.00E-12 | 35 | 100% | 0 | 0% | 1 | 35 | 100% | 1736 | 1770 + | 1736 |
| 54_MGAS9429_2.crt.6 | AP014585.1 Streptococcus pyogenes DNA, complete genome, strain: MTB314 | 35 | 18078 | 65.8 | 1.00E-12 | 35 | 100% | 0 | 0% | 1 | 35 | 100% | 16309 | 16343 - | 16309 |
| 54_MGAS9429_2.crt.6 | CP006366.1 Streptococcus pyogenes HSC5, complete genome | 35 | 32692 | 65.8 | 1.00E-12 | 35 | 100% | 0 | 0% | 1 | 35 | 100% | 30923 | 30957 - | 30923 |
| 54_MGAS9429_2.crt.6 | CP003901.1 Streptococcus pyogenes A20, complete genome | 35 | 29805 | 65.8 | 1.00E-12 | 35 | 100% | 0 | 0% | 1 | 35 | 100% | 28036 | 28070 - | 28036 |
| 54_MGAS9429_2.crt.6 | AP012491.2 Streptococcus pyogenes M1 476 DNA, complete genome | 35 | 29805 | 65.8 | 1.00E-12 | 35 | 100% | 0 | 0% | 1 | 35 | 100% | 28036 | 28070 - | 28036 |
| 54_MGAS9429_2.crt.7 | A07_1 | 35 | 31011 | 65.8 | 1.00E-12 | 35 | 100% | 0 | 0% | 1 | 35 | 100% | 8161 | 8195 + | 8161 |
| 54_MGAS9429_2.crt.7 | BA000034.2 Streptococcus pyogenes SSI-1 DNA, complete genome | 35 | 31011 | 65.8 | 1.00E-12 | 35 | 100% | 0 | 0% | 1 | 35 | 100% | 22817 | 22851 - | 22817 |
| 54_MGAS9429_2.crt.7 | CP008926.1 Streptococcus pyogenes strain ATCC 19615, complete genome | 35 | 31521 | 65.8 | 1.00E-12 | 35 | 100% | 0 | 0% | 1 | 35 | 100% | 22817 | 22851 - | 22817 |
| 54_MGAS9429_2.crt.8 | A07_1 | 33 | 31011 | 62.1 | 1.00E-11 | 33 | 100% | 0 | 0% | 1 | 33 | 100% | 12261 | 12293 + | 12261 |
| 54_MGAS9429_2.crt.8 | BA000034.2 Streptococcus pyogenes SSI-1 DNA, complete genome | 33 | 31011 | 62.1 | 1.00E-11 | 33 | 100% | 0 | 0% | 1 | 33 | 100% | 18719 | 18751 - | 18719 |
| 54_MGAS9429_2.crt.8 | CP008926.1 Streptococcus pyogenes strain ATCC 19615, complete genome | 33 | 31521 | 62.1 | 1.00E-11 | 33 | 100% | 0 | 0% | 1 | 33 | 100% | 18719 | 18751 - | 18719 |
| 55_MGAS2096_2.crt.1 | A07_1 | 34 | 31011 | 63.9 | 4.00E-12 | 34 | 100% | 0 | 0% | 1 | 34 | 100% | 12261 | 12294 - | 12261 |
| 55_MGAS2096_2.crt.1 | BA000034.2 Streptococcus pyogenes SSI-1 DNA, complete genome | 34 | 31011 | 63.9 | 4.00E-12 | 34 | 100% | 0 | 0% | 1 | 34 | 100% | 18718 | 18751 + | 18718 |
| 55_MGAS2096_2.crt.1 | CP008926.1 Streptococcus pyogenes strain ATCC 19615, complete genome | 34 | 31521 | 63.9 | 4.00E-12 | 34 | 100% | 0 | 0% | 1 | 34 | 100% | 18718 | 18751 + | 18718 |
| 55_MGAS2096_2.crt.2 | A07_1 | 36 | 31011 | 67.6 | 3.00E-13 | 36 | 100% | 0 | 0% | 1 | 36 | 100% | 8161 | 8196 - | 8161 |
| 55_MGAS2096_2.crt.2 | BA000034.2 Streptococcus pyogenes SSI-1 DNA, complete genome | 36 | 31011 | 67.6 | 3.00E-13 | 36 | 100% | 0 | 0% | 1 | 36 | 100% | 22816 | 22851 + | 22816 |
| 55_MGAS2096_2.crt.2 | CP008926.1 Streptococcus pyogenes strain ATCC 19615, complete genome | 36 | 31521 | 67.6 | 3.00E-13 | 36 | 100% | 0 | 0% | 1 | 36 | 100% | 22816 | 22851 + | 22816 |
| 55_MGAS2096_2.crt.3 | CP008695.1 Streptococcus pyogenes strain M23ND, complete genome | 35 | 37509 | 60.2 | 6.00E-11 | 34 | 97% | 0 | 0% | 1 | 35 | 100% | 36269 | 36303 - | 36269 |
| 55_MGAS2096_2.crt.4 | CP010449.1 Streptococcus pyogenes strain NGAS322, complete genome | 35 | 38059 | 65.8 | 1.00E-12 | 35 | 100% | 0 | 0% | 1 | 35 | 100% | 36361 | 36395 - | 36361 |
| 55_MGAS2096_2.crt.4 | CP007560.1 Streptococcus pyogenes strain NGAS743, complete genome | 35 | 33384 | 65.8 | 1.00E-12 | 35 | 100% | 0 | 0% | 1 | 35 | 100% | 1665 | 1699 + | 1665 |
| 55_MGAS2096_2.crt.4 | AE009949.1 Streptococcus pyogenes MGAS8232, complete genome | 35 | 39804 | 65.8 | 1.00E-12 | 35 | 100% | 0 | 0% | 1 | 35 | 100% | 1665 | 1699 + | 1665 |
| 55_MGAS2096_2.crt.4 | AM295007.1 Streptococcus pyogenes Manfredo complete genome | 35 | 39447 | 65.8 | 1.00E-12 | 35 | 100% | 0 | 0% | 1 | 35 | 100% | 37749 | 37783 - | 37749 |
| 55_MGAS2096_2.crt.4 | A07_1 | 35 | 37110 | 60.2 | 6.00E-11 | 34 | 97% | 0 | 0% | 1 | 35 | 100% | 35742 | 35776 - | 35742 |
| 55_MGAS2096_2.crt.4 | BA000034.2 Streptococcus pyogenes SSI-1 DNA, complete genome | 35 | 37117 | 60.2 | 6.00E-11 | 34 | 97% | 0 | 0% | 1 | 35 | 100% | 1335 | 1369 + | 1335 |
| 55_MGAS2096_2.crt.4 | AE014074.1 Streptococcus pyogenes MGAS315, complete genome | 35 | 36875 | 60.2 | 6.00E-11 | 34 | 97% | 0 | 0% | 1 | 35 | 100% | 35507 | 35541 - | 35507 |
| 55_MGAS2096_2.crt.4 | CP007041.1 Streptococcus pyogenes STAB902, complete genome | 35 | 36899 | 60.2 | 6.00E-11 | 34 | 97% | 0 | 0% | 1 | 35 | 100% | 1335 | 1369 + | 1335 |
| 55_MGAS2096_2.crt.5 | CP000056.1 Streptococcus pyogenes MGAS6180, complete genome | 35 | 35292 | 65.8 | 1.00E-12 | 35 | 100% | 0 | 0% | 1 | 35 | 100% | 29394 | 29428 + | 29394 |
| 55_MGAS2096_2.crt.5 | A09_1 | 35 | 30199 | 65.8 | 1.00E-12 | 35 | 100% | 0 | 0% | 1 | 35 | 100% | 8103 | 8137 - | 8103 |
| 55_MGAS2096_2.crt.5 | A07_1 | 35 | 37110 | 65.8 | 1.00E-12 | 35 | 100% | 0 | 0% | 1 | 35 | 100% | 23710 | 23744 + | 23710 |
| 55_MGAS2096_2.crt.5 | A06_1 | 35 | 30200 | 65.8 | 1.00E-12 | 35 | 100% | 0 | 0% | 1 | 35 | 100% | 8103 | 8137 - | 8103 |
| 55_MGAS2096_2.crt.5 | CP011069.1 Streptococcus pyogenes strain STAB09014, complete genome | 35 | 35292 | 65.8 | 1.00E-12 | 35 | 100% | 0 | 0% | 1 | 35 | 100% | 29394 | 29428 + | 29394 |
| 55_MGAS2096_2.crt.5 | BA000034.2 Streptococcus pyogenes SSI-1 DNA, complete genome | 35 | 37117 | 65.8 | 1.00E-12 | 35 | 100% | 0 | 0% | 1 | 35 | 100% | 13367 | 13401 - | 13367 |
| 55_MGAS2096_2.crt.5 | CP013672.1 Streptococcus pyogenes strain AP53, complete genome | 35 | 29027 | 65.8 | 1.00E-12 | 35 | 100% | 0 | 0% | 1 | 35 | 100% | 21483 | 21517 + | 21483 |
| 55_MGAS2096_2.crt.5 | CP014139.1 Streptococcus pyogenes strain MEW123, complete genome | 35 | 34850 | 65.8 | 1.00E-12 | 35 | 100% | 0 | 0% | 1 | 35 | 100% | 15233 | 15267 - | 15233 |
| 55_MGAS2096_2.crt.5 | CP010450.1 Streptococcus pyogenes strain NGAS638, complete genome | 35 | 35488 | 65.8 | 1.00E-12 | 35 | 100% | 0 | 0% | 1 | 35 | 100% | 29070 | 29104 + | 29070 |
| 55_MGAS2096_2.crt.5 | AE014074.1 Streptococcus pyogenes MGAS315, complete genome | 35 | 36875 | 65.8 | 1.00E-12 | 35 | 100% | 0 | 0% | 1 | 35 | 100% | 23486 | 23520 + | 23486 |
| 55_MGAS2096_2.crt.5 | CP011068.1 Streptococcus pyogenes strain STAB10015, complete genome | 35 | 35292 | 65.8 | 1.00E-12 | 35 | 100% | 0 | 0% | 1 | 35 | 100% | 29394 | 29428 + | 29394 |
| 55_MGAS2096_2.crt.5 | CP011535.2 Streptococcus pyogenes strain M28PF1, complete genome | 35 | 35292 | 65.8 | 1.00E-12 | 35 | 100% | 0 | 0% | 1 | 35 | 100% | 5865 | 5899 - | 5865 |
| 55_MGAS2096_2.crt.5 | CP011415.1 Streptococcus pyogenes strain D471, complete genome | 35 | 30199 | 65.8 | 1.00E-12 | 35 | 100% | 0 | 0% | 1 | 35 | 100% | 22063 | 22097 + | 22063 |
| 55_MGAS2096_2.crt.5 | CP011414.1 Streptococcus pyogenes JRS4, complete genome | 35 | 30199 | 65.8 | 1.00E-12 | 35 | 100% | 0 | 0% | 1 | 35 | 100% | 22063 | 22097 + | 22063 |
| 55_MGAS2096_2.crt.5 | CP007537.1 Streptococcus pyogenes strain AP1, complete genome | 35 | 29027 | 65.8 | 1.00E-12 | 35 | 100% | 0 | 0% | 1 | 35 | 100% | 7511 | 7545 - | 7511 |
| 55_MGAS2096_2.crt.5 | AP012335.1 Streptococcus pyogenes JRS4 DNA, complete genome | 35 | 30198 | 65.8 | 1.00E-12 | 35 | 100% | 0 | 0% | 1 | 35 | 100% | 22063 | 22097 + | 22063 |
| 55_MGAS2096_2.crt.5 | AE009949.1 Streptococcus pyogenes MGAS8232, complete genome | 35 | 30227 | 65.8 | 1.00E-12 | 35 | 100% | 0 | 0% | 1 | 35 | 100% | 22059 | 22093 + | 22059 |
| 55_MGAS2096_2.crt.5 | CP007041.1 Streptococcus pyogenes STAB902, complete genome | 35 | 36899 | 65.8 | 1.00E-12 | 35 | 100% | 0 | 0% | 1 | 35 | 100% | 13366 | 13400 - | 13366 |
| 55_MGAS2096_2.crt.5 | CP006366.1 Streptococcus pyogenes HSC5, complete genome | 35 | 37520 | 65.8 | 1.00E-12 | 35 | 100% | 0 | 0% | 1 | 35 | 100% | 31113 | 31147 + | 31113 |
| 55_MGAS2096_2.crt.5 | CP003068.1 Streptococcus pyogenes Alab49, complete genome | 35 | 29027 | 65.8 | 1.00E-12 | 35 | 100% | 0 | 0% | 1 | 35 | 100% | 21483 | 21517 + | 21483 |
| 55_MGAS2096_2.crt.5 | AM295007.1 Streptococcus pyogenes Manfredo complete genome | 35 | 28976 | 65.8 | 1.00E-12 | 35 | 100% | 0 | 0% | 1 | 35 | 100% | 7465 | 7499 - | 7465 |
| 55_MGAS2096_2.crt.6 | NOHIT | 33 | | | | | | | | | | | | | |
| 55_MGAS2096_2.crt.7 | NOHIT | 0 | | | | | | | | | | | | | |
| 55_MGAS2096_2.crt.8 | NOHIT | 37 | | | | | | | | | | | | | |
| 55_MGAS2096_2.crt.9 | NOHIT | 38 | | | | | | | | | | | | | |
| 58_KIH06_201.crt.1 | NOHIT | 34 | | | | | | | | | | | | | |
| 58_KIH06_201.crt.2 | NOHIT | 34 | | | | | | | | | | | | | |
| 58_KIH06_201.crt.3 | AE004092.2 Streptococcus pyogenes M1 GAS, complete genome | 33 | 29920 | 62.1 | 1.00E-11 | 33 | 100% | 0 | 0% | 1 | 33 | 100% | 22970 | 23002 + | 22970 |
| 58_KIH06_201.crt.3 | CP006366.1 Streptococcus pyogenes HSC5, complete genome | 33 | 39477 | 62.1 | 1.00E-11 | 33 | 100% | 0 | 0% | 1 | 33 | 100% | 27280 | 27312 + | 27280 |
| 6_MGAS6180.crt.1 | NOHIT | 30 | | | | | | | | | | | | | |
| 6_MGAS6180.crt.2 | NOHIT | 30 | | | | | | | | | | | | | |
| 6_MGAS6180.crt.3 | NOHIT | 30 | | | | | | | | | | | | | |
| 6_MGAS6180.crt.4 | NOHIT | 30 | | | | | | | | | | | | | |
| 7_MGAS5005.crt.1 | NOHIT | 30 | | | | | | | | | | | | | |
| 7_MGAS5005.crt.2 | NOHIT | 30 | | | | | | | | | | | | | |
| 7_MGAS5005.crt.3 | NOHIT | 30 | | | | | | | | | | | | | |
| 7_MGAS5005.crt.4 | NOHIT | 0 | | | | | | | | | | | | | |
| 7_MGAS5005.crt.5 | CP007561.1 Streptococcus pyogenes strain NGAS596, complete genome | 33 | 40047 | 62.1 | 1.00E-11 | 33 | 100% | 0 | 0% | 1 | 33 | 100% | 4156 | 4188 - | 4156 |
| 7_MGAS5005.crt.5 | CP008695.1 Streptococcus pyogenes strain M23ND, complete genome | 33 | 37509 | 62.1 | 1.00E-11 | 33 | 100% | 0 | 0% | 1 | 33 | 100% | 34682 | 34714 + | 34682 |
| 7_MGAS5005.crt.5 | AM295007.1 Streptococcus pyogenes Manfredo complete genome | 33 | 37666 | 62.1 | 1.00E-11 | 33 | 100% | 0 | 0% | 1 | 33 | 100% | 4183 | 4215 - | 4183 |
| 7_MGAS5005.crt.6 | NOHIT | 34 | | | | | | | | | | | | | |
| 7_MGAS5005.crt.7 | CP008695.1 Streptococcus pyogenes strain M23ND, complete genome | 33 | 37458 | 62.1 | 1.00E-11 | 33 | 100% | 0 | 0% | 1 | 33 | 100% | 21052 | 21084 + | 21052 |
| 7_MGAS5005.crt.8 | A08_1 | 35 | 36269 | 60.2 | 6.00E-11 | 34 | 97% | 0 | 0% | 1 | 35 | 100% | 28369 | 28403 - | 28369 |
| 7_MGAS5005.crt.8 | CP013838.1 Streptococcus pyogenes strain MGAS11027, complete genome | 35 | 51647 | 60.2 | 6.00E-11 | 34 | 97% | 0 | 0% | 1 | 35 | 100% | 13944 | 13978 + | 13944 |
| 7_MGAS5005.crt.8 | CP007560.1 Streptococcus pyogenes strain NGAS743, complete genome | 35 | 38050 | 60.2 | 6.00E-11 | 34 | 97% | 0 | 0% | 1 | 35 | 100% | 27864 | 27898 - | 27864 |
| 7_MGAS5005.crt.8 | CP007560.1 Streptococcus pyogenes strain NGAS743, complete genome | 35 | 39059 | 60.2 | 6.00E-11 | 34 | 97% | 0 | 0% | 1 | 35 | 100% | 14542 | 14576 + | 14542 |
| 7_MGAS5005.crt.8 | AE009949.1 Streptococcus pyogenes MGAS8232, complete genome | 35 | 39804 | 60.2 | 6.00E-11 | 34 | 97% | 0 | 0% | 1 | 35 | 100% | 12058 | 12092 + | 12058 |
| 7_MGAS5005.crt.8 | CP009612.1 Streptococcus pyogenes strain HKU360, complete genome | 35 | 38759 | 60.2 | 6.00E-11 | 34 | 97% | 0 | 0% | 1 | 35 | 100% | 26510 | 26544 - | 26510 |
| 7_MGAS5005.crt.8 | CP008695.1 Streptococcus pyogenes strain M23ND, complete genome | 35 | 37458 | 60.2 | 6.00E-11 | 34 | 97% | 0 | 0% | 1 | 35 | 100% | 12800 | 12834 + | 12800 |
| 7_MGAS5005.crt.8 | CP006366.1 Streptococcus pyogenes HSC5, complete genome | 35 | 39477 | 60.2 | 6.00E-11 | 34 | 97% | 0 | 0% | 1 | 35 | 100% | 24623 | 24657 - | 24623 |
| 7_MGAS5005.crt.8 | AM295007.1 Streptococcus pyogenes Manfredo complete genome | 35 | 39447 | 60.2 | 6.00E-11 | 34 | 97% | 0 | 0% | 1 | 35 | 100% | 27341 | 27375 - | 27341 |
| 8_MGAS9429.crt.1 | NOHIT | 38 | | | | | | | | | | | | | |
| 8_MGAS9429.crt.2 | NOHIT | 37 | | | | | | | | | | | | | |
| 8_MGAS9429.crt.3 | NOHIT | 0 | | | | | | | | | | | | | |
| 8_MGAS9429.crt.4 | NOHIT | 32 | | | | | | | | | | | | | |
| 8_MGAS9429.crt.5 | CP000056.1 Streptococcus pyogenes MGAS6180, complete genome | 34 | 35292 | 63.9 | 4.00E-12 | 34 | 100% | 0 | 0% | 1 | 34 | 100% | 29395 | 29428 - | 29395 |
| 8_MGAS9429.crt.5 | A09_1 | 34 | 30199 | 63.9 | 4.00E-12 | 34 | 100% | 0 | 0% | 1 | 34 | 100% | 8103 | 8136 + | 8103 |
| 8_MGAS9429.crt.5 | A07_1 | 34 | 37110 | 63.9 | 4.00E-12 | 34 | 100% | 0 | 0% | 1 | 34 | 100% | 23711 | 23744 - | 23711 |
| 8_MGAS9429.crt.5 | A06_1 | 34 | 30200 | 63.9 | 4.00E-12 | 34 | 100% | 0 | 0% | 1 | 34 | 100% | 8103 | 8136 + | 8103 |
| 8_MGAS9429.crt.5 | CP011069.1 Streptococcus pyogenes strain STAB09014, complete genome | 34 | 35292 | 63.9 | 4.00E-12 | 34 | 100% | 0 | 0% | 1 | 34 | 100% | 29395 | 29428 - | 29395 |
| 8_MGAS9429.crt.5 | BA000034.2 Streptococcus pyogenes SSI-1 DNA, complete genome | 34 | 37117 | 63.9 | 4.00E-12 | 34 | 100% | 0 | 0% | 1 | 34 | 100% | 13367 | 13400 + | 13367 |
| 8_MGAS9429.crt.5 | CP013672.1 Streptococcus pyogenes strain AP53, complete genome | 34 | 29027 | 63.9 | 4.00E-12 | 34 | 100% | 0 | 0% | 1 | 34 | 100% | 21484 | 21517 - | 21484 |
| 8_MGAS9429.crt.5 | CP014139.1 Streptococcus pyogenes strain MEW123, complete genome | 34 | 34850 | 63.9 | 4.00E-12 | 34 | 100% | 0 | 0% | 1 | 34 | 100% | 15233 | 15266 + | 15233 |
| 8_MGAS9429.crt.5 | CP010450.1 Streptococcus pyogenes strain NGAS638, complete genome | 34 | 35488 | 63.9 | 4.00E-12 | 34 | 100% | 0 | 0% | 1 | 34 | 100% | 29071 | 29104 - | 29071 |
| 8_MGAS9429.crt.5 | AE014074.1 Streptococcus pyogenes MGAS315, complete genome | 34 | 36875 | 63.9 | 4.00E-12 | 34 | 100% | 0 | 0% | 1 | 34 | 100% | 23487 | 23520 - | 23487 |
| 8_MGAS9429.crt.5 | CP011068.1 Streptococcus pyogenes strain STAB10015, complete genome | 34 | 35292 | 63.9 | 4.00E-12 | 34 | 100% | 0 | 0% | 1 | 34 | 100% | 29395 | 29428 - | 29395 |
| 8_MGAS9429.crt.5 | CP011535.2 Streptococcus pyogenes strain M28PF1, complete genome | 34 | 35292 | 63.9 | 4.00E-12 | 34 | 100% | 0 | 0% | 1 | 34 | 100% | 5865 | 5898 + | 5865 |
| 8_MGAS9429.crt.5 | CP011415.1 Streptococcus pyogenes strain D471, complete genome | 34 | 30199 | 63.9 | 4.00E-12 | 34 | 100% | 0 | 0% | 1 | 34 | 100% | 22064 | 22097 - | 22064 |
| 8_MGAS9429.crt.5 | CP011414.1 Streptococcus pyogenes JRS4, complete genome | 34 | 30199 | 63.9 | 4.00E-12 | 34 | 100% | 0 | 0% | 1 | 34 | 100% | 22064 | 22097 - | 22064 |
| 8_MGAS9429.crt.5 | CP007537.1 Streptococcus pyogenes strain AP1, complete genome | 34 | 29027 | 63.9 | 4.00E-12 | 34 | 100% | 0 | 0% | 1 | 34 | 100% | 7511 | 7544 + | 7511 |
| 8_MGAS9429.crt.5 | AP012335.1 Streptococcus pyogenes JRS4 DNA, complete genome | 34 | 30198 | 63.9 | 4.00E-12 | 34 | 100% | 0 | 0% | 1 | 34 | 100% | 22064 | 22097 - | 22064 |
| 8_MGAS9429.crt.5 | AE009949.1 Streptococcus pyogenes MGAS8232, complete genome | 34 | 30227 | 63.9 | 4.00E-12 | 34 | 100% | 0 | 0% | 1 | 34 | 100% | 22060 | 22093 - | 22060 |
| 8_MGAS9429.crt.5 | CP007041.1 Streptococcus pyogenes STAB902, complete genome | 34 | 36899 | 63.9 | 4.00E-12 | 34 | 100% | 0 | 0% | 1 | 34 | 100% | 13366 | 13399 + | 13366 |
| 8_MGAS9429.crt.5 | CP006366.1 Streptococcus pyogenes HSC5, complete genome | 34 | 37520 | 63.9 | 4.00E-12 | 34 | 100% | 0 | 0% | 1 | 34 | 100% | 31114 | 31147 - | 31114 |
| 8_MGAS9429.crt.5 | CP003068.1 Streptococcus pyogenes Alab49, complete genome | 34 | 29027 | 63.9 | 4.00E-12 | 34 | 100% | 0 | 0% | 1 | 34 | 100% | 21484 | 21517 - | 21484 |
| 8_MGAS9429.crt.5 | AM295007.1 Streptococcus pyogenes Manfredo complete genome | 34 | 28976 | 63.9 | 4.00E-12 | 34 | 100% | 0 | 0% | 1 | 34 | 100% | 7465 | 7498 + | 7465 |
| 8_MGAS9429.crt.6 | CP010449.1 Streptococcus pyogenes strain NGAS322, complete genome | 34 | 38059 | 63.9 | 4.00E-12 | 34 | 100% | 0 | 0% | 1 | 34 | 100% | 36361 | 36394 + | 36361 |
| 8_MGAS9429.crt.6 | CP007560.1 Streptococcus pyogenes strain NGAS743, complete genome | 34 | 33384 | 63.9 | 4.00E-12 | 34 | 100% | 0 | 0% | 1 | 34 | 100% | 1666 | 1699 - | 1666 |
| 8_MGAS9429.crt.6 | AE009949.1 Streptococcus pyogenes MGAS8232, complete genome | 34 | 39804 | 63.9 | 4.00E-12 | 34 | 100% | 0 | 0% | 1 | 34 | 100% | 1666 | 1699 - | 1666 |
| 8_MGAS9429.crt.6 | AM295007.1 Streptococcus pyogenes Manfredo complete genome | 34 | 39447 | 63.9 | 4.00E-12 | 34 | 100% | 0 | 0% | 1 | 34 | 100% | 37749 | 37782 + | 37749 |
| 8_MGAS9429.crt.7 | NOHIT | 34 | | | | | | | | | | | | | |
| 8_MGAS9429.crt.8 | CP000017.2 Streptococcus pyogenes MGAS5005, complete genome | 35 | 29805 | 65.8 | 1.00E-12 | 35 | 100% | 0 | 0% | 1 | 35 | 100% | 28036 | 28070 - | 28036 |
| 8_MGAS9429.crt.8 | CP012045.1 Streptococcus pyogenes strain HKU488, complete genome | 35 | 29804 | 65.8 | 1.00E-12 | 35 | 100% | 0 | 0% | 1 | 35 | 100% | 28035 | 28069 - | 28035 |
| 8_MGAS9429.crt.8 | CP008776.1 Streptococcus pyogenes strain 5448, complete genome | 35 | 29805 | 65.8 | 1.00E-12 | 35 | 100% | 0 | 0% | 1 | 35 | 100% | 1736 | 1770 + | 1736 |
| 8_MGAS9429.crt.8 | AP014585.1 Streptococcus pyogenes DNA, complete genome, strain: MTB314 | 35 | 18078 | 65.8 | 1.00E-12 | 35 | 100% | 0 | 0% | 1 | 35 | 100% | 16309 | 16343 - | 16309 |
| 8_MGAS9429.crt.8 | CP006366.1 Streptococcus pyogenes HSC5, complete genome | 35 | 32692 | 65.8 | 1.00E-12 | 35 | 100% | 0 | 0% | 1 | 35 | 100% | 30923 | 30957 - | 30923 |
| 8_MGAS9429.crt.8 | CP003901.1 Streptococcus pyogenes A20, complete genome | 35 | 29805 | 65.8 | 1.00E-12 | 35 | 100% | 0 | 0% | 1 | 35 | 100% | 28036 | 28070 - | 28036 |
| 8_MGAS9429.crt.8 | AP012491.2 Streptococcus pyogenes M1 476 DNA, complete genome | 35 | 29805 | 65.8 | 1.00E-12 | 35 | 100% | 0 | 0% | 1 | 35 | 100% | 28036 | 28070 - | 28036 |
| 8_MGAS9429.crt.9 | A07_1 | 35 | 31011 | 65.8 | 1.00E-12 | 35 | 100% | 0 | 0% | 1 | 35 | 100% | 8161 | 8195 + | 8161 |
| 8_MGAS9429.crt.9 | BA000034.2 Streptococcus pyogenes SSI-1 DNA, complete genome | 35 | 31011 | 65.8 | 1.00E-12 | 35 | 100% | 0 | 0% | 1 | 35 | 100% | 22817 | 22851 - | 22817 |
| 8_MGAS9429.crt.9 | CP008926.1 Streptococcus pyogenes strain ATCC 19615, complete genome | 35 | 31521 | 65.8 | 1.00E-12 | 35 | 100% | 0 | 0% | 1 | 35 | 100% | 22817 | 22851 - | 22817 |
| 8_MGAS9429.crt.10 | A07_1 | 33 | 31011 | 62.1 | 1.00E-11 | 33 | 100% | 0 | 0% | 1 | 33 | 100% | 12261 | 12293 + | 12261 |
| 8_MGAS9429.crt.10 | BA000034.2 Streptococcus pyogenes SSI-1 DNA, complete genome | 33 | 31011 | 62.1 | 1.00E-11 | 33 | 100% | 0 | 0% | 1 | 33 | 100% | 18719 | 18751 - | 18719 |
| 8_MGAS9429.crt.10 | CP008926.1 Streptococcus pyogenes strain ATCC 19615, complete genome | 33 | 31521 | 62.1 | 1.00E-11 | 33 | 100% | 0 | 0% | 1 | 33 | 100% | 18719 | 18751 - | 18719 |
| 9_MGAS10270.crt.1 | NOHIT | 37 | | | | | | | | | | | | | |
| 9_MGAS10270.crt.2 | NOHIT | 37 | | | | | | | | | | | | | |
| 9_MGAS10270.crt.3 | NOHIT | 0 | | | | | | | | | | | | | |
| 9_MGAS10270.crt.4 | A08_1 | 34 | 31416 | 63.9 | 4.00E-12 | 34 | 100% | 0 | 0% | 1 | 34 | 100% | 31149 | 31182 + | 31149 |
| 9_MGAS10270.crt.4 | A07_1 | 34 | 37110 | 63.9 | 4.00E-12 | 34 | 100% | 0 | 0% | 1 | 34 | 100% | 235 | 268 - | 235 |
| 9_MGAS10270.crt.4 | A07_1 | 34 | 31011 | 63.9 | 4.00E-12 | 34 | 100% | 0 | 0% | 1 | 34 | 100% | 30744 | 30777 + | 30744 |
| 9_MGAS10270.crt.4 | BA000034.2 Streptococcus pyogenes SSI-1 DNA, complete genome | 34 | 31011 | 63.9 | 4.00E-12 | 34 | 100% | 0 | 0% | 1 | 34 | 100% | 235 | 268 - | 235 |
| 9_MGAS10270.crt.4 | CP013838.1 Streptococcus pyogenes strain MGAS11027, complete genome | 34 | 35828 | 63.9 | 4.00E-12 | 34 | 100% | 0 | 0% | 1 | 34 | 100% | 35561 | 35594 + | 35561 |
| 9_MGAS10270.crt.4 | CP013672.1 Streptococcus pyogenes strain AP53, complete genome | 34 | 24636 | 63.9 | 4.00E-12 | 34 | 100% | 0 | 0% | 1 | 34 | 100% | 2347 | 2380 - | 2347 |
| 9_MGAS10270.crt.4 | CP014138.1 Streptococcus pyogenes strain MEW427, complete genome | 34 | 22381 | 63.9 | 4.00E-12 | 34 | 100% | 0 | 0% | 1 | 34 | 100% | 235 | 268 - | 235 |
| 9_MGAS10270.crt.4 | CP010449.1 Streptococcus pyogenes strain NGAS322, complete genome | 34 | 38059 | 63.9 | 4.00E-12 | 34 | 100% | 0 | 0% | 1 | 34 | 100% | 2334 | 2367 - | 2334 |
| 9_MGAS10270.crt.4 | CP007560.1 Streptococcus pyogenes strain NGAS743, complete genome | 34 | 39618 | 63.9 | 4.00E-12 | 34 | 100% | 0 | 0% | 1 | 34 | 100% | 235 | 268 - | 235 |
| 9_MGAS10270.crt.4 | CP007560.1 Streptococcus pyogenes strain NGAS743, complete genome | 34 | 33384 | 63.9 | 4.00E-12 | 34 | 100% | 0 | 0% | 1 | 34 | 100% | 33117 | 33150 + | 33117 |
| 9_MGAS10270.crt.4 | CP011415.1 Streptococcus pyogenes strain D471, complete genome | 34 | 39110 | 63.9 | 4.00E-12 | 34 | 100% | 0 | 0% | 1 | 34 | 100% | 235 | 268 - | 235 |
| 9_MGAS10270.crt.4 | CP011414.1 Streptococcus pyogenes JRS4, complete genome | 34 | 39110 | 63.9 | 4.00E-12 | 34 | 100% | 0 | 0% | 1 | 34 | 100% | 235 | 268 - | 235 |
| 9_MGAS10270.crt.4 | AP012335.1 Streptococcus pyogenes JRS4 DNA, complete genome | 34 | 39111 | 63.9 | 4.00E-12 | 34 | 100% | 0 | 0% | 1 | 34 | 100% | 235 | 268 - | 235 |
| 9_MGAS10270.crt.4 | CP009612.1 Streptococcus pyogenes strain HKU360, complete genome | 34 | 38759 | 63.9 | 4.00E-12 | 34 | 100% | 0 | 0% | 1 | 34 | 100% | 235 | 268 - | 235 |
| 9_MGAS10270.crt.4 | CP008695.1 Streptococcus pyogenes strain M23ND, complete genome | 34 | 37509 | 63.9 | 4.00E-12 | 34 | 100% | 0 | 0% | 1 | 34 | 100% | 37242 | 37275 + | 37242 |
| 9_MGAS10270.crt.4 | CP008926.1 Streptococcus pyogenes strain ATCC 19615, complete genome | 34 | 31521 | 63.9 | 4.00E-12 | 34 | 100% | 0 | 0% | 1 | 34 | 100% | 235 | 268 - | 235 |
| 9_MGAS10270.crt.4 | CP006366.1 Streptococcus pyogenes HSC5, complete genome | 34 | 39477 | 63.9 | 4.00E-12 | 34 | 100% | 0 | 0% | 1 | 34 | 100% | 235 | 268 - | 235 |
| 9_MGAS10270.crt.4 | CP003068.1 Streptococcus pyogenes Alab49, complete genome | 34 | 24636 | 63.9 | 4.00E-12 | 34 | 100% | 0 | 0% | 1 | 34 | 100% | 2347 | 2380 - | 2347 |
| 9_MGAS10270.crt.4 | CP000262.1 Streptococcus pyogenes MGAS10750, complete genome | 34 | 31381 | 63.9 | 4.00E-12 | 34 | 100% | 0 | 0% | 1 | 34 | 100% | 235 | 268 - | 235 |
| 9_MGAS10270.crt.5 | A07_1 | 34 | 37110 | 63.9 | 4.00E-12 | 34 | 100% | 0 | 0% | 1 | 34 | 100% | 35198 | 35231 - | 35198 |
| 9_MGAS10270.crt.5 | BA000034.2 Streptococcus pyogenes SSI-1 DNA, complete genome | 34 | 37117 | 63.9 | 4.00E-12 | 34 | 100% | 0 | 0% | 1 | 34 | 100% | 1880 | 1913 + | 1880 |
| 9_MGAS10270.crt.5 | CP010449.1 Streptococcus pyogenes strain NGAS322, complete genome | 34 | 38059 | 63.9 | 4.00E-12 | 34 | 100% | 0 | 0% | 1 | 34 | 100% | 35817 | 35850 - | 35817 |
| 9_MGAS10270.crt.5 | AE014074.1 Streptococcus pyogenes MGAS315, complete genome | 34 | 36875 | 63.9 | 4.00E-12 | 34 | 100% | 0 | 0% | 1 | 34 | 100% | 34963 | 34996 - | 34963 |
| 9_MGAS10270.crt.5 | CP007560.1 Streptococcus pyogenes strain NGAS743, complete genome | 34 | 33384 | 63.9 | 4.00E-12 | 34 | 100% | 0 | 0% | 1 | 34 | 100% | 2210 | 2243 + | 2210 |
| 9_MGAS10270.crt.5 | AE009949.1 Streptococcus pyogenes MGAS8232, complete genome | 34 | 39804 | 63.9 | 4.00E-12 | 34 | 100% | 0 | 0% | 1 | 34 | 100% | 2210 | 2243 + | 2210 |
| 9_MGAS10270.crt.5 | CP007041.1 Streptococcus pyogenes STAB902, complete genome | 34 | 36899 | 63.9 | 4.00E-12 | 34 | 100% | 0 | 0% | 1 | 34 | 100% | 1880 | 1913 + | 1880 |
| 9_MGAS10270.crt.5 | AM295007.1 Streptococcus pyogenes Manfredo complete genome | 34 | 39447 | 63.9 | 4.00E-12 | 34 | 100% | 0 | 0% | 1 | 34 | 100% | 37205 | 37238 - | 37205 |
| 9_MGAS10270.crt.6 | CP008695.1 Streptococcus pyogenes strain M23ND, complete genome | 32 | 27969 | 60.2 | 5.00E-11 | 32 | 100% | 0 | 0% | 1 | 32 | 100% | 10571 | 10602 - | 10571 |
